# Genomic analyses of high-grade neuroendocrine gynecological malignancies reveal a unique mutational landscape and therapeutic vulnerabilities

**DOI:** 10.1101/2020.11.17.387514

**Authors:** Haider Mahdi, Amy Joehlin-Price, Afshin Dowlati, Ata Abbas

**Affiliations:** Gynecologic Oncology Division, Cleveland Clinic, Cleveland, OH 44195; Department of Pathology, Cleveland Clinic, Cleveland, OH 44195; Division of Hematology and Oncology, Department of Medicine, Case Western Reserve University, Cleveland, OH 44106; University Hospitals Seidman Cancer Center, Cleveland, OH 44106; Developmental Therapeutics Program, Case Comprehensive Cancer Center, Case Western Reserve University School of Medicine, Cleveland, OH 44116

**Keywords:** Gynecologic neuroendocrine carcinoma, PARP inhibitors, CDK4/6 inhibitors, immunotherapy targets, YAP1

## Abstract

The high-grade neuroendocrine carcinoma of gynecologic origin (NEC-GYN) is a highly aggressive cancer affecting young women. The clinical management of NEC-GYN is often extrapolated from their counterpart, small cell carcinoma of the lung (SCLC), but, unfortunately, they have limited effect. In our NEC-GYN cohort, median progression-free survival (PFS) and overall survival (OS) were 1 and 12 months, respectively, indicating their highly lethal nature. Our comprehensive genomic analyses unveiled that NEC-GYN harbors a higher mutational burden with distinct mutational landscapes from SCLC. We identified 14 cancer driver genes (FDR <0.01) including the most frequently altered *KMT2C* (100%), *KNL1* (100%), *NCOR2* (100%), and *CCDC6* (93%) genes. Transcriptomic analyses identified several novel gene fusions in NEC-GYN. Furthermore, NEC-GYN exhibited a highly immunosuppressive state and uniquely belonged to the *YAP1* high molecular subtype that promotes multidrug resistance. Our study suggests an urgent need to reevaluate the therapeutic options and targets for NEC-GYN.

## Introduction

High-grade NECs of the cervix are rare, aggressive cancers accounting for about 1-1.5% of all cervical cancer^1,2^. Unlike the common type, squamous cell carcinoma, patients with cervical NEC are more likely to present with advanced or metastatic disease, resulting in poor prognosis. The 5-year survival is up to 36% for early-stage disease; however, the advanced-stage disease has <10% survival, with relapse rates exceeding 90%^1–4^. These cancers are likely to have a vascular invasion and nodal or visceral metastasis. Unfortunately, NEC of the cervix affects young women with a median age of 37^1–4^. High-grade NECs of other gynecologic origins are even rarer, share similar aggressive behavior and poor outcomes^4^. Furthermore, no prospective data are available to guide therapy in NEC-GYN. Therefore, an urgency exists in understanding the underlying pathobiology of NEC-GYN with hopes of developing better therapeutic options.

Current treatment considerations and guidelines for NEC-GYN are mostly extrapolated from studies conducted in SCLC. In patients with advanced-stage metastatic disease, standard therapy includes chemotherapy with platinum and etoposide^3–5^. In patients with recurrent disease, data to guide treatment decisions are entirely absent but again inferred from SCLC, using agents such as topotecan and paclitaxel. However, these regimens are both toxic and have limited activity. Most recently, our group and others have described subgroups of SCLC that may have different pathobiology and potentially distinct therapeutic vulnerabilities^6,7^. Therefore, we sought to comprehensively investigate NEC-GYN genomics to understand their oncologic drivers and determine their similarities and differences with SCLC to improve their clinical management.

## Results

### Patient characteristics, treatments and sequencing

Between 1998 and 2018, twenty-seven patients were diagnosed with high-grade NEC-GYN at Cleveland Clinic. Fifteen FFPE samples from 12 patients, including three samples at two different time points (see Table 1 for demographic and clinicopathologic annotations), were included. The majority of patients had advanced stage IIIC-IV (75%). Sites of origin included the cervix (58%), ovary (25%), and endometrium (17%). 58% of the patients received Cisplatin/Etoposide at the time diagnosis, and second line treatments at recurrence/progression were Topotecan (23%), liposomal Doxorubicin +/− Bevacizumab (23%), Immunotherapy (Nivolumab +/−Ipilimumab) (15.4%), etc. (Table 1). The median number of prior lines of therapy was 1. Median PFS and OS were 1 and 12 months, respectively (Table 1). Whole exome sequencing (WES) of 14 samples and stranded paired-end RNA-sequencing of 13 samples, including 12 samples with matched WES, were performed. The mean coverage for WES was 280.7× (range, 240.8–330×) and the average number of stranded paired-end reads for RNA-seq was 35.7 million (range, 23.1–46.3 million).

**Table 1.**
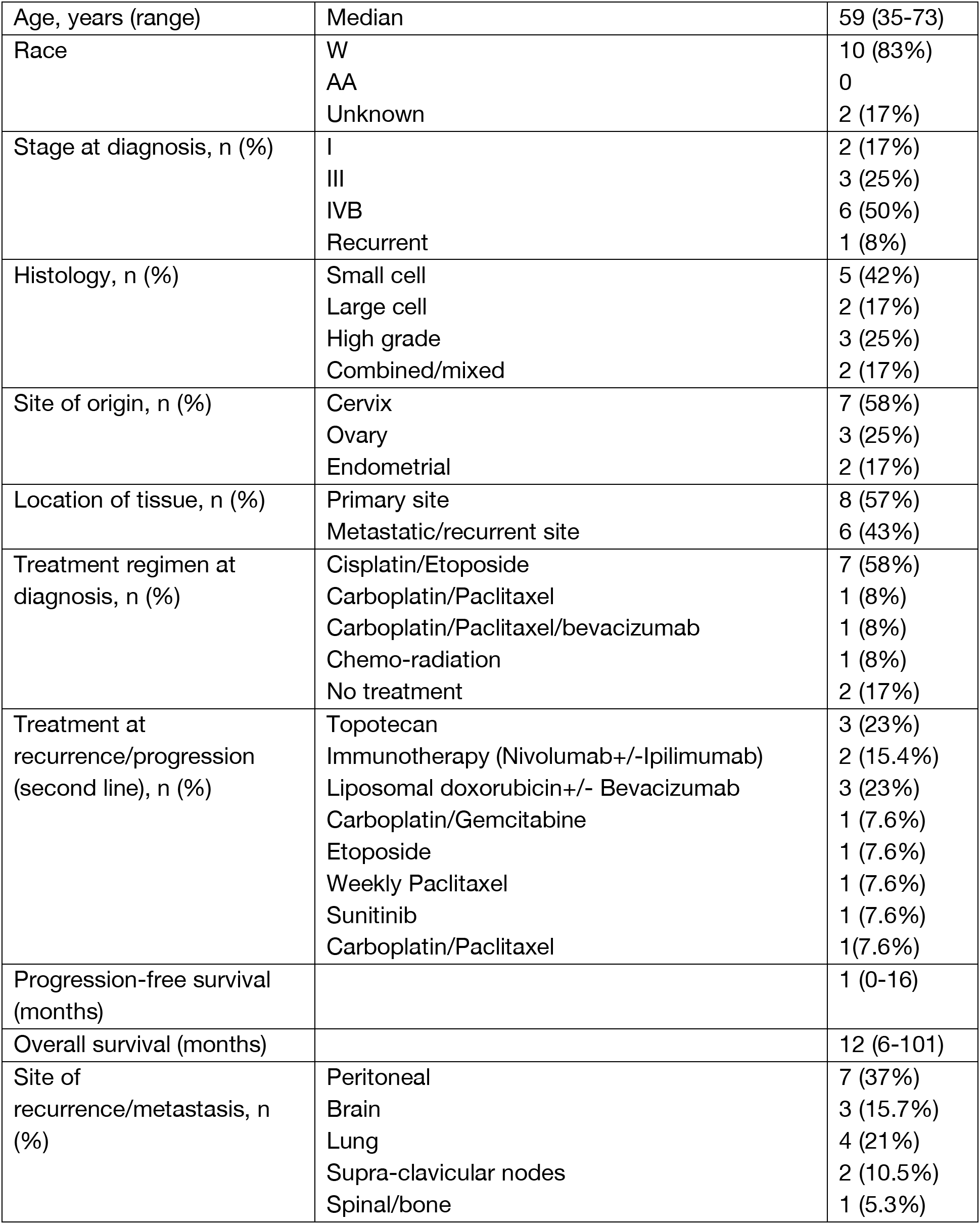
Patients’ clinical and pathologic characteristics, and treatment

### Mutational profile and tumor mutation burden in NEC-GYN

We discovered a unique mutational landscape in our NEC-GYN cohort (Figure 1, Supplementary Figure 1-3, Supplementary Table 1). The WES analyses detected mutations in *KMT2C*, *KNL1*, and *NCOR2* genes in all the tumor samples (100%) (Figure 1A, Supplementary Figure 4-6). Cervical NEC samples had a significantly higher number of mutations within individual genes compared with other NEC-GYN tumors (Supplementary Figure 7). The altered genes that were common between tumors from three gynecologic sites, surprisingly, enriched in the Hippo signaling pathway (Supplementary Figure 8). Using the oncodriveCLUST algorithm, we identified 14 cancer driver genes at a false discovery rate (FDR) of 0.01 (Figure1B, Supplementary Table 2). The oncodriveCLUST algorithm measures genes’ bias towards large mutation clustering and identify specific hot-spots where most of the variants in cancer-causing genes are enriched. The frequent driver genes were *CCDC6* (13 mutations in one cluster, 93% mutation frequency), *LATS2* (15 mutations in 2 clusters, 86%), and *CLTCL1* and *RNF43* (18 and 17 mutations respectively in 3 clusters, 86%) (Supplementary Table 2). Oncogenic pathway analysis revealed that the highest proportion of mutated genes were found in the cell cycle (73.3%), TGF-β (71.4%), RTK-RAS (44.7%), MYC (38.5), and PI3K (37.9%) pathways (Figure 1C, Supplementary Figure 9-10). Next, we compared the tumor mutation burden (TMB) of our cohort with 33 TCGA cohorts of various tumors. Surprisingly, NEC-GYN demonstrated the highest TMB compared to all TCGA cohorts examined (Figure 1D). Furthermore, we observed frequent mutations in various DNA-damage response (DDR) pathway genes including *MSH3* (79%), *FANCD2* (71%), *BRCA2* (64%), *BLM* (50%), etc. (Figure 1E), corroborating high TMB in NEC-GYN. Moreover, drug-gene interactions analysis using GDIdb identified clinically actionable (*KMT2C*, *MAP3K1*) and other potentially druggable targets (Supplementary Figure 11).

**Figure 1:**
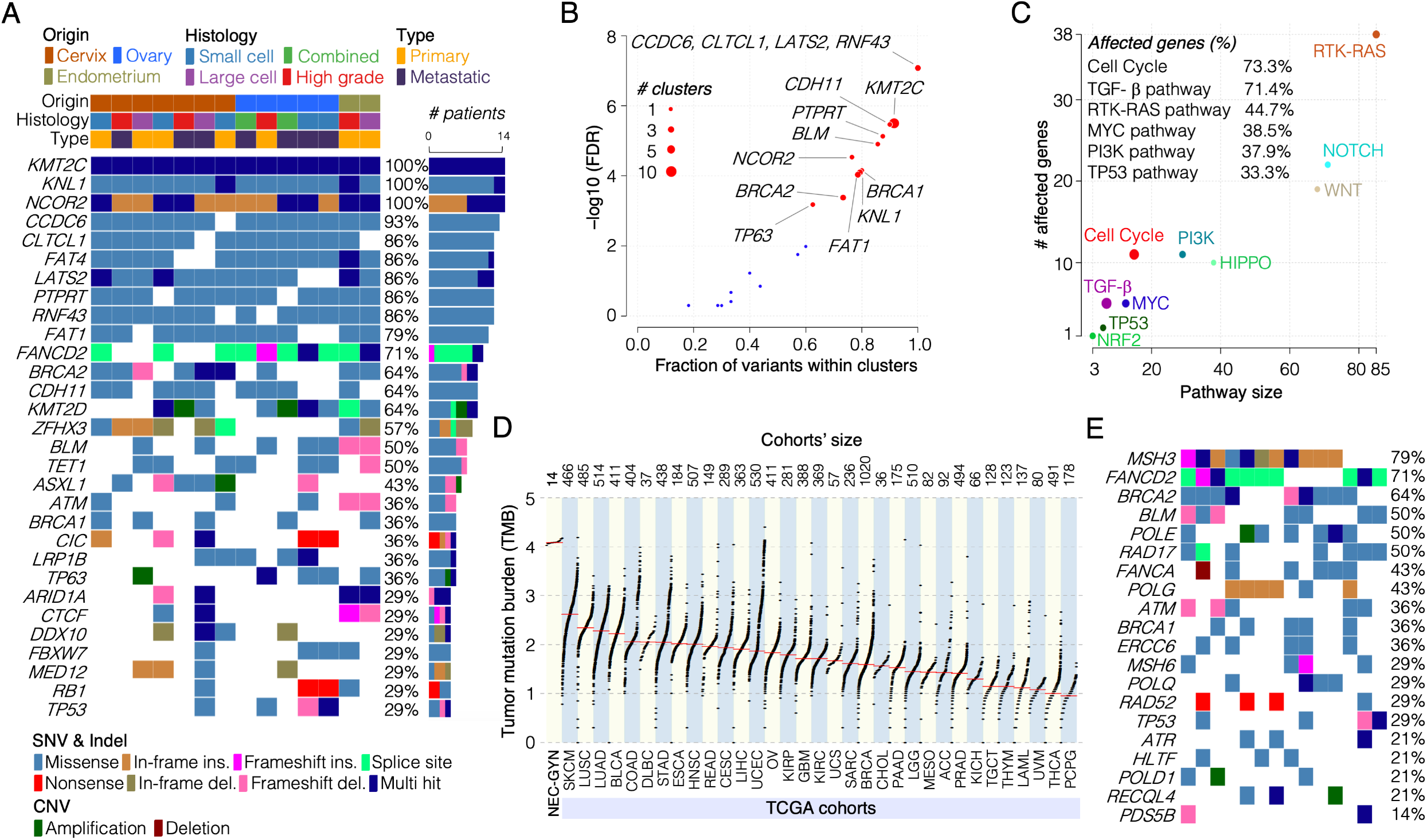
Unique mutational landscape in NEC-GYN. **(A)** Frequently altered genes in NEC of gynecologic origins detected by GATK HaplotypeCaller and XHMM. Cohort’s clinical features are given on the top and color codes representing various SNVs, Indel, and CNVs are given at the bottom **(B)** A scatter plot showing cancer driver genes (FDR <0.01) detected by OncodriveCLUST algorithm using positional clustering method. **(C)** Oncogenic signaling pathways associated with mutated genes in NEC-GYN are illustrated in a scatter plot. Percentage of genes altered in various oncogenic pathways are given. **(D)** Comparison of TMB in NEC-GYN with TCGA representing 33 tumors’ cohorts. The cohorts’ size is given on the top. **(E)** Mutation profile of DNA-damage repair genes in NEC-GYN cohort detected by GATK HaplotypeCaller and XHMM. Please refer to Figure 1A for color codes representing mutation types.

### Recurrent gene fusions, unique transcriptomic signatures and immunosuppressive tumor-microenvironment in NEC-GYN

Using transcriptomic data, we discovered several recurrent gene fusions, including *MALAT1-SMG* (53.8%), *EEF1A1-MALAT1* (30.8%), and *ASH1L-YY1AP1* (30.8%) (Figure 2A, Supplementary Table 3). Remarkably, the *MALAT1* gene partnered in >20% of all the fusion events (Figure 2A) and demonstrated extremely high expression in NEC-GYN compared to healthy cervix and ovary (Figure 2B, Supplementary Figure 12). Comparing transcriptomic profiles of NEC-GYN with TCGA cervical squamous cell and epithelial ovarian cancer, the non-neuroendocrine counterparts of the NEC-GYN, chromatin assembly and nucleosome organization were the top GO functions for significantly over-expressed genes in NEC-GYN (Supplementary Figure 13-14). Remarkably, under-expressed genes in NEC-GYN were enriched for protein modification and catabolic processes, and neutrophil-mediated immunity (Supplementary Figure 13-14).

**Figure 2:**
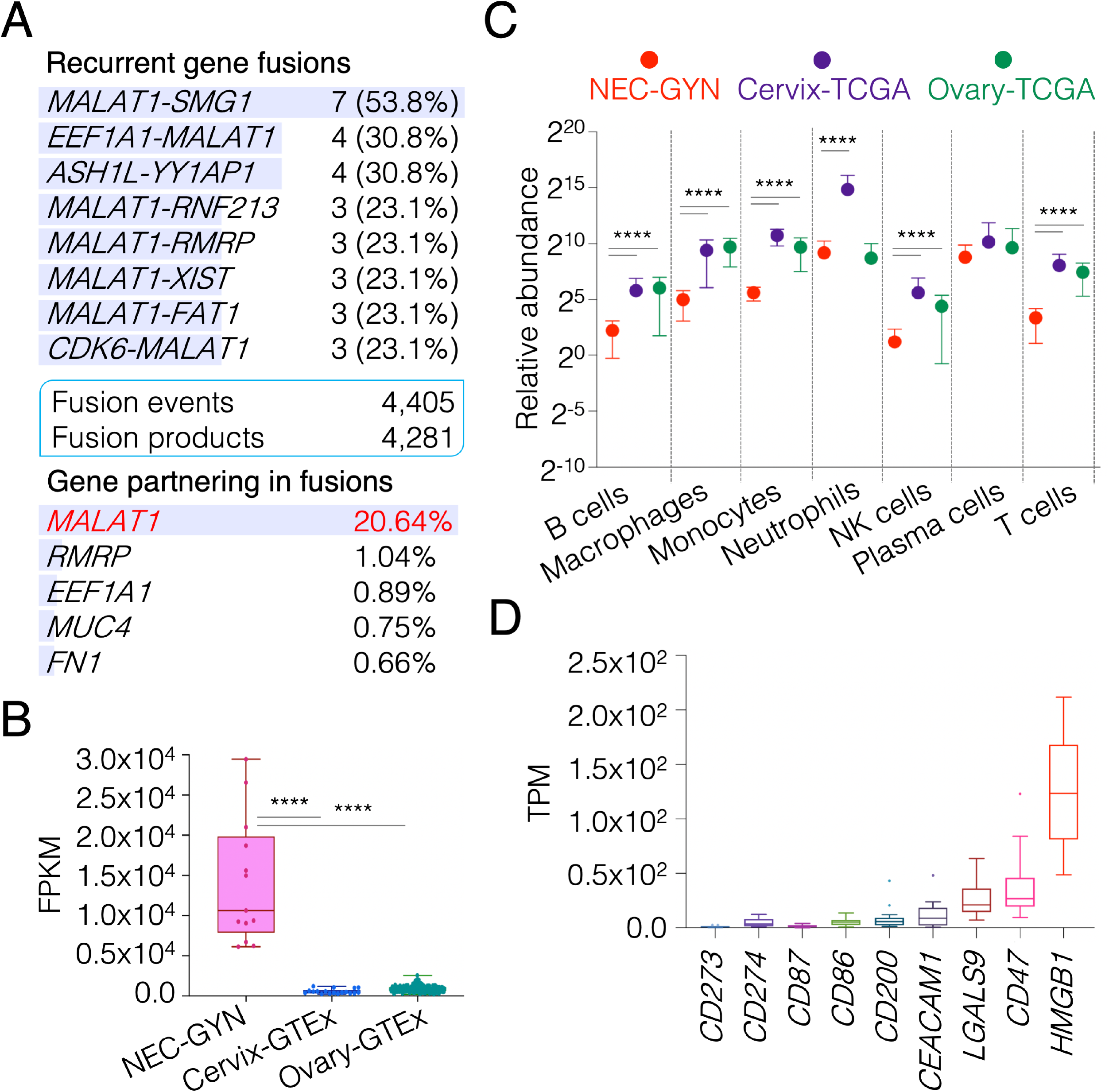
Transcriptional signature and tumor immune-microenvironment in NEC-GYN. **(A)** Recurrent gene fusions detected by using FusionCatcher in NEC-GYN. A total of 4405 fusion events were observed across NEC-GYN cohort that resulted in 4281 unique fusion products (see Supplementary table 3 for full list). Genes partnering in total fusion events (%) are given. **(B)** Expression profile of *MALAT1* gene in NEC-GYN, normal cervix, and normal ovary tissues (mean with SD, \*\*\*\**P* ≤ 0.0001 by two-tailed Mann–Whitney U test). **(C)** Comparison of relative abundance of immune cells in various tumors based on network-based deconvolution (ImSig) analysis (mean with SD, \*\**P* = 0.01, \*\*\*\**P* ≤ 0.0001 by two-tailed Mann–Whitney U test). **(D)** Relative expression levels of various immune checkpoint genes in NEC-GYN (box plot by Tukey method).

The tumor immune-microenvironment is crucial in determining immunotherapy outcomes. Using transcriptomic data, we analyzed immune cell gene signatures for profiling tumor microenvironment. Our data revealed significantly lower immune cell infiltrations in NEC-GYN compared to the ovarian or cervical non-endocrine cohorts, except plasma cells (Figure 2C). We then looked at the expression of various immune checkpoint genes and observed a high expression of *HMGB1*, followed by *CD47* and *LGALS9*, and low expression of *CD274* (PD-L1) and *CD273* (PD-L2) (Figure 2D).

### NEC-GYN exhibits distinct mutational profile from SCLC and belongs to *YAP1* high molecular subtype

Since NEC-GYN tumors are traditionally considered similar to SCLC, the prototypical high-grade neuroendocrine cancer, we compared our results with SCLC genomic mutation data obtained from cBioPortal, including data from a cohort of 90 SCLC patients from our laboratory. Astonishingly, the NEC-GYN mutation profile did not match SCLC as the most frequently mutated genes in NEC-GYN were among the least mutated in SCLC (Figure 3A). In SCLC, *TP53* and *RB1* genes are frequently mutated (88.9% and 75.7% respectively); however, *TP53* and *RB1* mutations in our NEC-GYN cohort were significantly lower (29%) (Figure 3A).

**Figure 3:**
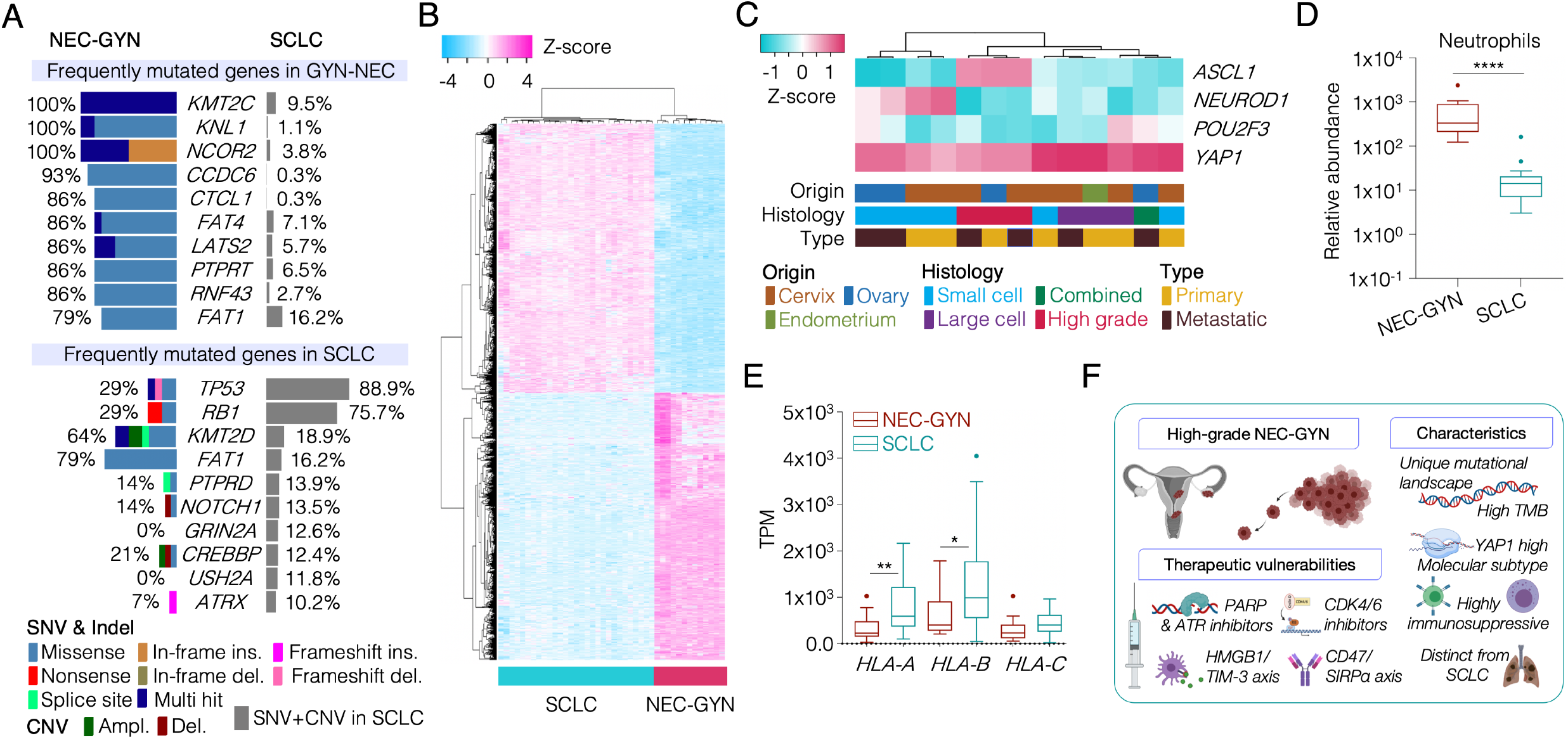
Comprehensive genomic and transcriptomic analyses between NEC-GYN and SCLC. **(A)** Comparison of frequently mutated genes (top 10 genes from both NEC-GYN and SCLC) between NEC-GYN and SCLC cohorts. **(B)** A heatmap representing distinct transcriptional signatures between NEC-GYN and SCLC cohorts. DEGs were sorted by *P*-adj values, and top 5000 gene were used for unsupervised hierarchical clustering. **(C)** Molecular subtypes of NEC-GYN defined by the expression of four key transcriptional regulators. **(D)** Comparison of relative abundance of neutrophils (ImSig analysis) between NYC-GYN and SCLC (Tukey, \*\*\*\**P* ≤ 0.0001 by two-tailed Mann–Whitney U test). **(E)** Expression patterns of HLA class-I genes in NEC-GYN and SCLC cohorts (Tukey, \**P* = 0.0262, \*\**P* = 0.0028 by two-tailed Mann–Whitney U test). **(F)** A cartoon depicting origin of NEC-GYN, unique characteristics, and therapeutic opportunities.

Compared to SCLC, NEC-GYN showed highly distinct transcriptomic patterns, as represented by the top 5,000 DEGs (p-adj >0.01) (Figure 3B, Supplementary Figure 15-16). SCLC is typically classified into four molecular subtypes, including *ASCL1* (the largest group), *NEUROD1*, *POU2F3*, and a small group represented by *YAP1* ^7^. Surprisingly, all of the NEC-GYN tumors, while still clustering into four groups, represent the *YAP1* high, a chemoresistant molecular subtype (Figure 3C). Looking into immune-microenvironment compared to SCLC, only neutrophils infiltration was significantly higher in NEC-GYN (*P* ≤ 0.0001) (Figure 3D, Supplementary Figure 17); however, *HLA-A* and *HLA-B* expression were significantly lower in our cohort (Figure 4E) demonstrating lower antigen presentation capacity.

## Discussion

Comprehensive genomic studies are limited in NEC-GYN. A pilot study with WES data of 5 neuroendocrine cervix cases reported recurrent mutation in *ATRX*, *EBRR4*, and AKT/mTOR signaling pathway genes^8^. Another study of NEC of the cervix reported mutation in *PIK3CA* (18%), *KRAS* (14%), and *TP53* (11%) genes using a limited gene-targeted panel^9^. A study of 10 cervical NEC cases found frequent mutations in *TP53* (40%) and *PIK3CA* (30%) genes, again using a targeted gene-panel^10^. More recently, Hillman et al. reported recurrent mutations in *PIK3CA* (26.7%) and *KMT2C* (20%) genes along with deletion in regions containing the *PTEN* gene (33%)^11^. Remarkably, no transcriptomic data are available for NEC-GYN. To the best of our knowledge, our study is the first comprehensive study having matched genomic and transcriptomic data that provides a novel opportunity to understand the pathobiology of NEC-GYN.

Our data demonstrated a unique mutational landscape in NEC-GYN with frequent mutations in COMPASS family members (*KMT2C*, 100%; and *KMT2D*, 64%) (Figure 1A) involved in epigenetic regulation of enhancers and their loss promote tumorigenesis^12^. Surprisingly, we observed frequent mutations in various DDR genes (Figure 1E) that possibly explain the incidence of very high TMB in our cohort compared to 33 TCGA cohorts. Providentially, DDR genes mutations confer therapeutic vulnerabilities in cancers^13–15^. A case report demonstrated promising results using PARP inhibitor in a small cell carcinoma of the cervix patient with somatic *BRCA2* gene mutation^16^. PARP inhibitor, rucaparib treatment resulted in 15☐months PFS and symptomatic improvement, was noteworthy for such a disease with very high mortality. A recent study evaluating PARP1 expression using IHC in pathological specimens of high-grade NEC of cervix reported PARP1 positivity for 91% of samples^17^. In our NEC-GYN cohort, the mutation rate in *BRCA2* (64%) and other genes involved in the DDR pathway was remarkably high (Figure 1E). Our data strongly suggests using PARP and other inhibitors targeting the DDR pathway in NEC-GYN. Furthermore, mutations in the cancer driver gene, *CCDC6* (93%), imply the use of PARP inhibitor^18^.

Transcriptional dysregulation is a hallmark of cancer, and oncogenic gene fusions are observed across several cancer types^19^. NEC-GYN transcriptomic signatures revealed modulation in chromatin assembly and nucleosome organization (Supplementary Figure 13-14). These transcriptomic data are consistent with the NEC-GYN mutational profile as *KMT2C*, *KMT2D*, and other epigenetic genes were frequently mutated. Surprisingly, gene fusions in NEC-GYN were mostly partnered with the *MALAT1* gene (Figure 2A). MALAT1 is one of the best-characterized lincRNA and a potentially viable druggable target^20^. Transcriptomic based tumor immune microenvironment analysis allows us to understand the abundance of the immune cells in tumor tissues. The presence of significantly lower immune cell infiltrations in our cohort than their non-neuroendocrine counterparts (Figure 2C) explains its immunosuppressive nature and perhaps the minimal effect of anti-PD1/PD-L1 immunotherapies in NEC-GYN^21^. Remarkably, other novel immunotherapeutic targets such as *HMGB1* and *CD47* expression are far greater compared to the *CD274* gene (codes for PD-L1) (Fig 2.D). HMGB1, Gal-9, CEACAM1, and PS are the four well-known ligands for TIM-3, a negative regulator of Type 1 immunity expressed on various immune cells. CD47 is a ligand for SIRPα receptor present on macrophages and involved in *‘don’t eat me’* signaling. Future clinical studies are needed to evaluate HMGB1/TIM-3 and CD47/SIRPα axes for potential immunotherapy targets, alone or in combination with chemotherapy, in NEC-GYN.

Treatment options for NEC-GYN are mostly inferred from studies conducted in SCLC. Astoundingly, the mutational landscape and transcriptional signatures are quite different in NEC-GYN compared to SCLC (Figure 3). SCLC is characterized by a very high *TP53* and *RB1* mutation rate that is believed to be responsible for the small cell phenotype. Conversely, *TP53* and *RB1* mutations in NEC-GYN were significantly lower (29%) then SCLC and did not distinctly correlate with their histology (Figure 3A, 1A). Unexpectedly, NEC-GYN showed a high expression of *YAP1*, which represents a small molecular subtype in SCLC. YAP1 is a component of the Hippo pathway responsible for multidrug resistance^22^. This unique YAP1 high molecular subtype possibly maintains NYC-GYN’s chemo-refractory nature that further facilitates immunosuppressive tumor microenvironment^23^. Lower antigen presentation capacity (Figure 3 E) also aid in immunosuppressive tumor-microenvironment in NEC-GYN and higher infiltration of neutrophils associated with poor clinical outcome^24^.

In conclusion, our data demonstrate a unique mutational landscape in NEC-GYN with a remarkably high TMB and suggest novel multi-modality therapeutic measures to address this highly fatal cancer (Figure 3F). Frequent mutations of DDR genes apprise PARP and ATR inhibitors-based therapies. Our finding of high *YAP1* expression along with frequent *RB1* wild-type status suggests a potential therapeutic vulnerability to CDK 4/6 inhibitors^6^. Our data provide a knowledge framework for future studies in prospective clinical settings to improve these highly lethal malignancies’ clinical outcomes affecting young women by using targeted therapies.

## Methods

For a detailed methodology, please refer to the *Supplementary methods*.

### Tumor samples, whole-exome and RNA-sequencing, and public database

Twenty-seven patients were diagnosed with high-grade NEC-GYN at Cleveland Clinic (between 1998 and 2018). Of them, we were able to retrieve 16 formalin-fixed paraffin-embedded (FFPE) samples that represented 13 patients, including three samples at two different time points. Response Evaluation Criteria in Solid Tumors (RECIST) 1.1 criteria was used assess objective response rate (ORR). PFS and OS were calculated with KM curves. All the samples were re-evaluated by an expert pathologist who marked the regions to dissect tissues for nucleic acids isolation. We successfully sequence fifteen FFPE samples from 12 patients, including three samples at two different time points. WES of 14 samples and stranded paired-end RNA-seq of 13 samples, including 12 samples with matched WES, were performed at Novogene Corporation Inc. (Sacramento, CA) using 150-bp paired-end format on a NovaSeq 6000 (Illumina, San Diego, CA) sequencer.

Data for frequently altered genes in SCLC were obtained from cBioPortal (https://www.cbioportal.org/ and https://www.cbioportal.org/sclc). Cervical and Ovarian cancer TCGA transcriptomic data were downloaded from the LinkedOmics portal (http://www.linkedomics.org/). Transcriptomic data from healthy human tissues were downloaded from GTEx (https://www.gtexportal.org/).

### WES analyses, variant calling, and CNV analysis

Sequence reads were mapped with GRCh38 using BWA-MEM with default parameters after FastQC and adoptor trimming using Trim Galore. Fallowing GATK best practices, variants were then called using GATK4 HaplotypeCaller with default parameters after removing PCR duplicates (using Picard tools) and subsequent realignment and base recalibration. MAF files were generated using the vcf2maf package. Ensembl Variant Effect Predictor (VEP) was used to filter variants. Non-deleterious missense mutations were further filtered using SIFT and PolyPhen-2. Using XHMM (eXome-Hidden Markov Model) with default parameters, CNVs were called using the hidden Markov model (HMM) Viterbi algorithm, and each called CNV was quantitatively genotyped using HMM forward-backward algorithm. GenomicRanges and Homo.sapiens R packages were used to get genes that fall under the deletion (DEL) and duplication (DUP) regions after converting hg38 coordinates to hg19 using the UCSC LiftOver.

### Mutational signatures, driver genes identification, and TMB comparison

The WES data were visualized using maftools in the R environment to plot MAF summary, transition and transversion mutations, somatic interactions, oncoplots, and lollipop plots for amino acid changes. Oncogenic signaling pathways were detected using the oncogenicPathways function of maftools. Based on oncodriveCLUST algorithm, cancer driver genes were detected based on the positional clustering method using the oncodrive function of maftools. A minimum of 5 mutations per gene cutoff was used, and the p-value was calculated by z-score. Tumor mutation burden was calculated and compared against TCGA cohorts using the tcgaCompare function of maftools. The mutation profile of DNA-damage repair genes was plotted using oncoplots function of maftools.

### RNA-seq analyses

RNA-seq reads were mapped against hg38 using STAR aligner with default parameters after FastQC, and adapter and quality trimming using TrimGalore. DESeq2 analysis with an adjusted P-value <0.001 was used to get a list of differentially expressed genes (DEGs). Top 5,000 significant DEGs (sorted by P-adj values) were used for unsupervised hierarchical clustering (NEC-GYN vs SCLC). Pathway analysis (GO biological process) was performed on Enrichr database. RSEM analyses were performed to calculate FPKM and TPM values with default parameters.

### Gene fusion and immune cell gene signature analyses

Novel and known somatic fusion genes were detected using FusionCatcher using default parameters. The fusion junctions were validated by using four different methods employing Bowtie, BLAT, STAR, and Bowtie2 aligners. Possible false positive and readthrough fusions were excluded.

The ImSig R package was used for analyzing immune cell gene signatures based on network-based deconvolution approach. Using ImSig algorithm with default parameter (correlation threshold, r = 0.7, over 75% genes overlap was observed), a table of the relative abundance of immune cells across samples was generated and plotted for visualizing relative abundance of various immune cells.

### Statistics

Two-tailed Mann–Whitney U tests were used to calculate P values. For box plots, box and whiskers graphs were plotted using either the Tukey method or minimum to maximum values. The middle line in the box indicates the median, whiskers indicate the highest and lowest values within 1.5×IQR (Inter-quartile distance between the 25th and 75th percentiles) up and down from the box, and dots plot values >1.5×IQR up and down from the box (for Figure 2D, Figure 3D-E). In Figure 2B, whiskers were plotted down to the minimum and up to the maximum value and individual value as a point superimposed on the graph. In Figure 2C, mean with SD was plotted.

### Data availability

The datasets generated during the current study are available in the SRA and GEO repositories with the following accession numbers.

SRA accession number for WES: PRJNA661624

GEO accession number for RNA-seq: GSE157601

## Acknowledgements

We thank Dr. Gary Wildey for critical discussion.

## Author contributions

H.M., A.D., and A.A. conceptualized the study. A.J.P. performed pathological evaluation; A.A. performed bioinformatic data analyses and interpretations; H.M. and A.D. provided clinical interpretation; H.M., A.D., and A.A. drafted the manuscript; and H.M. and A.A. supervised the study. All authors critically revised the manuscript and gave final approval.

## Supplementary information

Supplementary figures, supplementary tables, and supplementary methods are available online.

