## Supplementary Methods for "Genomic analyses of high-grade neuroendocrine gynecological malignancies reveal a unique mutational landscape and therapeutic vulnerabilities"

### Whole exome sequencing

1 μg genomic DNA was used for the WES library preparation. Sequencing libraries were generated using Agilent SureSelect Human All Exon kit (Agilent Technologies, Santa Clara, CA) following manufacturer's recommendations, and index codes were added to each sample. Briefly, fragmentation was carried out by the hydrodynamic shearing system (Covaris, Woburn, MA) to generate 180-280bp fragments. Remaining overhangs were converted into blunt ends via exonuclease/polymerase activities, and enzymes were removed. After adenylation of 3' ends of DNA fragments, adapter oligonucleotides were ligated. DNA fragments with ligated adapter molecules on both ends were selectively enriched in a PCR reaction. After the PCR reaction, the library was hybridized with the Liquid phase with a biotin-labeled probe, after which streptomycin-coated magnetic beads are used to capture the exons of genes. Captured libraries were enriched in a PCR reaction to add index tags to prepare for hybridization. Products were purified using the AMPure XP system (Beckman Coulter, Beverly, MA) and quantified using the Agilent high sensitivity DNA assay on the Agilent Bioanalyzer 2100 system. Sequencing was performed using 150-bp paired-end format on a NovaSeq 6000 (Illumina, San Diego, CA) sequencer.

### WES analyses and variant calling

FastQC (https://www.bioinformatics.babraham.ac.uk/projects/fastqc/) and adaptor trimming using Trim Galore (http://www.bioinformatics.babraham.ac.uk/projects/trim_galore/) were performed. We first generated the reference genome (hs38DH.fa) using GRCh38+ALT+decoy+HLA and then created the BWA index for mapping. Sequence reads were mapped with GRCh38 using BWA-MEM with default parameters to generate unsorted alignments (BAM) with ALT contigs aware mapping quality (1). Following GATK best practice (2,3), PCR duplicates were removed from sorted BAM files, and RG IDs were added using Picard tools (GitHub Repository http://broadinstitute.github.io/picard/). Subsequent realignment and base recalibration were performed using GATK4 (v 4.1.4.1). dbsnp_146.hg38.vcf, Mills_and_1000G_gold_standard.indels.hg38.vcf, and Homo_sapiens_assembly38.known_indels.vcf databases of known polymorphic sites were used to exclude regions around known polymorphisms. Variants were then called using GATK4 HaplotypeCaller with default parameters.

**Mutation Annotation Format (MAF) conversion and variants filtering**

VCF files were converted to MAF by mapping each variant to only one of all possible gene isoforms using the vcf2maf package (v 1.6.17) (https://github.com/ckandoth/vcf2maf). Ensembl Variant Effect Predictor (VEP) was used to determine the effect of variants (SNPs, insertions, deletions, CNVs, or structural variants) on genes, transcripts, and protein sequence, as well as regulatory regions (4). VEP is CLIA-compliant and uses HGVS variant format and Sequence Ontology nomenclature for variant effects. We used ExAC_nonTCGA.r0.3.1.sites.vep.vcf (germline variants called across thousands of normal samples excluding TCGA) for VEP (v 99.0) to filter variants. MAF files were further curated for missense mutations using SIFT (5) and PolyPhen-2 (6), and only deleterious missense mutations were kept.

**Copy number variations (CNVs) analysis**

CNVs were identified using XHMM (eXome-Hidden Markov Model, v 1.0) (7). First, the depth of coverage was calculated using GATK (v 3.8) with default parameters. The mean coverage for each exon interval for each sample was extracted and merged into a single sample-by-target matrix. GC content of each coding exons was calculated using GATK to exclude all exons with more than 90% or less than 10% GC content in the human reference sequence. Using Plink/Seq, a list of targets with low complexity was created based on the repeat-masked sequence fraction. Using XHMM 'matrix' command, the read-depth matrix was processed, and extreme GC content and low complexity lists were filtered out. PCA was run to determine the strongest independent ways (principal components) in which the data varies to normalize mean-centered data using this information. Next, we calculated the z-score of the per-sample read depth by centering relative to all target depths in that sample to remove any targets left with very high variance. Pre-normalized read depths, then taken and removed the same targets and samples that were removed during the normalization process. This matrix was used for annotation purposes in the subsequent CNV discovery and genotyping steps. CNVs were called using the hidden Markov model (HMM) Viterbi algorithm, and each called CNV was quantitatively genotyped using HMM forward-backward algorithm. XHMM output file (.xcnv) only contains chromosomal coordinates for deletion (DEL) and duplication (DUP). To get genes that fall under the DEL and DUP regions, we first prepared a bed file from .xcnv file using biomaRt (8) (R package, v 2.44.1). Next, we used GenomicRanges (9) (R package, v 1.40.0) and Homo.sapiens (R package, v 1.3.1, DOI: 10.18129/B9.bioc.Homo.sapiens) to get genes that fall under CNVs after converting hg38 coordinates to hg19 using the UCSC LiftOver tool as Homo.sapiens package is only compatible with hg19.

#### Mutational landscape, driver genes identification, and other analyses

To manage MAF files and visualize WES data, maftools (10) (R package, v 2.2.10) was used in the R environment. A copy number table of DEL and DUP was generated from CNVs data obtained from XHMM analysis and used along with the clinical annotation data table for generating various figures in maftools. MAF summary, transition and transversion mutations, somatic interactions, Oncoplots, and lollipop plots for amino acid changes were plotted using maftools. Oncoplots function that was used to generate Fig. 1A plotted SNV and CNV alterations together as 'multi-hit', if they happened to hit a given gene (a complete list of CNV data are included in Supplementary Table 1).

Cancer driver genes were detected based on the positional clustering method using the 'oncodrive' function of maftools. The oncodrive function is based on OncodriveCLUST algorithm (11) that measures genes' bias towards large mutation clustering. It uses a background model composed of coding-silent mutations that are not under selective pressure, therefore, provide the baseline clustering of somatic mutations. In principle, most of the variants in cancer-causing genes are enriched at a few specific hot-spots; on the basis of these positions, OncodriveCLUST can identify cancer genes. A minimum of 5 mutations per gene cutoff was used, and the p-value was calculated by z-score. Scatter plot in Fig. 1B was plotted with a false discovery rate (FDR) $<0.01$. Oncogenic Signaling Pathways were detected using the 'OncogenicPathways' function of maftools (Fig. 1C).

Tumor mutation burden (TMB) was calculated and compared against TCGA cohorts using the 'tcgaCompare' function of maftools. Mutation loads from 33 TCGA cohorts were used for this comparison (12) to generate Fig. 1D. We plotted Oncoplots for DNA-damage response (DDR) genes in Fig. 1E. A comprehensive DNA damage repair gene list was downloaded from Wood Laboratory at MD Anderson that is regularly curated (last curated on June 10$^{th}$, 2020; https://www.mdanderson.org/documents/Labs/Wood-Laboratory/human-dna-repair-genes.html), and the MAF file was subset for the DDR genes to use for Oncoplots (Fig 1E).

Potentially druggable gene categories and drug-gene interactions were detected using *drugInteractions* function of maftools. The *drugInteractions* uses drug–gene interactions and gene druggability information compiled from Drug Gene Interaction database (13) (http://www.dgidb.org).

**RNA-sequencing and analyses**

RNA was extracted from FFPE sections using the miRNeasy FFPE kit (Qiagen, Germantown, MD) according to the manufacturer's protocol. RNA quality was assessed via the Agilent 2100 Bioanalyzer (Agilent Technologies). Strand-specific RNA-seq library was prepared using NEBNext Ultra II Directional RNA Library Prep Kit (NEB, Ipswich, MA) according to the manufacturer's protocols. Briefly, the mRNA is fragmented randomly by adding a fragmentation buffer; then, the cDNA is synthesized by using an mRNA template and random hexamers primer, after which a custom second-strand synthesis buffer (Illumina), dNTPs, RNase H, and DNA polymerase I are added to initiate the second-strand synthesis. Terminal repair and sequencing adaptor ligation were performed, followed by size selection and PCR enrichment. Double-stranded cDNA libraries were purified using AMPure XP beads (Beckman Coulter) and quantified using the Agilent high sensitivity DNA assay on the Agilent Bioanalyzer 2100. Sequencing was performed using 150-bp paired-end format on a NovaSeq 6000 (Illumina) sequencer.

RNA-sequencing quality was checked by running FastQC, and TrimGalore was used for adapter and quality trimming. RNA-seq reads were mapped against hg38 using STAR (14) (v 2.7.0e) aligner with default parameters. DESeq2 (15) analysis with an adjusted P-value <0.001 was used to get a list of differentially expressed genes (DEGs). Top 5,000 significant DEGs (sorted by p-adj values) were used for unsupervised hierarchical clustering in Fig. 3B (NEC-GYN vs SCLC). Pathway analysis (GO biological process) was performed on Enrichr (https://amp.pharm.mssm.edu/Enrichr) database (16). RSEM (17) (v 1.3.2) analyses were performed to calculate FPKM and TPM values with default parameters.

**Fusion genes detection**

Novel and known somatic fusion genes were detected using FusionCatcher (18) (v 1.20). FusionCatcher is a powerful tool for finding somatic fusion genes in paired-end RNA-sequencing data. The fusion junctions were validated by using four different methods employing Bowtie, BLAT, STAR, and Bowtie2 aligners, and sequence analysis for ORFs (open reading frames) were also performed. Possible false positive and readthrough fusions were excluded.

**Immune cell gene signature analysis**

The 'ImSig' (R package, v 1.0.0) was used for immune cell gene signatures for profiling tumor microenvironment (19) . ImSig uses a set of immune gene signatures generated by a network-based deconvolution approach for seven immune cell types. Using ImSig algorithm with default parameter (correlation threshold, r = 0.7, over 75% genes overlap was observed), a table of the relative abundance of immune cells across samples was generated and plotted in Fig. 2C.
