## Supplementary Figures for "Genomic analyses of high-grade neuroendocrine gynecological malignancies reveal a unique mutational landscape and therapeutic vulnerabilities"

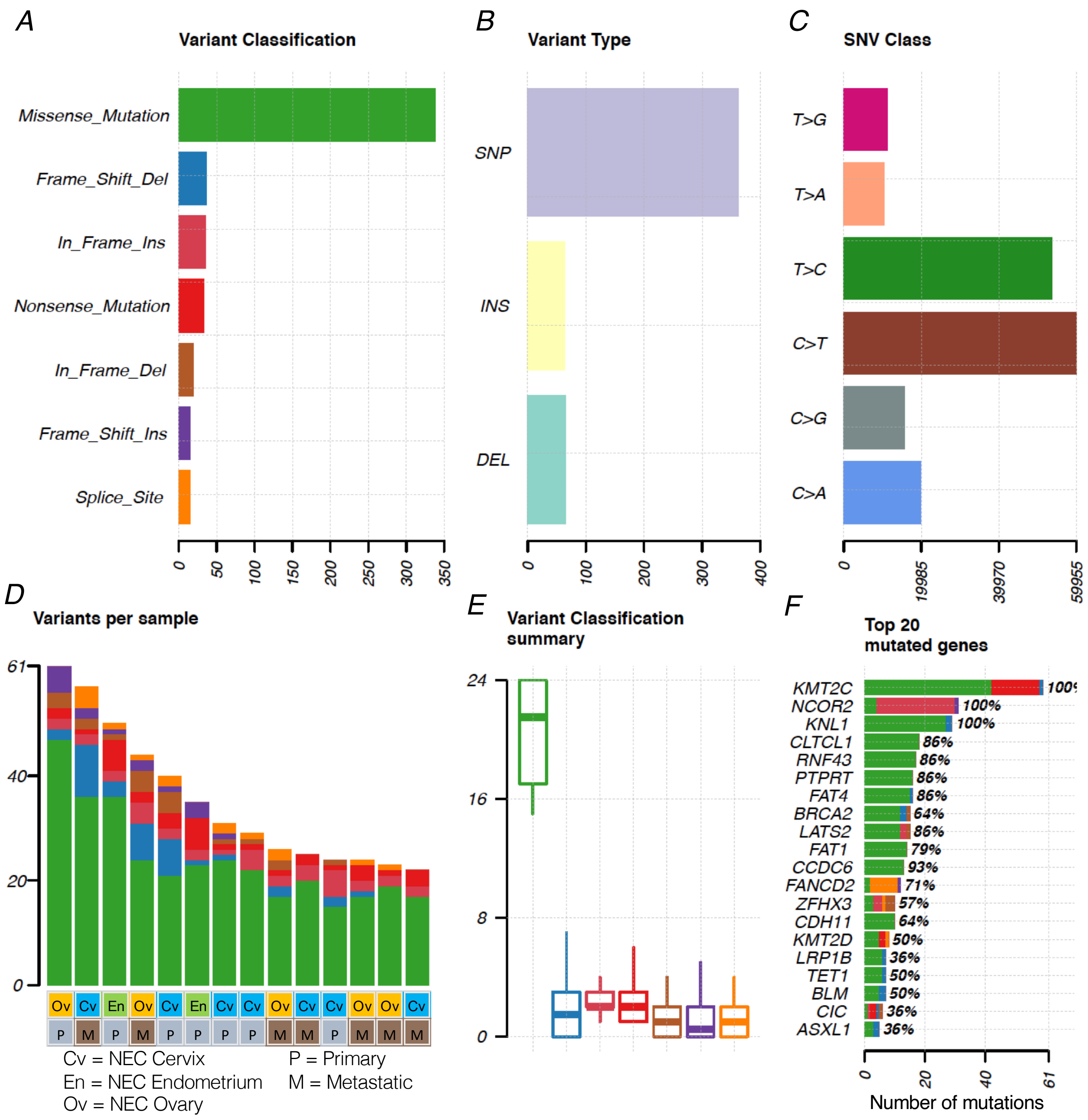


**Supplementary Fig. 1.** Mutational profile of neuroendocrine carcinoma of gynecologic origin (NEC-GYN) was detected by GATK Haplotype Caller. Figure panels representing **A)** various variants classifications; **B)** variant type; **C)** SNV class; **D)** numbers of variants in each samples along with **E)** variant classification summary; and **F)** frequently mutated genes in our NEC-GYN cohort.

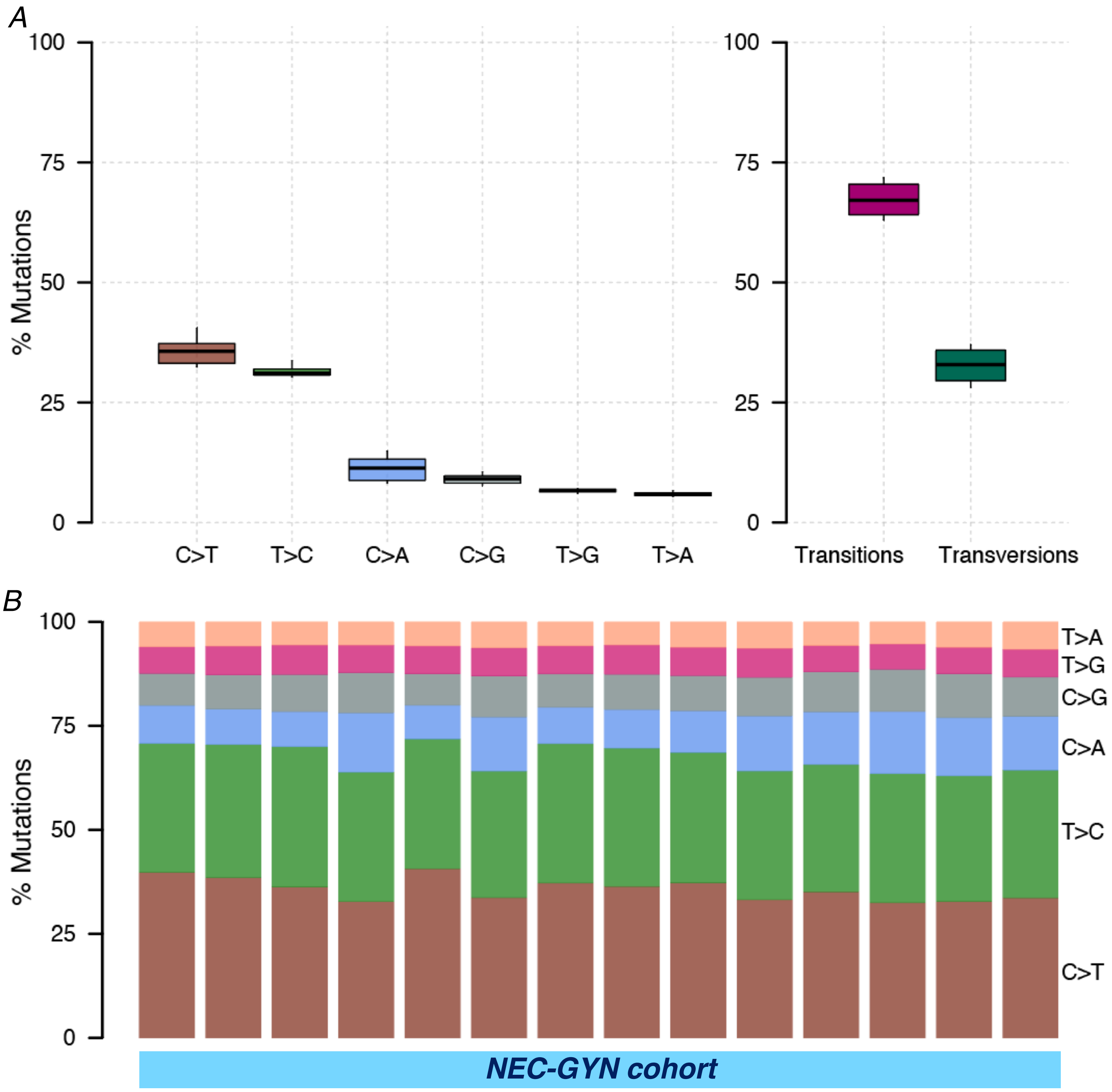


**Supplementary Fig. 2. A)** Percentages of various CNV classes in NEC-GYN (left) and transition & transversion mutations (right); and **B)** percentages of various CNV classes in various tumor sample in our cohort.

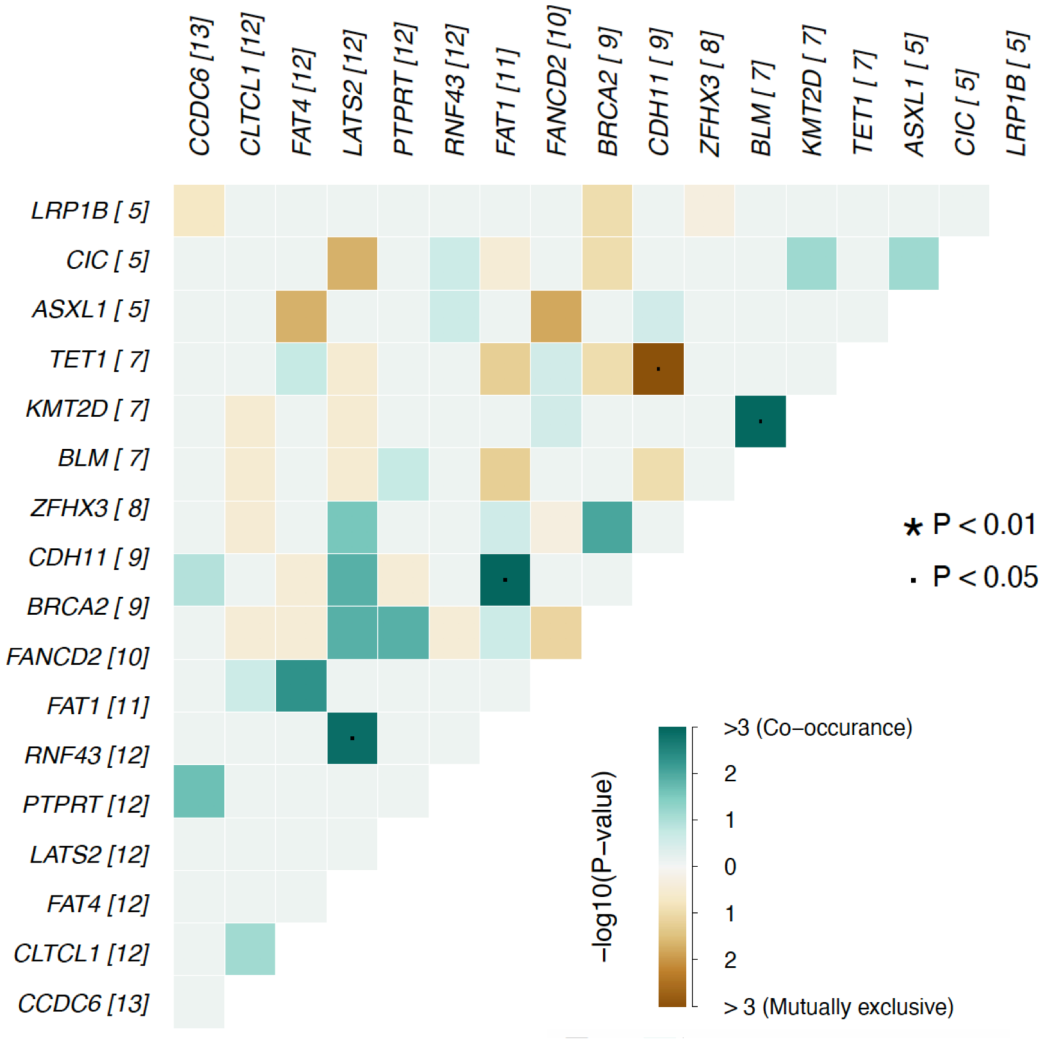


**Supplementary Fig. 3.** Mutually exclusive or co-occurring set of genes. Numbers in parenthesis indicate for the given gene mutated in total tumor samples out of 14 (cohort size). Pair-wise Fisher's Exact test was used to detect significant pair of genes.

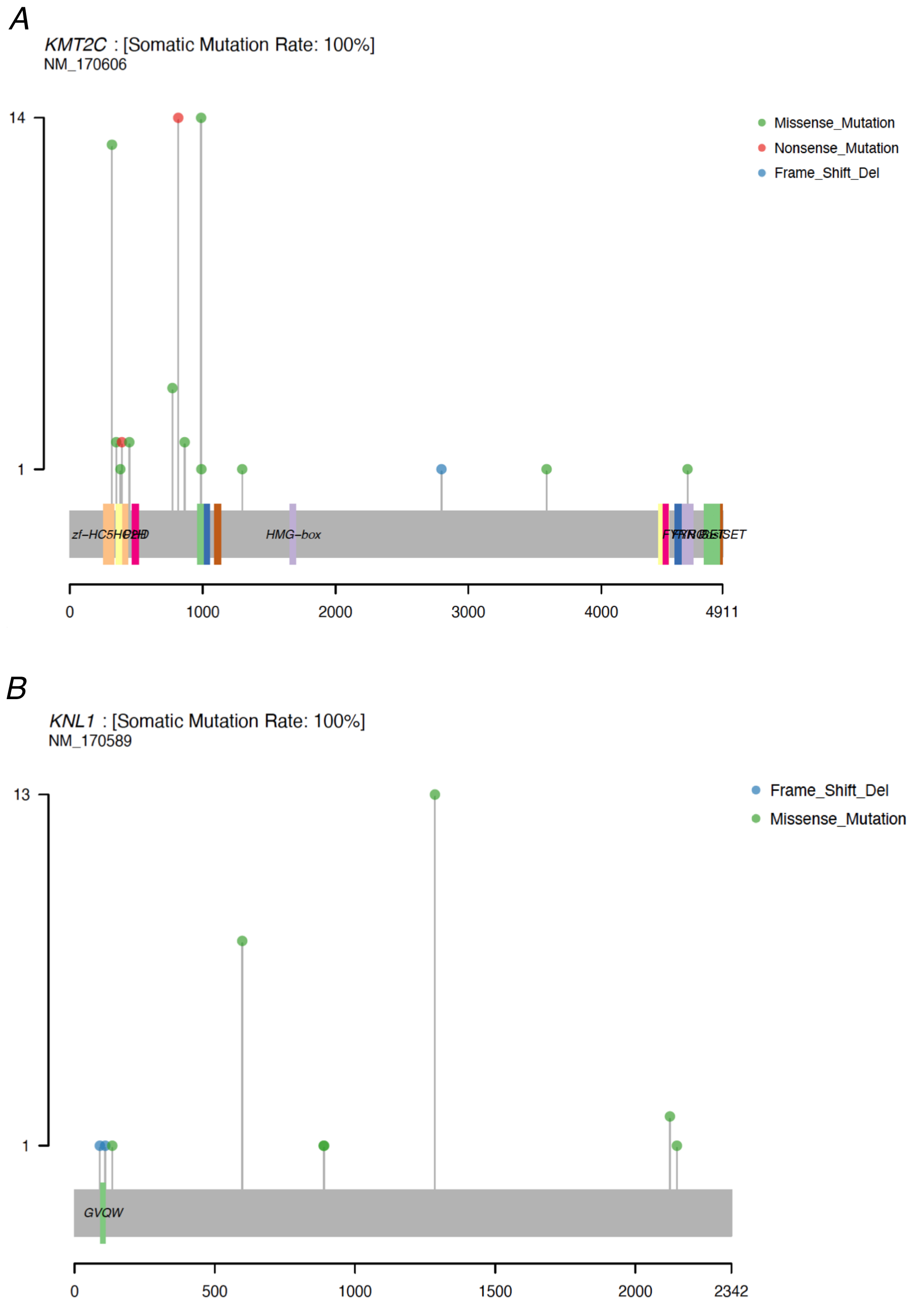


**Supplementary Fig. 4.** Lollipop plot showing the mutations (amino acid changes) in **A**) *KMT2C*; and **B**) *KNL1* gene in our cohort (NEC-GYN).

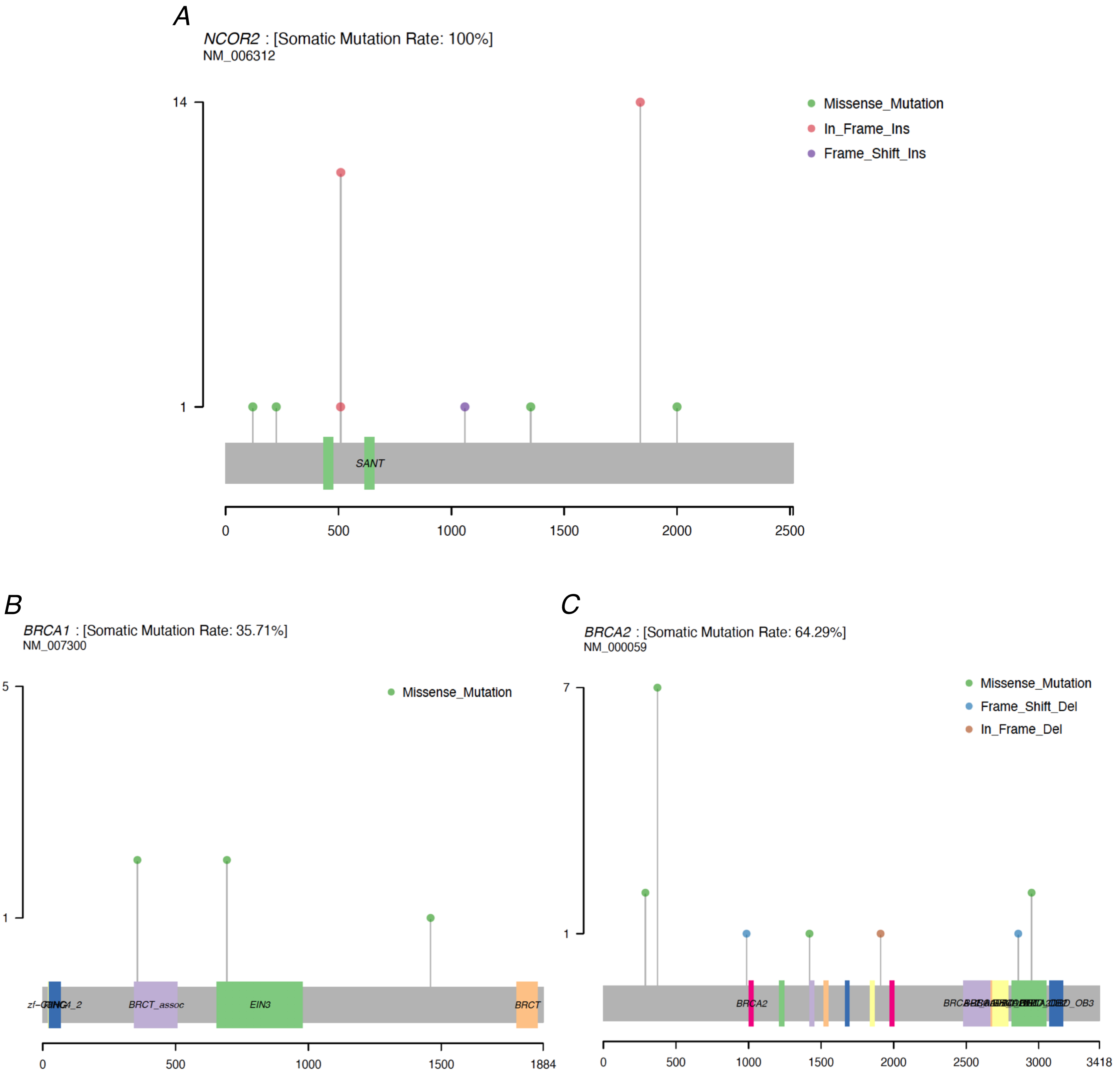


**Supplementary Fig. 5.** Lollipop plot showing the mutations (amino acid changes) in **A**) *NCOR2*; **B**) *BRCA1*; and **C**) *BRCA2* gene in NEC-GYN.

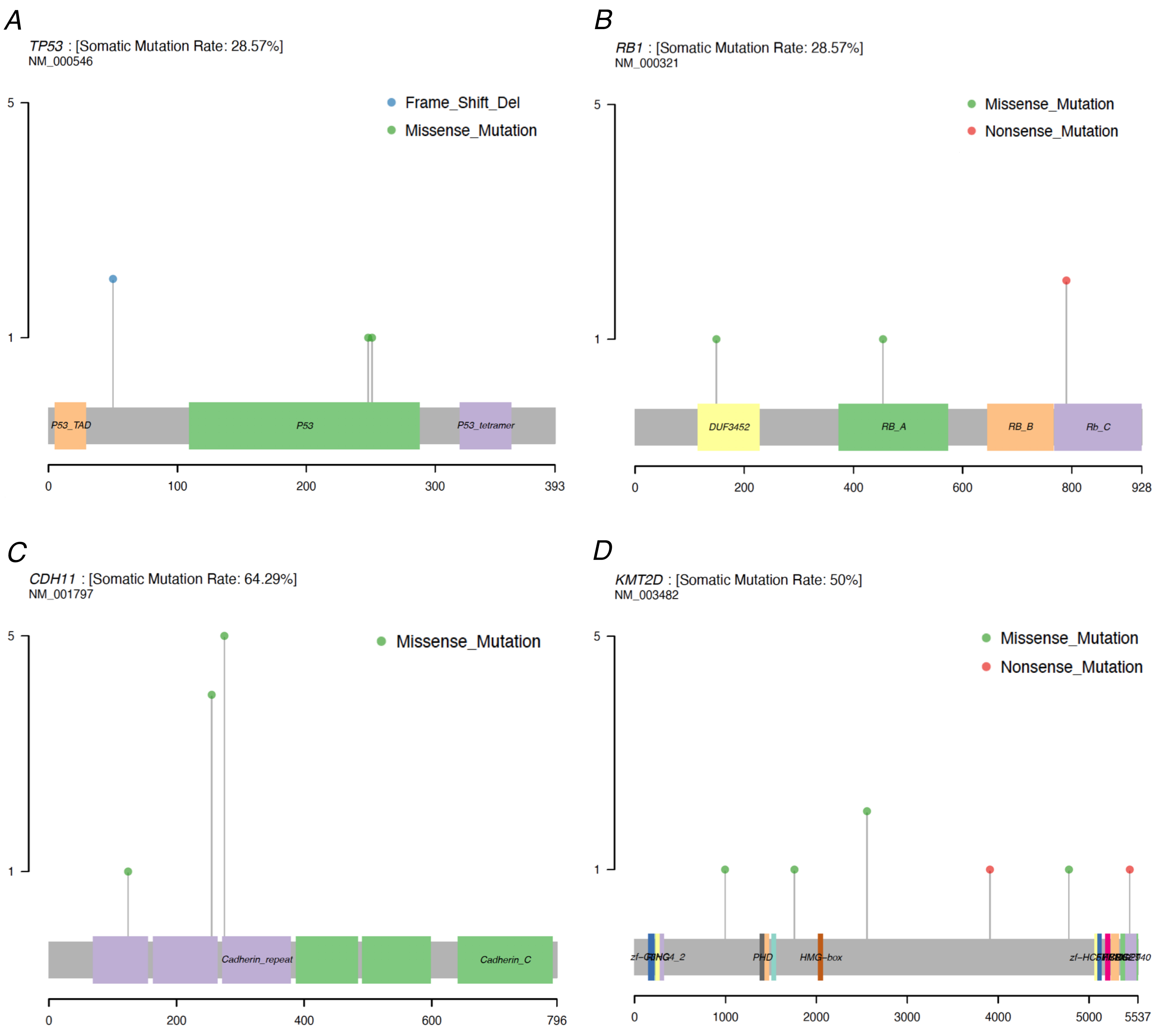


**Supplementary Fig. 6.** Lollipop plot showing the mutations (amino acid changes) in **A**) *TP53*; and **B**) *RB1* gene. *TP53* and *RB1* genes are highly mutated in SCLC but moderately mutated in our NEC-GYN cohort. Lollipop plot showing the mutations (amino acid changes) in **C**) *CDH11*; and **D**) *KMT2D* gene.

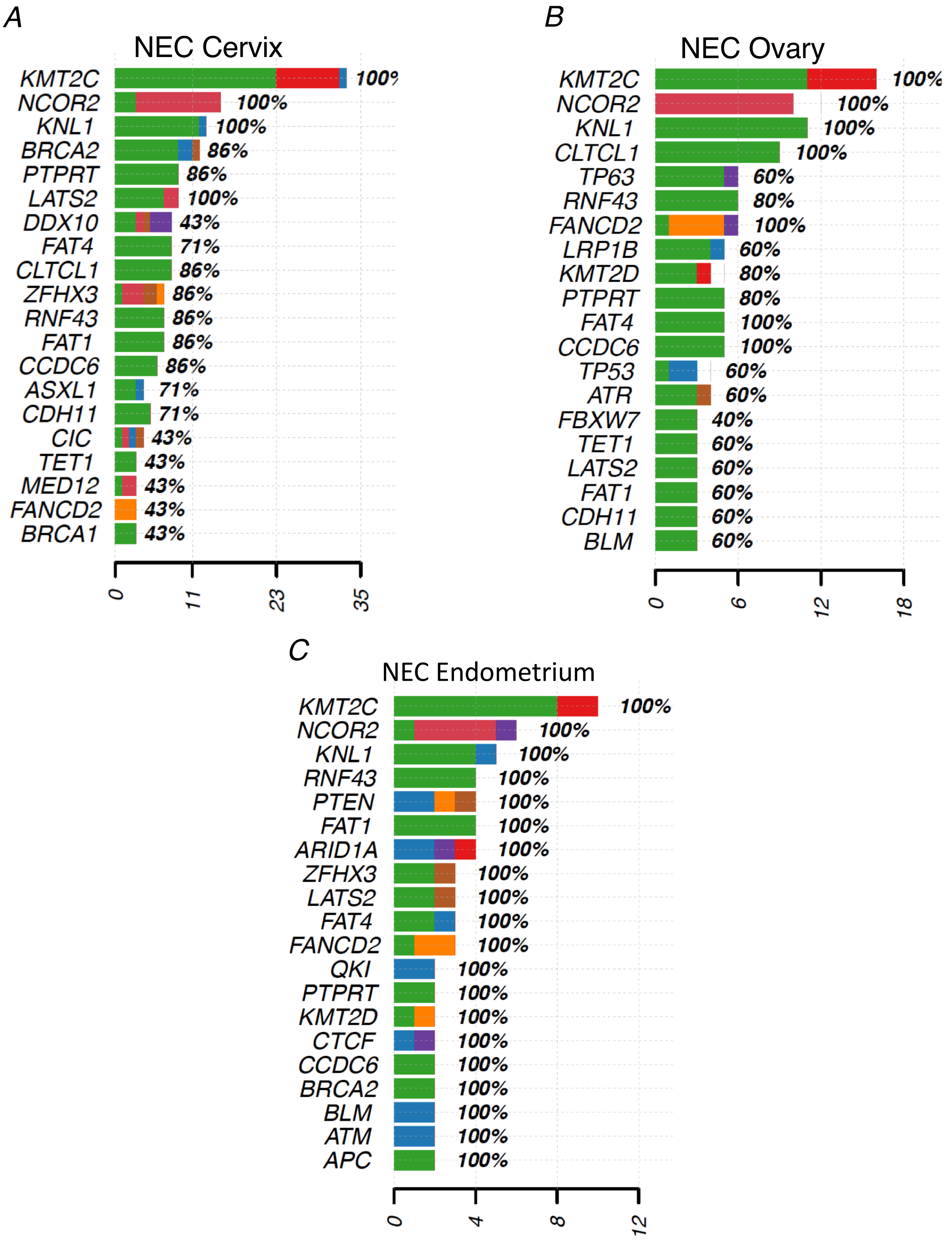


**Supplementary Fig. 7.** Cervical NEC samples showing higher number of mutations within individual genes compared to Ovary and Endometrial NEC. Frequently mutated genes in **A**) cervical; **B**) ovarian; and **C**) endometrial carcinoma. X-axis representing number of mutations. See Supplementary Fig.1 for color code for various variants.

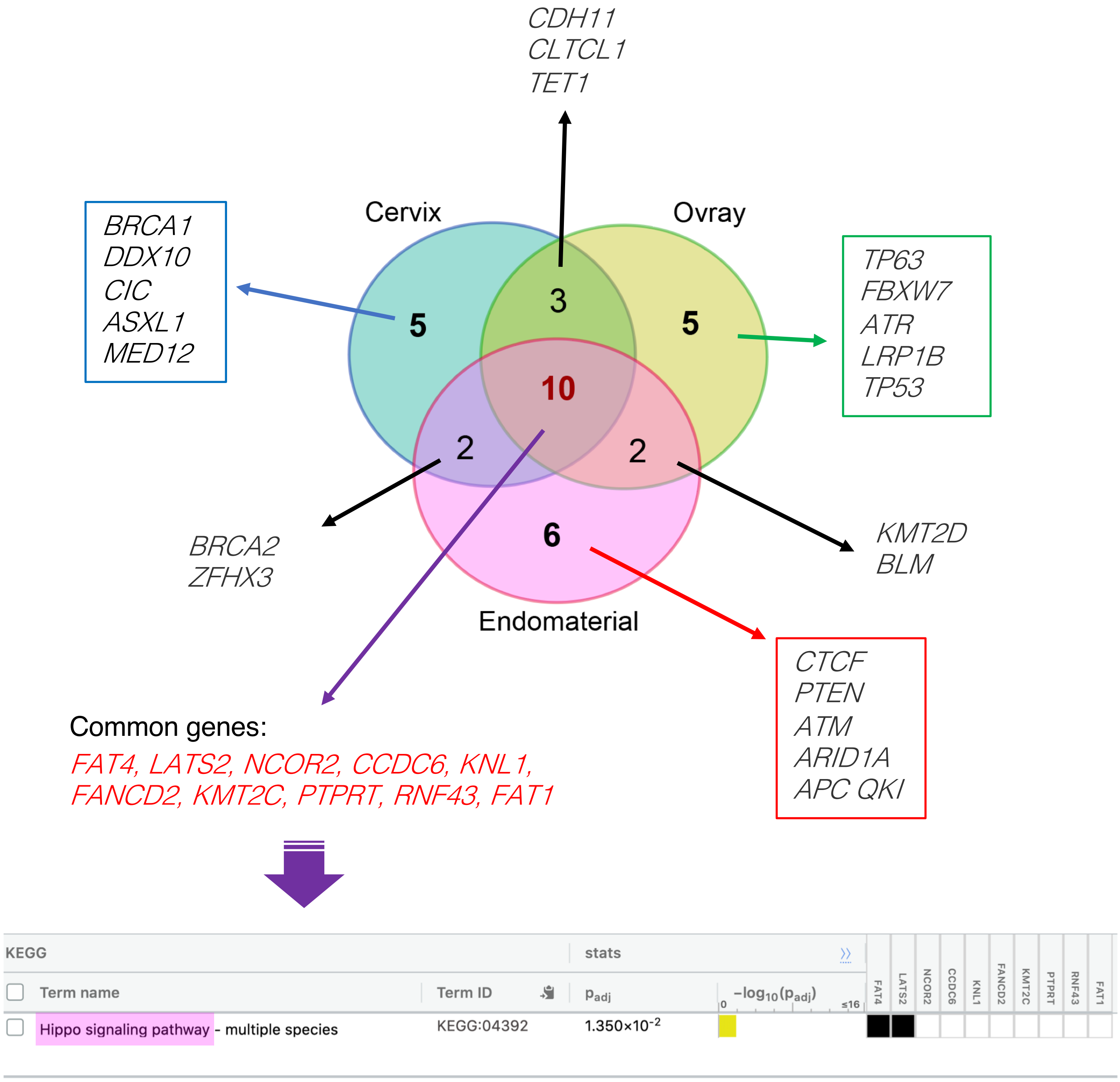


**Supplementary Fig. 8.** Venn diagram representing common mutated genes between Cervical, Ovarian, and Endometrial NEC (top 20 frequently mutated genes from each of three groups were used). The altered genes that were common between tumors from three gynecologic sites are significantly enriched for the Hippo signaling pathway (bottom).

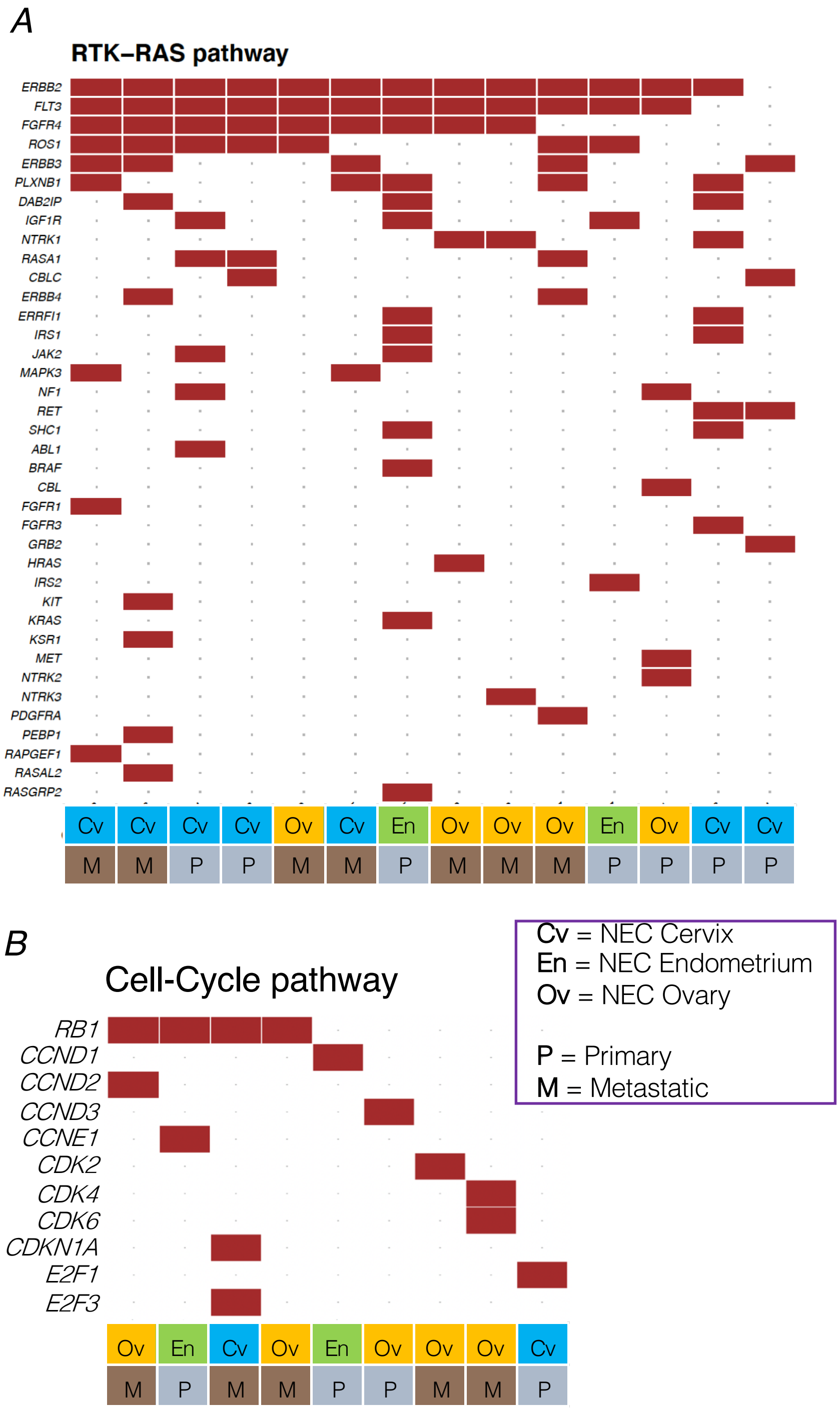


**Supplementary Fig. 9.** Oncogenic pathways affected in NEC-GYN. **A**) RTK-RAS pathway; and **B**) Cell-Cycle pathway.

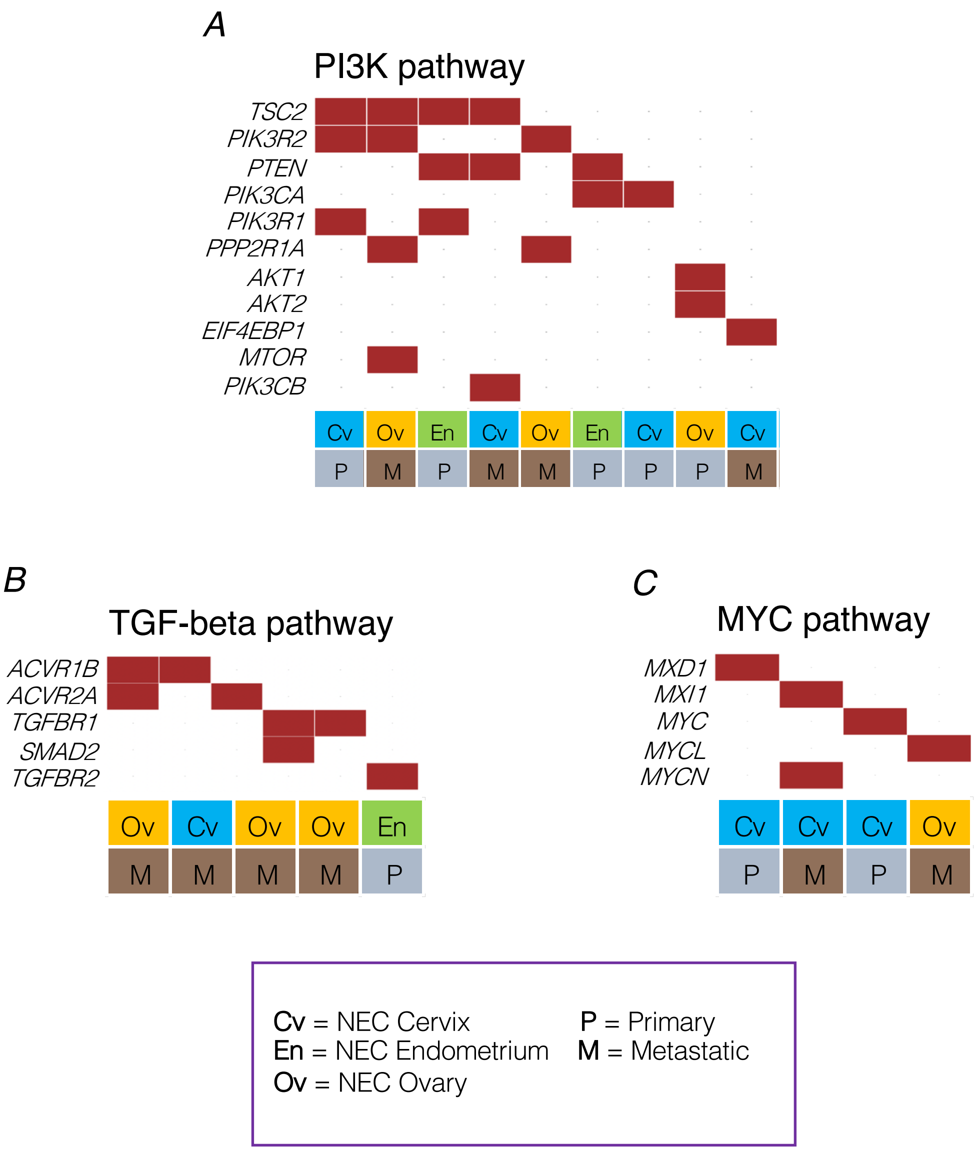


**Supplementary Fig. 10.** Oncogenic pathways affected in NEC-GYN. **A**) PI3K pathway; **B**) TGF-beta; and **C**) MYC pathway.

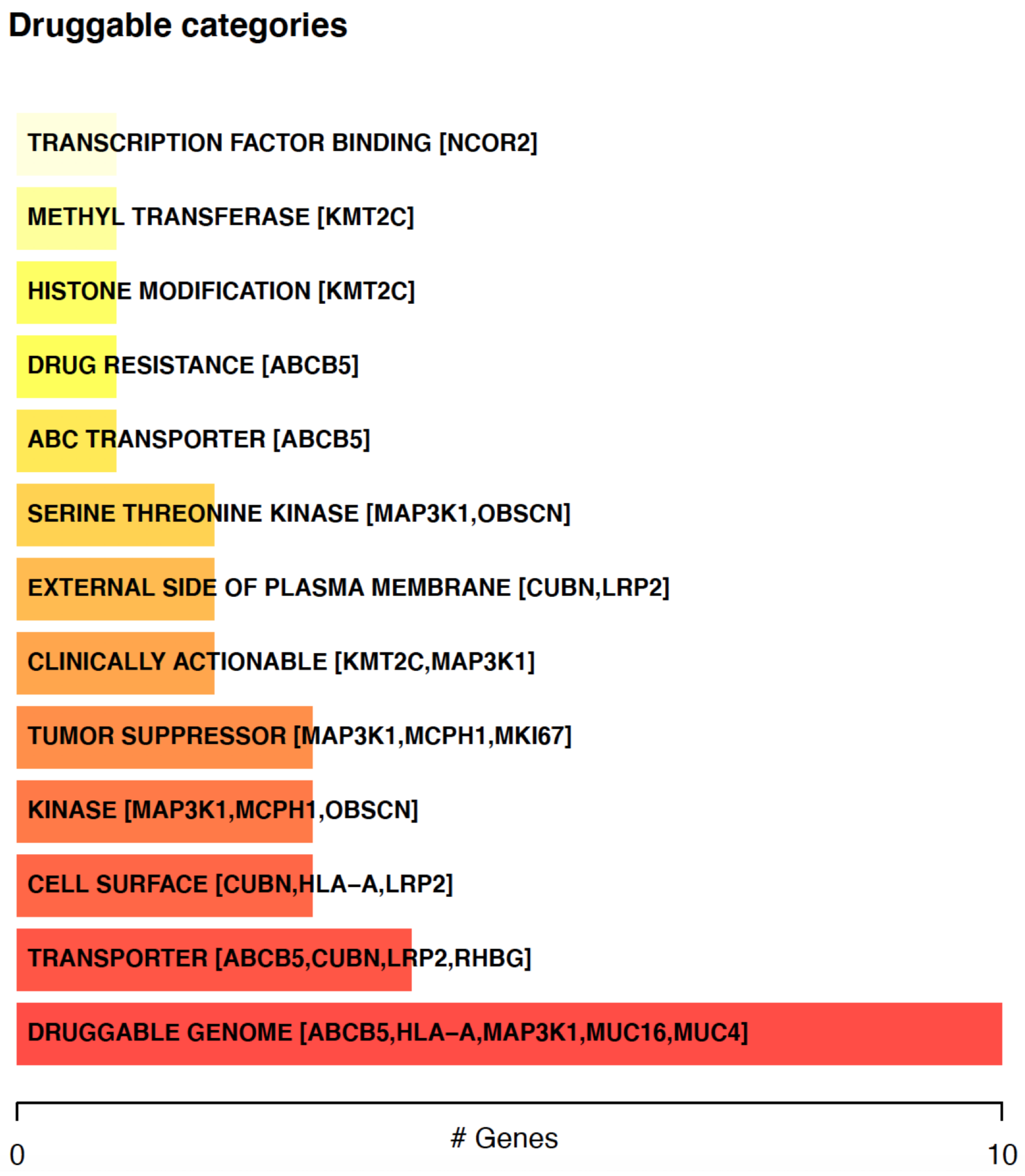


**Supplementary Fig. 11.** Various potentially druggable targets were identified by drug-gene interactions analysis using GDIdb.

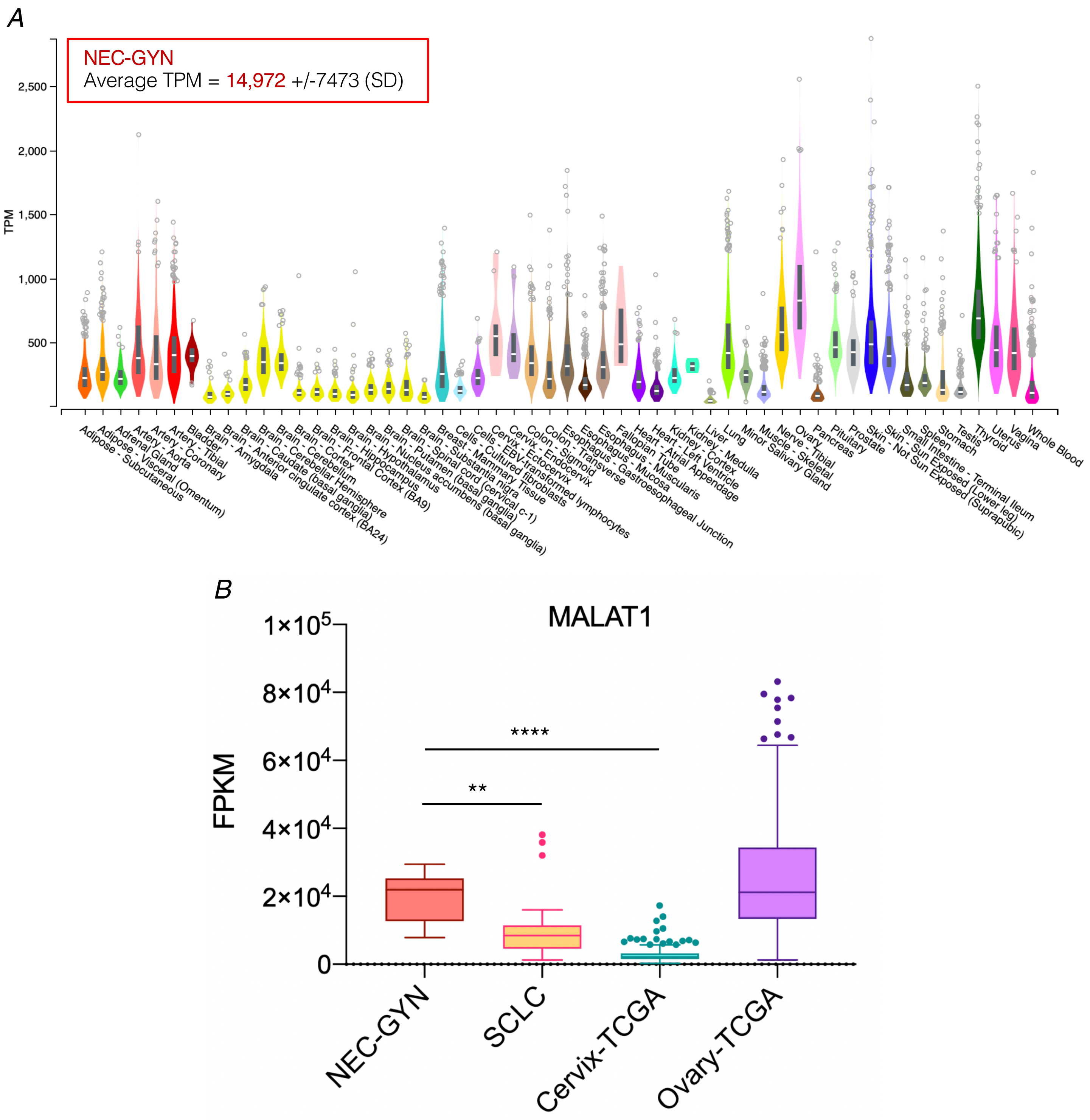


**Supplementary Fig. 12. A**) MALAT1 expression levels across various normal human tissues from GETx. TPM value of MALAT1 in NEC-GYN (shown in the box) is much higher than all of the GETx normal tissues. **B**) MALAT1 expression level in NEC-GYN compared to SCLC and TCGA cohort of cervical and ovarian cancers (Tukey, $^{**}P = 0.0012$, $^{****}P > 0.0001$ by two-tailed Mann–Whitney U test).

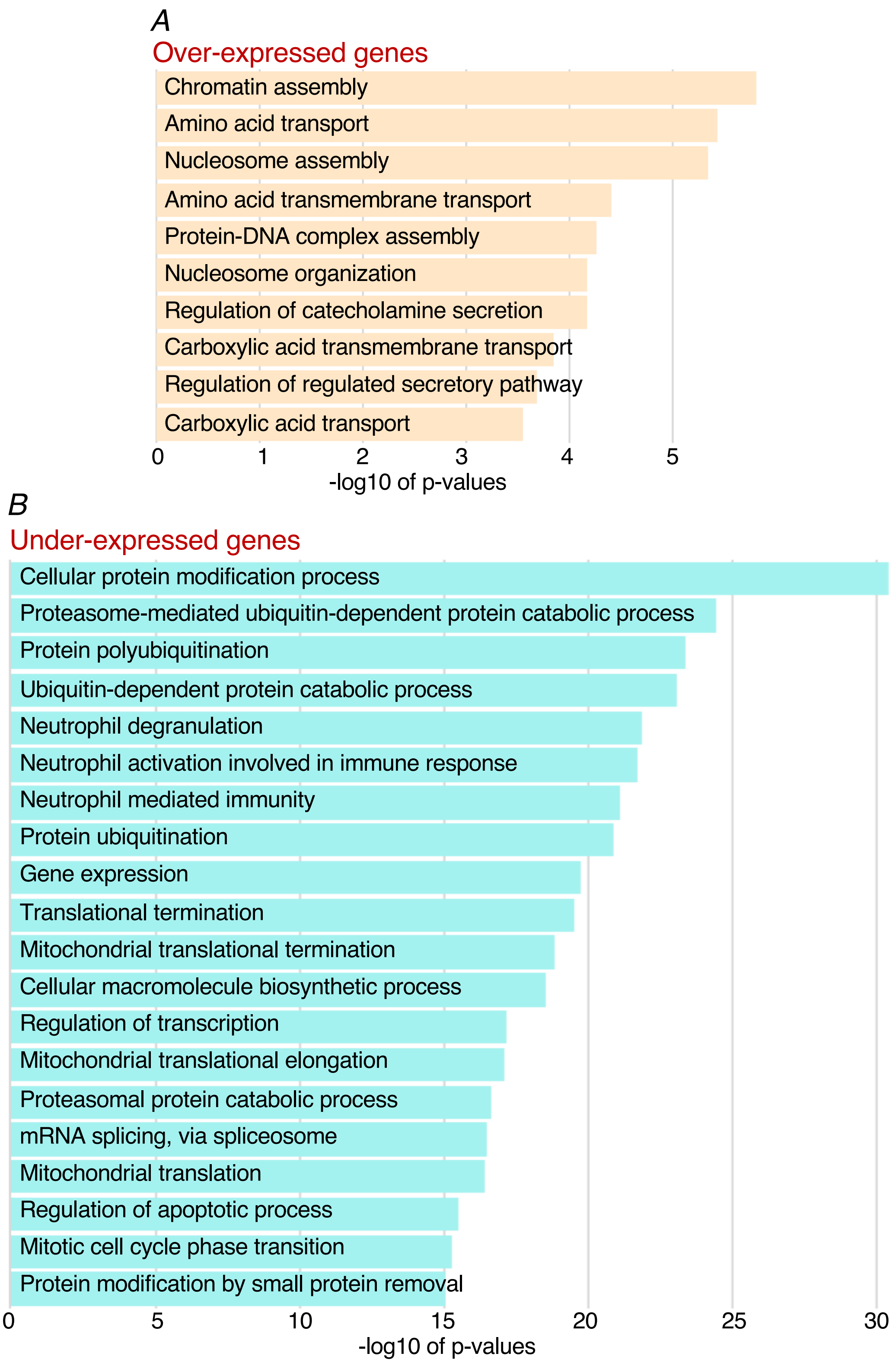


**Supplementary Fig. 13.** GO analysis of differentially expressed genes in NEC-GYN compared to Cervical cancer (TCGA). Significantly enriched pathways in **A**) over-expressed genes (4-folds cutoff); and **B**) under-expressed genes (10-folds cutoff).

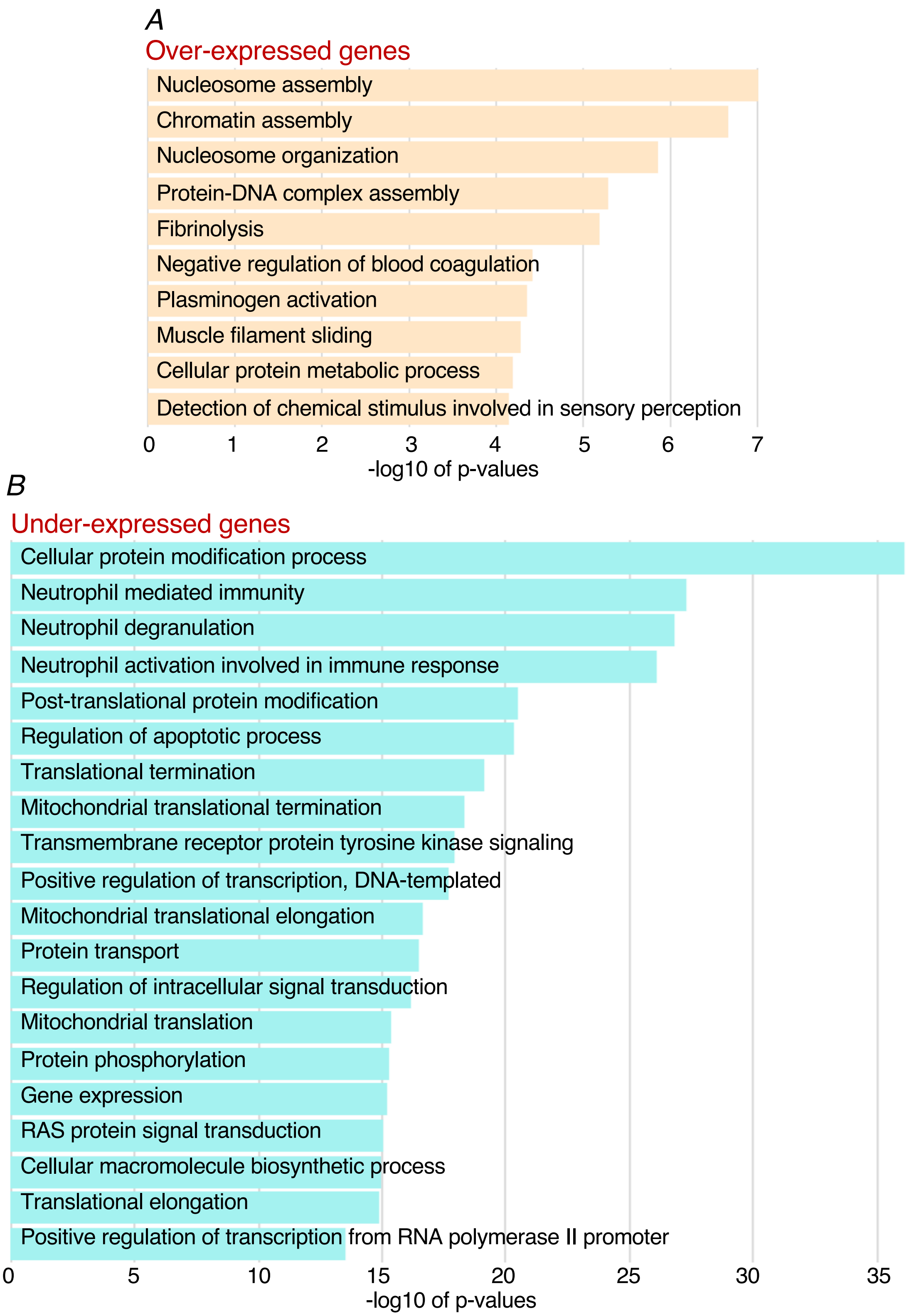


**Supplementary Fig. 14.** GO analysis of differentially expressed genes in NEC-GYN compared to Ovarian cancer (TCGA). Significantly enriched pathways in **A**) over-expressed genes (4-folds cutoff); and **B**) under-expressed genes (10-folds cutoff).

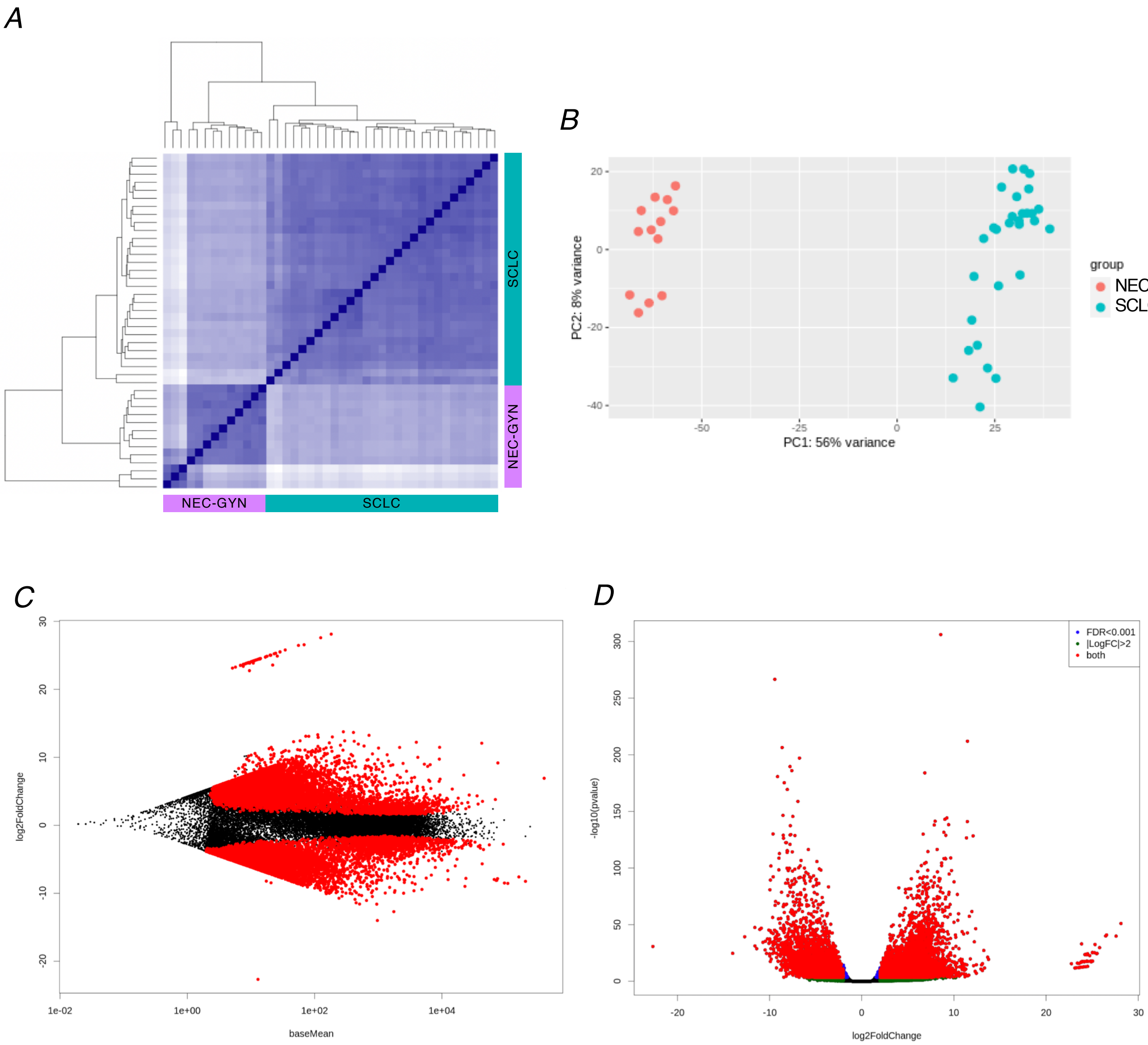


**Supplementary Fig. 15.** Comparison of NEC-GYN with SCLC. **A**) Samples clustering; and **B**) PCA plot are shown. Differential gene expressions in NEC-GYN vs SCLC are represented by **C**) MA plot; and **D**) Volcano plot.

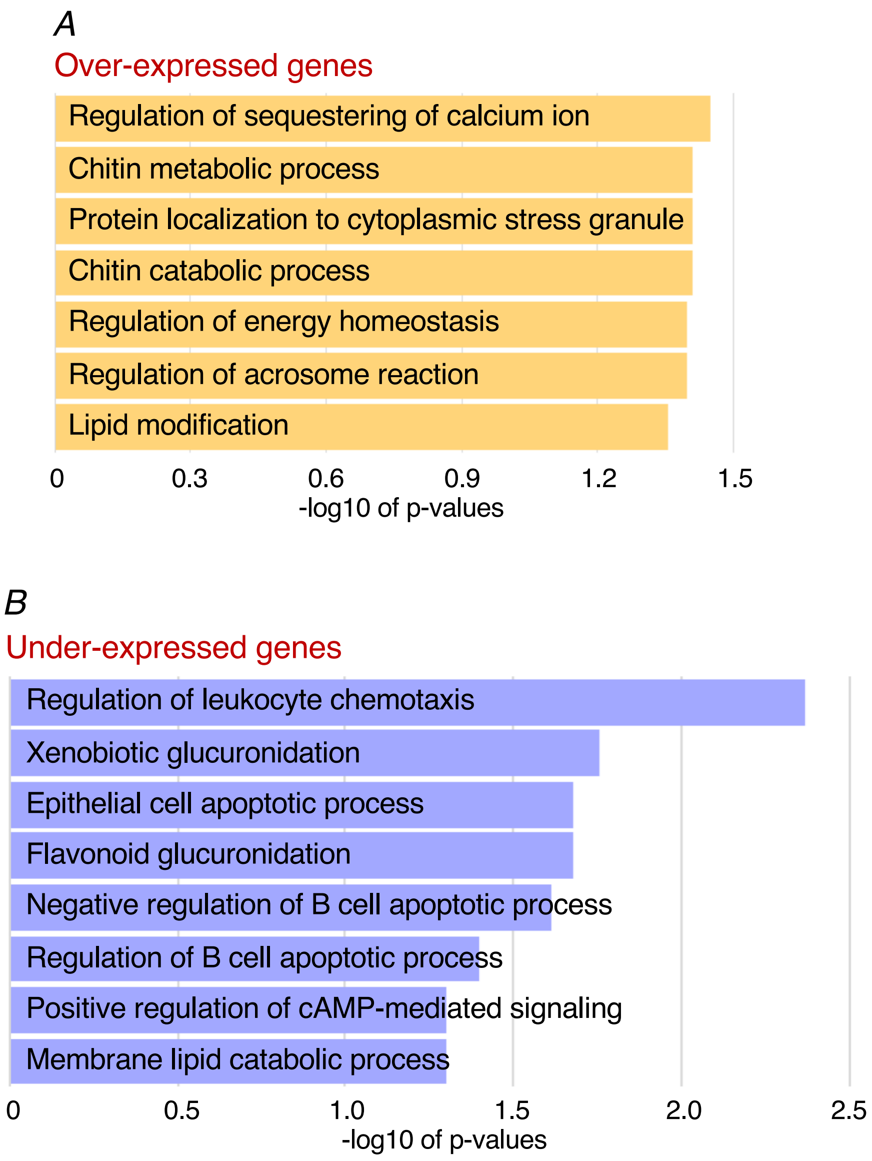


**Supplementary Fig. 16.** GO analysis of differentially expressed genes in NEC-GYN compared to SCLC (p-adj <0.001, fold-change >2). Significantly enriched pathways in **A**) over-expressed genes; and **B**) under-expressed genes.

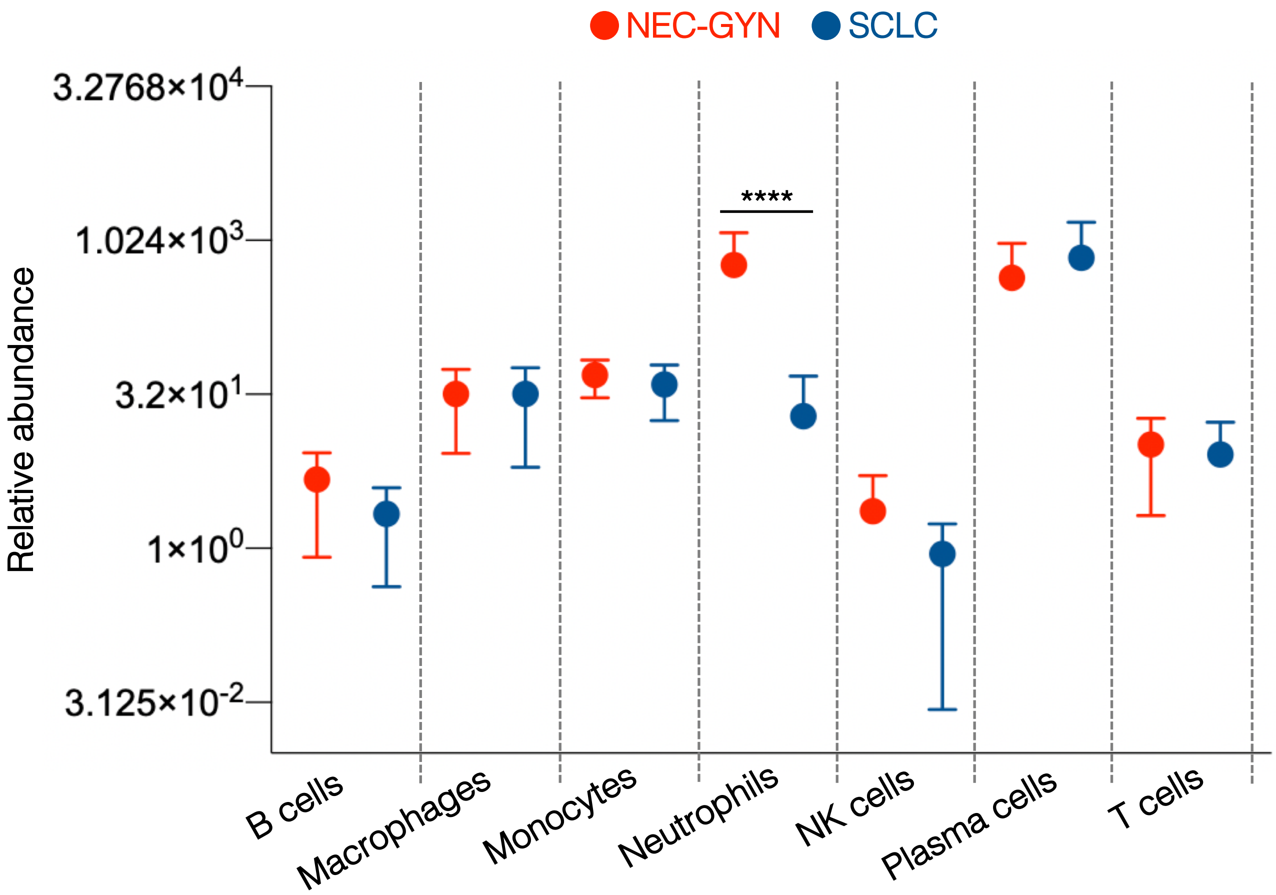


**Supplementary Fig. 17.** Comparison of relative abundance of immune cells in NEC-GYN and SCLC tumors based on network-based deconvolution (ImSig) analysis (mean with SD, ****P ≤ 0.0001 by two-tailed Mann–Whitney U test).
