## Supplementary Table 1 for "Genomic analyses of high-grade neuroendocrine gynecological malignancies reveal a unique mutational landscape and therapeutic vulnerabilities"

**Supplementary Table 1.** Copy number variations (CNV) detected by XHMM *(Table is sorted by chromosome intervals)*

| SAMPLE_ID | SAMPLE | CNV | INTERVAL | AFFECTED GENES |
|---|---|---|---|---|
| TKHS190002641 | NEC Ovary | DUP | chr1:10376545-10419749 | *KIF1B, PGD* |
| TKHS190002641 | NEC Ovary | DUP | chr1:109739833-109985172 | *AHCYL1, CSF1, GSTM3, EPS8L3* |
| TKHS190002657 | NEC Ovary | DEL | chr1:11502678-11787493 | *MAD2L2, FBXO2, FBXO6, DRAXIN, MTHFR, AGTRAP, DISP3, FBXO44* |
| TKHS190002657 | NEC Ovary | DEL | chr1:11858214-12207020 | *MIR4632, NPPB, PLOD1, MIIP, TNFRSF1B, KIAA2013, TNFRSF8, MFN2* |
| TKHS190002646 | NEC Cervix | DUP | chr1:11956917-11999095 | *PLOD1, MFN2* |
| TKHS190002641 | NEC Ovary | DUP | chr1:12774868-12893474 | *HNRNPCL1, PRAMEF10, PRAMEF12, PRAMEF4, PRAMEF11, HNRNPCL3, PRAMEF1, PRAMEF2* |
| TKHS190002646 | NEC Cervix | DUP | chr1:1309086-1629566 | *ACAP3, PUSL1, MIB2, MRPL20-AS1, DVL1, ATAD3C, SSU72, TMEM240, ANKRD65, MXRA8, INTS11, AURKAIP1, MRPL20, ATAD3A, TMEM88B, FNDC10, VWA1, CPTP, CCNL2, TAS1R3, ATAD3B* |
| TKHS190002653 | NEC Ovary | DEL | chr1:1310634-1336515 | *PUSL1, DVL1, INTS11, CPTP, TAS1R3* |
| TKHS190002657 | NEC Ovary | DEL | chr1:1315404-1387582 | *DVL1, MXRA8, INTS11, AURKAIP1, CPTP, CCNL2, TAS1R3* |
| TKHS190002657 | NEC Ovary | DEL | chr1:1436807-1545002 | *ATAD3C, SSU72, TMEM240, ATAD3A, VWA1, ATAD3B* |
| TKHS190002655 | NEC Cervix | DEL | chr1:145918432-145995561 | *NBPF10, NBPF20, TXNIP, LIX1L, ANKRD34A, POLR3GL, PEX11B, RBM8A* |
| TKHS190002656 | NEC Cervix | DUP | chr1:147252623-147271005 | *NA, NA, NA, CHD1L* |
| TKHS190002657 | NEC Ovary | DEL | chr1:150497530-150556793 | *MIR4257, FALEC, ECM1, ADAMTSL4, TARS2* |
| TKHS190002657 | NEC Ovary | DEL | chr1:152622834-152884773 | *C1orf68, LCE4A, LCE2B, LCE1A, LCE1B, LCE1C, LCE1D, LCE1E, LCE1F, LCE2A, LCE2C, LCE2D, LCE3A, SMCP, KPRP, LCE6A* |
| TKHS190002644 | NEC Cervix | DUP | chr1:153310611-153360838 | *PGLYRP3, PGLYRP4, S100A9* |
| TKHS190002644 | NEC Cervix | DUP | chr1:155023117-155058474 | *DCST1-AS1, DCST2, DCST1, ADAM15* |
| TKHS190002644 | NEC Cervix | DUP | chr1:155175386-155192843 | *GBAP1, MTX1, THBS3* |
| TKHS190002647 | NEC Endometrium | DEL | chr1:155202812-155241112 | *GBA, GBAP1, MTX1, THBS3* |
| TKHS190002643 | NEC Cervix | DUP | chr1:155203086-155240717 | *GBA, GBAP1, MTX1, THBS3* |
| TKHS190002641 | NEC Ovary | DUP | chr1:156566390-156652966 | *IQGAP3, NAXE, TTC24, GPATCH4, HAPLN2, BCAN* |
| TKHS190002647 | NEC Endometrium | DEL | chr1:156726906-156768239 | *HDGF, RRNAD1, PRCC, MRPL24, ISG20L2* |
| TKHS190002645 | NEC Endometrium | DUP | chr1:15727014-15738245 | *PLEKHM2, SLC25A34* |
| TKHS190002644 | NEC Cervix | DUP | chr1:161001020-161124462 | *TSTD1, KLHDC9, ARHGAP30, NIT1, F11R, PFDN2, USF1, NECTIN4, DEDD* |
| TKHS190002644 | NEC Cervix | DUP | chr1:161166848-161202480 | *NDUFS2, PPOX, B4GALT3, ADAMTS4* |
| TKHS190002657 | NEC Ovary | DEL | chr1:16125215-16207210 | *ARHGEF19, EPHA2* |
| TKHS190002650 | NEC Ovary | DUP | chr1:16129434-16206543 | *ARHGEF19, EPHA2* |
| TKHS190002643 | NEC Cervix | DEL | chr1:16922019-16946991 | *CROCC* |
| TKHS190002657 | NEC Ovary | DEL | chr1:16987046-16996164 | *ATP13A2* |
| TKHS190002642 | NEC Ovary | DUP | chr1:171781968-173050962 | *MIR3120, DNM3OS, DNM3, FASLG, SUCO, EEF1AKNMT, PIGC, TNFSF18, C1orf105* |
| TKHS190002657 | NEC Ovary | DEL | chr1:175093990-175128746 | *TNN* |

| | | | | |
|---|---|---|---|---|
| TKHS190002646 | NEC Cervix | DUP | chr1:1752904-1804581 | *GNB1, NADK* |
| TKHS190002641 | NEC Ovary | DUP | chr1:1752904-3816370 | *LOC100129534, NA, TTC34, MIR4251, RER1, TNFRSF14-AS1, TPRG1L, PRXL2B, ACTRT2, CCDC27, CALML6, MEGF6, FAAP20, GABRD, ARHGEF16, GNB1, TMEM52, HES5, SMIM1, PRDM16-DT, WRAP73, PEX10, PANK4, PRKCZ, TP73-AS1, LRRC47, PRDM16, SKI, NADK, MIR551A, TP73, MMEL1, MORN1, CFAP74, TNFRSF14, PLCH2, CEP104* |
| TKHS190002656 | NEC Cervix | DUP | chr1:179639839-180080661 | *FAM163A, TDRD5, TOR1AIP2, TOR1AIP1, CEP350* |
| TKHS190002653 | NEC Ovary | DEL | chr1:1923367-1959209 | *CFAP74* |
| TKHS190002646 | NEC Cervix | DUP | chr1:1923367-2229735 | *FAAP20, GABRD, PRKCZ, SKI, CFAP74* |
| TKHS190002641 | NEC Ovary | DUP | chr1:19267300-19304386 | *LOC100506730, AKR7A3, AKR7L, AKR7A2* |
| TKHS190002646 | NEC Cervix | DUP | chr1:19269272-19304386 | *LOC100506730, AKR7A3, AKR7L, AKR7A2* |
| TKHS190002657 | NEC Ovary | DEL | chr1:19270748-19327412 | *LOC100506730, AKR7A3, AKR7L, SLC66A1, AKR7A2* |
| TKHS190002656 | NEC Cervix | DUP | chr1:193081364-193212477 | *MIR1278, GLRX2, RO60, CDC73, B3GALT2* |
| TKHS190002654 | NEC Cervix | DEL | chr1:200857754-200913629 | *CAMSAP2, GPR25, INAVA* |
| TKHS190002653 | NEC Ovary | DUP | chr1:201009266-201110269 | *KIF21B, CACNA1S* |
| TKHS190002642 | NEC Ovary | DUP | chr1:201075495-201110269 | *CACNA1S* |
| TKHS190002641 | NEC Ovary | DEL | chr1:203020793-203217133 | *CHI3L1, CHIT1, ADORA1, MYBPH, MYOG, PPFIA4, TMEM183A* |
| TKHS190002642 | NEC Ovary | DUP | chr1:203048227-203225870 | *CHI3L1, CHIT1, ADORA1, MYBPH, MYOG, PPFIA4* |
| TKHS190002647 | NEC Endometrium | DEL | chr1:203165056-203503720 | *CHI3L1, CHIT1, ADORA1, FMOD, OPTC, MYBPH, PRELP, LINC01136, BTG2* |
| TKHS190002653 | NEC Ovary | DUP | chr1:203165261-203217133 | *CHI3L1, CHIT1, ADORA1, MYBPH* |
| TKHS190002654 | NEC Cervix | DUP | chr1:20342432-20354263 | *VWA5B1* |
| TKHS190002644 | NEC Cervix | DUP | chr1:204433316-204450017 | *PIK3C2B* |
| TKHS190002647 | NEC Endometrium | DEL | chr1:205256330-205531378 | *LEMD1-AS1, MIR135B, CDK18, KLHDC8A, NUAK2, LEMD1, TMCC2* |
| TKHS190002654 | NEC Cervix | DEL | chr1:205845325-206012649 | *SLC26A9, LOC284581* |
| TKHS190002644 | NEC Cervix | DUP | chr1:205928827-205935820 | *SLC26A9* |
| TKHS190002642 | NEC Ovary | DUP | chr1:208027243-208060837 | *PLXNA2* |
| TKHS190002656 | NEC Cervix | DUP | chr1:211987886-212080750 | *MIR3122, INTS7, DTL* |
| TKHS190002657 | NEC Ovary | DEL | chr1:21219955-21290198 | *ECE1* |
| TKHS190002656 | NEC Cervix | DUP | chr1:212796517-212951818 | *TATDN3, SPATA45, FLVCR1, FLVCR1-DT, VASH2* |
| TKHS190002656 | NEC Cervix | DUP | chr1:220073967-220145653 | *BPNT1, RNU5F-1, MIR194-1, MIR215, IARS2* |
| TKHS190002656 | NEC Cervix | DUP | chr1:228103134-228350057 | *MRPL55, IBA57, GUK1, GJC2, OBSCN-AS1, C1orf35, OBSCN* |
| TKHS190002648 | NEC Cervix | DEL | chr1:228277505-228338372 | *OBSCN* |
| TKHS190002650 | NEC Ovary | DUP | chr1:228277505-228375824 | *OBSCN* |
| TKHS190002657 | NEC Ovary | DEL | chr1:22896411-22965338 | *EPHB2* |
| TKHS190002656 | NEC Cervix | DUP | chr1:246889966-246945827 | *ZNF670-ZNF695, AHCTF1, ZNF695* |
| TKHS190002653 | NEC Ovary | DEL | chr1:2480183-2511683 | *PANK4, PLCH2* |

| | | | | |
|---|---|---|---|---|
| TKHS190002657 | NEC Ovary | DEL | chr1:26543162-26573361 | *MIR1976, RPS6KA1* |
| TKHS190002650 | NEC Ovary | DUP | chr1:32161947-32224121 | *DCDC2B, TXLNA, KPNA6, IQCC, TMEM234, CCDC28B, EIF3I* |
| TKHS190002650 | NEC Ovary | DUP | chr1:32274198-32651322 | *FAM229A, HDAC1, ZBTB8OS, LCK, BSDC1, RBBP4, MARCKSL1, ZBTB8A, ZBTB8B, TSSK3* |
| TKHS190002650 | NEC Ovary | DUP | chr1:32769880-35014888 | *MIR3605, TMEM35B, SMIM12, AZIN2, TMEM54, CSMD2, GJB4, HMGB4, RNF19B, A3GALT2, ZNF362, PHC2, AK2, FNDC5, GJA4, GJB3, GJB5, HPCA, CSMD2-AS1, TRIM62, KIAA1522, DLGAP3, S100PBP, LOC653160, ZSCAN20, C1orf94, YARS1, ZMYM6* |
| TKHS190002655 | NEC Cervix | DEL | chr1:33096707-33368622 | *MIR3605, AZIN2, A3GALT2, ZNF362, PHC2, TRIM62* |
| TKHS190002657 | NEC Ovary | DEL | chr1:33527196-33583830 | *CSMD2* |
| TKHS190002653 | NEC Ovary | DEL | chr1:3414560-3512128 | *MEGF6, ARHGEF16, PRDM16* |
| TKHS190002650 | NEC Ovary | DUP | chr1:35449867-35716321 | *C1orf216, NCDN, TFAP2E, PSMB2, KIAA0319L* |
| TKHS190002650 | NEC Ovary | DUP | chr1:36008690-36355433 | *COL8A2, AGO3, TRAPPC3, TEKT2, ADPRHL2, EVA1B, MAP7D1, SH3D21, STK40, THRAP3* |
| TKHS190002655 | NEC Cervix | DEL | chr1:36172464-36291546 | *MAP7D1, THRAP3* |
| TKHS190002657 | NEC Ovary | DEL | chr1:36319260-36355433 | *EVA1B, SH3D21, STK40* |
| TKHS190002646 | NEC Cervix | DUP | chr1:3634997-6701800 | *LINC01134, MIR4252, MIR4689, NA, ACOT7, PHF13, CCDC27, DFFB, ICMT, CHD5, NPHP4, LINC01777, ZBTB48, C1orf174, GPR153, SMIM1, RNF207, HES3, WRAP73, HES2, DNAJC11, AJAP1, TP73-AS1, PLEKHG5, LRRC47, RPL22, TP73, LINC01346, NOL9, TAS1R1, ESPN, KCNAB2, TNFRSF25, THAP3, CEP104, KLHL21* |
| TKHS190002650 | NEC Ovary | DUP | chr1:36463806-37483611 | *MIR4255, CSF3R, GRIK3, MRPS15, LINC01137, ZC3H12A* |
| TKHS190002650 | NEC Ovary | DUP | chr1:37989548-38016420 | *SF3A3, FHL3, UTP11* |
| TKHS190002641 | NEC Ovary | DUP | chr1:3857604-3900186 | *DFFB, C1orf174* |
| TKHS190002650 | NEC Ovary | DUP | chr1:39626507-39969568 | *PPIE, HEYL, MYCL, HPCAL4, TRIT1, OXCT2, BMP8B, NT5C1A, MFSD2A* |
| TKHS190002650 | NEC Ovary | DUP | chr1:40222574-40641925 | *ZMPSTE24, TMCO2, ZNF684, COL9A2, ZFP69, RLF, SMAP2, EXO5, ZFP69B, RIMS3* |
| TKHS190002657 | NEC Ovary | DEL | chr1:40301182-40374223 | *COL9A2, SMAP2* |
| TKHS190002641 | NEC Ovary | DUP | chr1:43272723-43313425 | *TMEM125, C1orf210, TIE1* |
| TKHS190002657 | NEC Ovary | DEL | chr1:43272723-43352772 | *TMEM125, C1orf210, MPL, TIE1* |
| TKHS190002646 | NEC Cervix | DUP | chr1:43307142-43338720 | *MPL, TIE1* |
| TKHS190002657 | NEC Ovary | DEL | chr1:43420153-43622003 | *SZT2, PTPRF, HYI* |
| TKHS190002650 | NEC Ovary | DUP | chr1:43578810-43622003 | *PTPRF* |
| TKHS190002657 | NEC Ovary | DEL | chr1:43992153-44248038 | *KLF17, CCDC24, DMAP1, SLC6A9, ERI3* |
| TKHS190002646 | NEC Cervix | DUP | chr1:44118908-44218755 | *KLF17, DMAP1* |
| TKHS190002646 | NEC Cervix | DUP | chr1:44760280-44762451 | *KIF2C* |
| TKHS190002657 | NEC Ovary | DEL | chr1:44802955-44832341 | *PLK3, BTBD19, TCTEX1D4, PTCH2* |
| TKHS190002657 | NEC Ovary | DEL | chr1:45621281-45623909 | *CCDC17* |
| TKHS190002657 | NEC Ovary | DEL | chr1:46032178-46036066 | *MAST2* |
| TKHS190002657 | NEC Ovary | DEL | chr1:46410398-46562843 | *MKNK1-AS1, DMBX1, KNCN, FAAH, FAAHP1, MKNK1* |
| TKHS190002657 | NEC Ovary | DEL | chr1:46799082-46941433 | *CYP4A11, CYP4B1, CYP4Z2P* |

| | | | | |
|---|---|---|---|---|
| TKHS190002657 | NEC Ovary | DEL | chr1:47142108-47187427 | *LINC00853, PDZK1IP1, CYP4A22* |
| TKHS190002657 | NEC Ovary | DEL | chr1:51294391-51321825 | *TTC39A* |
| TKHS190002657 | NEC Ovary | DEL | chr1:52353173-52362746 | *CC2D1B* |
| TKHS190002654 | NEC Cervix | DUP | chr1:53047858-53134429 | *PODN, SCP2, SLC1A7* |
| TKHS190002657 | NEC Ovary | DEL | chr1:54757734-54887119 | *LEXM, DHCR24, PARS2, TTC22* |
| TKHS190002657 | NEC Ovary | DEL | chr1:5863265-6142605 | *CHD5, NPHP4, KCNAB2* |
| TKHS190002641 | NEC Ovary | DUP | chr1:5863265-6701800 | *MIR4252, ACOT7, PHF13, ICMT, CHD5, NPHP4, ZBTB48, GPR153, RNF207, HES3, HES2, DNAJC11, PLEKHG5, RPL22, NOL9, TAS1R1, ESPN, KCNAB2, TNFRSF25, THAP3, KLHL21* |
| TKHS190002657 | NEC Ovary | DEL | chr1:6245110-6385666 | *ACOT7, GPR153, HES3* |
| TKHS190002646 | NEC Cervix | DUP | chr1:8945887-9129027 | *SLC2A7, SLC2A5, CA6, GPR157* |
| TKHS190002657 | NEC Ovary | DEL | chr1:8945887-9264869 | *SLC2A7, MIR34A, SLC2A5, CA6, GPR157, H6PD* |
| TKHS190002641 | NEC Ovary | DUP | chr1:9104419-9264869 | *MIR34A, GPR157, H6PD* |
| TKHS190002641 | NEC Ovary | DUP | chr1:925942-1649637 | *ACAP3, UBE2J2, PUSL1, B3GALT6, MIB2, SAMD11, MRPL20-AS1, DVL1, ATAD3C, LINC01342, TTLL10, NOC2L, SSU72, KLHL17, TMEM240, AGRN, C1QTNF12, RNF223, MIR200A, MIR200B, ANKRD65, SDF4, MXRA8, INTS11, C1orf159, AURKAIP1, MRPL20, ATAD3A, MIR429, HES4, SCNN1D, TMEM88B, FNDC10, VWA1, TNFRSF4, CPTP, CCNL2, TAS1R3, ATAD3B, PLEKHN1, PERM1, MMP23B, TNFRSF18, ISG15, CDK11B* |
| TKHS190002657 | NEC Ovary | DEL | chr1:948131-1244275 | *B3GALT6, LINC01342, TTLL10, NOC2L, KLHL17, AGRN, C1QTNF12, RNF223, MIR200A, MIR200B, SDF4, C1orf159, MIR429, HES4, TNFRSF4, PLEKHN1, PERM1, TNFRSF18, ISG15* |
| TKHS190002646 | NEC Cervix | DUP | chr1:952000-1244275 | *B3GALT6, LINC01342, TTLL10, NOC2L, KLHL17, AGRN, C1QTNF12, RNF223, MIR200A, MIR200B, SDF4, C1orf159, MIR429, HES4, TNFRSF4, PLEKHN1, PERM1, TNFRSF18, ISG15* |
| TKHS190002657 | NEC Ovary | DEL | chr1:9611753-9736042 | *TMEM201, CLSTN1, PIK3CD, PIK3CD-AS1* |
| TKHS190002653 | NEC Ovary | DEL | chr1:961293-1244497 | *B3GALT6, LINC01342, TTLL10, KLHL17, AGRN, C1QTNF12, RNF223, MIR200A, MIR200B, SDF4, C1orf159, MIR429, HES4, TNFRSF4, PLEKHN1, PERM1, TNFRSF18, ISG15* |
| TKHS190002643 | NEC Cervix | DUP | chr10:102380506-102398085 | *NFKB2, GBF1* |
| TKHS190002647 | NEC Endometrium | DUP | chr10:102472854-102475984 | *MFSD13A* |
| TKHS190002643 | NEC Cervix | DEL | chr10:122585271-122593598 | *DMBT1* |
| TKHS190002650 | NEC Ovary | DEL | chr10:129536241-129957326 | *MIR4297, EBF3, MGMT* |
| TKHS190002650 | NEC Ovary | DEL | chr10:131947530-132445779 | *DPYSL4, C10orf91, PWWP2B, JAKMIP3, STK32C, PPP2R2D, BNIP3, LRRC27* |
| TKHS190002657 | NEC Ovary | DUP | chr10:132149982-132445779 | *DPYSL4, C10orf91, PWWP2B, JAKMIP3, STK32C, LRRC27* |
| TKHS190002657 | NEC Ovary | DUP | chr10:132908468-133160569 | *LINC01168, CFAP46, ADGRA1, KNDC1* |
| TKHS190002657 | NEC Ovary | DUP | chr10:133263156-133312866 | *ADAM8, TUBGCP2, PRAP1, ZNF511* |
| TKHS190002650 | NEC Ovary | DEL | chr10:133263156-133352440 | *ADAM8, TUBGCP2, PRAP1, ZNF511, CALY* |
| TKHS190002650 | NEC Ovary | DEL | chr10:133398435-133532868 | *CYP2E1, SPRN, SCART1, MTG1* |
| TKHS190002656 | NEC Cervix | DUP | chr10:14839905-14934520 | *HSPA14, DCLRE1C, SUV39H2* |
| TKHS190002656 | NEC Cervix | DUP | chr10:21538833-21759618 | *DNAJC1, MLLT10* |
| TKHS190002656 | NEC Cervix | DUP | chr10:22316417-22389312 | *COMMD3-BMI1, COMMD3, BMI1, SPAG6* |
| TKHS190002656 | NEC Cervix | DUP | chr10:240056-356506 | *ZMYND11, DIP2C* |

| | | | | |
|---|---|---|---|---|
| TKHS190002656 | NEC Cervix | DUP | chr10:26770245-26823305 | *ABI1* |
| TKHS190002656 | NEC Cervix | DUP | chr10:27145428-27944962 | *YME1L1, RAB18, MKX, PTCHD3, LRRC37A6P, ARMC4, MASTL, ACBD5* |
| TKHS190002645 | NEC Endometrium | DEL | chr10:277325-662962 | *MIR5699, DIP2C, PRR26* |
| TKHS190002656 | NEC Cervix | DUP | chr10:32048464-32491997 | *NA, KIF5B, EPC1* |
| TKHS190002656 | NEC Cervix | DUP | chr10:35032435-35530036 | *CREM, CCNY, CUL2* |
| TKHS190002650 | NEC Ovary | DEL | chr10:44931998-45000859 | *DEPP1, C10orf25, TMEM72-AS1, TMEM72, ZNF22, RASSF4* |
| TKHS190002656 | NEC Cervix | DUP | chr10:5456087-5653050 | *NET1, CALML5, ASB13, CALML3* |
| TKHS190002650 | NEC Ovary | DEL | chr10:71615504-71682572 | *CDH23* |
| TKHS190002654 | NEC Cervix | DUP | chr10:71709098-71812890 | *C10orf105, CDH23, VSIR* |
| TKHS190002641 | NEC Ovary | DUP | chr11:116787482-116859604 | *APOA5, SIK3, APOA1, APOA4, APOC3, ZPR1* |
| TKHS190002641 | NEC Ovary | DUP | chr11:117175906-117203884 | *LOC100652768, PAFAH1B2, SIDT2, TAGLN* |
| TKHS190002646 | NEC Cervix | DUP | chr11:117179264-117192014 | *SIDT2* |
| TKHS190002641 | NEC Ovary | DUP | chr11:117504924-117839830 | *FXYD6-FXYD2, FXYD2, FXYD6, DSCAML1* |
| TKHS190002641 | NEC Ovary | DUP | chr11:118627305-118661990 | *TREH, PHLDB1* |
| TKHS190002643 | NEC Cervix | DUP | chr11:119020150-119111629 | *HYOU1, DPAGT1, SLC37A4, H2AX, HMBS, TRAPPC4, VPS11, C2CD2L* |
| TKHS190002641 | NEC Ovary | DUP | chr11:119023321-119048080 | *HYOU1, SLC37A4, TRAPPC4* |
| TKHS190002643 | NEC Cervix | DUP | chr11:119359024-119420905 | *USP2-AS1, THY1, USP2* |
| TKHS190002643 | NEC Cervix | DUP | chr11:12162156-12255750 | *MICAL2* |
| TKHS190002641 | NEC Ovary | DUP | chr11:130188869-130209823 | *ST14* |
| TKHS190002657 | NEC Ovary | DEL | chr11:130190454-130209823 | *ST14* |
| TKHS190002654 | NEC Cervix | DUP | chr11:17398339-17504697 | *USH1C, ABCC8* |
| TKHS190002643 | NEC Cervix | DUP | chr11:45847250-45870532 | *CRY2* |
| TKHS190002642 | NEC Ovary | DEL | chr11:487420-644654 | *MIR210HG, DEAF1, LRRC56, LOC143666, DRD4, LMNTD2, HRAS, IRF7, MIR210, CDHR5, PHRF1, RNH1, SCT, RASSF7, PTDSS2* |
| TKHS190002643 | NEC Cervix | DUP | chr11:61789795-61848284 | *MIR1908, FEN1, FADS1, MIR611, TMEM258, FADS2* |
| TKHS190002643 | NEC Cervix | DUP | chr11:62560069-62609696 | *EEF1G, EML3, TUT1, MTA2* |
| TKHS190002643 | NEC Cervix | DUP | chr11:62621259-62715379 | *HNRNPUL2-BSCL2, LRRN4CL, HNRNPUL2, GANAB, B3GAT3, BSCL2, GNG3, UBXN1, SNORA57, CSKMT, LBHD1, UQCC3, INTS5* |
| TKHS190002643 | NEC Cervix | DUP | chr11:62789023-62807666 | *NXF1, TMEM179B, STX5, TMEM223* |
| TKHS190002643 | NEC Cervix | DUP | chr11:64225499-64243876 | *FKBP2, DNAJC4, VEGFB, TRPT1, NUDT22* |
| TKHS190002644 | NEC Cervix | DUP | chr11:65107581-65210872 | *FAU, SPDYC, TM7SF2, VPS51, MRPL49, ZNHIT2, CAPN1, SYVN1* |
| TKHS190002643 | NEC Cervix | DUP | chr11:65114713-65133601 | *FAU, TM7SF2, MRPL49, ZNHIT2, SYVN1* |
| TKHS190002644 | NEC Cervix | DUP | chr11:65526124-65619932 | *KCNK7, ZNRD2, FAM89B, ZNRD2-AS1, EHBP1L1, PCNX3, LTBP3, MAP3K11, SCYL1* |
| TKHS190002642 | NEC Ovary | DEL | chr11:65546798-65580259 | *ZNRD2, FAM89B, ZNRD2-AS1, EHBP1L1, LTBP3* |
| TKHS190002642 | NEC Ovary | DEL | chr11:65593343-65628703 | *KCNK7, PCNX3, MAP3K11* |

| | | | | |
|---|---|---|---|---|
| TKHS190002643 | NEC Cervix | DUP | chr11:6604853-6641613 | *TPP1, ILK, TAF10, DCHS1* |
| TKHS190002643 | NEC Cervix | DUP | chr11:66565671-66593714 | *CCDC87, CTSF, CCS* |
| TKHS190002644 | NEC Cervix | DUP | chr11:66975363-67279519 | *GRK2, KDM2A, C11orf86, RHOD, SYT12* |
| TKHS190002644 | NEC Cervix | DUP | chr11:67431368-67442628 | *CORO1B, PTPRCAP, RPS6KB2* |
| TKHS190002644 | NEC Cervix | DUP | chr11:67585138-68019255 | *ALDH3B1, ALDH3B2, NUDT8, GSTP1, TBX10, DOC2GP, NDUFV1, FAM86C2P, UNC93B1, ACY3* |
| TKHS190002641 | NEC Ovary | DUP | chr11:695609-726981 | *DEAF1, TMEM80, EPS8L2* |
| TKHS190002643 | NEC Cervix | DUP | chr11:72757767-72780450 | *STARD10, ARAP1* |
| TKHS190002642 | NEC Ovary | DEL | chr11:774008-842508 | *LOC171391, CRACR2B, GATD1, CEND1, POLR2L, PIDD1, PNPLA2, RPLP2, SNORA52, SLC25A22, CD151* |
| TKHS190002643 | NEC Cervix | DUP | chr11:8683202-8917466 | *AKIP1, RPL27A, SNORA3A, DENND2B, SNORA3B* |
| TKHS190002648 | NEC Cervix | DUP | chr12:103666291-103677611 | *STAB2* |
| TKHS190002653 | NEC Ovary | DUP | chr12:108560765-108852678 | *ISCU, CORO1C, TMEM119, SSH1, SELPLG, SART3* |
| TKHS190002653 | NEC Ovary | DUP | chr12:109084953-109279930 | *ALKBH2, FOXN4, ACACB, UNG, USP30* |
| TKHS190002648 | NEC Cervix | DUP | chr12:109174132-109281799 | *FOXN4, ACACB* |
| TKHS190002648 | NEC Cervix | DUP | chr12:109579802-109855769 | *MIR4497, MVK, GLTP, TRPV4, FAM222A, FAM222A-AS1* |
| TKHS190002653 | NEC Ovary | DUP | chr12:109596426-109904147 | *MIR4497, MVK, GLTP, TRPV4, TCHP, FAM222A, FAM222A-AS1* |
| TKHS190002653 | NEC Ovary | DUP | chr12:111293446-111448302 | *SH2B3, PHETA1, CUX2* |
| TKHS190002657 | NEC Ovary | DEL | chr12:111304210-111363463 | *PHETA1, CUX2* |
| TKHS190002648 | NEC Cervix | DUP | chr12:113093539-113398846 | *SDS, IQCD, DTX1, PLBD2, CFAP73, TPCN1, DDX54, SLC8B1, RASAL1, RITA1* |
| TKHS190002657 | NEC Ovary | DEL | chr12:113162932-113176935 | *DDX54* |
| TKHS190002643 | NEC Cervix | DEL | chr12:113165644-113174936 | *DDX54* |
| TKHS190002653 | NEC Ovary | DUP | chr12:113172353-113226964 | *IQCD, TPCN1, DDX54, RITA1* |
| TKHS190002653 | NEC Ovary | DUP | chr12:117186477-117531013 | *FBXO21, KSR2, NOS1* |
| TKHS190002648 | NEC Cervix | DUP | chr12:120093000-120160023 | *MIR4498, GCN1, RAB35, BICDL1* |
| TKHS190002653 | NEC Ovary | DUP | chr12:120097245-120457179 | *PXN-AS1, MIR4498, GCN1, RAB35, COX6A1, SIRT4, GATC, MSI1, TRIAP1, PLA2G1B, PXN, RPLP0* |
| TKHS190002653 | NEC Ovary | DUP | chr12:120666875-120737988 | *MIR4700, ACADS, UNC119B, CABP1, MLEC* |
| TKHS190002648 | NEC Cervix | DUP | chr12:120687297-120769059 | *MIR4700, SPPL3, ACADS, UNC119B, MLEC* |
| TKHS190002648 | NEC Cervix | DUP | chr12:120994164-121005093 | *C12orf43, HNF1A* |
| TKHS190002648 | NEC Cervix | DUP | chr12:121221913-121250034 | *CAMKK2, P2RX4* |
| TKHS190002653 | NEC Ovary | DUP | chr12:121440816-121899889 | *TMEM120B, SETD1B, MORN3, HPD, LINC01089, RHOF, PSMD9, KDM2B, ORAI1* |
| TKHS190002648 | NEC Cervix | DUP | chr12:121775062-121849076 | *TMEM120B, SETD1B, HPD, LINC01089, RHOF* |
| TKHS190002653 | NEC Ovary | DUP | chr12:121968006-122207388 | *WDR66, MLXIP, LRRC43, IL31, BCL7A, B3GNT4* |
| TKHS190002648 | NEC Cervix | DUP | chr12:122022092-122184779 | *MLXIP, LRRC43, IL31, BCL7A* |
| TKHS190002648 | NEC Cervix | DUP | chr12:122848796-122858435 | *HIP1R* |

| | | | | |
|---|---|---|---|---|
| TKHS190002653 | NEC Ovary | DUP | chr12:122858838-122982586 | *ABCB9, ARL6IP4, OGFOD2, VPS37B, HIP1R* |
| TKHS190002648 | NEC Cervix | DUP | chr12:122976253-122986829 | *ARL6IP4, PITPNM2, OGFOD2* |
| TKHS190002657 | NEC Ovary | DEL | chr12:124340602-124437996 | *NCOR2* |
| TKHS190002648 | NEC Cervix | DUP | chr12:124343005-125125024 | *MIR5188, BRI3BP, DHX37, AACS, UBC, SCARB1, NCOR2* |
| TKHS190002653 | NEC Ovary | DUP | chr12:124344597-125125024 | *MIR5188, BRI3BP, DHX37, AACS, UBC, SCARB1, NCOR2* |
| TKHS190002653 | NEC Ovary | DUP | chr12:128543957-129337817 | *TMEM132D, SLC15A4, GLT1D1, TMEM132C* |
| TKHS190002657 | NEC Ovary | DEL | chr12:131843997-131929796 | *MMP17, PUS1, ULK1* |
| TKHS190002648 | NEC Cervix | DUP | chr12:132017535-132675517 | *LOC100130238, LRCOL1, P2RX2, DDX51, EP400P1, GALNT9, POLE, EP400, FBRSL1, SNORA49, NOC4L* |
| TKHS190002653 | NEC Ovary | DUP | chr12:132681138-132822128 | *PGAM5, ANKLE2, GOLGA3, POLE, PXMP2* |
| TKHS190002642 | NEC Ovary | DUP | chr12:157528-226514 | *IQSEC3, SLC6A12, SLC6A13* |
| TKHS190002642 | NEC Ovary | DUP | chr12:2504436-2691199 | *CACNA1C-AS1, CACNA1C* |
| TKHS190002653 | NEC Ovary | DUP | chr12:2504436-3696999 | *NA, CACNA1C-AS1, TSPAN9, FKBP4, FOXM1, ITFG2-AS1, ITFG2, PRMT8, TEAD4, TULP3, CACNA1C, RHNO1, NRIP2, CRACR2A* |
| TKHS190002653 | NEC Ovary | DUP | chr12:27766665-27969516 | *MANSC4, PTHLH, KLHL42* |
| TKHS190002656 | NEC Cervix | DEL | chr12:2820086-2827677 | *ITFG2, NRIP2* |
| TKHS190002642 | NEC Ovary | DUP | chr12:3021844-3638454 | *TSPAN9, PRMT8, TEAD4, CRACR2A* |
| TKHS190002653 | NEC Ovary | DUP | chr12:47717518-47791706 | *RAPGEF3, HDAC7, SLC48A1, ENDOU* |
| TKHS190002648 | NEC Cervix | DUP | chr12:47749390-47855801 | *RAPGEF3, HDAC7, SLC48A1, VDR* |
| TKHS190002656 | NEC Cervix | DEL | chr12:48716515-48778290 | *MIR4701, ADCY6, TEX49, CCNT1* |
| TKHS190002648 | NEC Cervix | DUP | chr12:48716515-48830692 | *MIR4701, ADCY6, TEX49, CACNB3, CCNT1, DDX23* |
| TKHS190002648 | NEC Cervix | DUP | chr12:49024578-49054751 | *KMT2D* |
| TKHS190002653 | NEC Ovary | DUP | chr12:49024578-49054751 | *KMT2D* |
| TKHS190002656 | NEC Cervix | DEL | chr12:49030893-49054140 | *KMT2D* |
| TKHS190002648 | NEC Cervix | DUP | chr12:49069734-49632880 | *TROAP, LOC100335030, TUBA1B, MCRS1, RHEBL1, KCNH3, PRPF40B, C1QL4, DHH, LMBR1L, PRPH, SPATS2, TUBA1A, DNAJC22, FAM186B, TUBA1C* |
| TKHS190002653 | NEC Ovary | DUP | chr12:49097672-49351502 | *TROAP, TUBA1B, C1QL4, LMBR1L, PRPH, TUBA1A, DNAJC22, TUBA1C* |
| TKHS190002653 | NEC Ovary | DUP | chr12:49543915-49634092 | *MCRS1, KCNH3, PRPF40B, FAM186B* |
| TKHS190002656 | NEC Cervix | DEL | chr12:49599469-49666207 | *PRPF40B, FAM186B, FMNL3* |
| TKHS190002648 | NEC Cervix | DUP | chr12:51993985-52077190 | *GRASP, NR4A1, ATG101, ACVR1B* |
| TKHS190002653 | NEC Ovary | DUP | chr12:51993985-52185587 | *KRT80, GRASP, NR4A1, ATG101, ACVR1B* |
| TKHS190002656 | NEC Cervix | DEL | chr12:52013687-52287322 | *KRT80, GRASP, C12orf80, LINC00592, NR4A1, KRT7, KRT81, KRT86, ATG101* |
| TKHS190002656 | NEC Cervix | DEL | chr12:53052455-53443827 | *SP7, TNS2, AMHR2, LOC283335, ZNF740, IGFBP6, ITGB7, PCBP2, CSAD, PFDN5, PRR13, RARG, C12orf10, SP1, AAAS, SOAT2, SPRYD3, MFSD5, ESPL1* |
| TKHS190002648 | NEC Cervix | DUP | chr12:53097718-53161389 | *IGFBP6, CSAD, SOAT2* |
| TKHS190002648 | NEC Cervix | DUP | chr12:53285924-53293474 | *ESPL1* |

| | | | | |
|---|---|---|---|---|
| TKHS190002642 | NEC Ovary | DUP | chr12:53285924-53293474 | *ESPL1* |
| TKHS190002653 | NEC Ovary | DUP | chr12:53320565-53442733 | *SP7, AMHR2, PCBP2, PRR13, SP1, AAAS* |
| TKHS190002653 | NEC Ovary | DUP | chr12:53485056-53524761 | *LOC100652999, ATF7, TARBP2, MAP3K12, NPFF* |
| TKHS190002653 | NEC Ovary | DUP | chr12:53722025-54011381 | *HOTAIR, HOXC13-AS, HOXC8, HOXC9, HOXC10, HOXC11, HOXC12, HOXC13, MIR196A2, CALCOCO1* |
| TKHS190002653 | NEC Ovary | DUP | chr12:54362661-54419198 | *ZNF385A, ITGA5, GPR84* |
| TKHS190002656 | NEC Cervix | DEL | chr12:54375844-54408235 | *ZNF385A, ITGA5* |
| TKHS190002642 | NEC Ovary | DUP | chr12:55636892-55699989 | *OR10P1, METTL7B, ITGA7* |
| TKHS190002653 | NEC Ovary | DUP | chr12:55716933-55749882 | *BLOC1S1-RDH5, GDF11, BLOC1S1, RDH5, CD63* |
| TKHS190002656 | NEC Cervix | DEL | chr12:55938915-55958607 | *DGKA, PMEL* |
| TKHS190002653 | NEC Ovary | DUP | chr12:55967857-56035331 | *CDK2, RAB5B, IKZF4, SUOX* |
| TKHS190002656 | NEC Cervix | DEL | chr12:56116788-56161415 | *MYL6B, ESYT1, MYL6, RPL41, ZC3H10* |
| TKHS190002656 | NEC Cervix | DEL | chr12:56224857-56254720 | *ANKRD52, SLC39A5, NABP2* |
| TKHS190002653 | NEC Ovary | DUP | chr12:57091125-57192970 | *LRP1, NAB2, STAT6* |
| TKHS190002656 | NEC Cervix | DEL | chr12:57091125-57269994 | *MIR1228, NXPH4, R3HDM2, STAC3, LRP1, NAB2, NDUFA4L2, SHMT2, STAT6* |
| TKHS190002648 | NEC Cervix | DUP | chr12:57141374-57232843 | *MIR1228, NXPH4, LRP1, SHMT2* |
| TKHS190002657 | NEC Ovary | DEL | chr12:57169140-57233662 | *MIR1228, NXPH4, LRP1, SHMT2* |
| TKHS190002653 | NEC Ovary | DUP | chr12:57221307-57236681 | *NXPH4, NDUFA4L2, SHMT2* |
| TKHS190002642 | NEC Ovary | DUP | chr12:57600324-57827123 | *AGAP2-AS1, TSFM, CTDSP2, CDK4, AVIL, OS9, ARHGEF25, AGAP2, CYP27B1, DTX3, B4GALNT1, EEF1AKMT3, MIR26A2, METTL1, TSPAN31, SLC26A10, PIP4K2C, MARCHF9* |
| TKHS190002648 | NEC Cervix | DUP | chr12:5967486-6075551 | *VWF* |
| TKHS190002653 | NEC Ovary | DUP | chr12:5967486-6493136 | *LTBR, MRPL51, TAPBPL, PLEKHG6, SCNN1A, CD27-AS1, VAMP1, TNFRSF1A, VWF, CD9, CD27* |
| TKHS190002653 | NEC Ovary | DUP | chr12:6526891-6563514 | *IFFO1, GAPDH, NOP2, NCAPD2* |
| TKHS190002653 | NEC Ovary | DUP | chr12:6695426-6970543 | *LRRC23, P3H3, C12orf57, PHB2, DSTNP2, ATN1, PIANP, ENO2, GPR162, GNB3, RPL13P5, LAG3, MIR141, MIR200C, COPS7A, PTMS, PTPN6, SCARNA12, TPI1, USP5, MLF2, CDCA3, SPSB2, CD4* |
| TKHS190002656 | NEC Cervix | DEL | chr12:6863830-6937561 | *LRRC23, DSTNP2, ATN1, ENO2, RPL13P5, TPI1, USP5, SPSB2* |
| TKHS190002656 | NEC Cervix | DEL | chr12:7067789-7150689 | *C1RL-AS1, C1RL, C1R, C1S, RBP5, CLSTN3* |
| TKHS190002653 | NEC Ovary | DUP | chr12:7090056-7190569 | *C1RL-AS1, C1RL, PEX5, C1R, RBP5, CLSTN3* |
| TKHS190002653 | NEC Ovary | DUP | chr12:99146-237290 | *IQSEC3, LOC574538, SLC6A12, SLC6A13* |
| TKHS190002642 | NEC Ovary | DUP | chr13:101057939-101083804 | *NALCN-AS1, NALCN* |
| TKHS190002657 | NEC Ovary | DUP | chr14:100123539-100343387 | *SLC25A29, DEGS2, SLC25A47, MIR345, EVL, WARS1, YY1* |
| TKHS190002642 | NEC Ovary | DUP | chr14:102039412-102050563 | *DYNC1H1* |
| TKHS190002641 | NEC Ovary | DEL | chr14:102039412-102229657 | *DYNC1H1, HSP90AA1, MOK, WDR20* |
| TKHS190002642 | NEC Ovary | DUP | chr14:102193463-102229657 | *MOK, WDR20* |
| TKHS190002641 | NEC Ovary | DEL | chr14:102939610-102964650 | *CDC42BPB* |

| | | | | |
|---|---|---|---|---|
| TKHS190002641 | NEC Ovary | DEL | chr14:103502882-103574170 | *CKB, TRMT61A, KLC1, MARK3, COA8, BAG5* |
| TKHS190002641 | NEC Ovary | DEL | chr14:104092633-105886061 | *ZBTB42, MIR203B, MIR4710, SIVA1, AHNAK2, CLBA1, PLD4, ADSS1, CRIP1, CRIP2, LINC00638, ELK2AP, AKT1, PACS2, NUDT14, KIF26A, CEP170B, TEDC1, BRF1, GPR132, JAG2, ASPG, TMEM179, C14orf180, MIR203A, CDCA4, INF2, TEX22, TMEM121, BTBD6, MTA1* |
| TKHS190002642 | NEC Ovary | DUP | chr14:104739655-104746385 | *ADSS1* |
| TKHS190002642 | NEC Ovary | DUP | chr14:104930742-105055420 | *AHNAK2, CLBA1, PLD4, GPR132, CDCA4* |
| TKHS190002647 | NEC Endometrium | DUP | chr14:105143482-105173261 | *NUDT14, JAG2* |
| TKHS190002657 | NEC Ovary | DUP | chr14:105173021-105217800 | *NUDT14, BRF1* |
| TKHS190002642 | NEC Ovary | DUP | *chr14:105838401-105888578* | |
| TKHS190002657 | NEC Ovary | DUP | *chr14:105845375-105886061* | |
| TKHS190002655 | NEC Cervix | DUP | chr14:106088122-106324688 | *ELK2AP* |
| TKHS190002653 | NEC Ovary | DEL | chr14:20992092-20995951 | *METTL17* |
| TKHS190002642 | NEC Ovary | DUP | chr14:23394068-23433732 | *MIR208B, MYH6, MYH7* |
| TKHS190002657 | NEC Ovary | DUP | chr14:24094766-24143489 | *FITM1, NRL, EMC9, PCK2, PSME1, PSME2, DCAF11* |
| TKHS190002657 | NEC Ovary | DUP | chr14:24147976-24159960 | *RNF31* |
| TKHS190002657 | NEC Ovary | DUP | chr14:24181711-24206723 | *TM9SF1, TSSK4, IPO4* |
| TKHS190002657 | NEC Ovary | DUP | chr14:24237245-24255517 | *TINF2, GMPR2, TGM1* |
| TKHS190002655 | NEC Cervix | DUP | chr14:24573637-24634160 | *CTSG, GZMH, GZMB* |
| TKHS190002657 | NEC Ovary | DUP | chr14:64727593-64742455 | *PLEKHG3* |
| TKHS190002641 | NEC Ovary | DEL | chr14:73251664-73265507 | *PAPLN* |
| TKHS190002657 | NEC Ovary | DUP | chr14:74494031-74505174 | *ISCA2, LTBP2* |
| TKHS190002657 | NEC Ovary | DUP | chr14:74521911-74586118 | *LTBP2* |
| TKHS190002657 | NEC Ovary | DUP | chr14:74906549-75009188 | *PGF, RPS6KL1, EIF2B2* |
| TKHS190002653 | NEC Ovary | DUP | chr15:29101599-29196564 | *FAM189A1, APBA2* |
| TKHS190002655 | NEC Cervix | DUP | chr15:40779545-40812902 | *DNAJC17, ZFYVE19* |
| TKHS190002650 | NEC Ovary | DEL | chr15:40779545-40812902 | *DNAJC17, ZFYVE19* |
| TKHS190002647 | NEC Endometrium | DUP | chr15:41512115-41513709 | *LTK* |
| TKHS190002650 | NEC Ovary | DEL | chr15:42675872-42719574 | *STARD9* |
| TKHS190002655 | NEC Cervix | DUP | chr15:42675872-42725251 | *CDAN1, STARD9* |
| TKHS190002655 | NEC Cervix | DUP | chr15:43601396-43614476 | *STRC* |
| TKHS190002657 | NEC Ovary | DUP | chr15:92444636-92949080 | *LINC00930, MIR3175, LINC01578, CHD2, C15orf32, FAM174B, ASB9P1, ST8SIA2* |
| TKHS190002647 | NEC Endometrium | DUP | chr16:1448681-1459187 | *CLCN7* |
| TKHS190002641 | NEC Ovary | DEL | chr16:1454411-1494616 | *CLCN7, PTX4, TELO2* |
| TKHS190002641 | NEC Ovary | DEL | chr16:15016129-15033399 | *PDXDC1* |
| TKHS190002653 | NEC Ovary | DUP | chr16:1520602-1542076 | *TMEM204, IFT140* |

| | | | | |
|---|---|---|---|---|
| TKHS190002641 | NEC Ovary | DEL | chr16:1523518-1674045 | *CRAMP1, TMEM204, IFT140* |
| TKHS190002655 | NEC Cervix | DUP | chr16:16033109-16203613 | *ABCC6, ABCC1* |
| TKHS190002645 | NEC Endometrium | DEL | chr16:16090405-16141281 | *ABCC1* |
| TKHS190002641 | NEC Ovary | DEL | chr16:1944837-1963839 | *SNORA64, NDUFB10, SNORA10, RPL3L, RPS2* |
| TKHS190002657 | NEC Ovary | DUP | chr16:1974181-1981179 | *TBL3, NOXO1* |
| TKHS190002646 | NEC Cervix | DEL | chr16:1974181-1986028 | *TBL3, NOXO1, GFER* |
| TKHS190002657 | NEC Ovary | DUP | chr16:2037757-2057182 | *NTHL1, TSC2, SLC9A3R2* |
| TKHS190002657 | NEC Ovary | DUP | chr16:2082436-2094210 | *MIR1225, PKD1, TSC2* |
| TKHS190002646 | NEC Cervix | DEL | chr16:2206086-2235572 | *E4F1, BRICD5, PGP, MLST8* |
| TKHS190002655 | NEC Cervix | DUP | chr16:22146236-22308225 | *VWA3A, EEF2K, POLR3E* |
| TKHS190002645 | NEC Endometrium | DEL | chr16:22184181-22264880 | *EEF2K* |
| TKHS190002657 | NEC Ovary | DUP | chr16:2228292-2235572 | *E4F1* |
| TKHS190002643 | NEC Cervix | DEL | chr16:23689238-23694927 | *ERN2, PLK1* |
| TKHS190002646 | NEC Cervix | DEL | chr16:2460283-2531073 | *NTN3, AMDHD2, ATP6V0C, TBC1D24, CEMP1, TEDC2* |
| TKHS190002653 | NEC Ovary | DUP | chr16:2784631-2786547 | *PRSS33* |
| TKHS190002653 | NEC Ovary | DUP | chr16:28974780-28986714 | *LAT, SPNS1* |
| TKHS190002656 | NEC Cervix | DEL | chr16:3024327-3026781 | *THOC6* |
| TKHS190002653 | NEC Ovary | DUP | chr16:30364900-30379534 | *SEPTIN1, ZNF48, TBC1D10B, MYLPF* |
| TKHS190002653 | NEC Ovary | DUP | chr16:30526326-30653103 | *ZNF689, ZNF785, ZNF688, ZNF747, PRR14, ZNF768, ZNF764* |
| TKHS190002650 | NEC Ovary | DEL | chr16:3067377-3205185 | *OR1F1, ZNF205, ZNF213, ZNF205-AS1, ZSCAN10, IL32* |
| TKHS190002657 | NEC Ovary | DUP | chr16:30756355-30761335 | *PHKG2, CCDC189* |
| TKHS190002655 | NEC Cervix | DUP | chr16:31277967-31325404 | *ITGAM* |
| TKHS190002650 | NEC Ovary | DEL | chr16:3355941-4007558 | *MTRNR2L4, TRAP1, ADCY9, CREBBP, ZNF597, DNASE1, NLRC3, CLUAP1, OR2C1, ZSCAN32, C16orf90, ZNF174, NAA60, SLX4* |
| TKHS190002657 | NEC Ovary | DUP | chr16:3563009-3657953 | *DNASE1, NLRC3, SLX4* |
| TKHS190002655 | NEC Cervix | DUP | chr16:3657720-3740549 | *TRAP1, CREBBP, DNASE1* |
| TKHS190002653 | NEC Ovary | DUP | chr16:4261779-4395339 | *CORO7-PAM16, VASN, PAM16, TFAP4, CORO7, GLIS2* |
| TKHS190002653 | NEC Ovary | DUP | chr16:4651963-4698062 | *ANKS3, MGRN1, NUDT16L1* |
| TKHS190002644 | NEC Cervix | DEL | chr16:50292676-50314406 | *ADCY7* |
| TKHS190002641 | NEC Ovary | DEL | chr16:50292676-50320030 | *ADCY7, BRD7* |
| TKHS190002653 | NEC Ovary | DUP | chr16:50548554-50674074 | *SNX20, NKD1* |
| TKHS190002641 | NEC Ovary | DEL | chr16:50621602-50712374 | *SNX20, NOD2, NKD1* |
| TKHS190002641 | NEC Ovary | DEL | chr16:55777-93325 | *MPG, RHBDF1, SNRNP25, NPRL3* |
| TKHS190002653 | NEC Ovary | DUP | chr16:56519746-56748426 | *MT1DP, MT1A, MT1B, MT1E, MT1F, MT1G, MT1H, MT1JP, MT1M, MT1L, MT1X, MT2A, MT3, BBS2, MT1IP, MT4, NUP93* |

| | | | | |
|---|---|---|---|---|
| TKHS190002644 | NEC Cervix | DEL | chr16:57680265-57724835 | *ADGRG3, DRC7* |
| TKHS190002644 | NEC Cervix | DEL | chr16:58487756-58506473 | *NDRG4* |
| TKHS190002657 | NEC Ovary | DUP | chr16:62538-85777 | *MPG, RHBDF1, NPRL3* |
| TKHS190002657 | NEC Ovary | DUP | chr16:665950-667782 | *WDR90* |
| TKHS190002642 | NEC Ovary | DUP | chr16:66847030-66912580 | *CDH16, PDP2, CA7* |
| TKHS190002642 | NEC Ovary | DUP | chr16:66940222-66988830 | *CES3, CES4A, CES2* |
| TKHS190002642 | NEC Ovary | DUP | chr16:67228355-67294171 | *KCTD19, PLEKHG4, TMEM208, FHOD1, SLC9A5* |
| TKHS190002641 | NEC Ovary | DEL | chr16:67877217-67931852 | *NRN1L, CTRL, EDC4, PSKH1, PSMB10* |
| TKHS190002642 | NEC Ovary | DUP | chr16:67877217-67931852 | *NRN1L, CTRL, EDC4, PSKH1, PSMB10* |
| TKHS190002657 | NEC Ovary | DUP | chr16:67879587-67930318 | *NRN1L, CTRL, EDC4, PSKH1* |
| TKHS190002646 | NEC Cervix | DEL | chr16:681438-697405 | *STUB1, FBXL16, JMJD8, WDR24* |
| TKHS190002650 | NEC Ovary | DEL | chr16:681793-717477 | *STUB1, FBXL16, JMJD8, METRN, WDR24* |
| TKHS190002657 | NEC Ovary | DUP | chr16:681793-730655 | *STUB1, CCDC78, FBXL16, JMJD8, CIAO3, ANTKMT, METRN, WDR24, HAGHL* |
| TKHS190002641 | NEC Ovary | DEL | chr16:69929448-69939940 | *WWP2, MIR140* |
| TKHS190002642 | NEC Ovary | DUP | chr16:69929448-69939940 | *WWP2, MIR140* |
| TKHS190002641 | NEC Ovary | DEL | chr16:70676881-70698811 | *VAC14, MTSS2* |
| TKHS190002641 | NEC Ovary | DEL | chr16:89722531-89739553 | *FANCA, ZNF276* |
| TKHS190002641 | NEC Ovary | DEL | chr16:89895990-89932679 | *TUBB3, TCF25, MC1R* |
| TKHS190002657 | NEC Ovary | DUP | chr17:1056035-1058867 | *ABR* |
| TKHS190002642 | NEC Ovary | DEL | chr17:1478087-1492487 | *MYO1C* |
| TKHS190002643 | NEC Cervix | DUP | chr17:1771029-1801024 | *SMYD4, SERPINF1* |
| TKHS190002643 | NEC Cervix | DUP | chr17:19341238-19362721 | *MIR1180, B9D1* |
| TKHS190002654 | NEC Cervix | DUP | chr17:19384462-19416175 | *MFAP4, RNF112* |
| TKHS190002654 | NEC Cervix | DUP | chr17:21287991-21304553 | *MAP2K3* |
| TKHS190002642 | NEC Ovary | DEL | chr17:2692571-2704564 | *CLUH* |
| TKHS190002650 | NEC Ovary | DEL | chr17:28554220-28575532 | *ALDOC, PIGS* |
| TKHS190002643 | NEC Cervix | DUP | chr17:28710871-28741987 | *NA, TLCD1, PROCA1, SNORD4B, SNORD4A, SNORD42B, SNORD42A, NEK8, RPL23A, RAB34* |
| TKHS190002644 | NEC Cervix | DUP | chr17:28735240-28749577 | *NEK8, TRAF4* |
| TKHS190002655 | NEC Cervix | DUP | chr17:35983801-36090058 | *CCL15-CCL14, CCL3, CCL14, CCL15, CCL18, CCL23* |
| TKHS190002650 | NEC Ovary | DEL | chr17:3953347-4034292 | *ZZEF1, ATP2A3* |
| TKHS190002657 | NEC Ovary | DUP | chr17:40021955-40031620 | *SNORD124, MED24* |
| TKHS190002643 | NEC Cervix | DUP | chr17:40035722-40097403 | *THRA, NR1D1, MED24* |
| TKHS190002647 | NEC Endometrium | DEL | chr17:42552934-42565047 | *HSD17B1, COASY* |

| | | | | |
|---|---|---|---|---|
| TKHS190002641 | NEC Ovary | DUP | chr17:4718612-4796296 | *VMO1, GLTPD2, MED11, ARRB2, PSMB6, CXCL16, ZMYND15, TM4SF5* |
| TKHS190002650 | NEC Ovary | DEL | chr17:4720393-4887790 | *VMO1, GLTPD2, MED11, ARRB2, MINK1, PLD2, PSMB6, CXCL16, ZMYND15, TM4SF5* |
| TKHS190002641 | NEC Ovary | DUP | chr17:4893286-4902494 | *C17orf107, CHRNE, MINK1* |
| TKHS190002650 | NEC Ovary | DEL | chr17:4902217-4981831 | *C17orf107, CHRNE, ENO3, CAMTA2, RNF167, GP1BA, PFN1, SLC25A11, SPAG7* |
| TKHS190002643 | NEC Cervix | DUP | chr17:4938154-4946844 | *RNF167, PFN1, SLC25A11* |
| TKHS190002657 | NEC Ovary | DUP | chr17:4952025-4960547 | *ENO3, SPAG7* |
| TKHS190002643 | NEC Cervix | DUP | chr17:4960022-4981359 | *CAMTA2, SPAG7* |
| TKHS190002643 | NEC Cervix | DUP | chr17:50186317-50199590 | *COL1A1* |
| TKHS190002644 | NEC Cervix | DUP | chr17:62666757-62681937 | *MRC2* |
| TKHS190002644 | NEC Cervix | DUP | chr17:63434402-63479912 | *CYB561, ACE* |
| TKHS190002644 | NEC Cervix | DUP | chr17:63824637-63838681 | *FTSJ3, PSMC5, SMARCD2* |
| TKHS190002644 | NEC Cervix | DUP | chr17:64001733-64160933 | *ERN1, ICAM2, TEX2, SNORA50C, SNORD104, PRR29* |
| TKHS190002643 | NEC Cervix | DUP | chr17:7013666-7017014 | *MIR497HG, RNASEK-C17orf49* |
| TKHS190002650 | NEC Ovary | DEL | chr17:7015062-7076152 | *MIR497HG, RNASEK-C17orf49, CLEC10A, SLC16A11, SLC16A13, BCL6B, MIR497* |
| TKHS190002641 | NEC Ovary | DUP | chr17:7015774-7043585 | *MIR497HG, RNASEK-C17orf49, SLC16A11, SLC16A13, BCL6B, MIR497* |
| TKHS190002655 | NEC Cervix | DUP | chr17:7075360-7076987 | *CLEC10A* |
| TKHS190002642 | NEC Ovary | DEL | chr17:7173659-7203593 | *DLG4, ASGR1* |
| TKHS190002641 | NEC Ovary | DUP | chr17:7173659-7236767 | *DLG4, DVL2, ACADVL, ASGR1, MIR324, PHF23* |
| TKHS190002654 | NEC Cervix | DUP | chr17:7193483-7194495 | *DLG4* |
| TKHS190002650 | NEC Ovary | DEL | chr17:7193483-7236767 | *DLG4, DVL2, ACADVL, MIR324, PHF23* |
| TKHS190002643 | NEC Cervix | DUP | chr17:7220464-7224552 | *ACADVL, MIR324* |
| TKHS190002643 | NEC Cervix | DUP | chr17:7229378-7244635 | *GABARAP, DVL2, CTDNEP1, PHF23* |
| TKHS190002641 | NEC Ovary | DUP | chr17:7252918-7437506 | *TMEM256-PLSCR3, CLDN7, KCTD11, EIF5A, SPEM2, ELP5, TMEM256, TMEM102, GPS2, TMEM95, SPEM1, YBX2, PLSCR3, NLGN2, SLC2A4, NEURL4, TNK1, ACAP1* |
| TKHS190002643 | NEC Cervix | DUP | chr17:7291093-7314597 | *EIF5A, GPS2, YBX2, NEURL4* |
| TKHS190002641 | NEC Ovary | DUP | chr17:7455794-7564864 | *SENP3-EIF4A1, CHRNB1, SENP3, TNFSF12-TNFSF13, POLR2A, ZBTB4, SLC35G6, TNFSF13, TNFSF12* |
| TKHS190002641 | NEC Ovary | DUP | chr17:74729256-74771858 | *MIR3615, NAT9, RAB37, SLC9A3R1* |
| TKHS190002644 | NEC Cervix | DUP | chr17:74744308-74788870 | *MIR3615, NAT9, RAB37, TMEM104, SLC9A3R1* |
| TKHS190002650 | NEC Ovary | DEL | chr17:7481988-7572864 | *SENP3-EIF4A1, SENP3, TNFSF12-TNFSF13, POLR2A, ZBTB4, SLC35G6, TNFSF13, TNFSF12* |
| TKHS190002641 | NEC Ovary | DUP | chr17:74872804-74964632 | *USH1G, FDXR, FADS6, HID1, OTOP3, OTOP2* |
| TKHS190002644 | NEC Cervix | DUP | chr17:74954138-75021079 | *HID1, CDR2L, MRPL58* |
| TKHS190002644 | NEC Cervix | DUP | chr17:75238914-75614360 | *LOC100287042, MIR3678, GGA3, TSEN54, GRB2, LLGL2, MRPS7, MIF4GD, CASKIN2, SLC25A19, MYO15B, TMEM94* |
| TKHS190002641 | NEC Ovary | DUP | chr17:75485428-75651265 | *SMIM6, TSEN54, LLGL2, CASKIN2, SMIM5, MYO15B, RECQL5, TMEM94* |
| TKHS190002653 | NEC Ovary | DEL | chr17:75491691-75504693 | *CASKIN2, TMEM94* |

| | | | | |
|---|---|---|---|---|
| TKHS190002643 | NEC Cervix | DUP | chr17:7550120-7587709 | *SENP3-EIF4A1, SENP3, SNORA67, TNFSF12-TNFSF13, SNORA48, SNORD10, TNFSF13, TNFSF12, MPDU1, CD68* |
| TKHS190002644 | NEC Cervix | DUP | chr17:75671816-75816912 | *MIR4738, GALK1, SAP30BP, H3-3B, ITGB4, UNK* |
| TKHS190002654 | NEC Cervix | DEL | chr17:75706008-75752356 | *SAP30BP, ITGB4* |
| TKHS190002641 | NEC Ovary | DUP | chr17:75752174-75828983 | *MIR4738, UNC13D, GALK1, H3-3B, ITGB4, UNK* |
| TKHS190002641 | NEC Ovary | DUP | chr17:75874208-75915139 | *TRIM65, MRPL38, FBF1, TRIM47* |
| TKHS190002644 | NEC Cervix | DUP | chr17:75949495-76002612 | *CDK3, ACOX1* |
| TKHS190002644 | NEC Cervix | DUP | chr17:76079439-76280953 | *RNF157-AS1, RNF157, FOXJ1, EXOC7, UBALD2, ZACN, QRICH2* |
| TKHS190002641 | NEC Ovary | DUP | chr17:76097796-76162002 | *RNF157-AS1, RNF157, FOXJ1, EXOC7* |
| TKHS190002641 | NEC Ovary | DUP | chr17:76468633-76540215 | *CYGB, AANAT, PRCD, RHBDF2* |
| TKHS190002650 | NEC Ovary | DEL | chr17:7661939-7703486 | *WRAP53, TP53* |
| TKHS190002644 | NEC Cervix | DUP | chr17:76625819-76643638 | *ST6GALNAC1* |
| TKHS190002644 | NEC Cervix | DUP | chr17:78122605-78205753 | *C17orf99, TMC6, AFMID, TMC8, TK1, SYNGR2* |
| TKHS190002641 | NEC Ovary | DUP | chr17:7831390-7858824 | *CYB5D1, DNAH2, KDM6B, NAA38, TMEM88* |
| TKHS190002642 | NEC Ovary | DEL | chr17:7845555-7855714 | *KDM6B, TMEM88* |
| TKHS190002643 | NEC Cervix | DUP | chr17:7930606-7934222 | *CNTROB, TRAPPC1* |
| TKHS190002644 | NEC Cervix | DUP | chr17:80140008-80201870 | *CARD14, EIF4A3* |
| TKHS190002644 | NEC Cervix | DUP | chr17:80251329-80415821 | *RNF213-AS1, SLC26A11, ENDOV, RNF213* |
| TKHS190002641 | NEC Ovary | DUP | chr17:8073657-8150706 | *MIR4314, ALOX12B, PER1, ALOXE3, HES7* |
| TKHS190002644 | NEC Cervix | DUP | chr17:81053668-81119580 | *BAIAP2, AATK* |
| TKHS190002641 | NEC Ovary | DUP | chr17:81379722-81548028 | *LOC100130370, MIR3186, MIR4740, FSCN2, BAHCC1, ACTG1, FAAP100* |
| TKHS190002650 | NEC Ovary | DEL | chr17:8143266-8255691 | *NA, MIR4521, LINC00324, PER1, PFAS, BORCS6, VAMP2, CTC1, TMEM107, AURKB* |
| TKHS190002641 | NEC Ovary | DUP | *chr17:81688704-81833852* | |
| TKHS190002644 | NEC Cervix | DUP | chr17:81824499-81941741 | *ALYREF, PPP1R27, MYADML2, NPB, MCRIP1, ARHGDIA, MAFG, P4HB, ANAPC11, SIRT7, PYCR1, PCYT2, MAFG-DT* |
| TKHS190002644 | NEC Cervix | DUP | chr17:82017309-82027011 | *CENPX, LRRC45, ASPSCR1* |
| TKHS190002641 | NEC Ovary | DUP | chr17:82030065-82091684 | *LRRC45, FASN, GPS1, DCXR, RAC3, RFNG, DUS1L* |
| TKHS190002644 | NEC Cervix | DUP | chr17:82033699-82053366 | *GPS1, DCXR, RAC3, RFNG* |
| TKHS190002653 | NEC Ovary | DEL | chr17:82081164-82085871 | *FASN* |
| TKHS190002646 | NEC Cervix | DUP | chr17:82082009-82084716 | *FASN* |
| TKHS190002641 | NEC Ovary | DUP | chr17:8254166-8319394 | *ARHGEF15, RANGRF, SLC25A35, PFAS* |
| TKHS190002644 | NEC Cervix | DUP | chr17:82741311-82907821 | *FN3K, TBCD, ZNF750* |
| TKHS190002645 | NEC Endometrium | DEL | chr18:12254384-14106409 | *LDLRAD4-AS1, MIR4526, MIR5190, PRELID3A, AFG3L2, CIDEA, FAM210A, ZNF519, MC2R, MC5R, CEP192, SPIRE1, PSMG2, PTPN2, LDLRAD4, CEP76, SEH1L, TUBB6, RNMT* |
| TKHS190002646 | NEC Cervix | DUP | chr18:158699-14837289 | *LINC01387, LOC100192426, LDLRAD4-AS1, MIR4317, MIR3156-2, MIR4526, MIR3976, MIR5190, DLGAP1-AS4, MYL12B, NDC80, MYL12A, PRELID3A, CETN1, RALBP1, AFG3L2, RAB31, CIDEA, ADCYAP1, FAM210A, APCDD1, LINC00526, ZNF519, RAB12, DLGAP1-AS3, EPB41L3, ANKRD12, MTCL1, SMCHD1, CLUL1, GNAL, DLGAP1-AS5, LAMA1, CYP4F35P, LINC00667,* |

| | | | | |
|---|---|---|---|---|
| | | | | *LRRC30, ANKRD62, IMPA2, ANKRD30B, POTEC, LINC00668, MC2R, MC5R, CXADRP3, ANKRD20A5P, NDUFV2, TYMSOS, C18orf61, CEP192, ENOSF1, LINC00470, SPIRE1, PSMG2, TWSG1, CHMP1B, PTPN2, PTPRM, PIEZO2, AKAIN1, CBX3P2, MIR3976HG, TMEM200C, METTL4, DLGAP1-AS1, MPPE1, TGIF1, LOC727896, TYMS, YES1, LDLRAD4, ZBTB14, ARHGAP28, CEP76, COLEC12, SEH1L, EMILIN2, TXNDC2, TUBB6, RNMT, MYOM1, NAPG, USP14, L3MBTL4, VAPA, DLGAP1, LPIN2, THOC1, PPP4R1* |
| TKHS190002646 | NEC Cervix | DUP | chr18:21395385-21518234 | *GREB1L* |
| TKHS190002646 | NEC Cervix | DUP | chr18:21659170-21857243 | *MIR320C1, ABHD3, MIR1-2, MIR133A1, MIB1* |
| TKHS190002646 | NEC Cervix | DUP | chr18:22993721-23753812 | *ANKRD29, RMC1, LAMA3, NPC1, RBBP8, TMEM241, RIOK3, CABLES1* |
| TKHS190002646 | NEC Cervix | DUP | chr18:23939223-24179835 | *OSBPL1A, TTC39C, CABYR, LAMA3* |
| TKHS190002645 | NEC Endometrium | DEL | chr18:2707563-3215223 | *SMCHD1, LOC727896, EMILIN2, MYOM1, LPIN2* |
| TKHS190002657 | NEC Ovary | DEL | chr18:3100320-3176134 | *MYOM1* |
| TKHS190002646 | NEC Cervix | DUP | chr18:31074683-31595276 | *DSG2-AS1, DSG4, DSC1, DSC2, DSG1, DSG2, DSG3, TTR* |
| TKHS190002646 | NEC Cervix | DUP | chr18:31917578-32393035 | *TRAPPC8, MEP1B, RNF138, RNF125, GAREM1* |
| TKHS190002650 | NEC Ovary | DUP | chr18:33245245-34877808 | *DTNA, CCDC178, ASXL3, NOL4* |
| TKHS190002645 | NEC Endometrium | DEL | chr18:346295-732965 | *CETN1, CLUL1, TYMSOS, ENOSF1, TYMS, YES1, COLEC12* |
| TKHS190002650 | NEC Ovary | DUP | chr18:36501932-36780188 | *TPGS2, FHOD3* |
| TKHS190002645 | NEC Endometrium | DUP | chr18:37160200-37321881 | *CELF4, KIAA1328* |
| TKHS190002650 | NEC Ovary | DUP | chr18:45912290-45939755 | *EPG5* |
| TKHS190002650 | NEC Ovary | DUP | chr18:46456519-46855435 | *LOXHD1, ST8SIA5, RNF165, PIAS2* |
| TKHS190002650 | NEC Ovary | DUP | chr18:47059556-48921910 | *MIR4743, ZBTB7C, SMAD2, SMAD7, IER3IP1, SKOR2, KATNAL2, HDHD2, CTIF* |
| TKHS190002650 | NEC Ovary | DUP | chr18:50270011-50819702 | *SKA1, CXXC1, MBD1, MAPK4, MRO* |
| TKHS190002645 | NEC Endometrium | DEL | chr18:5290858-5433556 | *EPB41L3, ZBTB14* |
| TKHS190002650 | NEC Ovary | DUP | chr18:56781533-57476723 | *LINC-ROR, WDR7, BOD1L2, ST8SIA3, ONECUT2* |
| TKHS190002650 | NEC Ovary | DUP | chr18:62231194-62575550 | *ZCCHC2, RELCH, TNFRSF11A* |
| TKHS190002645 | NEC Endometrium | DEL | chr18:6851034-10801428 | *LOC100192426, RALBP1, RAB31, APCDD1, RAB12, ANKRD12, MTCL1, LAMA1, LRRC30, LINC00668, NDUFV2, TWSG1, PTPRM, PIEZO2, ARHGAP28, TXNDC2, NAPG, VAPA, PPP4R1* |
| TKHS190002657 | NEC Ovary | DEL | chr18:6942079-7012138 | *LAMA1* |
| TKHS190002650 | NEC Ovary | DUP | chr18:70057742-70642723 | *RTTN, SOCS6* |
| TKHS190002650 | NEC Ovary | DUP | chr18:76923071-77268902 | *GALR1, MBP, ZNF236* |
| TKHS190002653 | NEC Ovary | DEL | chr18:80034496-80133192 | *RBFADN, ADNP2, RBFA* |
| TKHS190002657 | NEC Ovary | DUP | chr19:10093349-10133701 | *SNORD105B, DNMT1, P2RY11, PPAN, SNORD105, PPAN-P2RY11, ANGPTL6, EIF3G* |
| TKHS190002657 | NEC Ovary | DUP | chr19:1037625-1043473 | *ABCA7, CNN2* |
| TKHS190002647 | NEC Endometrium | DUP | chr19:10459400-10467621 | *PDE4A* |
| TKHS190002657 | NEC Ovary | DUP | chr19:1063760-1086006 | *ABCA7, ARHGAP45* |
| TKHS190002641 | NEC Ovary | DEL | chr19:10858316-10928410 | *CARM1, C19orf38, YIPF2* |
| TKHS190002657 | NEC Ovary | DUP | chr19:10858316-10929702 | *CARM1, C19orf38, YIPF2, TIMM29* |

| | | | | |
|---|---|---|---|---|
| TKHS190002646 | NEC Cervix | DEL | chr19:10908039-10928410 | *CARM1, YIPF2* |
| TKHS190002657 | NEC Ovary | DUP | chr19:11147134-11202483 | *SPC24, KANK2, DOCK6* |
| TKHS190002643 | NEC Cervix | DUP | chr19:11166558-11202483 | *KANK2, DOCK6* |
| TKHS190002657 | NEC Ovary | DUP | chr19:1256995-1440462 | *CIRBP, CIRBP-AS1, EFNA2, GAMT, DAZAP1, NDUFS7, FAM174C, RPS15, PWWP3A, MIDN* |
| TKHS190002647 | NEC Endometrium | DUP | chr19:12701752-12715739 | *SNORD41, TNPO2* |
| TKHS190002657 | NEC Ovary | DUP | chr19:12706041-12847687 | *RNASEH2A, MAST1, SNORD41, HOOK2, TNPO2, JUNB, GET3, BEST2, PRDX2, TRIR, RTBDN* |
| TKHS190002643 | NEC Cervix | DUP | chr19:12772780-12810208 | *RNASEH2A, HOOK2, JUNB, PRDX2* |
| TKHS190002657 | NEC Ovary | DUP | chr19:12880972-13100748 | *KLF1, DNASE2, DAND5, FARSA, SYCE2, GCDH, LYL1, NFIX, RAD23A, CALR, GADD45GIP1* |
| TKHS190002646 | NEC Cervix | DEL | chr19:12947848-13090390 | *DAND5, NFIX, RAD23A, GADD45GIP1* |
| TKHS190002657 | NEC Ovary | DUP | chr19:13758868-13904913 | *LOC284454, C19orf53, NANOS3, MIR181C, MIR181D, ZSWIM4, C19orf57, CCDC130, MRI1* |
| TKHS190002647 | NEC Endometrium | DUP | chr19:13809070-13831050 | *ZSWIM4* |
| TKHS190002647 | NEC Endometrium | DUP | chr19:13919130-13930289 | *CC2D1A* |
| TKHS190002657 | NEC Ovary | DUP | chr19:13979447-14093237 | *MISP3, RLN3, PALM3, PRKACA, RFX1, C19orf67, SAMD1, IL27RA* |
| TKHS190002657 | NEC Ovary | DUP | chr19:14402597-14446561 | *DDX39A, PKN1, ADGRE5* |
| TKHS190002647 | NEC Endometrium | DUP | chr19:14952928-15111439 | *OR1I1, CCDC105, CASP14, SLC1A6, SYDE1* |
| TKHS190002647 | NEC Endometrium | DUP | chr19:15170331-15192519 | *NOTCH3* |
| TKHS190002657 | NEC Ovary | DUP | chr19:15452045-15458655 | *RASAL3* |
| TKHS190002657 | NEC Ovary | DUP | chr19:17283225-17333800 | *ANKLE1, ANO8, MRPL34, DDA1, ABHD8* |
| TKHS190002644 | NEC Cervix | DUP | chr19:17305831-17333800 | *ANO8, MRPL34, DDA1* |
| TKHS190002647 | NEC Endometrium | DUP | chr19:1784819-1791861 | *ATP8B3* |
| TKHS190002657 | NEC Ovary | DUP | chr19:18143763-18532814 | *MIR3188, MIR3189, IFI30, LRRC25, SSBP4, MAST3, FKBP8, LSM4, JUND, PDE4C, ISYNA1, PIK3R2, PGPEP1, RAB3A, LOC729966, IQCN, ELL, MPV17L2, GDF15* |
| TKHS190002641 | NEC Ovary | DEL | chr19:18193282-18220515 | *PDE4C, RAB3A, MPV17L2* |
| TKHS190002646 | NEC Cervix | DEL | chr19:18432973-18465575 | *SSBP4, ISYNA1, ELL* |
| TKHS190002657 | NEC Ovary | DUP | chr19:18667577-18788319 | *COMP, CRTC1, KLHL26* |
| TKHS190002657 | NEC Ovary | DUP | chr19:18832210-18920909 | *CERS1, COPE, GDF1, DDX49, UPF1, HOMER3* |
| TKHS190002647 | NEC Endometrium | DUP | chr19:19120765-19133306 | *TMEM161A* |
| TKHS190002647 | NEC Endometrium | DUP | chr19:1918127-1987692 | *SCAMP4, CSNK1G2, CSNK1G2-AS1, BTBD2* |
| TKHS190002647 | NEC Endometrium | DUP | chr19:19260813-19268754 | *HAPLN4, TM6SF2* |
| TKHS190002657 | NEC Ovary | DUP | chr19:2039613-2426088 | *MIR1227, MIR4321, IZUMO4, JSRP1, MOB3A, TIMM13, AMH, MKNK2, TMPRSS9, PEAK3, OAZ1, LSM7, PLEKHJ1, SPPL2B, LINGO3, SF3A2, DOT1L, AP3D1* |
| TKHS190002650 | NEC Ovary | DEL | chr19:2132471-2247022 | *MIR1227, PLEKHJ1, SF3A2, DOT1L, AP3D1* |
| TKHS190002647 | NEC Endometrium | DUP | chr19:291285-327989 | *MIER2, PLPP2* |
| TKHS190002654 | NEC Cervix | DUP | chr19:2980100-2991984 | *TLE6* |

| | | | | |
|---|---|---|---|---|
| TKHS190002657 | NEC Ovary | DUP | chr19:3055664-3062927 | *TLE5* |
| TKHS190002647 | NEC Endometrium | DUP | chr19:3435083-3539260 | *C19orf71, MFSD12, SMIM24, NFIC, FZR1, DOHH* |
| TKHS190002657 | NEC Ovary | DUP | chr19:35262546-35295981 | *MAG, LSR, HAMP, USF2* |
| TKHS190002657 | NEC Ovary | DUP | chr19:35449715-35547447 | *TMEM147-AS1, TMEM147, GAPDHS, FFAR2, SBSN, KRTDAP, DMKN* |
| TKHS190002657 | NEC Ovary | DUP | chr19:35738282-35752488 | *U2AF1L4, PSENEN, LIN37, IGFLR1, KMT2B* |
| TKHS190002643 | NEC Cervix | DUP | chr19:35743809-35759991 | *HSPB6, PROSER3, U2AF1L4, PSENEN, LIN37, IGFLR1* |
| TKHS190002647 | NEC Endometrium | DUP | chr19:36016158-36035592 | *THAP8, CLIP3* |
| TKHS190002647 | NEC Endometrium | DUP | chr19:3637410-3653589 | *PIP5K1C* |
| TKHS190002657 | NEC Ovary | DUP | chr19:3643243-3738663 | *PIP5K1C, TJP3* |
| TKHS190002657 | NEC Ovary | DUP | chr19:38616195-38734389 | *MAP4K1, CAPN12, EIF3K, ACTN4* |
| TKHS190002657 | NEC Ovary | DUP | chr19:38869943-38907707 | *RINL, SIRT2, NFKBIB* |
| TKHS190002657 | NEC Ovary | DUP | chr19:38917724-38922078 | *SARS2* |
| TKHS190002657 | NEC Ovary | DUP | chr19:39388339-39392522 | *PAF1, MED29* |
| TKHS190002643 | NEC Cervix | DUP | chr19:39388560-39397757 | *PAF1, MED29* |
| TKHS190002645 | NEC Endometrium | DUP | chr19:42729123-43194594 | *PSG8-AS1, PSG8, PSG1, PSG2, PSG3, PSG4, PSG5, PSG6, PSG7, PSG11, PSG10P* |
| TKHS190002657 | NEC Ovary | DUP | chr19:4408903-4448415 | *CHAF1A, MIR4746, UBXN6* |
| TKHS190002647 | NEC Endometrium | DUP | chr19:4428664-4445623 | *CHAF1A, UBXN6* |
| TKHS190002657 | NEC Ovary | DUP | chr19:4499491-5040011 | *DPP9-AS1, MIR4747, PLIN3, SEMA6B, LRG1, TNFAIP8L1, TICAM1, KDM4B, MIR7-3HG, UHRF1, MIR7-3, PLIN5, FEM1A, MYDGF, ARRDC5, PLIN4, HDGFL2, DPP9* |
| TKHS190002657 | NEC Ovary | DUP | chr19:45406697-45500710 | *CD3EAP, PPM1N, ERCC1, FOSB, RTN2* |
| TKHS190002657 | NEC Ovary | DUP | chr19:45759414-46040416 | *SIX5, DMPK, DMWD, IRF2BP1, FOXA3, MYPOP, NANOS2, BHMG1, IGFL4, NOVA2, CCDC61, MIR769, RSPH6A, SYMPK, PGLYRP1* |
| TKHS190002647 | NEC Endometrium | DUP | chr19:4651870-4685771 | *DPP9-AS1, TNFAIP8L1, MYDGF, DPP9* |
| TKHS190002641 | NEC Ovary | DEL | chr19:4658005-4684862 | *DPP9-AS1, MYDGF, DPP9* |
| TKHS190002653 | NEC Ovary | DUP | chr19:474621-519423 | *ODF3L2, MADCAM1, TPGS1* |
| TKHS190002657 | NEC Ovary | DUP | chr19:47480958-47514940 | *NAPA-AS1, KPTN, NAPA* |
| TKHS190002657 | NEC Ovary | DUP | chr19:48311553-48363531 | *EMP3, TMEM143, CCDC114* |
| TKHS190002657 | NEC Ovary | DUP | chr19:48702957-48806717 | *FUT1, FUT2, FGF21, MAMSTR, IZUMO1, RASIP1, BCAT2* |
| TKHS190002641 | NEC Ovary | DEL | chr19:4891046-5215412 | *MIR4747, KDM4B, UHRF1, PTPRS, ARRDC5* |
| TKHS190002657 | NEC Ovary | DUP | chr19:48965508-48993112 | *FTL, GYS1* |
| TKHS190002657 | NEC Ovary | DUP | chr19:49070063-49166215 | *HRC, KCNA7, TRPM4, C19orf73, LIN7B, SNRNP70, PPFIA3* |
| TKHS190002647 | NEC Endometrium | DUP | chr19:49171578-49189091 | *TRPM4* |
| TKHS190002657 | NEC Ovary | DUP | chr19:49374717-49450939 | *GFY, PTH2, CCDC155, DKKL1, PIH1D1, SLC17A7* |
| TKHS190002657 | NEC Ovary | DUP | chr19:49513882-49685037 | *MIR5088, FCGRT, PRMT1, IRF3, NOSIP, PRRG2, RCN3, PRR12, SCAF1, RRAS, BCL2L12* |
| TKHS190002657 | NEC Ovary | DUP | chr19:49801390-49836962 | *AP2A1, FUZ, MED25* |

| | | | | |
|---|---|---|---|---|
| TKHS190002647 | NEC Endometrium | DUP | chr19:49932422-49960935 | *MIR4751, SIGLEC11, ATF5* |
| TKHS190002647 | NEC Endometrium | DUP | chr19:50399371-50407415 | *POLD1* |
| TKHS190002657 | NEC Ovary | DUP | chr19:50451280-50519415 | *FAM71E1, JOSD2, EMC10, MYBPC2, ASPDH, LRRC4B* |
| TKHS190002657 | NEC Ovary | DUP | chr19:50711371-50860127 | *LINC01869, GPR32, KLK3, KLK1, SHANK1, KLK15, CLEC11A, SNORD88A, SNORD88B, SNORD88C, C19orf48, ACP4* |
| TKHS190002657 | NEC Ovary | DUP | chr19:51022545-51060093 | *KLK11, KLK13, KLK12* |
| TKHS190002647 | NEC Endometrium | DUP | chr19:51643779-51719928 | *SIGLEC14, SPACA6, HAS1, MIRLET7E, MIR125A, MIR99B* |
| TKHS190002657 | NEC Ovary | DUP | chr19:55132915-55174999 | *DNAAF3, SYT5, TNNI3, TNNT1* |
| TKHS190002647 | NEC Endometrium | DUP | chr19:55236913-55274622 | *PPP6R1, HSPBP1* |
| TKHS190002657 | NEC Ovary | DUP | chr19:55437605-55689950 | *SBK3, ZNF865, U2AF2, ZNF524, ZNF784, ZNF579, SSC5D, CCDC106, EPN1, ZNF580, ZNF581, NAT14, SBK2, SHISA7, ISOC2, FIZ1, ZNF628* |
| TKHS190002657 | NEC Ovary | DUP | chr19:5654061-5707143 | *MICOS13, RPL36, HSD11B1L, SAFB, LONP1* |
| TKHS190002647 | NEC Endometrium | DUP | chr19:5700789-5708403 | *LONP1* |
| TKHS190002657 | NEC Ovary | DUP | chr19:5776161-5791141 | *PRR22, CATSPERD, DUS3L* |
| TKHS190002643 | NEC Cervix | DUP | chr19:58545425-58547906 | *TRIM28* |
| TKHS190002647 | NEC Endometrium | DUP | chr19:6002721-6147675 | *LOC100128568, RFX2, ACSBG2* |
| TKHS190002657 | NEC Ovary | DUP | chr19:7468825-7507141 | *NA, ARHGEF18, TEX45, PEX11G* |
| TKHS190002647 | NEC Endometrium | DUP | chr19:7539918-7551107 | *PNPLA6* |
| TKHS190002657 | NEC Ovary | DUP | chr19:7846543-7981186 | *TGFBR3L, TIMM44, EVI5L, ELAVL1, LYPLA2P2, CTXN1, MAP2K7, SNAPC2, LRRC8E, PRR36* |
| TKHS190002657 | NEC Ovary | DUP | chr19:8131554-8251249 | *CERS4, FBN3* |
| TKHS190002647 | NEC Endometrium | DUP | chr19:814434-843626 | *PRTN3, AZU1, PLPPR3* |
| TKHS190002657 | NEC Ovary | DUP | chr19:8303855-8325096 | *KANK3, NDUFA7, CD320, RPS28* |
| TKHS190002657 | NEC Ovary | DUP | chr19:860861-877180 | *MED16, CFD* |
| TKHS190002647 | NEC Endometrium | DUP | chr19:8895668-8981138 | *MUC16* |
| TKHS190002641 | NEC Ovary | DUP | chr2:176002160-176190142 | *HOXD-AS2, HOXD1, HOXD3, HOXD4, HOXD8, HOXD9, HOXD10, HOXD11, HOXD12, HOXD13, EVX2, HAGLR, MIR10B, LNPK* |
| TKHS190002641 | NEC Ovary | DUP | chr2:218356885-218436626 | *CATIP, MIR26B, CTDSP1, SLC11A1, VIL1* |
| TKHS190002657 | NEC Ovary | DEL | chr2:218386635-218431042 | *MIR26B, CTDSP1, SLC11A1, VIL1* |
| TKHS190002646 | NEC Cervix | DUP | chr2:218404297-218435434 | *CTDSP1, VIL1* |
| TKHS190002654 | NEC Cervix | DUP | chr2:218882161-219010085 | *LOC100129175, CRYBA2, LINC00608, CFAP65, MIR375, FEV, WNT10A, CDK5R2* |
| TKHS190002654 | NEC Cervix | DUP | chr2:219169823-219230405 | *ABCB6, ZFAND2B, SLC23A3, CNPPD1, ANKZF1, ATG9A, RETREG2, NHEJ1* |
| TKHS190002643 | NEC Cervix | DUP | chr2:219210230-219233433 | *ABCB6, ANKZF1, ATG9A* |
| TKHS190002641 | NEC Ovary | DUP | chr2:219246000-219309332 | *DNAJB2, MIR153-1, PTPRN, TUBA4A, TUBA4B, STK16* |
| TKHS190002654 | NEC Cervix | DUP | chr2:219251565-219451807 | *SPEG, DES, DNPEP, DNAJB2, RESP18, MIR153-1, PTPRN, TUBA4A, TUBA4B* |
| TKHS190002657 | NEC Ovary | DEL | chr2:219284632-219298103 | *DNAJB2, MIR153-1, PTPRN* |
| TKHS190002657 | NEC Ovary | DEL | chr2:219441558-219641728 | *MIR3132, SPEG, STK11IP, TMEM198, OBSL1, GMPPA, INHA, ASIC4, SLC4A3, CHPF* |

| | | | | |
|---|---|---|---|---|
| TKHS190002654 | NEC Cervix | DUP | chr2:219479783-219568324 | *MIR3132, SPEG, TMEM198, OBSL1, GMPPA, ASIC4, CHPF* |
| TKHS190002644 | NEC Cervix | DEL | chr2:219556454-219568324 | *OBSL1* |
| TKHS190002657 | NEC Ovary | DEL | chr2:232456200-232524844 | *ALPI, PRSS56, ECEL1* |
| TKHS190002644 | NEC Cervix | DEL | chr2:233457221-233462735 | *DGKD* |
| TKHS190002654 | NEC Cervix | DUP | *chr2:233789497-233831540* | |
| TKHS190002653 | NEC Ovary | DUP | chr2:237324774-237381499 | *COL6A3* |
| TKHS190002654 | NEC Cervix | DUP | chr2:238238827-238260953 | *HES6, PER2* |
| TKHS190002654 | NEC Cervix | DUP | chr2:240572423-241033875 | *CAPN10, AGXT, LOC200772, SNED1, GPR35, AQP12A, KIF1A, RNPEPL1, AQP12B, MAB21L4* |
| TKHS190002644 | NEC Cervix | DEL | chr2:240723975-240785100 | *KIF1A* |
| TKHS190002654 | NEC Cervix | DUP | chr2:241491061-241508035 | *STK25, FARP2* |
| TKHS190002644 | NEC Cervix | DUP | chr2:24820724-24830825 | *ADCY3, CENPO* |
| TKHS190002644 | NEC Cervix | DUP | chr2:26476891-26503845 | *OTOF* |
| TKHS190002643 | NEC Cervix | DUP | chr2:27035135-27080270 | *OST4, EMILIN1, TMEM214, AGBL5* |
| TKHS190002643 | NEC Cervix | DUP | chr2:27131672-27232694 | *PREB, TCF23, PRR30, ATRAID, CAD, SLC5A6* |
| TKHS190002644 | NEC Cervix | DUP | chr2:27207258-27234685 | *ATRAID, CAD, SLC5A6* |
| TKHS190002644 | NEC Cervix | DUP | chr2:27342826-27380526 | *ZNF513, GTF3C2, EIF2B4, SNX17* |
| TKHS190002644 | NEC Cervix | DUP | chr2:27437262-27449031 | *KRTCAP3, IFT172, NRBP1* |
| TKHS190002644 | NEC Cervix | DUP | chr2:28602821-28783978 | *PLB1, SPDYA, PPP1CB* |
| TKHS190002644 | NEC Cervix | DUP | chr2:28999181-29133816 | *TOGARAM2, PCARE, CLIP4* |
| TKHS190002644 | NEC Cervix | DUP | chr2:31372228-31403144 | *XDH* |
| TKHS190002644 | NEC Cervix | DUP | chr2:70921098-70985300 | *VAX2, ATP6V1B1, ANKRD53* |
| TKHS190002644 | NEC Cervix | DUP | chr2:74432467-74531371 | *INO80B-WBP1, MRPL53, LBX2-AS1, DQX1, WBP1, HTRA2, TLX2, AUP1, RTKN, TTC31, MOGS, INO80B, PCGF1, CCDC142, LBX2* |
| TKHS190002641 | NEC Ovary | DEL | chr2:96828567-96986313 | *ANKRD23, CNNM3, ANKRD39, FAM178B, SEMA4C* |
| TKHS190002653 | NEC Ovary | DUP | chr2:96863682-96986313 | *FAM178B, SEMA4C* |
| TKHS190002641 | NEC Ovary | DEL | chr2:97647067-97760914 | *ACTR1B, COX5B, TMEM131, C2orf92, ZAP70* |
| TKHS190002644 | NEC Cervix | DUP | chr2:9845220-9986282 | *GRHL1, TAF1B* |
| TKHS190002654 | NEC Cervix | DUP | chr20:10641460-10664014 | *JAG1* |
| TKHS190002654 | NEC Cervix | DUP | chr20:1266654-1320435 | *FKBP1A-SDCBP2, SDCBP2, SNPH* |
| TKHS190002657 | NEC Ovary | DEL | chr20:17614737-17636729 | *RRBP1* |
| TKHS190002654 | NEC Cervix | DUP | chr20:18542296-19886780 | *C20orf78, LOC100130264, LINC00653, SEC23B, SCP2D1, LINC00652, SMIM26, RIN2, SLC24A3, DTD1* |
| TKHS190002654 | NEC Cervix | DUP | chr20:1922313-2865495 | *SNORD119, TMEM239, MIR1292, NOP56, TMC2, C20orf141, SIRPA, STK35, IDH3B, TGM6, PDYN, CPXM1, EBF4, PTPRA, VPS16, PCED1A, SNRPB, SNORA51, SNORD110, TGM3, ZNF343* |
| TKHS190002654 | NEC Cervix | DUP | chr20:20050470-20640878 | *CFAP61, INSM1, CRNKL1, RALGAPA2* |
| TKHS190002654 | NEC Cervix | DUP | chr20:229278-417342 | *RBCK1, C20orf96, DEFB129, DEFB132, TRIB3, SOX12, NRSN2, ZCCHC3* |

| | | | | |
|---|---|---|---|---|
| TKHS190002655 | NEC Cervix | DUP | chr20:2317068-2463227 | *SNORD119, TGM6, SNRPB, TGM3* |
| TKHS190002654 | NEC Cervix | DUP | chr20:24958928-25294292 | *LOC284798, VSX1, APMAP, PYGB, ACSS1, CST7, ENTPD6* |
| TKHS190002654 | NEC Cervix | DUP | chr20:3021309-3213226 | *UBOX5-AS1, UBOX5, GNRH2, ITPA, OXT, AVP, PTPRA, FASTKD5, MRPS26, DDRGK1, LZTS3* |
| TKHS190002654 | NEC Cervix | DUP | chr20:31368651-31722218 | *MCTS2P, MIR3193, HM13-AS1, DEFB118, LINC00028, DEFB119, DEFB121, DEFB122, DEFB123, DEFB124, REM1, ID1, BCL2L1, HM13, COX4I2* |
| TKHS190002644 | NEC Cervix | DEL | chr20:31819581-31845542 | *FOXS1, MYLK2* |
| TKHS190002654 | NEC Cervix | DUP | chr20:31819581-32316874 | *DUSP15, TSPY26P, CCM2L, TTLL9, FOXS1, POFUT1, HCK, XKR7, PLAGL2, PDRG1, MYLK2, KIF3B, TM9SF4* |
| TKHS190002654 | NEC Cervix | DUP | chr20:32366384-32703458 | *NOL4L, NOL4L-DT, COMMD7, ASXL1, C20orf203* |
| TKHS190002654 | NEC Cervix | DUP | chr20:32846618-33302412 | *BPIFB6, BPIFA3, BPIFA2, SUN5, BPIFB4, MAPRE1, BPIFA4P, BPIFB3, BPIFA1, BPIFB2, BPIFB1* |
| TKHS190002655 | NEC Cervix | DUP | chr20:33086021-33103014 | *BPIFB4* |
| TKHS190002654 | NEC Cervix | DUP | chr20:33408507-33680416 | *C20orf144, ACTL10, E2F1, NECAB3, SNTA1, CBFA2T2* |
| TKHS190002654 | NEC Cervix | DUP | chr20:34534628-34709524 | *PIGU, TP53INP2, DYNLRB1, MAP1LC3A* |
| TKHS190002654 | NEC Cervix | DUP | chr20:35384624-35669685 | *RBM12, CEP250, C20orf173, ERGIC3, UQCC1, SPAG4, GDF5, CPNE1, NFS1* |
| TKHS190002654 | NEC Cervix | DUP | chr20:35931249-36011257 | *CNBD2, PHF20, SCAND1* |
| TKHS190002654 | NEC Cervix | DUP | chr20:36195329-36634589 | *TGIF2-RAB5IF, MYL9, EPB41L1, DLGAP4, AAR2, RAB5IF, TGIF2, SLA2* |
| TKHS190002654 | NEC Cervix | DUP | chr20:3659909-3889728 | *CENPB, HSPA12B, SPEF1, C20orf27, AP5S1, MAVS, GFRA4, SIGLEC1, PANK2, ADAM33, CDC25B* |
| TKHS190002654 | NEC Cervix | DUP | chr20:37732879-38246126 | *VSTM2L, CTNNBL1, RPRD1B, TGM2, KIAA1755, TTI1* |
| TKHS190002654 | NEC Cervix | DUP | chr20:38623359-38725114 | *SLC32A1, ARHGAP40* |
| TKHS190002654 | NEC Cervix | DUP | chr20:38771559-38962407 | *PPP1R16B, DHX35, ACTR5, FAM83D* |
| TKHS190002650 | NEC Ovary | DUP | chr20:41162958-41358542 | *ZHX3, PLCG1, LPIN3* |
| TKHS190002641 | NEC Ovary | DEL | chr20:41169457-41412263 | *ZHX3, PLCG1, LPIN3, CHD6, EMILIN3* |
| TKHS190002642 | NEC Ovary | DUP | chr20:41172193-41416794 | *ZHX3, PLCG1, LPIN3, CHD6, EMILIN3* |
| TKHS190002650 | NEC Ovary | DUP | chr20:43686852-44186705 | *GTSF1L, MYBL2, JPH2, TOX2* |
| TKHS190002650 | NEC Ovary | DUP | chr20:45894823-46013796 | *MMP9, PLTP, CTSA, ZNF335, PCIF1* |
| TKHS190002641 | NEC Ovary | DEL | chr20:45949485-46013796 | *MMP9, ZNF335* |
| TKHS190002654 | NEC Cervix | DUP | chr20:45971210-46387437 | *MMP9, SLC35C2, SLC12A5, NCOA5, ELMO2, ZNF335, CDH22, CD40* |
| TKHS190002641 | NEC Ovary | DEL | chr20:46040373-46354965 | *SLC35C2, SLC12A5, NCOA5, CDH22, CD40* |
| TKHS190002650 | NEC Ovary | DUP | chr20:47224317-47298947 | *ZMYND8* |
| TKHS190002650 | NEC Ovary | DUP | chr20:48625885-48651583 | *PREX1* |
| TKHS190002654 | NEC Cervix | DUP | chr20:49874705-50818282 | *TRERNA1, CEBPB, RIPOR3, SLC9A8, LINC01270, TMEM189, TMEM189-UBE2V1, BCAS4, RNF114, PTPN1, SNAI1, MIR645, UBE2V1, PARD6B, SPATA2* |
| TKHS190002653 | NEC Ovary | DUP | chr20:51671115-51792352 | *ATP9A, SALL4* |
| TKHS190002654 | NEC Cervix | DUP | chr20:57366807-57659697 | *NA, CTCFL, PCK1, RBM38, PMEPA1, ZBP1, RAE1* |
| TKHS190002654 | NEC Cervix | DUP | chr20:5863024-5952528 | *CHGB, SHLD1, TRMT6, MCM8* |
| TKHS190002654 | NEC Cervix | DUP | chr20:604591-968138 | *SLC52A3, SRXN1, RSPO4, ANGPT4, TCF15, FAM110A, SCRT2* |

| | | | | |
|---|---|---|---|---|
| TKHS190002644 | NEC Cervix | DEL | chr20:61844668-61929842 | *CDH4* |
| TKHS190002654 | NEC Cervix | DUP | chr20:61844668-62327669 | *CDH4, MIR1257, ADRM1, HRH3, LSM14B, SS18L1, MTG2, LAMA5, PSMA7, TAF4, OSBPL2* |
| TKHS190002654 | NEC Cervix | DUP | chr20:62346506-62395014 | *LAMA5, RPS21, CABLES2* |
| TKHS190002654 | NEC Cervix | DUP | chr20:62465340-62882414 | *UCKL1-AS1, MIR941-1, MIR941-3, MIR1914, RGS19, SAMD10, ABHD16B, LKAAEAR1, PRPF6, NPBWR2, C20orf204, MYT1, OPRL1, SOX18, UCKL1, ZNF512B, TCEA2, MIR647, TPD52L2, DNAJC5* |
| TKHS190002644 | NEC Cervix | DEL | chr20:62944739-63278995 | *LINC00029, LINC01056, FLJ16779, MIR3196, BHLHE23, NKAIN4, MIR124-3, YTHDF1, GID8, ARFGAP1, SLC17A9, LINC01749, HAR1A, HAR1B, BIRC7* |
| TKHS190002654 | NEC Cervix | DUP | chr20:62944739-63666079 | *LINC00029, LINC01056, FLJ16779, MIR4326, MIR3196, RTEL1-TNFRSF6B, CHRNA4, BHLHE23, NKAIN4, EEF1A2, GMEB2, KCNQ2, MIR124-3, STMN3, RTEL1, YTHDF1, GID8, ARFGAP1, PTK6, COL20A1, SLC17A9, LINC01749, SRMS, HAR1A, HAR1B, FNDC11, PPDPF, BIRC7, HELZ2* |
| TKHS190002644 | NEC Cervix | DEL | chr20:63326096-63442531 | *CHRNA4, KCNQ2, COL20A1* |
| TKHS190002654 | NEC Cervix | DUP | chr20:63692805-63993485 | *UCKL1-AS1, MIR941-1, MIR941-3, MIR1914, RTEL1-TNFRSF6B, ARFRP1, ZBTB46, SAMD10, ABHD16B, PRPF6, RTEL1, LIME1, UCKL1, SLC2A4RG, ZNF512B, MIR647, TPD52L2, DNAJC5, ZGPAT, TNFRSF6B* |
| TKHS190002654 | NEC Cervix | DUP | chr20:64066915-64240448 | *RGS19, LKAAEAR1, NPBWR2, MYT1, OPRL1, TCEA2* |
| TKHS190002642 | NEC Ovary | DUP | chr21:30319403-30561289 | *MIR4327, KRTAP13-1, KRTAP15-1, KRTAP13-4, KRTAP19-1, KRTAP13-2, KRTAP13-3, KRTAP23-1, KRTAP19-2, KRTAP19-3, KRTAP19-4, KRTAP19-5, KRTAP19-6, KRTAP19-7, KRTAP26-1, KRTAP27-1* |
| TKHS190002641 | NEC Ovary | DEL | chr21:41740838-41859228 | *RIPK4, PRDM15* |
| TKHS190002641 | NEC Ovary | DEL | chr21:41918105-42111710 | *UMODL1-AS1, C2CD2, ZBTB21, UMODL1* |
| TKHS190002646 | NEC Cervix | DEL | chr21:42271070-42444643 | *UBASH3A, TMPRSS3, TFF1, TFF2, TFF3, ABCG1* |
| TKHS190002641 | NEC Ovary | DEL | chr21:42765381-42862394 | *WDR4, PDE9A* |
| TKHS190002641 | NEC Ovary | DUP | chr21:44107417-44230089 | *ICOSLG, PWP2, GATD3A* |
| TKHS190002654 | NEC Cervix | DUP | chr21:44108984-44711514 | *LRRC3-DT, TSPEAR-AS2, ICOSLG, DNMT3L, AIRE, KRTAP12-2, KRTAP12-1, KRTAP10-10, KRTAP10-4, KRTAP10-6, KRTAP10-7, KRTAP10-9, KRTAP10-1, KRTAP10-11, KRTAP10-2, KRTAP10-5, KRTAP10-8, KRTAP10-3, KRTAP12-3, KRTAP12-4, KRTAP10-12, PFKL, TSPEAR, PWP2, TRPM2, CFAP410, LRRC3, GATD3A* |
| TKHS190002657 | NEC Ovary | DEL | chr21:44304275-44339194 | *PFKL, CFAP410* |
| TKHS190002641 | NEC Ovary | DUP | chr21:44508869-44638173 | *TSPEAR-AS2, KRTAP10-10, KRTAP10-4, KRTAP10-6, KRTAP10-7, KRTAP10-9, KRTAP10-1, KRTAP10-2, KRTAP10-5, KRTAP10-8, KRTAP10-3, TSPEAR* |
| TKHS190002653 | NEC Ovary | DUP | chr21:45981851-46182816 | *FTCD, COL6A1, COL6A2, SPATC1L* |
| TKHS190002654 | NEC Cervix | DUP | chr21:45999153-46112824 | *COL6A1, COL6A2* |
| TKHS190002641 | NEC Ovary | DEL | chr21:46001253-46275325 | *FTCD, MCM3AP-AS1, COL6A1, COL6A2, LSS, SPATC1L, MCM3AP* |
| TKHS190002646 | NEC Cervix | DEL | chr21:46138880-46209625 | *FTCD, LSS, SPATC1L* |
| TKHS190002647 | NEC Endometrium | DUP | chr22:20139194-20149640 | *ZDHHC8, CCDC188* |
| TKHS190002654 | NEC Cervix | DUP | chr22:21000647-21030351 | *TUBA3FP, THAP7-AS1, SLC7A4, THAP7, P2RX6* |
| TKHS190002647 | NEC Endometrium | DUP | chr22:24184766-24231530 | *GGT5, SUSD2* |
| TKHS190002647 | NEC Endometrium | DUP | chr22:25768115-26457967 | *SEZ6L, ASPHD2, MYO18B, HPS4* |
| TKHS190002648 | NEC Cervix | DEL | chr22:30893638-30935322 | *MORC2-AS1, MORC2, OSBP2* |
| TKHS190002643 | NEC Cervix | DUP | chr22:36145473-36266033 | *APOL2, APOL4, APOL3, APOL1* |
| TKHS190002643 | NEC Cervix | DUP | chr22:36314145-36518910 | *TXN2, MYH9, FOXRED2, EIF3D* |
| TKHS190002643 | NEC Cervix | DUP | chr22:37832958-37886845 | *ANKRD54, EIF3L, MIR658, MIR659* |

| | | | | |
|---|---|---|---|---|
| TKHS190002643 | NEC Cervix | DUP | chr22:38670884-38729662 | *CBY1, TOMM22, GTPBP1, JOSD1* |
| TKHS190002647 | NEC Endometrium | DEL | chr22:42127781-42129185 | *NDUFA6-DT, CYP2D6* |
| TKHS190002647 | NEC Endometrium | DUP | chr22:43171819-43231875 | *TTLL12, SCUBE1* |
| TKHS190002643 | NEC Cervix | DUP | chr22:45527847-45564992 | *FBLN1* |
| TKHS190002643 | NEC Cervix | DUP | chr22:50229404-50248145 | *HDAC10, TUBGCP6* |
| TKHS190002648 | NEC Cervix | DEL | chr22:50316174-50423614 | *DENND6B, PPP6R2* |
| TKHS190002647 | NEC Endometrium | DUP | chr22:50603223-50740694 | *MAPK8IP2, ARSA, ACR, SHANK3* |
| TKHS190002645 | NEC Endometrium | DUP | chr3:100629598-100695001 | *ADGRG7* |
| TKHS190002644 | NEC Cervix | DUP | chr3:121911574-122022077 | *ILDR1, SLC15A2* |
| TKHS190002644 | NEC Cervix | DUP | chr3:122414536-122577853 | *DTX3L, PARP15, KPNA1, WDR5B, PARP9* |
| TKHS190002644 | NEC Cervix | DUP | chr3:122680859-122718958 | *PARP14* |
| TKHS190002645 | NEC Endometrium | DUP | chr3:126914680-127034017 | *PLXNA1, CHCHD6* |
| TKHS190002648 | NEC Cervix | DEL | chr3:126988594-127030064 | *PLXNA1* |
| TKHS190002653 | NEC Ovary | DUP | chr3:128123584-128625653 | *DNAJB8, LINC01565, GATA2, DNAJB8-AS1, EEFSEC, RPN1, RUVBL1, LOC90246* |
| TKHS190002644 | NEC Cervix | DUP | chr3:134357705-134365909 | *AMOTL2* |
| TKHS190002644 | NEC Cervix | DUP | chr3:136854461-136995556 | *NCK1, IL20RB, SLC35G2* |
| TKHS190002644 | NEC Cervix | DEL | chr3:14819072-14922548 | *FGD5-AS1, FGD5* |
| TKHS190002644 | NEC Cervix | DUP | chr3:149145354-149221792 | *CP, HPS3* |
| TKHS190002644 | NEC Cervix | DUP | chr3:158571301-158734037 | *MLF1, LXN, RARRES1, GFM1* |
| TKHS190002644 | NEC Cervix | DUP | chr3:172127028-172820237 | *ECT2, GHSR, NCEH1, FNDC3B, TNFSF10* |
| TKHS190002654 | NEC Cervix | DEL | chr3:172506492-172760263 | *ECT2, NCEH1, TNFSF10* |
| TKHS190002648 | NEC Cervix | DEL | chr3:184164492-184189277 | *AP2M1, DVL3, ABCF3* |
| TKHS190002656 | NEC Cervix | DEL | chr3:184180169-184193452 | *AP2M1, ABCF3* |
| TKHS190002654 | NEC Cervix | DUP | chr3:184234266-184236749 | *VWA5B2* |
| TKHS190002644 | NEC Cervix | DUP | chr3:184299851-184354314 | *CLCN2, FAM131A, EIF4G1, PSMD2, SNORD66* |
| TKHS190002656 | NEC Cervix | DEL | chr3:184308708-184383642 | *CLCN2, FAM131A, EIF4G1, POLR2H, PSMD2, SNORD66, THPO, CHRD* |
| TKHS190002644 | NEC Cervix | DUP | chr3:186787127-187236567 | *ADIPOQ-AS1, RPL39L, RTP1, EIF4A2, MASP1, RFC4, SNORA63, SNORA4, ST6GAL1, ADIPOQ* |
| TKHS190002644 | NEC Cervix | DUP | chr3:189310386-190429731 | *MIR944, CLDN16, TMEM207, TPRG1, P3H2, TP63, CLDN1* |
| TKHS190002644 | NEC Cervix | DEL | chr3:38097744-38122605 | *DLEC1* |
| TKHS190002644 | NEC Cervix | DEL | chr3:46989511-47019840 | *NRADDP, NBEAL2, SETD2* |
| TKHS190002657 | NEC Ovary | DEL | chr3:47407130-47426169 | *SCAP, PTPN23* |
| TKHS190002644 | NEC Cervix | DEL | chr3:48565636-48569760 | *COL7A1* |
| TKHS190002644 | NEC Cervix | DEL | chr3:48586074-48593696 | *COL7A1* |

| | | | | |
|---|---|---|---|---|
| TKHS190002644 | NEC Cervix | DEL | chr3:48605770-48642467 | *CELSR3, TMEM89, SLC26A6, UQCRC1* |
| TKHS190002645 | NEC Endometrium | DEL | chr3:49096029-49104733 | *QARS1* |
| TKHS190002645 | NEC Endometrium | DEL | chr3:49129839-49132867 | *LAMB2* |
| TKHS190002644 | NEC Cervix | DEL | chr3:49530797-49663886 | *BSN-DT, DAG1, BSN* |
| TKHS190002644 | NEC Cervix | DEL | chr3:50193265-50257690 | *SLC38A3, GNAI2, GNAT1* |
| TKHS190002657 | NEC Ovary | DEL | chr3:50275012-50292466 | *LSMEM2, IFRD2, SEMA3B* |
| TKHS190002644 | NEC Cervix | DEL | chr3:50367812-50434429 | *CACNA2D2* |
| TKHS190002654 | NEC Cervix | DUP | chr3:50375644-50434429 | *CACNA2D2* |
| TKHS190002644 | NEC Cervix | DEL | chr3:51712473-51937736 | *IQCF4, IQCF1, GRM2, IQCF2, IQCF5, IQCF3, IQCF6, RRP9* |
| TKHS190002654 | NEC Cervix | DUP | chr3:52278201-52396538 | *GLYCTK, DNAH1, MIR135A1, WDR82* |
| TKHS190002644 | NEC Cervix | DEL | chr3:52287094-52396538 | *GLYCTK, DNAH1, MIR135A1* |
| TKHS190002642 | NEC Ovary | DUP | chr3:52331148-52361352 | *DNAH1* |
| TKHS190002654 | NEC Cervix | DUP | chr3:52435603-52442304 | *SEMA3G* |
| TKHS190002644 | NEC Cervix | DEL | chr3:52435603-52529334 | *NISCH, STAB1, SEMA3G, NT5DC2, TNNC1* |
| TKHS190002654 | NEC Cervix | DUP | chr3:52470859-52528935 | *NISCH, STAB1, NT5DC2* |
| TKHS190002657 | NEC Ovary | DEL | chr3:52484513-52536906 | *NISCH, STAB1, SMIM4, NT5DC2* |
| TKHS190002654 | NEC Cervix | DUP | chr3:53495167-53735503 | *CACNA1D* |
| TKHS190002654 | NEC Cervix | DUP | chr3:97895959-97967593 | *CRYBG3, RIOX2* |
| TKHS190002654 | NEC Cervix | DUP | chr4:139719323-139889306 | *MGST2, MAML3* |
| TKHS190002641 | NEC Ovary | DUP | chr4:2898431-2933663 | *MFSD10, ADD1* |
| TKHS190002642 | NEC Ovary | DEL | chr4:2928171-2933663 | *MFSD10, ADD1* |
| TKHS190002641 | NEC Ovary | DUP | chr4:6287093-6348010 | *PPP2R2C, WFS1* |
| TKHS190002644 | NEC Cervix | DUP | chr4:73740659-74099120 | *PPBPP2, CXCL1, CXCL2, CXCL3, CXCL8, PF4, PF4V1, PPBP, CXCL6, CXCL5* |
| TKHS190002643 | NEC Cervix | DUP | chr4:7774739-7984893 | *AFAP1, ABLIM2, AFAP1-AS1* |
| TKHS190002654 | NEC Cervix | DUP | chr4:8008042-8080802 | *ABLIM2* |
| TKHS190002643 | NEC Cervix | DUP | chr4:8366961-8416521 | *ACOX3* |
| TKHS190002641 | NEC Ovary | DUP | chr4:932043-1002485 | *SLC26A1, DGKQ, GAK, IDUA, TMEM175* |
| TKHS190002653 | NEC Ovary | DUP | chr5:1032152-1074671 | *MIR4635, SLC12A7, NKD2* |
| TKHS190002646 | NEC Cervix | DEL | chr5:1032152-1076812 | *MIR4635, SLC12A7, NKD2* |
| TKHS190002641 | NEC Ovary | DEL | chr5:1032152-1321819 | *MIR4457, MIR4635, SLC12A7, SLC6A19, SLC6A18, TERT, CLPTM1L, NKD2* |
| TKHS190002642 | NEC Ovary | DUP | chr5:116292451-116504944 | *COMMD10, SEMA6A* |
| TKHS190002642 | NEC Ovary | DUP | chr5:1210444-1282624 | *SLC6A19, SLC6A18, TERT* |
| TKHS190002646 | NEC Cervix | DEL | chr5:1232219-1321819 | *MIR4457, SLC6A18, TERT, CLPTM1L* |

| | | | | |
|---|---|---|---|---|
| TKHS190002653 | NEC Ovary | DUP | chr5:134115321-134146521 | *TCF7* |
| TKHS190002654 | NEC Cervix | DUP | chr5:14479919-14504593 | *TRIO* |
| TKHS190002644 | NEC Cervix | DUP | chr5:149003738-149063022 | *SH3TC2* |
| TKHS190002643 | NEC Cervix | DUP | chr5:151663455-151676188 | *SPARC* |
| TKHS190002650 | NEC Ovary | DUP | chr5:176337062-177096357 | *MIR4281, GPRIN1, CLTB, HIGD2A, FGFR4, RNF44, FAF2, ZNF346, EIF4E1B, TSPAN17, ARL10, HK3, SIMC1, NOP16, UIMC1, CDHR2, KIAA1191, SNCB, UNC5A* |
| TKHS190002641 | NEC Ovary | DEL | chr5:176530582-176896159 | *MIR4281, GPRIN1, RNF44, EIF4E1B, TSPAN17, HK3, CDHR2, SNCB, UNC5A* |
| TKHS190002650 | NEC Ovary | DUP | chr5:177267562-177441053 | *RGS14, LMAN2, F12, PRELID1, GRK6, PRR7-AS1, PFN3, RAB24, NSD1, SLC34A1, MXD3* |
| TKHS190002648 | NEC Cervix | DEL | chr5:177291954-177404664 | *RGS14, LMAN2, F12, PRELID1, PFN3, RAB24, NSD1, SLC34A1, MXD3* |
| TKHS190002641 | NEC Ovary | DEL | chr5:177291954-177441053 | *RGS14, LMAN2, F12, PRELID1, GRK6, PRR7-AS1, PFN3, RAB24, NSD1, SLC34A1, MXD3* |
| TKHS190002650 | NEC Ovary | DUP | chr5:180619665-180631778 | *FLT4* |
| TKHS190002641 | NEC Ovary | DEL | chr5:306602-454051 | *PDCD6, EXOC3, EXOC3-AS1, AHRR* |
| TKHS190002642 | NEC Ovary | DUP | chr5:58454583-58459092 | *PLK2* |
| TKHS190002653 | NEC Ovary | DUP | chr5:58455539-58459959 | *PLK2* |
| TKHS190002643 | NEC Cervix | DEL | chr5:716808-833807 | *ZDHHC11* |
| TKHS190002648 | NEC Cervix | DUP | chr6:100393487-100449699 | *SIM1* |
| TKHS190002648 | NEC Cervix | DUP | chr6:136557758-137009492 | *NHEG1, SLC35D3, MAP3K5, PEX7, IL20RA* |
| TKHS190002648 | NEC Cervix | DUP | chr6:138170662-138512339 | *MIR3145, HEBP2, ARFGEF3, NHSL1, PBOV1* |
| TKHS190002648 | NEC Cervix | DUP | chr6:16143699-16328310 | *GMPR, MYLIP, ATXN1* |
| TKHS190002641 | NEC Ovary | DUP | chr6:16143699-17110880 | *GMPR, MYLIP, STMND1, ATXN1* |
| TKHS190002648 | NEC Cervix | DUP | chr6:167136240-167359412 | *CCR6, GPR31, TCP10L2, UNC93A, TCP10, TTLL2* |
| TKHS190002648 | NEC Cervix | DUP | chr6:167951186-168608239 | *HGC6.3, DACT2, KIF25, AFDN, SMOC2, FRMD1* |
| TKHS190002643 | NEC Cervix | DUP | chr6:26092685-26368912 | *BTN3A2, H1-3, H1-4, H1-6, H2AC8, H2AC7, H2BC5, HFE, H2AC6, H2BC8, H2BC7, H2BC6, H2BC9, H2BC10, H2BC4, H3C4, H3C6, H3C8, H4C4, H4C6, H4C3, H4C8, H4C5, H4C7, H3C7* |
| TKHS190002641 | NEC Ovary | DUP | chr6:27248847-27255131 | *PRSS16* |
| TKHS190002641 | NEC Ovary | DUP | *chr6:29440010-31980801* | |
| TKHS190002646 | NEC Cervix | DUP | *chr6:29727957-30075789* | |
| TKHS190002646 | NEC Cervix | DUP | *chr6:30556131-31148728* | |
| TKHS190002646 | NEC Cervix | DUP | *chr6:31540322-31972757* | |
| TKHS190002648 | NEC Cervix | DUP | *chr6:31766245-31792895* | |
| TKHS190002641 | NEC Ovary | DUP | *chr6:32026983-32316414* | |
| TKHS190002646 | NEC Cervix | DUP | *chr6:32039356-32217067* | |
| TKHS190002643 | NEC Cervix | DUP | *chr6:32169966-32188796* | |
| TKHS190002648 | NEC Cervix | DUP | *chr6:32202076-32223928* | |
| TKHS190002641 | NEC Ovary | DUP | chr6:32641972-33711686 | *HLA-DQA1, ITPR3, LINC00336, GGNBP1, BAK1, UQCC2* |
| TKHS190002643 | NEC Cervix | DUP | *chr6:32841531-32852502* | |
| TKHS190002646 | NEC Cervix | DUP | chr6:33168955-33575441 | *BAK1* |

| | | | | |
|---|---|---|---|---|
| TKHS190002643 | NEC Cervix | DUP | *chr6:33201246-33205912* | |
| TKHS190002643 | NEC Cervix | DUP | *chr6:33287614-33290450* | |
| TKHS190002643 | NEC Cervix | DUP | *chr6:33412264-33438919* | |
| TKHS190002648 | NEC Cervix | DUP | chr6:33442889-33687327 | *ITPR3, LINC00336, GGNBP1, BAK1* |
| TKHS190002641 | NEC Ovary | DUP | chr6:34527332-34544455 | *SPDEF, PACSIN1* |
| TKHS190002648 | NEC Cervix | DUP | chr6:35242622-35462914 | *FANCE, SCUBE3, DEF6, PPARD, ZNF76* |
| TKHS190002641 | NEC Ovary | DUP | chr6:35242622-35466345 | *FANCE, SCUBE3, DEF6, PPARD, ZNF76* |
| TKHS190002648 | NEC Cervix | DUP | chr6:35480312-35506173 | *TEAD3, TULP1* |
| TKHS190002641 | NEC Ovary | DUP | chr6:36052699-36291555 | *MAPK14, BRPF3, PNPLA1, MAPK13* |
| TKHS190002648 | NEC Cervix | DUP | chr6:36131271-36330690 | *BRPF3, PNPLA1, BNIP5, MAPK13* |
| TKHS190002641 | NEC Ovary | DUP | chr6:37170303-37360574 | *TMEM217, PIM1, TBC1D22B, RNF8* |
| TKHS190002641 | NEC Ovary | DUP | chr6:37462829-37696811 | *MIR4462, CCDC167, CMTR1, MDGA1* |
| TKHS190002648 | NEC Cervix | DUP | chr6:37484810-37696811 | *MIR4462, CCDC167, MDGA1* |
| TKHS190002641 | NEC Ovary | DUP | chr6:39026546-39316947 | *DNAH8, GLP1R, SAYSD1, KCNK16, KCNK5, KCNK17* |
| TKHS190002648 | NEC Cervix | DUP | chr6:39887486-41201008 | *APOBEC2, ADCY10P1, OARD1, TSPO2, UNC5CL, DAAM2, TREML1, LINC00951, MOCS1, NFYA, TREM2, LRFN2, TDRG1, TREML2* |
| TKHS190002648 | NEC Cervix | DUP | chr6:41335877-41597242 | *FOXP4, NCR2* |
| TKHS190002641 | NEC Ovary | DUP | chr6:41336087-41597242 | *FOXP4, NCR2* |
| TKHS190002641 | NEC Ovary | DUP | chr6:41653319-42185797 | *TOMM6, FRS3, TAF8, USP49, GUCA1A, GUCA1B, PRICKLE4, MDFI, PGC, C6orf132, BYSL, TFEB, CCND3, MED20* |
| TKHS190002648 | NEC Cervix | DUP | chr6:42128672-42185797 | *GUCA1A, GUCA1B, C6orf132* |
| TKHS190002646 | NEC Cervix | DUP | chr6:42935537-42966657 | *CNPY3, GNMT, PEX6* |
| TKHS190002641 | NEC Ovary | DUP | chr6:42935537-42966657 | *CNPY3, GNMT, PEX6* |
| TKHS190002648 | NEC Cervix | DUP | chr6:42962212-42968498 | *GNMT, PEX6* |
| TKHS190002641 | NEC Ovary | DUP | chr6:43056307-43521300 | *DNPH1, SLC22A7, LRRC73, CUL9, ZNF318, YIPF3, CRIP3, MRPL2, PTK7, DLK2, SRF, TTBK1, ABCC10, KLC4, TJAP1, POLR1C* |
| TKHS190002648 | NEC Cervix | DUP | chr6:43128977-43512563 | *DNPH1, SLC22A7, LRRC73, CUL9, ZNF318, YIPF3, CRIP3, PTK7, DLK2, SRF, TTBK1, ABCC10, TJAP1* |
| TKHS190002646 | NEC Cervix | DUP | chr6:43146618-43306124 | *DNPH1, SLC22A7, CUL9, CRIP3, PTK7, SRF, TTBK1* |
| TKHS190002641 | NEC Ovary | DUP | chr6:44002681-44287825 | *MIR4647, CAPN11, TCTE1, SLC29A1, SPATS1, C6orf223, HSP90AB1, SLC35B2, TMEM151B, NFKBIE, TMEM63B, MRPL14* |
| TKHS190002646 | NEC Cervix | DUP | chr6:44135017-44183235 | *CAPN11, TMEM63B* |
| TKHS190002648 | NEC Cervix | DUP | chr6:44146847-44230665 | *CAPN11, SLC29A1, TMEM63B* |
| TKHS190002648 | NEC Cervix | DUP | chr6:44273066-44310443 | *TCTE1, SPATS1, TMEM151B, AARS2* |
| TKHS190002641 | NEC Ovary | DEL | chr7:100090399-100198582 | *MIR4658, STAG3, COPS6, GPC2, CNPY4, MBLAC1, LAMTOR4, MIR106B, MIR25, MIR93, MCM7, MAP11, TAF6, GAL3ST4, AP4M1* |
| TKHS190002641 | NEC Ovary | DEL | chr7:100590174-100606630 | *PCOLCE-AS1, FBXO24, PCOLCE* |
| TKHS190002641 | NEC Ovary | DEL | chr7:100755226-100819889 | *EPHB4, ZAN* |
| TKHS190002641 | NEC Ovary | DEL | chr7:101211587-101216786 | *PLOD3* |
| TKHS190002653 | NEC Ovary | DUP | chr7:101321671-101622200 | *COL26A1, IFT22, MYL10* |

| | | | | |
|---|---|---|---|---|
| TKHS190002642 | NEC Ovary | DUP | chr7:101553326-101624264 | *COL26A1, MYL10* |
| TKHS190002657 | NEC Ovary | DUP | chr7:128807360-128840160 | *FLNC, CCDC136* |
| TKHS190002653 | NEC Ovary | DUP | chr7:132174778-132198636 | *PLXNA4* |
| TKHS190002657 | NEC Ovary | DUP | chr7:135362784-135422353 | *CNOT4* |
| TKHS190002643 | NEC Cervix | DEL | chr7:142065715-142094863 | *MGAM* |
| TKHS190002650 | NEC Ovary | DEL | *chr7:142384379-142520138* | |
| TKHS190002655 | NEC Cervix | DUP | *chr7:142384544-142507810* | |
| TKHS190002657 | NEC Ovary | DUP | chr7:149250159-149288691 | *ZNF783, LOC155060, ZNF212* |
| TKHS190002657 | NEC Ovary | DUP | chr7:149492859-150340609 | *ZBED6CL, ZNF746, ATP6V0E2, ZNF467, SSPO, ATP6V0E2-AS1, RARRES2, ZNF862, ACTR3C, LRRC61, ZNF767P, KRBA1* |
| TKHS190002641 | NEC Ovary | DEL | chr7:149721380-149832923 | *ZNF467, SSPO, KRBA1* |
| TKHS190002646 | NEC Cervix | DEL | chr7:149721380-149879677 | *ATP6V0E2, ZNF467, SSPO, ATP6V0E2-AS1, ZNF862, KRBA1* |
| TKHS190002657 | NEC Ovary | DUP | chr7:150856471-150977913 | *AOC1, KCNH2* |
| TKHS190002657 | NEC Ovary | DUP | chr7:151011788-151239746 | *ABCF2, CDK5, FASTK, ABCB8, AGAP3, ASB10, GBX1, ATG9B, NOS3, CHPF2, SLC4A2, SMARCD3, MIR671, TMUB1, ASIC3* |
| TKHS190002646 | NEC Cervix | DEL | chr7:151074399-151079021 | *FASTK, SLC4A2* |
| TKHS190002646 | NEC Cervix | DEL | chr7:151222521-151239746 | *ABCF2, CHPF2, SMARCD3, MIR671* |
| TKHS190002657 | NEC Ovary | DUP | chr7:154991006-155358179 | *HTR5A-AS1, PAXIP1-AS1, PAXIP1, HTR5A, INSIG1* |
| TKHS190002657 | NEC Ovary | DUP | chr7:2524655-2538944 | *MIR4648, BRAT1, LFNG* |
| TKHS190002641 | NEC Ovary | DEL | chr7:2649913-2700755 | *AMZ1, TTYH3* |
| TKHS190002650 | NEC Ovary | DEL | chr7:3969257-4178586 | *SDK1* |
| TKHS190002657 | NEC Ovary | DUP | chr7:4158448-4178586 | *SDK1* |
| TKHS190002657 | NEC Ovary | DUP | chr7:44110015-44114261 | *MIR4649, AEBP1* |
| TKHS190002657 | NEC Ovary | DUP | chr7:44707582-44801422 | *OGDH, PPIA, ZMIZ2* |
| TKHS190002657 | NEC Ovary | DUP | chr7:5489807-5623179 | *RNF216, ACTB, FSCN1, MIR589, FBXL18* |
| TKHS190002641 | NEC Ovary | DEL | chr7:5603257-5623179 | *RNF216, FSCN1* |
| TKHS190002657 | NEC Ovary | DUP | chr7:56068482-56084244 | *SUMF2, PHKG1, PSPH* |
| TKHS190002643 | NEC Cervix | DUP | chr7:5919849-5960864 | *RSPH10B, CCZ1* |
| TKHS190002648 | NEC Cervix | DEL | chr7:6139092-6171314 | *USP42, CYTH3* |
| TKHS190002643 | NEC Cervix | DEL | chr7:6763928-6805055 | *CCZ1B, RSPH10B2* |
| TKHS190002657 | NEC Ovary | DUP | chr7:729663-926644 | *ADAP1, SUN1, GET4, DNAAF5* |
| TKHS190002657 | NEC Ovary | DUP | chr7:73303826-73459718 | *TRIM50, NSUN5, FZD9, FKBP6, BAZ1B* |
| TKHS190002641 | NEC Ovary | DEL | chr7:74093250-74601297 | *LIMK1, RFC2, MIR590, EIF4H, CLIP2, LAT2, GTF2IRD1* |
| TKHS190002657 | NEC Ovary | DUP | chr7:74249739-74254389 | *RFC2* |
| TKHS190002650 | NEC Ovary | DUP | chr7:74317547-74538760 | *CLIP2, GTF2IRD1* |
| TKHS190002646 | NEC Cervix | DUP | chr7:75007994-75066792 | *NA, RCC1L* |

| | | | | |
|---|---|---|---|---|
| TKHS190002642 | NEC Ovary | DUP | chr7:87193198-87444975 | *TP53TG1, ABCB4, CROT, TMEM243, DMTF1* |
| TKHS190002642 | NEC Ovary | DUP | chr7:90160805-99624180 | *TMEM225B, MIR3609, NA, ATP5MF-PTCD1, MIR4652, ARPC1B, LOC101409256, AKAP9, SLC25A13, CDK6, BET1, ARPC1A, CPSF4, PDAP1, COL1A2, CYP51A1, BHLHA15, DLX5, DLX6, DYNC1I1, SAMD9L, ZSCAN25, FAM200A, TMEM130, LMTK2, PEG10, ZKSCAN5, HEPACAM2, FAM133B, BRI3, TECPR1, PTCD1, STEAP2, STEAP1, GNG11, GNGT1, DLX6-AS1, ZNF789, CCZ1P-OR7E38P, LRRD1, NA, KPNA7, ASNS, NPTX2, OCM2, PDK4, ASB4, PEX1, CDK14, PON1, PON2, PON3, ANKIB1, SAMD9, PPP1R9A, VPS50, BAIAP2L1, SDHAF3, SMURF1, MIR489, GATAD1, CASD1, TAC1, MIR591, MIR653, ZNF655, MTERF1, SEM1, TFPI2, CFAP69, CALCR, TRRAP, FZD1, RBM48, ZNF394, MYH16, NA, GTPBP10, KRIT1, BUD31, SGCE, CLDN12, ATP5MF* |
| TKHS190002641 | NEC Ovary | DEL | chr7:98848056-99035716 | *MIR3609, TMEM130, SMURF1, TRRAP* |
| TKHS190002642 | NEC Ovary | DUP | chr7:99848166-101216786 | *PCOLCE-AS1, SAP25, MIR4653, MIR4658, MUC12, POP7, STAG3, COPS6, AP1S1, MUC17, EPHB4, EPO, PPP1R35, GPC2, NYAP1, CNPY4, MBLAC1, FBXO24, GNB2, PILRB, PILRA, AGFG2, MOGAT3, GJC3, CASTOR3, NAT16, LAMTOR4, C7orf61, UFSP1, LRCH4, MIR106B, MIR25, MIR93, MCM7, ACHE, SPDYE3, SERPINE1, PCOLCE, ACTL6B, SRRT, PMS2P1, ZCWPW1, MAP11, MEPCE, AZGP1, SLC12A9, MOSPD3, GIGYF1, AZGP1P1, CYP3A43, TAF6, TFR2, TRIP6, VGF, ZAN, ZNF3, ZKSCAN1, ZSCAN21, PVRIG, GAL3ST4, OR2AE1, TSC22D4, TRIM56, TRIM4, PLOD3, AP4M1* |
| TKHS190002648 | NEC Cervix | DUP | chr8:101543241-104499519 | *MIR3151, BAALC-AS1, MIR5680, CTHRC1, BAALC-AS2, DPYS, DCAF13, LRP12, ODF1, RRM2B, UBR5, AZIN1, ATP6V1C1, KLF10, BAALC, GRHL2, SLC25A32, DCSTAMP, FZD6, NCALD, RIMS2* |
| TKHS190002648 | NEC Cervix | DUP | chr8:108239958-109498771 | *TMEM74, EIF3E, ENY2, TRHR, NUDCD1, PKHD1L1, EMC2* |
| TKHS190002648 | NEC Cervix | DUP | chr8:10834666-13115657 | *FAM66D, FAM66A, MIR3926-2, MIR3926-1, LOC100506990, LINC00681, DLC1, CTSB, TDH, FDFT1, DEFB109A, DEFB130A, NEIL2, GATA4, XKR6, LOC340357, USP17L2, FAM90A25P, LOC392196, USP17L7, PINX1, TRMT9B, DEFB135, DEFB136, DEFB134, BLK, LOC649352, FAM86B2, MTMR9, MIR598, ZNF705D, FAM90A2P, LOC729732, SOX7, FAM167A, SLC35G5, LINC00208, FAM167A-AS1, FAM86B1, LONRF1* |
| TKHS190002648 | NEC Cervix | DUP | chr8:112247020-113019179 | *MIR2053, CSMD3* |
| TKHS190002643 | NEC Cervix | DEL | chr8:11331107-11561452 | *TDH, BLK, FAM167A, SLC35G5, FAM167A-AS1* |
| TKHS190002648 | NEC Cervix | DUP | chr8:116657215-119783633 | *MIR3610, COLEC10, MAL2, SLC30A8, EXT1, SAMD12, AARD, CCN3, TNFRSF11B, ENPP2, SAMD12-AS1, RAD21, RAD21-AS1, TAF2, UTP23, EIF3H, MED30* |
| TKHS190002642 | NEC Ovary | DUP | chr8:119562857-119638780 | *ENPP2* |
| TKHS190002648 | NEC Cervix | DUP | chr8:119853047-120283778 | *DEPTOR, COL14A1, DSCC1* |
| TKHS190002642 | NEC Ovary | DUP | chr8:123019195-123231757 | *FAM83A-AS1, ZHX1-C8orf76, MIR4663, DERL1, C8orf76, FAM83A, TBC1D31* |
| TKHS190002642 | NEC Ovary | DUP | chr8:123429983-123737334 | *FBXO32, ANXA13, KLHL38, WDYHV1* |
| TKHS190002648 | NEC Cervix | DUP | chr8:123973434-124095038 | *FER1L6-AS1, FER1L6* |
| TKHS190002643 | NEC Cervix | DEL | chr8:124010594-124095038 | *FER1L6-AS1, FER1L6* |
| TKHS190002642 | NEC Ovary | DUP | chr8:124013431-124076325 | *FER1L6-AS1, FER1L6* |
| TKHS190002648 | NEC Cervix | DUP | chr8:124547000-125021945 | *ZNF572, LINC00964, NDUFB9, SQLE, MTSS1* |
| TKHS190002642 | NEC Ovary | DUP | chr8:124565662-125000071 | *ZNF572, LINC00964, NDUFB9, SQLE, MTSS1* |
| TKHS190002648 | NEC Cervix | DUP | chr8:130821342-132087739 | *HHLA1, ADCY8, EFR3A, OC90* |
| TKHS190002643 | NEC Cervix | DEL | chr8:132003236-132184367 | *HHLA1, EFR3A, KCNQ3, OC90* |
| TKHS190002648 | NEC Cervix | DUP | chr8:132777829-132868227 | *PHF20L1, TG* |
| TKHS190002642 | NEC Ovary | DUP | chr8:132836540-133116716 | *PHF20L1, SLA, TG* |

| | | | | |
|---|---|---|---|---|
| TKHS190002653 | NEC Ovary | DUP | chr8:132844156-133464957 | *NDRG1, PHF20L1, ST3GAL1, SLA, TG, CCN4* |
| TKHS190002643 | NEC Cervix | DEL | chr8:138137147-138606452 | *COL22A1, FAM135B* |
| TKHS190002648 | NEC Cervix | DUP | chr8:138137147-138844158 | *COL22A1, FAM135B* |
| TKHS190002642 | NEC Ovary | DUP | chr8:138138986-138195307 | *FAM135B* |
| TKHS190002648 | NEC Cervix | DUP | chr8:140458281-142879813 | *LNCOC1, MIR4472-1, PTP4A3, LYPD2, CYP11B1, TSNARE1, DENND3, ARC, AGO2, GML, GPR20, MROH5, THEM6, CHRAC1, LY6K, SLURP1, SLC45A4, PTK2, ADGRB1, LINC00051, LYNX1, LINC01300, PSCA, TRAPPC9, LY6D, JRK* |
| TKHS190002642 | NEC Ovary | DUP | chr8:141150286-143380035 | *CDC42P3, LY6E-DT, LNCOC1, MIR4472-1, PTP4A3, RHPN1, TOP1MT, LYPD2, CYP11B1, CYP11B2, TSNARE1, DENND3, ARC, GLI4, GML, GPR20, C8orf31, ZFP41, GPIHBP1, MROH5, LY6E, LY6H, THEM6, LY6K, SLURP1, SLC45A4, ADGRB1, LINC00051, LYNX1, LINC01300, RHPN1-AS1, ZNF696, PSCA, LY6D, JRK* |
| TKHS190002643 | NEC Cervix | DEL | chr8:141217090-143380035 | *CDC42P3, LY6E-DT, LNCOC1, MIR4472-1, PTP4A3, RHPN1, TOP1MT, LYPD2, CYP11B1, CYP11B2, TSNARE1, ARC, GLI4, GML, GPR20, C8orf31, ZFP41, GPIHBP1, MROH5, LY6E, LY6H, THEM6, LY6K, SLURP1, SLC45A4, ADGRB1, LINC00051, LYNX1, LINC01300, RHPN1-AS1, ZNF696, PSCA, LY6D, JRK* |
| TKHS190002648 | NEC Cervix | DUP | chr8:143269397-143560771 | *RHPN1, TOP1MT, ZC3H3, GLI4, ZFP41, MAFA, RHPN1-AS1, GSDMD, ZNF696* |
| TKHS190002653 | NEC Ovary | DUP | chr8:143558378-143616220 | *EEF1D, MROH6, PYCR3, TSTA3, GSDMD, TIGD5, NAPRT* |
| TKHS190002648 | NEC Cervix | DUP | chr8:143577269-144264831 | *MIR937, FAM83H-AS1, CCDC166, MIR4664, CYC1, EEF1D, MAPK15, PUF60, SCRIB, OPLAH, ZNF707, BREA2, FAM83H, GRINA, NRBP2, WDR97, SPATC1, HGH1, PLEC, EXOSC4, PYCR3, MIR661, TSTA3, MROH1, SHARPIN, EPPK1, MAF1, PARP10, TIGD5, GPAA1, NAPRT, ZNF623* |
| TKHS190002642 | NEC Ovary | DUP | chr8:143608062-143927992 | *MIR937, FAM83H-AS1, CCDC166, MIR4664, MAPK15, PUF60, SCRIB, ZNF707, BREA2, FAM83H, NRBP2, PLEC, PYCR3, TSTA3, EPPK1, ZNF623* |
| TKHS190002643 | NEC Cervix | DEL | chr8:143691073-143932032 | *MIR937, FAM83H-AS1, CCDC166, MIR4664, MAPK15, PUF60, SCRIB, ZNF707, BREA2, FAM83H, NRBP2, PLEC, EPPK1* |
| TKHS190002654 | NEC Cervix | DEL | chr8:143716378-143730582 | *MAPK15, FAM83H* |
| TKHS190002653 | NEC Ovary | DUP | chr8:143812817-144096029 | *MIR937, CYC1, PUF60, SCRIB, OPLAH, GRINA, NRBP2, SPATC1, PLEC, EXOSC4, MIR661, EPPK1, PARP10, GPAA1* |
| TKHS190002641 | NEC Ovary | DEL | chr8:143840706-143932032 | *NRBP2, PLEC, EPPK1* |
| TKHS190002654 | NEC Cervix | DEL | chr8:143926977-143946392 | *PLEC, MIR661* |
| TKHS190002643 | NEC Cervix | DEL | chr8:143983167-144200541 | *CYC1, OPLAH, GRINA, WDR97, SPATC1, HGH1, EXOSC4, MROH1, SHARPIN, MAF1, PARP10, GPAA1* |
| TKHS190002642 | NEC Ovary | DUP | chr8:143983167-144517190 | *MIR939, MIR1234, MFSD3, CYC1, ADCK5, BOP1, FBXL6, OPLAH, GPT, GRINA, CPSF1, HSF1, WDR97, TMEM249, SPATC1, TONSL, CYHR1, VPS28, HGH1, EXOSC4, SLC39A4, SCX, MROH1, SLC52A2, SHARPIN, SCRT1, MAF1, PARP10, PPP1R16A, DGAT1, GPAA1, FOXH1, KIFC2, RECQL4* |
| TKHS190002641 | NEC Ovary | DEL | chr8:144264225-144475756 | *MIR939, MIR1234, ADCK5, BOP1, FBXL6, CPSF1, HSF1, TMEM249, TONSL, CYHR1, VPS28, HGH1, SLC39A4, SCX, SLC52A2, SCRT1, DGAT1, FOXH1, KIFC2* |
| TKHS190002648 | NEC Cervix | DUP | chr8:144497195-144517190 | *MFSD3, GPT, PPP1R16A, RECQL4* |
| TKHS190002642 | NEC Ovary | DUP | chr8:144537721-144946206 | *COMMD5, ZNF517, ZNF250, RPL8, ZNF7, ZNF16, ARHGAP39, ZNF34, ZNF251* |
| TKHS190002648 | NEC Cervix | DUP | chr8:144580846-145054157 | *ZNF252P, TMED10P1, ZNF252P-AS1, COMMD5, ZNF517, ZNF250, RPL8, C8orf33, ZNF7, ZNF16, ARHGAP39, ZNF34, ZNF251* |
| TKHS190002653 | NEC Ovary | DUP | chr8:144580846-145054157 | *ZNF252P, TMED10P1, ZNF252P-AS1, COMMD5, ZNF517, ZNF250, RPL8, C8orf33, ZNF7, ZNF16, ARHGAP39, ZNF34, ZNF251* |
| TKHS190002648 | NEC Cervix | DUP | chr8:17299862-20220402 | *NAT2, LOC100128993, FGL1, PSD3, LPL, ASAH1, PCM1, PDGFRL, ATP6V1B2, INTS10, CSGALNACT1, MTUS1, SH2D4A, SLC7A2, SLC18A1, NAT1, MTMR7* |
| TKHS190002644 | NEC Cervix | DEL | chr8:21702805-22211728 | *NPM2, LGI3, DMTN, XPO7, GFRA2, HR, SFTPC, FAM160B2, BMP1, NUDT18, REEP4, FGF17, DOK2* |
| TKHS190002648 | NEC Cervix | DUP | chr8:21702805-25410202 | *LOC100507156, SORBS3, NPM2, ADAM28, NKX2-6, PEBP4, EGR3, R3HCC1, LGI3, DMTN, XPO7, RHOBTB2, SLC39A14, LOC254896, GFRA2, ADAMDEC1, LOC286059, LOC389641, LOXL2, MIR320A, NEFM, NEFL, NKX3-1, SLC25A37, C8orf58, PIWIL2, PPP2R2A, PPP3CC, HR, BIN3, CCAR2,* |

| | | | | |
|---|---|---|---|---|
| | | | | *PDLIM2, SFTPC, FAM160B2, BMP1, POLR3D, STC1, NUDT18, DOCK5, BIN3-IT1, REEP4, ADAM7, TNFRSF10D, TNFRSF10C, TNFRSF10B, TNFRSF10A, FGF17, DOK2, CHMP7, ENTPD4, PHYHIP* |
| TKHS190002643 | NEC Cervix | DEL | chr8:22120342-22138793 | *HR, REEP4* |
| TKHS190002653 | NEC Ovary | DUP | chr8:22554507-22616248 | *SORBS3, C8orf58, CCAR2, PDLIM2* |
| TKHS190002650 | NEC Ovary | DEL | chr8:23005960-23197204 | *RHOBTB2, LOC254896, LOC286059, TNFRSF10D, TNFRSF10C, TNFRSF10B, TNFRSF10A* |
| TKHS190002648 | NEC Cervix | DUP | chr8:25861309-29263034 | *MIR4287, MIR4288, MIR3622A, MIR3622B, PNMA2, CHRNA2, CLU, ADRA1A, ESCO2, FBXO16, DPYSL2, EPHX2, EXTL3, PTK2B, TRIM35, KIF13B, SCARA5, NUGGC, SCARA3, PNOC, ELP3, PPP2R2A, CCDC25, INTS9, PBK, ZNF395, EBF2, BNIP3L, HMBOX1, FZD3, STMN4* |
| TKHS190002644 | NEC Cervix | DEL | chr8:26770149-27538692 | *CHRNA2, ADRA1A, EPHX2, PTK2B, TRIM35, STMN4* |
| TKHS190002643 | NEC Cervix | DEL | chr8:27239259-27525473 | *CHRNA2, EPHX2, PTK2B, TRIM35, STMN4* |
| TKHS190002650 | NEC Ovary | DEL | chr8:27397585-27544244 | *CHRNA2, EPHX2, PTK2B* |
| TKHS190002648 | NEC Cervix | DUP | chr8:30065935-30545175 | *RBPMS-AS1, MIR548O2, DCTN6, RBPMS, LEPROTL1, SARAF, MBOAT4* |
| TKHS190002648 | NEC Cervix | DUP | chr8:32548727-33597515 | *NRG1, DUSP26, RNF122, TTI2, MAK16, FUT10* |
| TKHS190002648 | NEC Cervix | DUP | chr8:3284144-4637558 | *CSMD1* |
| TKHS190002648 | NEC Cervix | DUP | chr8:37695980-39274350 | *ERLIN2, PLPBP, GOT1L1, LETM2, ADRB3, EIF4EBP1, HTRA4, ADAM32, FGFR1, DDHD2, ADGRA2, LSM1, RNF5P1, C8orf86, NSD3, BRF2, PLEKHA2, STAR, TACC1, LOC728024, ZNF703, RAB11FIP1, TM2D2, PLPP5, ADAM9, ASH2L, BAG4* |
| TKHS190002648 | NEC Cervix | DUP | chr8:39987871-40674931 | *IDO2, TCIM, ZMAT4* |
| TKHS190002648 | NEC Cervix | DUP | chr8:413080-3052647 | *TDRP, ERICH1, CLN8, FBXO25, LOC286083, NA, CSMD1, MIR596, MYOM2, DLGAP2, ARHGEF10, KBTBD11* |
| TKHS190002648 | NEC Cervix | DUP | chr8:41609655-42850997 | *AP3M2, CHRNB3, SMIM19, GPAT4, NKX6-3, DKK4, ANK1, IKBKB, PLAT, POLB, THAP1, MIR486-1, SLC20A2, VDAC3, KAT6A, RNF170, CHRNA6* |
| TKHS190002648 | NEC Cervix | DUP | chr8:47279862-49075247 | *CEBPD, SPIDR, MCM4, PPDPFL, PRKDC, SNAI2, UBE2V2, EFCAB1* |
| TKHS190002643 | NEC Cervix | DEL | chr8:47779004-47854214 | *PRKDC* |
| TKHS190002644 | NEC Cervix | DEL | chr8:52136590-52180312 | *ST18* |
| TKHS190002648 | NEC Cervix | DUP | chr8:52137421-52180312 | *ST18* |
| TKHS190002648 | NEC Cervix | DUP | chr8:54135284-56316347 | *SBF1P1, XKR4, TMEM68, SDR16C5, SNORD54, MRPL15, LYN, MOS, PLAG1, RP1, RPS20, SOX17, CHCHD7, TGS1* |
| TKHS190002648 | NEC Cervix | DUP | chr8:58585898-61607322 | *LOC100130298, CLVS1, ASPH, CHD7, RAB2A, CA8, NSMAF, TOX* |
| TKHS190002648 | NEC Cervix | DUP | chr8:65708976-66516076 | *LINC00967, RRS1-AS1, ADHFE1, CRH, RRS1, VXN, PDE7A, TRIM55, DNAJC5B, MTFR1* |
| TKHS190002648 | NEC Cervix | DUP | chr8:66612819-66841110 | *C8orf44-SGK3, SGK3, PTTG3P, MYBL1, C8orf44, VCPIP1* |
| TKHS190002648 | NEC Cervix | DUP | chr8:67056519-67251450 | *ARFGEF1, COPS5, CSPP1* |
| TKHS190002648 | NEC Cervix | DUP | chr8:6747670-7056697 | *GS1-24F4.2, DEFA1, DEFA4, DEFA5, DEFA6, DEFB1, DEFT1P, XKR5, DEFA8P, DEFA9P, DEFA10P, AGPAT5, DEFA11P, DEFA1B* |
| TKHS190002648 | NEC Cervix | DUP | chr8:68083240-72055600 | *MSC-AS1, LINC01592, NCOA2, C8orf34, EYA1, SULF1, TRAM1, C8orf34-AS1, LACTB2-AS1, XKR9, LACTB2, PRDM14, PREX2, SLCO5A1, TRPA1, MSC* |
| TKHS190002648 | NEC Cervix | DUP | chr8:74825250-75016705 | *PI15, CRISPLD1* |
| TKHS190002648 | NEC Cervix | DUP | chr8:80050445-80993227 | *MIR5708, PAG1, ZNF704, ZBTB10, TPD52* |
| TKHS190002654 | NEC Cervix | DEL | chr8:86468676-86626080 | *RMDN1, CNGB3, CPNE3* |
| TKHS190002648 | NEC Cervix | DUP | chr8:86486484-86579252 | *RMDN1, CNGB3, CPNE3* |
| TKHS190002654 | NEC Cervix | DEL | chr8:94372170-94512089 | *FSBP, RAD54B, VIRMA* |

| | | | | |
|---|---|---|---|---|
| TKHS190002648 | NEC Cervix | DUP | chr8:9720374-9772422 | *MIR597, TNKS* |
| TKHS190002648 | NEC Cervix | DUP | chr8:99135015-99442635 | *VPS13B* |
| TKHS190002648 | NEC Cervix | DEL | chr9:115052740-115091018 | *TNC* |
| TKHS190002643 | NEC Cervix | DUP | chr9:121282452-121321401 | *GSN* |
| TKHS190002648 | NEC Cervix | DEL | chr9:121301963-121348149 | *STOM, GSN* |
| TKHS190002644 | NEC Cervix | DUP | chr9:121318406-121348149 | *STOM, GSN* |
| TKHS190002648 | NEC Cervix | DEL | chr9:122378433-122392544 | *PTGS1* |
| TKHS190002643 | NEC Cervix | DUP | chr9:127168679-127212719 | *RALGPS1* |
| TKHS190002648 | NEC Cervix | DEL | chr9:127168679-127218769 | *RALGPS1* |
| TKHS190002648 | NEC Cervix | DEL | chr9:127407112-127473931 | *SNORA65, SLC2A8, RPL12, ZNF79, LRSAM1* |
| TKHS190002648 | NEC Cervix | DEL | chr9:127707032-127896332 | *SH2D3C, MIR3960, MIR4672, CDK9, PTRH1, TTC16, ENG, AK1, FPGS, TOR2A, CFAP157, ST6GALNAC6* |
| TKHS190002657 | NEC Ovary | DUP | chr9:127808800-127824974 | *ENG, FPGS* |
| TKHS190002648 | NEC Cervix | DEL | chr9:128265788-128525423 | *MIR219B, SLC27A4, TRUB2, GLE1, GOLGA2, SWI5, MIR219A2, ODF2, COQ4, CERCAM, URM1* |
| TKHS190002648 | NEC Cervix | DEL | chr9:128604326-128684007 | *SET, SPTAN1, WDR34* |
| TKHS190002648 | NEC Cervix | DEL | chr9:128705734-128949243 | *LOC100506100, ZER1, ENDOG, DOLK, NUP188, PHYHD1, PKN3, SPOUT1, TBC1D13, LRRC8A, ZDHHC12, KYAT1* |
| TKHS190002648 | NEC Cervix | DEL | chr9:129098249-129108077 | *CRAT* |
| TKHS190002648 | NEC Cervix | DEL | chr9:129612347-129807363 | *ASB6, TOR1B, NTMT1, C9orf50, PRRX2, PTGES* |
| TKHS190002648 | NEC Cervix | DEL | chr9:129858050-129878440 | *USP20* |
| TKHS190002650 | NEC Ovary | DUP | chr9:129868005-129878440 | *USP20* |
| TKHS190002648 | NEC Cervix | DEL | *chr9:130407571-130422726* | |
| TKHS190002648 | NEC Cervix | DEL | chr9:131518445-131650242 | *POMT1, RAPGEF1, UCK1* |
| TKHS190002654 | NEC Cervix | DUP | chr9:132166832-132241044 | *NTNG2* |
| TKHS190002654 | NEC Cervix | DUP | chr9:133514636-133644320 | *DBH, MYMK, FAM163B, ADAMTSL2* |
| TKHS190002646 | NEC Cervix | DEL | chr9:133568603-133734173 | *DBH-AS1, DBH, SARDH, FAM163B, ADAMTSL2* |
| TKHS190002654 | NEC Cervix | DUP | chr9:134690912-134796901 | *COL5A1* |
| TKHS190002648 | NEC Cervix | DEL | chr9:134883302-135522177 | *PPP1R26-AS1, OLFM1, C9orf116, C9orf62, FCN1, FCN2, LCN1, LOC401557, MRPS2, PPP1R26* |
| TKHS190002648 | NEC Cervix | DEL | chr9:135742772-136745914 | *MIR4673, MIR4674, UBAC1, AGPAT2, ENTR1, NACC2, DIPK1B, CAMSAP1, C9orf163, LCN6, QSOX2, PMPCA, GPSM1, DKFZP434A062, MIR126, LCN10, NOTCH1, EGFL7, INPP5E, KCNT1, CARD9, SNAPC4, SNORA17A, SNORA17B, DNLZ, LHX3, SNHG7, TMEM250, SEC16A* |
| TKHS190002653 | NEC Ovary | DEL | chr9:136899594-137051129 | *ABCA2, FUT7, LCN12, PAXX, LCNL1, C9orf139, FBXW5, NPDC1, PTGDS, TRAF2, C8G, CLIC3, ENTPD2* |
| TKHS190002653 | NEC Ovary | DEL | chr9:137158379-137233722 | *MIR3621, SLC34A3, NDOR1, TPRN, GRIN1, ANAPC2, CYSRT1, LRRC26, RNF224, RNF208, SSNA1, TMEM203* |
| TKHS190002657 | NEC Ovary | DUP | chr9:137167774-137175837 | *MIR3621, GRIN1, ANAPC2, LRRC26* |
| TKHS190002657 | NEC Ovary | DUP | chr9:137233004-137252525 | *STPG3-AS1, TUBB4B, SLC34A3, FAM166A, STPG3* |
| TKHS190002648 | NEC Cervix | DEL | chr9:137493013-137577603 | *PNPLA7, MRPL41, DPH7* |
| TKHS190002648 | NEC Cervix | DEL | chr9:137957595-138118081 | *CACNA1B* |

| | | | | |
|---|---|---|---|---|
| TKHS190002654 | NEC Cervix | DEL | chr9:33441990-33466709 | *AQP3, NOL6* |
| TKHS190002644 | NEC Cervix | DUP | chr9:33441990-33467262 | *AQP3, NOL6* |
| TKHS190002643 | NEC Cervix | DUP | chr9:34636997-34648456 | *SIGMAR1, GALT* |
| TKHS190002643 | NEC Cervix | DUP | chr9:35074108-35079524 | *FANCG* |
| TKHS190002643 | NEC Cervix | DUP | chr9:35658576-35675567 | *CCDC107, CA9, ARHGEF39* |
| TKHS190002643 | NEC Cervix | DUP | chr9:35739628-35750741 | *GBA2, RGP1* |
| TKHS190002654 | NEC Cervix | DUP | chrX:103078560-103785768 | *RAB40A, TMEM31, BEX3, TCEAL5, MORF4L2-AS1, GLRA4, TCEAL9, RAB9B, PLP1, NXF3, BEX4, TCEAL7, TCEAL4, BEX2, TCEAL3, TCEAL8, TCEAL1, MORF4L2* |
| TKHS190002648 | NEC Cervix | DEL | chrX:152850265-153058544 | *PNMA5, PNMA3, NSDHL, ZNF185* |
| TKHS190002657 | NEC Ovary | DUP | chrX:152922720-153797814 | *MIR718, MIR3202-2, MIR3202-1, BCAP31, PNCK, TEX28, DNASE1L1, EMD, ABCD1, FLNA, G6PD, OPN1MW, SRPK3, GDI1, SNORA70, HCFC1, IDH3G, IRAK1, L1CAM, ARHGAP4, MECP2, PLXNB3, ATP6AP1, AVPR2, PLXNA3, PDZD4, OPN1LW, RENBP, FAM3A, RPL10, SLC6A8, SSR4, TAZ, NAA10, UBL4A, TMEM187, LAGE3, SLC10A3, TKTL1, IKBKG, FAM50A* |
| TKHS190002646 | NEC Cervix | DEL | chrX:153454394-154018787 | *MIR3202-2, MIR3202-1, BCAP31, PNCK, DUSP9, ABCD1, SRPK3, HCFC1, IDH3G, IRAK1, L1CAM, ARHGAP4, ATP2B3, PLXNB3, AVPR2, HAUS7, PDZD4, RENBP, BGN, SLC6A8, SSR4, NAA10, TMEM187, CCNQ* |
| TKHS190002648 | NEC Cervix | DEL | chrX:153457137-153580298 | *ATP2B3, HAUS7, BGN* |
| TKHS190002654 | NEC Cervix | DUP | chrX:153808342-153913051 | *L1CAM, ARHGAP4, AVPR2, PDZD4* |
| TKHS190002648 | NEC Cervix | DEL | chrX:153905532-154546163 | *MIR718, MIR3202-2, MIR3202-1, TEX28, DNASE1L1, EMD, FLNA, G6PD, OPN1MW, GDI1, SNORA70, HCFC1, IRAK1, ARHGAP4, MECP2, ATP6AP1, AVPR2, PLXNA3, OPN1LW, RENBP, FAM3A, RPL10, TAZ, NAA10, UBL4A, TMEM187, LAGE3, SLC10A3, TKTL1, IKBKG, FAM50A* |
| TKHS190002657 | NEC Ovary | DUP | chrX:153918832-154462310 | *MIR718, MIR3202-2, MIR3202-1, TEX28, DNASE1L1, EMD, FLNA, OPN1MW, GDI1, SNORA70, HCFC1, IRAK1, ARHGAP4, MECP2, ATP6AP1, PLXNA3, OPN1LW, RENBP, RPL10, TAZ, NAA10, TMEM187, TKTL1, FAM50A* |
| TKHS190002641 | NEC Ovary | DEL | chrX:154325339-154380019 | *EMD, FLNA, TKTL1* |
| TKHS190002646 | NEC Cervix | DEL | chrX:154325339-154463519 | *DNASE1L1, EMD, FLNA, GDI1, SNORA70, ATP6AP1, PLXNA3, RPL10, TAZ, TKTL1, FAM50A* |
| TKHS190002643 | NEC Cervix | DUP | chrX:154435139-154442428 | *GDI1, ATP6AP1* |
| TKHS190002657 | NEC Ovary | DUP | chrX:154468823-154546163 | *G6PD, PLXNA3, FAM3A, UBL4A, LAGE3, SLC10A3, IKBKG* |
| TKHS190002654 | NEC Cervix | DUP | chrX:15517929-15537305 | *BMX* |
| TKHS190002654 | NEC Cervix | DUP | chrX:40049916-40076962 | *BCOR* |
| TKHS190002643 | NEC Cervix | DUP | chrX:47199048-47241699 | *CDK16, UBA1, USP11, INE1* |
| TKHS190002646 | NEC Cervix | DEL | chrX:48559614-48706627 | *TBC1D25, RBM3, WDR13, SUV39H1, WAS* |
| TKHS190002646 | NEC Cervix | DEL | chrX:49065454-49193463 | *WDR45, PRAF2, GPKOW, PRICKLE3, PLP2, SYP, MAGIX, CCDC120* |
| TKHS190002645 | NEC Endometrium | DUP | chrX:50285164-50404184 | *DGKK, CCNB3* |
| TKHS190002648 | NEC Cervix | DEL | chrX:53193437-53215976 | *KDM5C* |
| TKHS190002648 | NEC Cervix | DEL | chrX:71223708-71290791 | *GJB1, NONO, BCYRN1, ZMYM3* |
| TKHS190002650 | NEC Ovary | DUP | chrX:71247734-71253255 | *BCYRN1, ZMYM3* |
| TKHS190002654 | NEC Cervix | DUP | chrX:72129831-72143574 | *RTL5, NHSL2* |
| TKHS190002646 | NEC Cervix | DEL | chrX:9894358-10474750 | *CLDN34, CLCN4, SHROOM2, MID1, WWC3* |
