## Supplementary Table 2 for "Genomic analyses of high-grade neuroendocrine gynecological malignancies reveal a unique mutational landscape and therapeutic vulnerabilities"

**Supplementary Table 2.** List of cancer driver genes detected based on positional clustering

| Gene | Total Mutations | Mutated Samples | Clusters | Mutations in clusters | Z-score | p-value | FDR | Fraction of mutations in clusters |
|---|---|---|---|---|---|---|---|---|
| *CCDC6* | 13 | 13 | 1 | 13 | 5.54615385 | 1.46E-08 | 8.40E-08 | 1 |
| *CLTCL1* | 18 | 12 | 3 | 18 | 5.54615385 | 1.46E-08 | 8.40E-08 | 1 |
| *LATS2* | 15 | 12 | 2 | 15 | 5.54615385 | 1.46E-08 | 8.40E-08 | 1 |
| *RNF43* | 17 | 12 | 3 | 17 | 5.54615385 | 1.46E-08 | 8.40E-08 | 1 |
| *KMT2C* | 59 | 14 | 8 | 54 | 4.82907432 | 6.86E-07 | 3.15E-06 | 0.91525424 |
| *CDH11* | 10 | 9 | 2 | 9 | 4.77692308 | 8.90E-07 | 3.41E-06 | 0.9 |
| *PTPRT* | 16 | 12 | 2 | 14 | 4.58461539 | 2.27E-06 | 7.47E-06 | 0.875 |
| *BLM* | 7 | 7 | 2 | 6 | 4.44725275 | 4.35E-06 | 1.25E-05 | 0.85714286 |
| *NCOR2* | 34 | 14 | 2 | 26 | 4.23278084 | 1.15E-05 | 2.95E-05 | 0.76470588 |
| *BRCA1* | 5 | 5 | 2 | 4 | 4.00769231 | 3.07E-05 | 7.05E-05 | 0.8 |
| *KNL1* | 29 | 14 | 3 | 23 | 3.95464191 | 3.83E-05 | 8.01E-05 | 0.79310345 |
| *FAT1* | 14 | 11 | 3 | 11 | 3.8978022 | 4.85E-05 | 9.30E-05 | 0.78571429 |
| *BRCA2* | 15 | 9 | 3 | 11 | 3.4948718 | 0.00023715 | 0.00041956 | 0.73333333 |
| *TP63* | 8 | 4 | 2 | 5 | 3.34835165 | 0.00040647 | 0.00066777 | 0.625 |
| *CTCF* | 5 | 4 | 1 | 3 | 2.46923077 | 0.00677019 | 0.01038096 | 0.6 |
| *TET1* | 7 | 7 | 1 | 4 | 2.24945055 | 0.01224192 | 0.01759776 | 0.57142857 |
| *TP53* | 5 | 4 | 1 | 2 | 1.7 | 0.04456546 | 0.06029445 | 0.4 |
| *FAT4* | 16 | 12 | 1 | 7 | 1.21923077 | 0.11137831 | 0.14231673 | 0.4375 |
| *ASXL1* | 6 | 5 | 1 | 2 | 0.93076923 | 0.17598648 | 0.21303626 | 0.33333333 |
| *CIC* | 6 | 5 | 1 | 2 | 0.41794872 | 0.33799231 | 0.38869115 | 0.33333333 |
| *KMT2D* | 11 | 7 | 1 | 2 | 0.05164835 | 0.47940445 | 0.50119556 | 0.18181818 |
| *LRP1B* | 7 | 5 | 1 | 2 | 0.05164835 | 0.47940445 | 0.50119556 | 0.28571429 |
| *ZFHX3* | 10 | 8 | 1 | 3 | -0.0094017 | 0.50375068 | 0.50375068 | 0.3 |
