## Supplementary Table 3 for "Genomic analyses of high-grade neuroendocrine gynecological malignancies reveal a unique mutational landscape and therapeutic vulnerabilities"

**Supplementary Table 3.** List of fusion genes

| Fusion genes | % |
|---|---|
| *MALAT1--SMG1* | 53.8 |
| *AD000090.1--MALAT1* | 53.8 |
| *AC098591.3--RLIM* | 46.2 |
| *AC099535.1--UBA52* | 38.5 |
| *EEF1A1--MALAT1* | 30.8 |
| *ACTB--MALAT1* | 30.8 |
| *ASH1L--YY1AP1* | 30.8 |
| *DYNC1H1--MALAT1* | 23.1 |
| *AC091807.1--IMPDH1* | 23.1 |
| *AC007621.1--UBE2V1* | 23.1 |
| *MALAT1--TPT1* | 23.1 |
| *XIST--MALAT1* | 23.1 |
| *MALAT1--XIST* | 23.1 |
| *MALAT1--RNF213* | 23.1 |
| *MALAT1--RMRP* | 23.1 |
| *MALAT1--FAT1* | 23.1 |
| *CDK6--MALAT1* | 23.1 |
| *AHNAK--MALAT1* | 23.1 |
| *AC091951.3--HERC2* | 23.1 |
| *ZNF480--ZNF665* | 15.4 |
| *MALAT1--ZNF638* | 15.4 |
| *MALAT1--VPS13A* | 15.4 |
| *MALAT1--TCF4* | 15.4 |
| *MALAT1--SPTBN1* | 15.4 |
| *MALAT1--RPSA* | 15.4 |
| *KAT6A--MALAT1* | 15.4 |
| *HNRNPU--MALAT1* | 15.4 |
| *DST--MALAT1* | 15.4 |
| *CR381653.1--ZNF717* | 15.4 |
| *CLTC--MALAT1* | 15.4 |
| *CCT3--MALAT1* | 15.4 |
| *AC093878.1--CAMSAP1* | 15.4 |
| *AC024270.1--FABP5* | 15.4 |
| *UBC--MALAT1* | 15.4 |
| *SET--LMOD3* | 15.4 |
| *RPPH1--ACTB* | 15.4 |
| *RANBP2--MALAT1* | 15.4 |
| *PARG--AC022400.4* | 15.4 |
| *NEAT1--XIST* | 15.4 |
| *NAP1L1--MALAT1* | 15.4 |
| *MROH1--MALAT1* | 15.4 |
| *MALAT1--VMP1* | 15.4 |
| *MALAT1--TTC3* | 15.4 |
| *MALAT1--TNRC6A* | 15.4 |
| *MALAT1--TAOK1* | 15.4 |
| *MALAT1--SYNE2* | 15.4 |
| *MALAT1--SREBF2* | 15.4 |
| *MALAT1--SMC3* | 15.4 |
| *MALAT1--ROCK1* | 15.4 |
| *MALAT1--RGPD2* | 15.4 |
| *MALAT1--RDX* | 15.4 |
| *MALAT1--RALGAPA1* | 15.4 |
| *MALAT1--POLR2A* | 15.4 |
| *MALAT1--PCM1* | 15.4 |
| *MALAT1--NKTR* | 15.4 |
| *MALAT1--KIAA1328* | 15.4 |
| *MALAT1--JAK2* | 15.4 |
| *MALAT1--JAK1* | 15.4 |
| *MALAT1--HNRNPU* | 15.4 |
| *MALAT1--HNRNPA3* | 15.4 |
| *MALAT1--HNRNPA2B1* | 15.4 |
| *MALAT1--EEF1A1* | 15.4 |
| *MALAT1--CP* | 15.4 |
| *MALAT1--COL1A2* | 15.4 |
| *MALAT1--CKAP5* | 15.4 |
| *MALAT1--CANX* | 15.4 |
| *MALAT1--BIRC6* | 15.4 |
| *MALAT1--ALK* | 15.4 |
| *MALAT1--ACTG1* | 15.4 |
| *MALAT1--ACIN1* | 15.4 |
| *LNPEP--MALAT1* | 15.4 |
| *LINC02241--C1QTNF3-AMACR* | 15.4 |
| *LINC01923--NDUFV1* | 15.4 |
| *IL9R--AL591424.1* | 15.4 |
| *HSP90AA1--MALAT1* | 15.4 |
| *HNRNPH1--MALAT1* | 15.4 |
| *FOXP1--MALAT1* | 15.4 |
| *EEF1A1--NEAT1* | 15.4 |
| *COL3A1--MALAT1* | 15.4 |
| *CNOT4--MALAT1* | 15.4 |
| *CANX--MALAT1* | 15.4 |
| *C1QTNF3-AMACR--LINC02241* | 15.4 |
| *BTN2A2--BTN2A3P* | 15.4 |
| *BRD4--MALAT1* | 15.4 |
| *BMS1P4-AGAP5--TIMM23B* | 15.4 |
| *AL591424.1--IL9R* | 15.4 |
| *AL589743.5--TOMM40* | 15.4 |
| *AL157871.3--NDUFB3* | 15.4 |
| *AC064847.1--ENO3* | 15.4 |
| *ZZEF1--FLNC* | 7.7 |
| *ZZEF1--ACP6* | 7.7 |
| *ZYG11A--KCNH7* | 7.7 |
| *ZSWIM4--INTS3* | 7.7 |
| *ZSCAN29--PHF3* | 7.7 |
| *ZSCAN29--FAM149A* | 7.7 |
| *ZRANB1--RPS14* | 7.7 |
| *ZRANB1--MUC4* | 7.7 |
| *ZPR1--DGKA* | 7.7 |
| *ZNRF3--RREB1* | 7.7 |
| *ZNF91--ZNF525* | 7.7 |
| *ZNF91--CBX5* | 7.7 |
| *ZNF850--AC010624.4* | 7.7 |
| *ZNF841--ZNF721* | 7.7 |
| *ZNF816-ZNF321P--AC023934.1* | 7.7 |
| *ZNF814--RSRP1* | 7.7 |
| *ZNF813--ELL2* | 7.7 |
| *ZNF8--DNM1L* | 7.7 |
| *ZNF790--TMEM176A* | 7.7 |
| *ZNF788P--RCAN3* | 7.7 |
| *ZNF782--OLA1* | 7.7 |
| *ZNF781--ZNF20* | 7.7 |
| *ZNF767P--LARS* | 7.7 |
| *ZNF766--ZNF415* | 7.7 |
| *ZNF763--ZNF442* | 7.7 |
| *ZNF761--ZNF808* | 7.7 |
| *ZNF737--NCOR1* | 7.7 |
| *ZNF721--ZNF716* | 7.7 |
| *ZNF721--TRAF7* | 7.7 |
| *ZNF721--MYH11* | 7.7 |
| *ZNF721--AC010332.2* | 7.7 |
| *ZNF714--YTHDF3* | 7.7 |
| *ZNF71--SIN3A* | 7.7 |
| *ZNF701--ZNF347* | 7.7 |
| *ZNF69--AC022415.2* | 7.7 |
| *ZNF682--PLEKHG5* | 7.7 |
| *ZNF680--ZNF471* | 7.7 |
| *ZNF664--TRMT12* | 7.7 |
| *ZNF664--IPO9* | 7.7 |
| *ZNF664--GCC2* | 7.7 |

| | |
|---|---|
| ZNF66--ACAP2 | 7.7 |
| ZNF658B--ZNF334 | 7.7 |
| ZNF654--GNAS | 7.7 |
| ZNF644--NPM1 | 7.7 |
| ZNF641--ASS1 | 7.7 |
| ZNF638--ZFP30 | 7.7 |
| ZNF638--TIMP2 | 7.7 |
| ZNF629--TARS2 | 7.7 |
| ZNF626--AC021451.1 | 7.7 |
| ZNF625--C12ORF57 | 7.7 |
| ZNF623--LCOR | 7.7 |
| ZNF609--HSP90B1 | 7.7 |
| ZNF609--ADCY1 | 7.7 |
| ZNF608--TMEM260 | 7.7 |
| ZNF608--MALAT1 | 7.7 |
| ZNF605--FN1 | 7.7 |
| ZNF594--TOPORS | 7.7 |
| ZNF592--LINC-PINT | 7.7 |
| ZNF586--ZNF417 | 7.7 |
| ZNF584--IFIH1 | 7.7 |
| ZNF57--AGL | 7.7 |
| ZNF561--DNAH7 | 7.7 |
| ZNF559--MALAT1 | 7.7 |
| ZNF558--TCF7L1 | 7.7 |
| ZNF551--ZNF530 | 7.7 |
| ZNF551--RGP1 | 7.7 |
| ZNF530--KCNMB3 | 7.7 |
| ZNF526--PPP1R12B | 7.7 |
| ZNF518B--ARL6IP1 | 7.7 |
| ZNF518A--RNF40 | 7.7 |
| ZNF460--DNAJC7 | 7.7 |
| ZNF438--ZBTB20 | 7.7 |
| ZNF436--MALAT1 | 7.7 |
| ZNF43--PYCR1 | 7.7 |
| ZNF428--MIR100HG | 7.7 |
| ZNF426--ZFP90 | 7.7 |
| ZNF426--SETX | 7.7 |
| ZNF41--XIST | 7.7 |
| ZNF397--ERGIC1 | 7.7 |
| ZNF383--ZNF420 | 7.7 |
| ZNF37A--COPS3 | 7.7 |
| ZNF37A--COG4 | 7.7 |
| ZNF33A--SYNE2 | 7.7 |
| ZNF337--ZNF586 | 7.7 |
| ZNF330--TTC3 | 7.7 |

| | |
|---|---|
| ZNF326--FAM193A | 7.7 |
| ZNF281--STIL | 7.7 |
| ZNF277--PCM1 | 7.7 |
| ZNF276--TSPYL5 | 7.7 |
| ZNF266--ZNF30 | 7.7 |
| ZNF24--ZNF852 | 7.7 |
| ZNF236--SCAF8 | 7.7 |
| ZNF234--CTCFL | 7.7 |
| ZNF224--PLSCR1 | 7.7 |
| ZNF217--ANKS1A | 7.7 |
| ZNF208--ZNF267 | 7.7 |
| ZNF208--HIPK2 | 7.7 |
| ZNF180--MALAT1 | 7.7 |
| ZNF169--VPS13C | 7.7 |
| ZNF154--ZNF530 | 7.7 |
| ZNF148--WDR76 | 7.7 |
| ZNF141--TMEM135 | 7.7 |
| ZNF141--FXYD5 | 7.7 |
| ZNF14--IGF2 | 7.7 |
| ZNF14--HNMT | 7.7 |
| ZNF137P--ZNF83 | 7.7 |
| ZNF121--ZNF473 | 7.7 |
| ZMYND8--C17ORF64 | 7.7 |
| ZMYND15--DNAH1 | 7.7 |
| ZMYND11--MALAT1 | 7.7 |
| ZMYM4--RBM5 | 7.7 |
| ZMYM2--MAP4K4 | 7.7 |
| ZMYM2--MALAT1 | 7.7 |
| ZMPSTE24--AC087190.3 | 7.7 |
| ZMIZ2--FBXO36-IT1 | 7.7 |
| ZMIZ1--XIST | 7.7 |
| ZMIZ1--RMRP | 7.7 |
| ZMIZ1--LAMA1 | 7.7 |
| ZMIZ1--CCDC14 | 7.7 |
| ZKSCAN1--MALAT1 | 7.7 |
| ZKSCAN1--ASCL1 | 7.7 |
| ZKSCAN1--AGO1 | 7.7 |
| ZIM2--SPTBN1 | 7.7 |
| ZIC5--MIIP | 7.7 |
| ZHX2--TIMP2 | 7.7 |
| ZFYVE16--UTRN | 7.7 |
| ZFP92--ZNF629 | 7.7 |
| ZFP90--ZNF433 | 7.7 |
| ZFP36L1--ATXN2 | 7.7 |
| ZFP30--ZNF420 | 7.7 |

| | |
|---|---|
| ZFAND6--TAF15 | 7.7 |
| ZFAND5--USP47 | 7.7 |
| ZEB2--IGH@ | 7.7 |
| ZDHHC11B--ARID2 | 7.7 |
| ZDBF2--CFAP410 | 7.7 |
| ZCCHC8--COX7B | 7.7 |
| ZC3H7A--FOXD1 | 7.7 |
| ZC3H14--PSME4 | 7.7 |
| ZC3H14--CCT7 | 7.7 |
| ZC3H13--RPS29 | 7.7 |
| ZC3H13--FOCAD | 7.7 |
| ZBTB8OS--TCP11L1 | 7.7 |
| ZBTB48--MALAT1 | 7.7 |
| ZBTB26--METTL8 | 7.7 |
| ZBTB21--DCAF10 | 7.7 |
| ZBTB20--COL3A1 | 7.7 |
| ZBTB12BP--ZBTB12 | 7.7 |
| ZBTB1--DNAH5 | 7.7 |
| ZBED9--SEL1L | 7.7 |
| ZBED6--ZSCAN31 | 7.7 |
| ZBED6--HIST1H1D | 7.7 |
| ZBED6--ERCC5 | 7.7 |
| Z83843.1--PEG3 | 7.7 |
| YY1AP1--COL6A2 | 7.7 |
| YWHAZ--SPTAN1 | 7.7 |
| YWHAZ--DST | 7.7 |
| YWHAG--TMSB4X | 7.7 |
| YWHAE--UBR4 | 7.7 |
| YWHAE--DIDO1 | 7.7 |
| YWHAE--CFAP44 | 7.7 |
| YWHAE--ALMS1 | 7.7 |
| YTHDF3--PISD | 7.7 |
| YTHDF3--ITGA9 | 7.7 |
| YTHDC1--PLXNA3 | 7.7 |
| YTHDC1--MUC4 | 7.7 |
| YTHDC1--GOLGB1 | 7.7 |
| YME1L1--TRAF6 | 7.7 |
| YME1L1--TBC1D32 | 7.7 |
| YME1L1--KRAS | 7.7 |
| YLPM1--YTHDF3 | 7.7 |
| YIPF6--STARD10 | 7.7 |
| YIPF5--HSPD1 | 7.7 |
| YES1--SURF4 | 7.7 |
| YES1--DOCK1 | 7.7 |
| YBX1--GNLY | 7.7 |

| | |
|---|---|
| *YAP1--ZMYM2* | 7.7 |
| *YAP1--AC004951.1* | 7.7 |
| *YAE1--DYNC1LI2* | 7.7 |
| *XRN2--TLN1* | 7.7 |
| *XRN2--MALAT1* | 7.7 |
| *XRCC6--SPATA13* | 7.7 |
| *XPOT--KAT6A* | 7.7 |
| *XPO5--CD63* | 7.7 |
| *XPO5--CALR* | 7.7 |
| *XPO5--ACTB* | 7.7 |
| *XPO1--SPAST* | 7.7 |
| *XPNPEP1--ADRM1* | 7.7 |
| *XIST--ZNF652* | 7.7 |
| *XIST--TP73-AS1* | 7.7 |
| *XIST--SECISBP2* | 7.7 |
| *XIST--PLXNB2* | 7.7 |
| *XIST--PLCG2* | 7.7 |
| *XIST--HMGN1* | 7.7 |
| *XIST--G6PD* | 7.7 |
| *XIST--EPS15* | 7.7 |
| *XIST--EEF1A1* | 7.7 |
| *XIST--CDC42* | 7.7 |
| *XIST--CCT3* | 7.7 |
| *XIST--AC004922.1* | 7.7 |
| *XIAP--EIF3J-DT* | 7.7 |
| *XIAP--BLOC1S5-TXNDC5* | 7.7 |
| *WWTR1--B2M* | 7.7 |
| *WWOX--ZNF267* | 7.7 |
| *WWOX--PSMA1* | 7.7 |
| *WWOX--FN1* | 7.7 |
| *WWOX--ACTG1* | 7.7 |
| *WWC3--SUN2* | 7.7 |
| *WTAP--ACTG1* | 7.7 |
| *WSB2--CACNA2D1* | 7.7 |
| *WRAP73--SKIL* | 7.7 |
| *WNK1--SMARCA2* | 7.7 |
| *WNK1--GTF2I* | 7.7 |
| *WIF1--CPLANE1* | 7.7 |
| *WDSUB1--AKR1B10* | 7.7 |
| *WDR82--PARP3* | 7.7 |
| *WDR74--UTRN* | 7.7 |
| *WDR74--ROCK1* | 7.7 |
| *WDR74--MUC4* | 7.7 |
| *WDR74--HNRNPUL2-BSCL2* | 7.7 |
| *WDR74--FTL* | 7.7 |
| *WDR74--EGR1* | 7.7 |
| *WDR74--DNAJC10* | 7.7 |
| *WDR74--DDX5* | 7.7 |
| *WDR74--AP005263.1* | 7.7 |
| *WDR74--AEBP1* | 7.7 |
| *WDR72--CCDC162P* | 7.7 |
| *WDR64--MIR100HG* | 7.7 |
| *WDR6--KLHL11* | 7.7 |
| *WDR6--GNAS* | 7.7 |
| *WDR6--CNTNAP1* | 7.7 |
| *WDR6--A4GNT* | 7.7 |
| *WDR45B--SILC1* | 7.7 |
| *WDR33--PHLDA1* | 7.7 |
| *WDR26--FOXJ3* | 7.7 |
| *WDR12--HAGHL* | 7.7 |
| *WDFY3--UBE3A* | 7.7 |
| *WDFY3--POLR3F* | 7.7 |
| *WASHC4--GGNBP2* | 7.7 |
| *WAPL--KLF3* | 7.7 |
| *WAC-AS1--MALAT1* | 7.7 |
| *VWF--LONP2* | 7.7 |
| *VWF--BCL2L11* | 7.7 |
| *VWA8--TIMM8A* | 7.7 |
| *VTI1B--GPR157* | 7.7 |
| *VSNL1--PAFAH1B1* | 7.7 |
| *VPS45--PHRF1* | 7.7 |
| *VPS39--GOLGA8R* | 7.7 |
| *VPS37A--NEB* | 7.7 |
| *VPS13C--EEF1A1* | 7.7 |
| *VPS13C--DDX27* | 7.7 |
| *VPS13A--TLE4* | 7.7 |
| *VPS13A--SPDL1* | 7.7 |
| *VPS13A--KIF2A* | 7.7 |
| *VPS13A--FAM193A* | 7.7 |
| *VMP1--CANX* | 7.7 |
| *VMP1--ASXL1* | 7.7 |
| *VMO1--NEB* | 7.7 |
| *VMA21--DNMT1* | 7.7 |
| *VKORC1L1--ARID1B* | 7.7 |
| *VIRMA--ZNF558* | 7.7 |
| *VIM--TMEM38B* | 7.7 |
| *VIM--POGZ* | 7.7 |
| *VIM--PIK3R2* | 7.7 |
| *VIM--MALAT1* | 7.7 |
| *VIM--KMT2C* | 7.7 |
| *VIM--EIF4A1* | 7.7 |
| *VIM--ANP32B* | 7.7 |
| *VIM--AD000090.1* | 7.7 |
| *VIL1--RBM33* | 7.7 |
| *VIL1--MALAT1* | 7.7 |
| *VIL1--CCT5* | 7.7 |
| *VEZT--GNA13* | 7.7 |
| *VCPIP1--SYNJ2BP-COX16* | 7.7 |
| *VCPIP1--KIAA0895* | 7.7 |
| *VCP--LMBR1* | 7.7 |
| *VCP--CCDC47* | 7.7 |
| *VCL--PDGFRA* | 7.7 |
| *VCAN--TAF15* | 7.7 |
| *VCAN--NR4A3* | 7.7 |
| *VAMP3--RAB11FIP5* | 7.7 |
| *VAMP3--CYP17A1* | 7.7 |
| *UTRN--MALAT1* | 7.7 |
| *UTRN--COL1A1* | 7.7 |
| *UTRN--ALKBH7* | 7.7 |
| *USP9X--MAPK8IP3* | 7.7 |
| *USP9X--IBTK* | 7.7 |
| *USP9X--GPR155* | 7.7 |
| *USP8--TMOD3* | 7.7 |
| *USP7--C2CD3* | 7.7 |
| *USP53--UBA52* | 7.7 |
| *USP40--MALAT1* | 7.7 |
| *USP37--ZSWIM5* | 7.7 |
| *USP37--FNDC3A* | 7.7 |
| *USP34--SOS1* | 7.7 |
| *USP34--SMG1* | 7.7 |
| *USP34--NBAS* | 7.7 |
| *USP34--GIGYF1* | 7.7 |
| *USP34--ARID1B* | 7.7 |
| *USP28--YLPM1* | 7.7 |
| *USP28--PTMA* | 7.7 |
| *USP24--GUSB* | 7.7 |
| *USP20--SERPINB6* | 7.7 |
| *USP15--RANBP3* | 7.7 |
| *USP1--PPIA* | 7.7 |
| *USH1C--TRA@* | 7.7 |
| *USF3--FNDC3B* | 7.7 |
| *URI1--PHF14* | 7.7 |
| *UQCRFS1--KHDC4* | 7.7 |

| | |
|---|---|
| UQCRB--TRA@ | 7.7 |
| UQCRB--PODXL | 7.7 |
| UQCC2--MALAT1 | 7.7 |
| UNG--NAPA | 7.7 |
| UNG--DYM | 7.7 |
| UNC80--PPP3CB | 7.7 |
| UNC80--MALAT1 | 7.7 |
| UMPS--DISP2 | 7.7 |
| ULK4--MRAS | 7.7 |
| ULK1--ZNF292 | 7.7 |
| UGT2B15--XIST | 7.7 |
| UGDH--DST | 7.7 |
| UFM1--CLCN7 | 7.7 |
| UFC1--HSPD1 | 7.7 |
| UCP2--AL021155.5 | 7.7 |
| UCHL5--AL390957.1 | 7.7 |
| UBXN4--VPS13A | 7.7 |
| UBR4--SYMPK | 7.7 |
| UBR4--RPS11 | 7.7 |
| UBR4--RPL26 | 7.7 |
| UBR4--MALAT1 | 7.7 |
| UBR4--CHD2 | 7.7 |
| UBR3--PANK2 | 7.7 |
| UBR3--CPLANE1 | 7.7 |
| UBR2--DDX17 | 7.7 |
| UBIAD1--STK24 | 7.7 |
| UBE4A--ENPP4 | 7.7 |
| UBE3A--RANBP2 | 7.7 |
| UBE3A--AC068580.4 | 7.7 |
| UBE2Z--MALAT1 | 7.7 |
| UBE2V1--AC007621.1 | 7.7 |
| UBE2Q2--PTPRD | 7.7 |
| UBE2D3--PHKG2 | 7.7 |
| UBC--ZZEF1 | 7.7 |
| UBC--ZFPM1 | 7.7 |
| UBC--VIM | 7.7 |
| UBC--SLC25A3 | 7.7 |
| UBC--PRRC2B | 7.7 |
| UBC--MARS | 7.7 |
| UBC--LMAN1 | 7.7 |
| UBC--FAM214A | 7.7 |
| UBC--CIITA | 7.7 |
| UBC--CAPZB | 7.7 |
| UBC--BAZ2B | 7.7 |
| UBC--ADAMTS12 | 7.7 |

| | |
|---|---|
| UBB--SYNPO2 | 7.7 |
| UBAP1--PLEKHM3 | 7.7 |
| UBAC1--ZFP36 | 7.7 |
| UBA52--PDPR | 7.7 |
| UBA52--AC099535.1 | 7.7 |
| UBA3--NEB | 7.7 |
| UBA2--LIFR | 7.7 |
| UBA1--LRBA | 7.7 |
| UBA1--BACH1 | 7.7 |
| UAP1--PRPF4B | 7.7 |
| TYK2--NEFH | 7.7 |
| TXNRD1--NF1 | 7.7 |
| TXNRD1--MYCBP2 | 7.7 |
| TXNRD1--IFT46 | 7.7 |
| TXNRD1--FTH1 | 7.7 |
| TXNL1--OARD1 | 7.7 |
| TXNIP--MUC4 | 7.7 |
| TXNIP--IGK@ | 7.7 |
| TXN--MALAT1 | 7.7 |
| TXLNA--PRKAG1 | 7.7 |
| TWSG1--HSPD1 | 7.7 |
| TUT7--VARS | 7.7 |
| TUT4--ADCY9 | 7.7 |
| TULP3--UNC13B | 7.7 |
| TUBGCP4--SLC44A1 | 7.7 |
| TUBB6--CDV3 | 7.7 |
| TUBA1C--SMC4 | 7.7 |
| TUBA1B--CHST11 | 7.7 |
| TTYH3--IGF2BP1 | 7.7 |
| TTN--TRA@ | 7.7 |
| TTN--TCF4 | 7.7 |
| TTN--SMC2 | 7.7 |
| TTN--RALY | 7.7 |
| TTI2--MIR100HG | 7.7 |
| TTF2--FMNL3 | 7.7 |
| TTC3--RMRP | 7.7 |
| TTC28--ULK4 | 7.7 |
| TTC21B--EPAS1 | 7.7 |
| TTC14--INSR | 7.7 |
| TSPYL1--MAPK8IP3 | 7.7 |
| TSPAN14--ERCC1 | 7.7 |
| TSN--VPS13B | 7.7 |
| TSIX--PHC1 | 7.7 |
| TSHZ2--INF2 | 7.7 |
| TSG101--ATG16L2 | 7.7 |

| | |
|---|---|
| TSEN34--PLCE1 | 7.7 |
| TSC22D2--XIST | 7.7 |
| TSC2--RPL31 | 7.7 |
| TRPV4--ARHGAP33 | 7.7 |
| TRPV1--WIZ | 7.7 |
| TRNAU1AP--CALR | 7.7 |
| TRIT1--VPS13D | 7.7 |
| TRIP12--IL13RA1 | 7.7 |
| TRIP12--FYCO1 | 7.7 |
| TRIP12--EIF3B | 7.7 |
| TRIOBP--UBA1 | 7.7 |
| TRIOBP--MALAT1 | 7.7 |
| TRIOBP--IGF1R | 7.7 |
| TRIOBP--ARHGAP35 | 7.7 |
| TRIO--CDKAL1 | 7.7 |
| TRIO--ATXN2L | 7.7 |
| TRIO--AP000295.1 | 7.7 |
| TRIM52--ARIH2 | 7.7 |
| TRIM44--ZNF607 | 7.7 |
| TRIM25--EML4 | 7.7 |
| TRIM2--FTH1 | 7.7 |
| TRIM13--ARL3 | 7.7 |
| TRIL--FN1 | 7.7 |
| TRIB3--JAG1 | 7.7 |
| TRG@--DNAH14 | 7.7 |
| TRAPPC9--GABBR1 | 7.7 |
| TRAPPC8--RALGAPA1 | 7.7 |
| TRAPPC8--PPIP5K1 | 7.7 |
| TRAPPC3--GTF2F1 | 7.7 |
| TRAPPC10--TCF12 | 7.7 |
| TRAP1--STT3B | 7.7 |
| TRANK1--INO80B-WBP1 | 7.7 |
| TRAM1--HDAC7 | 7.7 |
| TRAK2--SLC25A44 | 7.7 |
| TRAK1--RAD54L | 7.7 |
| TRAK1--CSF1R | 7.7 |
| TRAIP--SPTAN1 | 7.7 |
| TRAF3--TRPC4AP | 7.7 |
| TRA@--SYNE2 | 7.7 |
| TRA@--GOSR1 | 7.7 |
| TRA@--ANO10 | 7.7 |
| TRA2B--C17ORF113 | 7.7 |
| TRA2A--MALAT1 | 7.7 |
| TPT1--PCF11 | 7.7 |

| | |
|---|---|
| *TPT1--MCPH1* | 7.7 |
| *TPT1--AAK1* | 7.7 |
| *TPR--MYH9* | 7.7 |
| *TPR--CKAP5* | 7.7 |
| *TPP2--HTT* | 7.7 |
| *TPP2--EHD2* | 7.7 |
| *TPP2--AGL* | 7.7 |
| *TPM3--MALAT1* | 7.7 |
| *TPM3--CCAR1* | 7.7 |
| *TPM2--TAOK1* | 7.7 |
| *TPM2--RCHY1* | 7.7 |
| *TPM1--WDR60* | 7.7 |
| *TPM1--QSER1* | 7.7 |
| *TPI1--TPMT* | 7.7 |
| *TPI1--RMRP* | 7.7 |
| *TPI1--CPD* | 7.7 |
| *TPI1--ANKRD6* | 7.7 |
| *TPGS2--ACTN1* | 7.7 |
| *TPD52L2--VCP* | 7.7 |
| *TPCN1--EDN2* | 7.7 |
| *TP53BP1--DOCK7* | 7.7 |
| *TOR1AIP1--TTBK2* | 7.7 |
| *TOP2B--CELSR3* | 7.7 |
| *TOP2A--WWOX* | 7.7 |
| *TOP1--P2RY11* | 7.7 |
| *TOP1--MYH15* | 7.7 |
| *TOLLIP--AC021087.5* | 7.7 |
| *TOB1--OSGIN1* | 7.7 |
| *TNS3--AKR1C4* | 7.7 |
| *TNS1--WDR70* | 7.7 |
| *TNS1--PDE5A* | 7.7 |
| *TNRC6C--PRSS35* | 7.7 |
| *TNRC6C--MUC4* | 7.7 |
| *TNRC6B--SCP2* | 7.7 |
| *TNRC6B--MALAT1* | 7.7 |
| *TNRC6A--FOXA2* | 7.7 |
| *TNRC6A--ACTR2* | 7.7 |
| *TNRC18--FGD5* | 7.7 |
| *TNPO2--MALAT1* | 7.7 |
| *TNPO1--STAT1* | 7.7 |
| *TNKS2--SCARNA5* | 7.7 |
| *TNKS2--HERC2* | 7.7 |
| *TNK2--MAGI2* | 7.7 |
| *TNIP1--ZMYM4* | 7.7 |
| *TNFRSF21--HIST1H1B* | 7.7 |

| | |
|---|---|
| *TNFRSF14--SDAD1* | 7.7 |
| *TNC--NUMA1* | 7.7 |
| *TMTC3--ADCY2* | 7.7 |
| *TMPO--COL1A1* | 7.7 |
| *TMEM59--GANC* | 7.7 |
| *TMEM43--PPP1R15B* | 7.7 |
| *TMEM38B--YME1L1* | 7.7 |
| *TMEM38B--ACTG1* | 7.7 |
| *TMEM259--ARF4* | 7.7 |
| *TMEM229B--RPGRIP1L* | 7.7 |
| *TMEM209--ZNF699* | 7.7 |
| *TMEM184C--MALAT1* | 7.7 |
| *TMEM18--TNKS1BP1* | 7.7 |
| *TMEM176B--COMTD1* | 7.7 |
| *TMEM165--HSPB1* | 7.7 |
| *TMEM161B-AS1--VPS13A* | 7.7 |
| *TMEM131--AC021087.5* | 7.7 |
| *TMEM117--PCLO* | 7.7 |
| *TMEM108--RAB10* | 7.7 |
| *TMEM107--FBXO31* | 7.7 |
| *TMEM107--ARHGAP28* | 7.7 |
| *TMED10--H19* | 7.7 |
| *TMCO6--CTNNAL1* | 7.7 |
| *TMCO4--KAT6A* | 7.7 |
| *TMCO1--IVNS1ABP* | 7.7 |
| *TM9SF3--MALAT1* | 7.7 |
| *TM9SF2--LRIG2* | 7.7 |
| *TM9SF2--CCDC80* | 7.7 |
| *TLN1--CACNA1A* | 7.7 |
| *TLE5--POLR2A* | 7.7 |
| *TLE5--MALAT1* | 7.7 |
| *TLE3--CTSK* | 7.7 |
| *TLCD4-RWDD3--STAB1* | 7.7 |
| *TLCD4--STAB1* | 7.7 |
| *TJAP1--LMF1* | 7.7 |
| *TIRAP--KLF6* | 7.7 |
| *TIMM23B--BMS1P4-AGAP5* | 7.7 |
| *TIMM23B--AC022400.4* | 7.7 |
| *TICRR--PFDN5* | 7.7 |
| *TICRR--PABPC1* | 7.7 |
| *THUMPD3--HTATSF1* | 7.7 |
| *THRAP3--MALAT1* | 7.7 |
| *THOC7--SAT1* | 7.7 |
| *THOC5--MUC16* | 7.7 |

| | |
|---|---|
| *THOC1--SGK1* | 7.7 |
| *THBS2--AL445685.3* | 7.7 |
| *THAP6--AC006978.1* | 7.7 |
| *THAP5--WDR45B* | 7.7 |
| *TGIF2-RAB5IF--ZNF672* | 7.7 |
| *TGIF1--MAK16* | 7.7 |
| *TGIF1--FTL* | 7.7 |
| *TGFBR1--PGK1* | 7.7 |
| *TFG--PHACTR4* | 7.7 |
| *TF--MALAT1* | 7.7 |
| *TF--ADGRV1* | 7.7 |
| *TEX261--EIF3A* | 7.7 |
| *TEX2--CCDC6* | 7.7 |
| *TET2--ZNF117* | 7.7 |
| *TET2--TRAPPC10* | 7.7 |
| *TES--TRPC4AP* | 7.7 |
| *TENT5A--EIF2AK1* | 7.7 |
| *TENT4B--FTH1* | 7.7 |
| *TENT4A--OSGEP* | 7.7 |
| *TEAD2--PABPC1* | 7.7 |
| *TDRD3--CSPP1* | 7.7 |
| *TDRD3--ACTG1* | 7.7 |
| *TCTN1--WDR70* | 7.7 |
| *TCP1--LBH* | 7.7 |
| *TCF7--C3ORF62* | 7.7 |
| *TCF4--WBP4* | 7.7 |
| *TCF4--SMG1* | 7.7 |
| *TCF4--ITSN2* | 7.7 |
| *TCF25--PRPF8* | 7.7 |
| *TCEAL4--HDAC7* | 7.7 |
| *TBX2--RB1* | 7.7 |
| *TBX15--EPRS* | 7.7 |
| *TBL1X--PRRC2C* | 7.7 |
| *TBCE--ATP5MC2* | 7.7 |
| *TBCA--DDX5* | 7.7 |
| *TBC1D9B--SQSTM1* | 7.7 |
| *TBC1D9B--MALAT1* | 7.7 |
| *TBC1D9B--ETV6* | 7.7 |
| *TBC1D4--MALAT1* | 7.7 |
| *TBC1D32--LUCAT1* | 7.7 |
| *TASP1--NCOR2* | 7.7 |
| *TASOR2--WRNIP1* | 7.7 |
| *TAS2R63P--TAS2R30* | 7.7 |
| *TARS--TCF3* | 7.7 |
| *TARDBP--KIAA1109* | 7.7 |

| | |
|---|---|
| *TAOK1--RHEB* | 7.7 |
| *TAOK1--COL1A1* | 7.7 |
| *TANGO6--MTCH2* | 7.7 |
| *TANC2--NHLRC3* | 7.7 |
| *TANC2--HSPG2* | 7.7 |
| *TAF1D--AL033519.5* | 7.7 |
| *TAF15--VPS13A* | 7.7 |
| *TAF15--SF1* | 7.7 |
| *TAF11--WNK1* | 7.7 |
| *TAF1--UBAP1* | 7.7 |
| *TAB3--MKNK2* | 7.7 |
| *SYTL4--MBIP* | 7.7 |
| *SYT14--INTS10* | 7.7 |
| *SYNRG--CHCHD10* | 7.7 |
| *SYNPO--EIF4A2* | 7.7 |
| *SYNJ2--AL358113.1* | 7.7 |
| *SYNE3--DNAH14* | 7.7 |
| *SYNE2--MALAT1* | 7.7 |
| *SYNE2--LGALS3BP* | 7.7 |
| *SYNE2--HIST1H2AG* | 7.7 |
| *SYNE2--FAM168A* | 7.7 |
| *SYNE1--FTL* | 7.7 |
| *SYNE1--ATP5F1A* | 7.7 |
| *SYCP2--PTPRB* | 7.7 |
| *SWAP70--CDK13* | 7.7 |
| *SUZ12--TRB@* | 7.7 |
| *SUZ12--NF1* | 7.7 |
| *SUZ12--ASCL1* | 7.7 |
| *SUPT6H--RBL2* | 7.7 |
| *SUMO3--NUP153* | 7.7 |
| *SUMO2--IL6ST* | 7.7 |
| *SULF1--ADCY4* | 7.7 |
| *SUFU--ADGRD1* | 7.7 |
| *STXBP5--SHPRH* | 7.7 |
| *STXBP5--LYST* | 7.7 |
| *STX7--XIST* | 7.7 |
| *STX3--SETBP1* | 7.7 |
| *STRN4--SON* | 7.7 |
| *STRADA--ATP2B1* | 7.7 |
| *STN1--ZNF292* | 7.7 |
| *STMN1--USP7* | 7.7 |
| *STMN1--ARHGAP22* | 7.7 |
| *STK4--WWOX* | 7.7 |
| *STK4--CTDSP1* | 7.7 |
| *STK38--RBBP8* | 7.7 |

| | |
|---|---|
| *STK36--ZMYM2* | 7.7 |
| *STK32B--UBC* | 7.7 |
| *STK24--EPRS* | 7.7 |
| *STK24--C9ORF85* | 7.7 |
| *STIL--BDP1* | 7.7 |
| *STAT5B--NPEPPS* | 7.7 |
| *STAT1--PEX6* | 7.7 |
| *STAT1--NEAT1* | 7.7 |
| *STAT1--ANP32B* | 7.7 |
| *STARD7--ZNF300* | 7.7 |
| *STARD4--CD46* | 7.7 |
| *STARD3--GEN1* | 7.7 |
| *STARD10--STAG1* | 7.7 |
| *STAMBP--LONP2* | 7.7 |
| *STAG3L5P-PVRIG2P-PILRB--JAM3* | 7.7 |
| *STAG3--GTF2IRD2B* | 7.7 |
| *STAG3--C19MC* | 7.7 |
| *ST8SIA4--MOV10* | 7.7 |
| *ST5--SBF2* | 7.7 |
| *ST3GAL6--SLC16A1-AS1* | 7.7 |
| *ST18--USP34* | 7.7 |
| *ST13--XPNPEP1* | 7.7 |
| *SSR1--AKR1C4* | 7.7 |
| *SSBP3--UBC* | 7.7 |
| *SSBP3--SYBU* | 7.7 |
| *SSBP1--PRRC1* | 7.7 |
| *SSBP1--MGAM* | 7.7 |
| *SRSF9--PDCD4* | 7.7 |
| *SRSF4--MESD* | 7.7 |
| *SRSF4--MALAT1* | 7.7 |
| *SRSF3--RARS2* | 7.7 |
| *SRSF2--CLCN7* | 7.7 |
| *SRSF11--CTCF* | 7.7 |
| *SRSF10--ZNF528* | 7.7 |
| *SRRT--RMRP* | 7.7 |
| *SRRM2--TRMT11* | 7.7 |
| *SRRM2--SLC29A2* | 7.7 |
| *SRRM2--HUWE1* | 7.7 |
| *SRRM2--HGSNAT* | 7.7 |
| *SRRM2--GTF2I* | 7.7 |
| *SRRM2--GAPDH* | 7.7 |
| *SRRM2--COL18A1* | 7.7 |
| *SRRM2--ATM* | 7.7 |
| *SRPRA--FANCI* | 7.7 |

| | |
|---|---|
| *SRP72--YTHDF2* | 7.7 |
| *SRGAP1--SH3BP2* | 7.7 |
| *SRC--UNC119* | 7.7 |
| *SQSTM1--PCNT* | 7.7 |
| *SPTBN1--SLC26A8* | 7.7 |
| *SPTBN1--NUP214* | 7.7 |
| *SPTBN1--HELZ2* | 7.7 |
| *SPTBN1--FKTN* | 7.7 |
| *SPTAN1--STXBP5* | 7.7 |
| *SPTAN1--RAB11FIP1* | 7.7 |
| *SPTAN1--CARM1* | 7.7 |
| *SPPL3--XBP1* | 7.7 |
| *SPPL2A--NBPF9* | 7.7 |
| *SPIN4--ALCAM* | 7.7 |
| *SPIN3--GTF2I* | 7.7 |
| *SPICE1--TTC17* | 7.7 |
| *SPG7--EIF4B* | 7.7 |
| *SPG7--COL6A2* | 7.7 |
| *SPEF2--SRP72* | 7.7 |
| *SPECC1L-ADORA2A--PPWD1* | 7.7 |
| *SPECC1L-ADORA2A--ADA2* | 7.7 |
| *SPECC1L--PPWD1* | 7.7 |
| *SPC24--IL20RB* | 7.7 |
| *SPATS2L--HYMAI* | 7.7 |
| *SPATS2--PDGFRL* | 7.7 |
| *SPART--PTPRC* | 7.7 |
| *SPART--GRHL2* | 7.7 |
| *SPARC--SPCS1* | 7.7 |
| *SPARC--NACA4P* | 7.7 |
| *SPARC--EHBP1* | 7.7 |
| *SPAG9--RPL10* | 7.7 |
| *SP3--OLA1* | 7.7 |
| *SP140L--FBRSL1* | 7.7 |
| *SP100--ZNF91* | 7.7 |
| *SP1--XRCC5* | 7.7 |
| *SP1--KARS* | 7.7 |
| *SOX4--RFX6* | 7.7 |
| *SOX4--B3GALNT2* | 7.7 |
| *SOX18--WDR74* | 7.7 |
| *SOS2--WDR36* | 7.7 |
| *SOS1--TGFBR1* | 7.7 |
| *SORL1--MALAT1* | 7.7 |
| *SORCS1--A2M* | 7.7 |
| *SON--SSR1* | 7.7 |

| | |
|---|---|
| *SON--PFKL* | 7.7 |
| *SON--EEF1B2* | 7.7 |
| *SOD3--CACNA1H* | 7.7 |
| *SNX9--UQCRB* | 7.7 |
| *SNX6--NCL* | 7.7 |
| *SNX30--HADHB* | 7.7 |
| *SNX27--MALAT1* | 7.7 |
| *SNX25--TPR* | 7.7 |
| *SNX21--ITFG1* | 7.7 |
| *SNX13--DELE1* | 7.7 |
| *SNU13--C12ORF4* | 7.7 |
| *SNU13--AC091060.1* | 7.7 |
| *SNTB1--AC253536.7* | 7.7 |
| *SNRPN--SLC30A9* | 7.7 |
| *SNRNP27--CHSY1* | 7.7 |
| *SNRNP200--MALAT1* | 7.7 |
| *SMURF2--NCAPG2* | 7.7 |
| *SMTN--TANK* | 7.7 |
| *SMPDL3A--PTS* | 7.7 |
| *SMIM7--MALAT1* | 7.7 |
| *SMG1--SNX1* | 7.7 |
| *SMG1--SLC39A10* | 7.7 |
| *SMG1--RSL24D1* | 7.7 |
| *SMG1--PIKFYVE* | 7.7 |
| *SMG1--GTF2I* | 7.7 |
| *SMG1--GSN* | 7.7 |
| *SMG1--DSP* | 7.7 |
| *SMG1--CCL3* | 7.7 |
| *SMG1--CACNA2D1* | 7.7 |
| *SMG1--C2CD5* | 7.7 |
| *SMCHD1--PARG* | 7.7 |
| *SMC6--ZBTB22* | 7.7 |
| *SMC4--SNU13* | 7.7 |
| *SMC4--MUC4* | 7.7 |
| *SMC4--KRCC1* | 7.7 |
| *SMC3--PDCD4* | 7.7 |
| *SMC2--DNAH14* | 7.7 |
| *SMC1A--MDN1* | 7.7 |
| *SMARCC2--RPPH1* | 7.7 |
| *SMARCC2--CLCN2* | 7.7 |
| *SMARCC2--AKR1C3* | 7.7 |
| *SMARCC1--UBC* | 7.7 |
| *SMARCC1--BRWD1* | 7.7 |
| *SMARCA5--MALAT1* | 7.7 |
| *SMARCA4--ZFP36L1* | 7.7 |

| | |
|---|---|
| *SMARCA4--RELL1* | 7.7 |
| *SMARCA4--AL031777.3* | 7.7 |
| *SMARCA2--HNRNPA3* | 7.7 |
| *SMARCA1--MALAT1* | 7.7 |
| *SMARCA1--EEF1B2* | 7.7 |
| *SMAD4--SH3GLB2* | 7.7 |
| *SMAD3--AC007114.2* | 7.7 |
| *SMAD1--SPTBN1* | 7.7 |
| *SLTM--RNF141* | 7.7 |
| *SLTM--CNOT8* | 7.7 |
| *SLTM--AC107029.2* | 7.7 |
| *SLMAP--TSC2* | 7.7 |
| *SLC9A7--HYOU1* | 7.7 |
| *SLC9A3R1--SPEN* | 7.7 |
| *SLC8A1--RBM25* | 7.7 |
| *SLC7A6--CNTN3* | 7.7 |
| *SLC7A11--METTL7B* | 7.7 |
| *SLC6A9--SREBF1* | 7.7 |
| *SLC6A15--RASSF3* | 7.7 |
| *SLC4A7--TBK1* | 7.7 |
| *SLC4A7--ANKRD17* | 7.7 |
| *SLC40A1--MLYCD* | 7.7 |
| *SLC40A1--FUBP1* | 7.7 |
| *SLC39A5--FAM234A* | 7.7 |
| *SLC39A10--AC092943.1* | 7.7 |
| *SLC38A9--NUP153* | 7.7 |
| *SLC35F5--MALAT1* | 7.7 |
| *SLC25A46--RMRP* | 7.7 |
| *SLC25A40--GTF3C1* | 7.7 |
| *SLC25A4--VAT1* | 7.7 |
| *SLC25A3--TM9SF2* | 7.7 |
| *SLC25A3--TAPT1* | 7.7 |
| *SLC25A3--ASAP1* | 7.7 |
| *SLC25A16--ATP13A1* | 7.7 |
| *SLC25A13--TENM4* | 7.7 |
| *SLC24A3--LUC7L2* | 7.7 |
| *SLC24A1--MIR4435-2HG* | 7.7 |
| *SLC1A5--ZC3H13* | 7.7 |
| *SLC1A3--EHF* | 7.7 |
| *SLC19A2--PRKDC* | 7.7 |
| *SLC16A1--IGKC* | 7.7 |
| *SLC16A1--IGK@* | 7.7 |
| *SLC16A1--ADGRL1* | 7.7 |
| *SLC12A6--ZEB2* | 7.7 |

| | |
|---|---|
| *SLC12A2--IGSF10* | 7.7 |
| *SLC12A2--GOLGA2* | 7.7 |
| *SIPA1L3--LRRN4* | 7.7 |
| *SIPA1L2--TUG1* | 7.7 |
| *SIPA1L1--KLB* | 7.7 |
| *SIK3--MALAT1* | 7.7 |
| *SIDT2--CFL2* | 7.7 |
| *SIAE--DHX9* | 7.7 |
| *SHTN1--RMND5A* | 7.7 |
| *SHROOM4--PPP1R3F* | 7.7 |
| *SHROOM4--NFAT5* | 7.7 |
| *SHROOM3--MALAT1* | 7.7 |
| *SHROOM3--ACTB* | 7.7 |
| *SHPRH--KCNB2* | 7.7 |
| *SHLD2--RANGAP1* | 7.7 |
| *SHLD2--FN1* | 7.7 |
| *SHISA6--PPFIBP2* | 7.7 |
| *SHISA5--MALAT1* | 7.7 |
| *SHISA4--IGF1R* | 7.7 |
| *SHISA2--SPPL2B* | 7.7 |
| *SHC1--GRIK5* | 7.7 |
| *SH3PXD2A--EDIL3* | 7.7 |
| *SH3GLB1--VCAN* | 7.7 |
| *SH3GLB1--E2F1* | 7.7 |
| *SH2D3C--KTN1* | 7.7 |
| *SGSM2--FBXO41* | 7.7 |
| *SFSWAP--RYBP* | 7.7 |
| *SFSWAP--FSIP2* | 7.7 |
| *SFPQ--TPM1* | 7.7 |
| *SFPQ--RBIS* | 7.7 |
| *SFPQ--PRPF6* | 7.7 |
| *SF3B2--CEP350* | 7.7 |
| *SF3B2--ACIN1* | 7.7 |
| *SF3B1--ZNF362* | 7.7 |
| *SF3B1--CELSR1* | 7.7 |
| *SETX--RAC1* | 7.7 |
| *SETDB1--MALAT1* | 7.7 |
| *SETD5--CSGALNACT2* | 7.7 |
| *SETBP1--WDR74* | 7.7 |
| *SET--PRDM2* | 7.7 |
| *SET--MALAT1* | 7.7 |
| *SET--IPO5* | 7.7 |
| *SET--AC019155.1* | 7.7 |
| *SESTD1--CIC* | 7.7 |
| *SESN3--SIK2* | 7.7 |

| | |
|---|---|
| *SERPINE2--ATIC* | 7.7 |
| *SERPINA1--MAN1A2* | 7.7 |
| *SERPINA1--EEF2* | 7.7 |
| *SERP1--TMSB4X* | 7.7 |
| *SERP1--ERGIC3* | 7.7 |
| *SERBP1--NT5C* | 7.7 |
| *SEPTIN2--CNOT3* | 7.7 |
| *SEPTIN2--BCOR* | 7.7 |
| *SEPTIN11--KDM4B* | 7.7 |
| *SEPTIN11--FTH1* | 7.7 |
| *SEPHS2--UBQLN2* | 7.7 |
| *SENP6--B4GALT5* | 7.7 |
| *SEMA6A--MYO1E* | 7.7 |
| *SEMA6A--CALCA* | 7.7 |
| *SEMA3C--TNRC6B* | 7.7 |
| *SEMA3C--SMG6* | 7.7 |
| *SELENON--ATRX* | 7.7 |
| *SELENOF--EPS15L1* | 7.7 |
| *SEC63--WWOX* | 7.7 |
| *SEC63--RTN1* | 7.7 |
| *SEC63--BANK1* | 7.7 |
| *SEC61A1--CPT2* | 7.7 |
| *SEC31A--MGAT4A* | 7.7 |
| *SEC31A--COPB2* | 7.7 |
| *SEC24D--MASTL* | 7.7 |
| *SEC23B--PAPOLA* | 7.7 |
| *SEC23B--CPLX2* | 7.7 |
| *SEC23A--ACTA1* | 7.7 |
| *SEC14L1--CASP3* | 7.7 |
| *SEC13--SNX33* | 7.7 |
| *SDHC--IGFN1* | 7.7 |
| *SDF4--RIMS2* | 7.7 |
| *SDF4--REV1* | 7.7 |
| *SCYL1--GTF3C5* | 7.7 |
| *SCN9A--MALAT1* | 7.7 |
| *SCN8A--ILRUN* | 7.7 |
| *SCG3--NAA15* | 7.7 |
| *SCG2--RMRP* | 7.7 |
| *SCG2--NEAT1* | 7.7 |
| *SCFD1--NOP53* | 7.7 |
| *SCD5--AC027097.2* | 7.7 |
| *SCD--FGFR1* | 7.7 |
| *SCD--APOC3* | 7.7 |
| *SCARNA9--ZNF676* | 7.7 |
| *SCARNA7--NPAS2* | 7.7 |

| | |
|---|---|
| *SCARNA7--EXOSC10* | 7.7 |
| *SCARNA5--SLC22A17* | 7.7 |
| *SCARNA5--NEAT1* | 7.7 |
| *SCARNA5--AMD1* | 7.7 |
| *SCARNA2--LIMA1* | 7.7 |
| *SCAMP1--CCDC162P* | 7.7 |
| *SCAI--SRRM2* | 7.7 |
| *SCAF4--RALGAPB* | 7.7 |
| *SCAF11--AC117386.2* | 7.7 |
| *SBNO1--NUCB1* | 7.7 |
| *SAT1--RACGAP1* | 7.7 |
| *SAT1--MED8* | 7.7 |
| *SAT1--CREB1* | 7.7 |
| *SASS6--LAMB1* | 7.7 |
| *SART3--SMOC2* | 7.7 |
| *SAP130--B2M* | 7.7 |
| *SAMM50--FAM104B* | 7.7 |
| *SAFB2--PTK2* | 7.7 |
| *SACS--NEAT1* | 7.7 |
| *SACS--MLLT6* | 7.7 |
| *S100A6--STARD13* | 7.7 |
| *S100A11--CAP1* | 7.7 |
| *RUNX1--FAM111B* | 7.7 |
| *RUFY3--NAGA* | 7.7 |
| *RUBCN--FYTTD1* | 7.7 |
| *RTTN--CCN2* | 7.7 |
| *RTN3--H19* | 7.7 |
| *RTL6--C16ORF96* | 7.7 |
| *RTKN--ZNF347* | 7.7 |
| *RTF1--SLC3A2* | 7.7 |
| *RTEL1-TNFRSF6B--TTC3* | 7.7 |
| *RSRC2--MALAT1* | 7.7 |
| *RSL24D1--GJA1* | 7.7 |
| *RSF1--TMED10* | 7.7 |
| *RSF1--KIAA1109* | 7.7 |
| *RSAD2--NEB* | 7.7 |
| *RRP8--MALAT1* | 7.7 |
| *RRP36--MIR205HG* | 7.7 |
| *RRP15--CRKL* | 7.7 |
| *RRBP1--SMARCA1* | 7.7 |
| *RRBP1--MYDGF* | 7.7 |
| *RRBP1--LCOR* | 7.7 |
| *RRAS--LMAN1* | 7.7 |
| *RPS6KB1--MALAT1* | 7.7 |

| | |
|---|---|
| *RPS6KA3--HSP90AA1* | 7.7 |
| *RPS6--PAX5* | 7.7 |
| *RPS4X--RGS3* | 7.7 |
| *RPS3--UQCRC2* | 7.7 |
| *RPS29--MALAT1* | 7.7 |
| *RPS27A--PITPNM3* | 7.7 |
| *RPS24--RMRP* | 7.7 |
| *RPS20--FTSJ3* | 7.7 |
| *RPS18--PTPN21* | 7.7 |
| *RPS17--TBL1XR1* | 7.7 |
| *RPS17--MALAT1* | 7.7 |
| *RPS15--PRPF31* | 7.7 |
| *RPPH1--ZBTB16* | 7.7 |
| *RPPH1--TRAPPC1* | 7.7 |
| *RPPH1--SEPTIN9* | 7.7 |
| *RPPH1--PRKAR1A* | 7.7 |
| *RPPH1--OGDH* | 7.7 |
| *RPPH1--MYH10* | 7.7 |
| *RPPH1--MUC4* | 7.7 |
| *RPPH1--LPXN* | 7.7 |
| *RPPH1--FTL* | 7.7 |
| *RPPH1--DST* | 7.7 |
| *RPPH1--CNOT1* | 7.7 |
| *RPPH1--CEP350* | 7.7 |
| *RPPH1--ARMC8* | 7.7 |
| *RPPH1--AP001267.5* | 7.7 |
| *RPPH1--AC138409.2* | 7.7 |
| *RPN2--HAUS6* | 7.7 |
| *RPN1--MKI67* | 7.7 |
| *RPLP0--MALAT1* | 7.7 |
| *RPL8--WDR74* | 7.7 |
| *RPL8--SOGA1* | 7.7 |
| *RPL5--FMNL2* | 7.7 |
| *RPL38--RMRP* | 7.7 |
| *RPL37A--MALAT1* | 7.7 |
| *RPL37--CRABP2* | 7.7 |
| *RPL35A--YRDC* | 7.7 |
| *RPL34--FAM91A1* | 7.7 |
| *RPL32--MKI67* | 7.7 |
| *RPL32--CD24* | 7.7 |
| *RPL31--IGK@* | 7.7 |
| *RPL31--FTL* | 7.7 |
| *RPL27A--MALAT1* | 7.7 |
| *RPL24--TTC37* | 7.7 |
| *RPL23AP82--DHX9* | 7.7 |

| | | | | | | | |
|---|---|---|---|---|---|---|---|
| *RPL23--APEH* | 7.7 | | *RMRP--PDS5A* | 7.7 | | *RBM47--NAMPT* | 7.7 |
| *RPL23--AC013549.4* | 7.7 | | *RMRP--NUP153* | 7.7 | | *RBM47--HUWE1* | 7.7 |
| *RPL17--DOCK6* | 7.7 | | *RMRP--MALAT1* | 7.7 | | *RBM39--MALAT1* | 7.7 |
| *RPL13A--UBN2* | 7.7 | | *RMRP--HSF1* | 7.7 | | *RBM39--HIST1H2BG* | 7.7 |
| *RPL13A--ANKRD17* | 7.7 | | *RMRP--HEATR1* | 7.7 | | *RBM39--G6PD* | 7.7 |
| *RPL13--ELN* | 7.7 | | *RMRP--GPCPD1* | 7.7 | | *RBM38--BBS2* | 7.7 |
| *RPL11--EFCAB14* | 7.7 | | *RMRP--GNAS* | 7.7 | | *RBM24--FTH1* | 7.7 |
| *RPL11--C21ORF58* | 7.7 | | *RMRP--FTH1* | 7.7 | | *RBM15--TXNIP* | 7.7 |
| *RPL11--BRD4* | 7.7 | | *RMRP--C2CD5* | 7.7 | | *RBM14-RBM4--KIF1C* | 7.7 |
| *RPL11--ATXN7* | 7.7 | | *RMRP--ATP2B1* | 7.7 | | *RBM12--AP2A1* | 7.7 |
| *RPL11--ARFIP2* | 7.7 | | *RIOK3--SVIL* | 7.7 | | *RBL1--ANKRD44* | 7.7 |
| *RPL10--UBC* | 7.7 | | *RIMS2--VPS13B* | 7.7 | | *RBFOX2--PAXBP1* | 7.7 |
| *RPGRIP1L--SLC12A7* | 7.7 | | *RIMKLB--LAMC1* | 7.7 | | *RBFOX2--KCNJ2* | 7.7 |
| *RPGR--TLK1* | 7.7 | | *RICTOR--AP3B1* | 7.7 | | *RBFA--MLLT10* | 7.7 |
| *RPGR--COX4I1* | 7.7 | | *RIC1--GNAS* | 7.7 | | *RBBP6--AL645922.1* | 7.7 |
| *RPAP1--CUL3* | 7.7 | | *RHOU--EBF1* | 7.7 | | *RBBP4--TCF4* | 7.7 |
| *ROCK1--SET* | 7.7 | | *RHOQ--UROS* | 7.7 | | *RBBP4--GPI* | 7.7 |
| *ROBO2--COL1A2* | 7.7 | | *RHOA--TPI1* | 7.7 | | *RBBP4--FTH1* | 7.7 |
| *RO60--HNRNPC* | 7.7 | | *RHOA--KNDC1* | 7.7 | | *RB1--EP300* | 7.7 |
| *RNPS1--TTN* | 7.7 | | *RFX7--TUBGCP5* | 7.7 | | *RASSF6--KAT2B* | 7.7 |
| *RNPC3--PTPRA* | 7.7 | | *RFC1--SEC24A* | 7.7 | | *RARS2--OAZ1* | 7.7 |
| *RNF34--ZEB2* | 7.7 | | *REV1--SON* | 7.7 | | *RAPH1--NSD3* | 7.7 |
| *RNF213--RAD23B* | 7.7 | | *REV1--SH3BGRL* | 7.7 | | *RAP1GAP--ATP5F1B* | 7.7 |
| *RNF213--PPP5C* | 7.7 | | *RET--CDC42BPB* | 7.7 | | *RANGAP1--CAPN12* | 7.7 |
| *RNF213--HNRNPA3* | 7.7 | | *REST--MALAT1* | 7.7 | | *RANBP2--MUC1* | 7.7 |
| *RNF213--ATP2C1* | 7.7 | | *RERE--RALGAPB* | 7.7 | | *RANBP2--LIPF* | 7.7 |
| *RNF213--ALKBH5* | 7.7 | | *RERE--PRRC2A* | 7.7 | | *RANBP2--FOXK1* | 7.7 |
| *RNF213--AD000090.1* | 7.7 | | *RERE--DDX17* | 7.7 | | *RAN--MALAT1* | 7.7 |
| *RNF170--EPAS1* | 7.7 | | *REPS1--IGFBP7* | 7.7 | | *RAN--BTAF1* | 7.7 |
| *RNF152--UBA6* | 7.7 | | *REPS1--CENPT* | 7.7 | | *RALY--EP300* | 7.7 |
| *RNF149--UBE4B* | 7.7 | | *RELN--SETD5* | 7.7 | | *RALY--CFL1* | 7.7 |
| *RNF133--EFTUD2* | 7.7 | | *RECK--PREPL* | 7.7 | | *RALGAPA2--PSIP1* | 7.7 |
| *RNF13--GNB1* | 7.7 | | *REC8--XIST* | 7.7 | | *RALGAPA2--LPCAT3* | 7.7 |
| *RNF123--UBA52* | 7.7 | | *RC3H2--RPL8* | 7.7 | | *RALGAPA1--VCL* | 7.7 |
| *RNF115--PIAS3* | 7.7 | | *RC3H2--HOXC6* | 7.7 | | *RALGAPA1--MAPK7* | 7.7 |
| *RNF10--HNRNPA1* | 7.7 | | *RC3H2--CPSF6* | 7.7 | | *RALBP1--THRAP3* | 7.7 |
| *RNASEH2A--MUC5B* | 7.7 | | *RC3H1--TMF1* | 7.7 | | *RALB--SPATS2L* | 7.7 |
| *RMRP--ZFAND3* | 7.7 | | *RC3H1--PPT1* | 7.7 | | *RALA--NAA20* | 7.7 |
| *RMRP--XIST* | 7.7 | | *RC3H1--MALAT1* | 7.7 | | *RAD54L2--FTL* | 7.7 |
| *RMRP--USP24* | 7.7 | | *RBPMS--ATXN2L* | 7.7 | | *RAD54L2--DPY19L2* | 7.7 |
| *RMRP--UBALD2* | 7.7 | | *RBPJ--C8ORF33* | 7.7 | | *RAD51D--GTF2I* | 7.7 |
| *RMRP--TPX2* | 7.7 | | *RBMXL1--RPS17* | 7.7 | | *RAD51AP1--VHL* | 7.7 |
| *RMRP--SUGP2* | 7.7 | | *RBMX2--ENAH* | 7.7 | | *RAD23B--LAMA2* | 7.7 |
| *RMRP--SMARCAL1* | 7.7 | | *RBM5--TCERG1* | 7.7 | | *RAD21--ELOVL5* | 7.7 |
| *RMRP--SBNO1* | 7.7 | | *RBM5--ABI1* | 7.7 | | *RAD21--ADAR* | 7.7 |

| | |
|---|---|
| RAD18--COL1A2 | 7.7 |
| RAD17--DMTF1 | 7.7 |
| RAD17--ANO6 | 7.7 |
| RACK1--STARD9 | 7.7 |
| RACK1--RMRP | 7.7 |
| RACK1--AC108704.2 | 7.7 |
| RAC1--HSPA9 | 7.7 |
| RABEP1--WDR74 | 7.7 |
| RAB8B--NT5C2 | 7.7 |
| RAB8A--AXIN2 | 7.7 |
| RAB8A--AC004805.1 | 7.7 |
| RAB7A--UBC | 7.7 |
| RAB7A--ACAD9 | 7.7 |
| RAB6B--BACH1 | 7.7 |
| RAB3GAP2--PLIN3 | 7.7 |
| RAB3GAP2--CES1 | 7.7 |
| RAB3GAP1--PPP6R2 | 7.7 |
| RAB2A--VIPAS39 | 7.7 |
| RAB2A--FBRSL1 | 7.7 |
| RAB21--PHF20L1 | 7.7 |
| RAB18--SYNE2 | 7.7 |
| RAB12--NOP58 | 7.7 |
| RAB11A--CHD7 | 7.7 |
| QSOX2--PKP4 | 7.7 |
| QRICH2--CNOT1 | 7.7 |
| QKI--SF3B1 | 7.7 |
| PWWP2A--EIF4G2 | 7.7 |
| PVT1--MYC | 7.7 |
| PUM3--ATP11C | 7.7 |
| PUM1--SRRM2 | 7.7 |
| PTPRS--RAD21 | 7.7 |
| PTPRN2--CUX1 | 7.7 |
| PTPRG--MALAT1 | 7.7 |
| PTPRF--PTCH2 | 7.7 |
| PTPRF--LSP1 | 7.7 |
| PTPRF--FLNC | 7.7 |
| PTPRC--BTBD2 | 7.7 |
| PTPRA--MALAT1 | 7.7 |
| PTPN3--FN1 | 7.7 |
| PTPN14--SYNM | 7.7 |
| PTPN12--ZNF236 | 7.7 |
| PTPN11--CREBBP | 7.7 |
| PTP4A2--SRSF10 | 7.7 |
| PTOV1--AC063952.2 | 7.7 |
| PTMS--PSMC4 | 7.7 |

| | |
|---|---|
| PTMS--P4HA2 | 7.7 |
| PTMS--GTF2A1L | 7.7 |
| PTMA--NEFM | 7.7 |
| PTMA--NEAT1 | 7.7 |
| PTMA--MALAT1 | 7.7 |
| PTMA--KIF21A | 7.7 |
| PTMA--CDC25A | 7.7 |
| PTK7--PRCP | 7.7 |
| PTGES3--FTH1 | 7.7 |
| PTCH2--FTL | 7.7 |
| PTCH1--ECPAS | 7.7 |
| PTCH1--DMGDH | 7.7 |
| PTBP3--GAL | 7.7 |
| PTBP2--IGHMBP2 | 7.7 |
| PTBP1--SH2B1 | 7.7 |
| PTBP1--COPZ1 | 7.7 |
| PTAR1--EIF2B1 | 7.7 |
| PSMG4--STAG1 | 7.7 |
| PSME4--RPL14 | 7.7 |
| PSME4--MALAT1 | 7.7 |
| PSME3--RPL37A | 7.7 |
| PSME1--SELENOT | 7.7 |
| PSME1--MAPK1IP1L | 7.7 |
| PSMD2--PRR34-AS1 | 7.7 |
| PSMD2--GAP43 | 7.7 |
| PSMD13--ZNF664 | 7.7 |
| PSMD12--AHNAK | 7.7 |
| PSMC4--ATP13A3 | 7.7 |
| PSMB4--CTCF | 7.7 |
| PSMA3--STIL | 7.7 |
| PSMA3--FREM2 | 7.7 |
| PSIP1--SAR1A | 7.7 |
| PSIP1--IGF2 | 7.7 |
| PSIP1--CCDC171 | 7.7 |
| PSD3--TUBB | 7.7 |
| PSAT1--UBR4 | 7.7 |
| PSAP--NPLOC4 | 7.7 |
| PSAP--KRT7 | 7.7 |
| PSAP--FCHSD2 | 7.7 |
| PRRT1--PPM1G | 7.7 |
| PRRG3--RBM6 | 7.7 |
| PRRC2C--ZC3H7A | 7.7 |
| PRRC2C--TBC1D14 | 7.7 |
| PRRC2C--SMC5 | 7.7 |
| PRRC2C--RP1 | 7.7 |

| | |
|---|---|
| PRRC2C--NFAT5 | 7.7 |
| PRRC2C--MALAT1 | 7.7 |
| PRRC2C--ENTPD4 | 7.7 |
| PRRC2C--DNAJA1 | 7.7 |
| PRRC2C--CYP4B1 | 7.7 |
| PRRC2C--CHST11 | 7.7 |
| PRRC2B--NEFM | 7.7 |
| PRRC2B--MALAT1 | 7.7 |
| PRRC2A--STOM | 7.7 |
| PRRC2A--MALAT1 | 7.7 |
| PRR5-ARHGAP8--TIMP1 | 7.7 |
| PRPSAP1--HECTD1 | 7.7 |
| PRPF8--RNF38 | 7.7 |
| PRPF8--RIF1 | 7.7 |
| PRPF8--PPARG | 7.7 |
| PRPF8--MALAT1 | 7.7 |
| PRPF4B--SERPINA1 | 7.7 |
| PRPF38B--TRB@ | 7.7 |
| PRPF3--ZMYND8 | 7.7 |
| PROCR--RHOB | 7.7 |
| PROCA1--PDCD10 | 7.7 |
| PROC--NR2F2 | 7.7 |
| PRNP--ZBTB22 | 7.7 |
| PRNP--SYNPO2 | 7.7 |
| PRMT7--RPPH1 | 7.7 |
| PRMT5--PTCH2 | 7.7 |
| PRKDC--MALAT1 | 7.7 |
| PRKDC--ACTG1 | 7.7 |
| PRKCSH--RNF145 | 7.7 |
| PRKAB2--STARD7 | 7.7 |
| PRKAB2--HNRNPC | 7.7 |
| PRIM1--UGGT1 | 7.7 |
| PRICKLE2--BPNT1 | 7.7 |
| PREPL--IGK@ | 7.7 |
| PREP--FAM172A | 7.7 |
| PRELID2--MALAT1 | 7.7 |
| PRDM2--EIF5 | 7.7 |
| PQBP1--TALDO1 | 7.7 |
| PPWD1--ESD | 7.7 |
| PPRC1--PPP4R3A | 7.7 |
| PPP4R3B--SETD4 | 7.7 |
| PPP4R3A--CYP4F11 | 7.7 |
| PPP4R1--UQCRFS1 | 7.7 |
| PPP2R3B--MROH1 | 7.7 |

| | |
|---|---|
| *PPP2R1A--PKP4* | 7.7 |
| *PPP2R1A--CCDC82* | 7.7 |
| *PPP1R3B--SERPINH1* | 7.7 |
| *PPP1R12C--PLBD1* | 7.7 |
| *PPP1R12C--DDX39B* | 7.7 |
| *PPP1R12B--SS18L1* | 7.7 |
| *PPP1CC--SUPT16H* | 7.7 |
| *PPP1CA--RP1L1* | 7.7 |
| *PPM1L--NLRP3* | 7.7 |
| *PPM1L--EFHD2* | 7.7 |
| *PPM1G--HMGCS1* | 7.7 |
| *PPM1G--ABCC2* | 7.7 |
| *PPIP5K2--CDK16* | 7.7 |
| *PPIP5K1--LGR4* | 7.7 |
| *PPIL3--MTA2* | 7.7 |
| *PPIH--YBX1* | 7.7 |
| *PPIG--PDPK1* | 7.7 |
| *PPIG--B3GALNT1* | 7.7 |
| *PPIG--AL627309.5* | 7.7 |
| *PPIB--SLC35G1* | 7.7 |
| *PPIB--LAMB3* | 7.7 |
| *PPIA--RRBP1* | 7.7 |
| *PPFIBP1--WDFY3* | 7.7 |
| *PPARG--ACTG1* | 7.7 |
| *POU2F1--SPIN1* | 7.7 |
| *POSTN--CLIP4* | 7.7 |
| *POSTN--C1R* | 7.7 |
| *POMT2--RCC1* | 7.7 |
| *POM121C--RBAK* | 7.7 |
| *POLR3G--AARS* | 7.7 |
| *POLR3E--ERGIC1* | 7.7 |
| *POLR3A--P4HA2* | 7.7 |
| *POLR2E--WDR74* | 7.7 |
| *POLR2B--AP002373.1* | 7.7 |
| *POLR2A--KPNB1* | 7.7 |
| *POLR2A--KANK3* | 7.7 |
| *POLR1B--RAP1GAP2* | 7.7 |
| *POLR1A--COMMD9* | 7.7 |
| *POLG--BRWD1* | 7.7 |
| *POLD2--COL1A2* | 7.7 |
| *POLA1--NEB* | 7.7 |
| *POGZ--RNF111* | 7.7 |
| *POGK--EXO1* | 7.7 |
| *PNPT1--SCARNA9* | 7.7 |
| *PNISR--PTAR1* | 7.7 |

| | |
|---|---|
| *PLXND1--GFM1* | 7.7 |
| *PLXNB3--GTF2I* | 7.7 |
| *PLXNB2--RPPH1* | 7.7 |
| *PLXNB2--PTPRN2* | 7.7 |
| *PLXNB2--NUAK2* | 7.7 |
| *PLXNB1--PRPF8* | 7.7 |
| *PLXNA1--UBC* | 7.7 |
| *PLXNA1--NOP9* | 7.7 |
| *PLSCR1--SERPINH1* | 7.7 |
| *PLPP5--DDX5* | 7.7 |
| *PLP2--MUC4* | 7.7 |
| *PLOD1--ZNF91* | 7.7 |
| *PLOD1--PARP3* | 7.7 |
| *PLLP--MALAT1* | 7.7 |
| *PLEKHO1--COL11A1* | 7.7 |
| *PLEKHM3--FAS* | 7.7 |
| *PLEKHH1--VMP1* | 7.7 |
| *PLEKHG3--PHF2* | 7.7 |
| *PLEC--CHSY1* | 7.7 |
| *PLCL1--SSR1* | 7.7 |
| *PLCG1--KIF26A* | 7.7 |
| *PLA2G2A--ZNF766* | 7.7 |
| *PKP3--MALAT1* | 7.7 |
| *PKP3--EIF4G2* | 7.7 |
| *PKP1--BET1L* | 7.7 |
| *PKD1--ARHGAP26* | 7.7 |
| *PITRM1--CCT6A* | 7.7 |
| *PITRM1--AL358933.1* | 7.7 |
| *PITPNM2--PSMD14* | 7.7 |
| *PITPNA--ZNF281* | 7.7 |
| *PITPNA--EEF1A1* | 7.7 |
| *PIP5K1C--TUBB* | 7.7 |
| *PINK1-AS--TM9SF2* | 7.7 |
| *PIKFYVE--DOCK6* | 7.7 |
| *PIK3C3--WDR74* | 7.7 |
| *PIK3C2G--CDC42BPA* | 7.7 |
| *PIK3C2B--CALR* | 7.7 |
| *PIGW--MACC1* | 7.7 |
| *PIGS--APH1A* | 7.7 |
| *PIGB--TCF4* | 7.7 |
| *PIF1--PLK1* | 7.7 |
| *PIEZO1--MAZ* | 7.7 |
| *PICALM--SYTL2* | 7.7 |
| *PICALM--MYH9* | 7.7 |
| *PIBF1--SPIDR* | 7.7 |

| | |
|---|---|
| *PIBF1--KLF5* | 7.7 |
| *PIAS3--NUP98* | 7.7 |
| *PHTF2--PTPN12* | 7.7 |
| *PHTF1--NDUFV1* | 7.7 |
| *PHPT1--BIRC6* | 7.7 |
| *PHLDA1--DVL1* | 7.7 |
| *PHKB--PHF3* | 7.7 |
| *PHKB--KIAA1958* | 7.7 |
| *PHIP--TUBB* | 7.7 |
| *PHIP--SPTBN1* | 7.7 |
| *PHIP--PAWR* | 7.7 |
| *PHIP--COL24A1* | 7.7 |
| *PHGDH--ZNF469* | 7.7 |
| *PHF3--TUBA1B* | 7.7 |
| *PHF20--LARP4* | 7.7 |
| *PHF19--P2RX7* | 7.7 |
| *PHF14--VPS13D* | 7.7 |
| *PHF14--NCOR1* | 7.7 |
| *PHF13--MTA3* | 7.7 |
| *PHF12--DNAH5* | 7.7 |
| *PHC3--CAT* | 7.7 |
| *PHACTR1--MSN* | 7.7 |
| *PGRMC2--STMN2* | 7.7 |
| *PGM3--VPS13A* | 7.7 |
| *PGM2--SMG1* | 7.7 |
| *PGM2--BRWD1* | 7.7 |
| *PGK1--ZRANB3* | 7.7 |
| *PGD--FAM117B* | 7.7 |
| *PGBD1--ZNF276* | 7.7 |
| *PEX26--ACADVL* | 7.7 |
| *PEX2--TUBB* | 7.7 |
| *PEX19--SMG1* | 7.7 |
| *PERM1--YTHDF2* | 7.7 |
| *PELI2--PTCD3* | 7.7 |
| *PEG3--HNRNPU* | 7.7 |
| *PDXDC1--HTT* | 7.7 |
| *PDS5B--CHST11* | 7.7 |
| *PDS5A--SERPINI1* | 7.7 |
| *PDS5A--NIN* | 7.7 |
| *PDLIM2--BECN1* | 7.7 |
| *PDK2--SYNJ1* | 7.7 |
| *PDIA6--ARFGEF2* | 7.7 |
| *PDIA3--HR* | 7.7 |
| *PDHB--GPX2* | 7.7 |
| *PDE8A--SCAF8* | 7.7 |

| | |
|---|---|
| *PDE7A--TPM2* | 7.7 |
| *PDE5A--NLRC4* | 7.7 |
| *PDCD2L--AK2* | 7.7 |
| *PCSK5--NBR1* | 7.7 |
| *PCNX4--PHF8* | 7.7 |
| *PCNX4--MUC4* | 7.7 |
| *PCNX4--CNNM4* | 7.7 |
| *PCNP--SERPINF1* | 7.7 |
| *PCNA--NEAT1* | 7.7 |
| *PCMTD1--PTPRG* | 7.7 |
| *PCMT1--PKP4* | 7.7 |
| *PCM1--STT3A* | 7.7 |
| *PCM1--RHOXF2B* | 7.7 |
| *PCM1--EEF1A1* | 7.7 |
| *PCLO--ZCCHC14* | 7.7 |
| *PCLO--SMARCAD1* | 7.7 |
| *PCLO--CACNA2D1* | 7.7 |
| *PCLO--AC124312.3* | 7.7 |
| *PCID2--SETX* | 7.7 |
| *PCGF5--EGR1* | 7.7 |
| *PCED1A--PHLPP1* | 7.7 |
| *PCBP2--MKLN1* | 7.7 |
| *PCBP1--EZH1* | 7.7 |
| *PCBD1--MAST3* | 7.7 |
| *PC--MDM2* | 7.7 |
| *PBXIP1--PDCD11* | 7.7 |
| *PBX1--PTMS* | 7.7 |
| *PAXIP1--RSF1* | 7.7 |
| *PAXIP1--NSMAF* | 7.7 |
| *PAWR--SGCA* | 7.7 |
| *PATJ--OAS3* | 7.7 |
| *PATJ--MALAT1* | 7.7 |
| *PASK--EFHD2* | 7.7 |
| *PARVA--TEAD1* | 7.7 |
| *PARP9--CUL4B* | 7.7 |
| *PARP16--SDHAF1* | 7.7 |
| *PARP14--RMRP* | 7.7 |
| *PARP14--PPP2CB* | 7.7 |
| *PARP1--EDRF1* | 7.7 |
| *PARG--BMS1* | 7.7 |
| *PARD3--SGMS1* | 7.7 |
| *PAPOLA--SYNE2* | 7.7 |
| *PANK3--MUC4* | 7.7 |
| *PANK3--GKAP1* | 7.7 |
| *PAN2--KLK11* | 7.7 |

| | |
|---|---|
| *PAM--AKAP1* | 7.7 |
| *PALLD--TAGLN* | 7.7 |
| *PALLD--SLC35B4* | 7.7 |
| *PALLD--SH3BP5L* | 7.7 |
| *PAICS--MEGF8* | 7.7 |
| *PAFAH1B2--CD276* | 7.7 |
| *PAFAH1B1--NSUN5* | 7.7 |
| *PACRGL--CLTC* | 7.7 |
| *PABPC4--LDHA* | 7.7 |
| *PABPC4--CCT5* | 7.7 |
| *PABPC1--TMEM106C* | 7.7 |
| *PABPC1--SENP7* | 7.7 |
| *PABPC1--MALAT1* | 7.7 |
| *PABPC1--GANAB* | 7.7 |
| *PABPC1--GAK* | 7.7 |
| *PA2G4--S1PR3* | 7.7 |
| *PA2G4--MALAT1* | 7.7 |
| *OTUD7B--IER3IP1* | 7.7 |
| *OSTC--LIG3* | 7.7 |
| *OSGIN2--COL6A3* | 7.7 |
| *OSGIN1--CHD6* | 7.7 |
| *OSBPL9--FAF1* | 7.7 |
| *OSBPL8--NAP1L1* | 7.7 |
| *OS9--VPS13D* | 7.7 |
| *OS9--TBCK* | 7.7 |
| *OS9--TAF7L* | 7.7 |
| *ORAI2--CNNM4* | 7.7 |
| *OR7H1P--OR7E55P* | 7.7 |
| *OR2A14--OR2A15P* | 7.7 |
| *ONECUT2--SHROOM3* | 7.7 |
| *ONECUT2--ADAM9* | 7.7 |
| *OMA1--ITGAV* | 7.7 |
| *OIP5-AS1--DIS3* | 7.7 |
| *OGT--SMG7* | 7.7 |
| *OGN--COL1A2* | 7.7 |
| *OGDHL--GOLM1* | 7.7 |
| *OGDH--PSMD7* | 7.7 |
| *OGA--COL1A1* | 7.7 |
| *ODF2L--CP* | 7.7 |
| *OBSCN--CHGB* | 7.7 |
| *OAZ2--NOL10* | 7.7 |
| *NYNRIN--KIAA1549* | 7.7 |
| *NXPE3--TMEM106A* | 7.7 |
| *NVL--TUBB* | 7.7 |
| *NVL--MALAT1* | 7.7 |

| | |
|---|---|
| *NUTM2A-AS1--MUC4* | 7.7 |
| *NUP62--ACTB* | 7.7 |
| *NUP58--DBI* | 7.7 |
| *NUP214--TTN-AS1* | 7.7 |
| *NUP214--DHX9* | 7.7 |
| *NUP210--ADAMTS1* | 7.7 |
| *NUP205--STMP1* | 7.7 |
| *NUP155--MALAT1* | 7.7 |
| *NUP153--DGCR8* | 7.7 |
| *NUP153--AL513165.2* | 7.7 |
| *NUP133--YTHDC1* | 7.7 |
| *NUMA1--ZNF483* | 7.7 |
| *NUFIP2--EIF4G1* | 7.7 |
| *NUF2--ZMYND8* | 7.7 |
| *NUCKS1--SCAMP5* | 7.7 |
| *NUCKS1--LETM1* | 7.7 |
| *NUCKS1--CPD* | 7.7 |
| *NUCKS1--ATP8B2* | 7.7 |
| *NTRK2--FAT1* | 7.7 |
| *NTHL1--TRIM25* | 7.7 |
| *NT5DC2--SNRPB* | 7.7 |
| *NT5C3A--TSPYL4* | 7.7 |
| *NT5C3A--ALG8* | 7.7 |
| *NSMF--DNMT1* | 7.7 |
| *NSF--MBNL1* | 7.7 |
| *NSD3--CAMSAP2* | 7.7 |
| *NSD1--ZFP36L1* | 7.7 |
| *NSD1--RPL14* | 7.7 |
| *NSD1--MALAT1* | 7.7 |
| *NRDE2--RMRP* | 7.7 |
| *NRDC--ZNF644* | 7.7 |
| *NRCAM--NEB* | 7.7 |
| *NR6A1--ATP5F1B* | 7.7 |
| *NR4A1--ACTG1* | 7.7 |
| *NR2F2--SV2B* | 7.7 |
| *NR2F2--DAZAP2* | 7.7 |
| *NR2F1-AS1--ULK4* | 7.7 |
| *NQO1--NADK* | 7.7 |
| *NQO1--HSPA8* | 7.7 |
| *NQO1--BBS5* | 7.7 |
| *NPTN--ESYT1* | 7.7 |
| *NPM1--PAX8* | 7.7 |
| *NPM1--LINC01193* | 7.7 |
| *NPM1--BRWD1* | 7.7 |
| *NPLOC4--RAI1* | 7.7 |

| | |
|---|---|
| *NPIPB5--ZNF347* | 7.7 |
| *NPHP4--MALAT1* | 7.7 |
| *NPEPL1--EEF1A1* | 7.7 |
| *NPC1--AHSA2P* | 7.7 |
| *NPAT--USP9X* | 7.7 |
| *NOTUM--RERE* | 7.7 |
| *NOTCH3--CACNA1A* | 7.7 |
| *NOTCH2--SMG1* | 7.7 |
| *NOTCH2--MUC4* | 7.7 |
| *NOTCH2--FTL* | 7.7 |
| *NORAD--ZNF551* | 7.7 |
| *NOL9--AC138811.2* | 7.7 |
| *NOC2L--NUP62* | 7.7 |
| *NMNAT3--RNF213* | 7.7 |
| *NMNAT1--AC044839.1* | 7.7 |
| *NME6--GAS6* | 7.7 |
| *NLRP12--ZC3H13* | 7.7 |
| *NKIRAS2--NEB* | 7.7 |
| *NKD1--NSMAF* | 7.7 |
| *NISCH--SMG1* | 7.7 |
| *NIPBL-DT--TUT4* | 7.7 |
| *NIPBL--VPS13C* | 7.7 |
| *NIPBL--ATP2A2* | 7.7 |
| *NIBAN1--DSP* | 7.7 |
| *NHP2--RPL30* | 7.7 |
| *NHLRC3--NIPBL* | 7.7 |
| *NHLRC2--NAGK* | 7.7 |
| *NFX1--LDLR* | 7.7 |
| *NFX1--AUXG01000058.1* | 7.7 |
| *NFKBIZ--UTP15* | 7.7 |
| *NFIB--PSIP1* | 7.7 |
| *NFIA--TBC1D25* | 7.7 |
| *NFIA--SH3BP4* | 7.7 |
| *NFE2L2--IGKC* | 7.7 |
| *NFE2L2--ABCD3* | 7.7 |
| *NFE2L1--XIST* | 7.7 |
| *NFAT5--UTRN* | 7.7 |
| *NFAT5--MALAT1* | 7.7 |
| *NFAT5--EIF3K* | 7.7 |
| *NF1--PACSIN2* | 7.7 |
| *NF1--LPP* | 7.7 |
| *NEURL4--NDUFC1* | 7.7 |
| *NES--TOR1AIP2* | 7.7 |
| *NEO1--AC124312.3* | 7.7 |
| *NEMF--FGFR2* | 7.7 |
| *NEK4--PEBP1* | 7.7 |
| *NEFL--KAT6A* | 7.7 |
| *NEDD1--TNRC6B* | 7.7 |
| *NEBL--MED13L* | 7.7 |
| *NEB--XIST* | 7.7 |
| *NEB--USP31* | 7.7 |
| *NEB--NIPBL* | 7.7 |
| *NEB--MALAT1* | 7.7 |
| *NEB--ETV5* | 7.7 |
| *NEB--BAG2* | 7.7 |
| *NEAT1--USP34* | 7.7 |
| *NEAT1--UHMK1* | 7.7 |
| *NEAT1--UCHL1* | 7.7 |
| *NEAT1--SPATA6* | 7.7 |
| *NEAT1--RNF213* | 7.7 |
| *NEAT1--RMRP* | 7.7 |
| *NEAT1--RBBP4* | 7.7 |
| *NEAT1--RARRES1* | 7.7 |
| *NEAT1--PTPN22* | 7.7 |
| *NEAT1--PHF3* | 7.7 |
| *NEAT1--NCOA2* | 7.7 |
| *NEAT1--MUC4* | 7.7 |
| *NEAT1--MAP3K7* | 7.7 |
| *NEAT1--LUC7L3* | 7.7 |
| *NEAT1--LDHA* | 7.7 |
| *NEAT1--KPNB1* | 7.7 |
| *NEAT1--KAT6B* | 7.7 |
| *NEAT1--HMGXB4* | 7.7 |
| *NEAT1--GTF2I* | 7.7 |
| *NEAT1--GART* | 7.7 |
| *NEAT1--FTL* | 7.7 |
| *NEAT1--CNPY4* | 7.7 |
| *NEAT1--CDK12* | 7.7 |
| *NEAT1--CCNT1* | 7.7 |
| *NEAT1--ABCD3* | 7.7 |
| *NDUFS5--CSE1L* | 7.7 |
| *NDUFS1--SCN9A* | 7.7 |
| *NDUFS1--RSPO4* | 7.7 |
| *NDUFS1--DMXL2* | 7.7 |
| *NDUFB5--GLUD1* | 7.7 |
| *NDUFB3--AL157871.3* | 7.7 |
| *NDUFA6--OSBPL6* | 7.7 |
| *NDUFA10--HDAC4* | 7.7 |
| *NDST1--SESN2* | 7.7 |
| *NDRG1--DPP8* | 7.7 |
| *NDNF--COPS6* | 7.7 |
| *NDFIP1--LMNB1* | 7.7 |
| *NDE1--RC3H2* | 7.7 |
| *NCOR2--SRGAP2* | 7.7 |
| *NCOR2--EP400* | 7.7 |
| *NCOR1--PLEC* | 7.7 |
| *NCOR1--PIK3CA* | 7.7 |
| *NCOA6--PSPH* | 7.7 |
| *NCOA3--SUSD6* | 7.7 |
| *NCOA3--ARHGAP29* | 7.7 |
| *NCOA3--ADD1* | 7.7 |
| *NCOA1--STK38L* | 7.7 |
| *NCL--PHB2* | 7.7 |
| *NCL--PALM3* | 7.7 |
| *NCL--PABPC4* | 7.7 |
| *NCL--MMP15* | 7.7 |
| *NCL--KIF1C* | 7.7 |
| *NCL--ACTG1* | 7.7 |
| *NCKAP1--MALAT1* | 7.7 |
| *NCKAP1--CLCN6* | 7.7 |
| *NCF1B--NCF1* | 7.7 |
| *NCAPH2--RNF213* | 7.7 |
| *NCAPD2--ACTB* | 7.7 |
| *NBPF9--PRSS23* | 7.7 |
| *NBPF9--NEAT1* | 7.7 |
| *NBPF8--NEAT1* | 7.7 |
| *NBPF3--SCAF11* | 7.7 |
| *NBPF13P--AC239859.1* | 7.7 |
| *NBN--EVI2B* | 7.7 |
| *NBEAL1--NEAT1* | 7.7 |
| *NAV2--EPB41L5* | 7.7 |
| *NAV2--CHST10* | 7.7 |
| *NAT10--ZNF808* | 7.7 |
| *NAT10--RCAN1* | 7.7 |
| *NAT10--PCNX1* | 7.7 |
| *NARF--YLPM1* | 7.7 |
| *NAPG--TRIM34* | 7.7 |
| *NAP1L1--TMCO6* | 7.7 |
| *NAP1L1--MARK1* | 7.7 |
| *NAMPT--NEAT1* | 7.7 |
| *NAGPA--GFM1* | 7.7 |
| *NACA--COL5A2* | 7.7 |
| *NAA50--DSP* | 7.7 |
| *NAA16--TAB2* | 7.7 |

| | |
|---|---|
| *N4BP2--UHRF2* | 7.7 |
| *N4BP1--MSL3* | 7.7 |
| *MYOM3--SFT2D3* | 7.7 |
| *MYO9B--ZNF385A* | 7.7 |
| *MYO9B--IBTK* | 7.7 |
| *MYO6--TMSB4X* | 7.7 |
| *MYO6--DAPK1* | 7.7 |
| *MYO6--APLP2* | 7.7 |
| *MYO5C--NEBL* | 7.7 |
| *MYO3B--ISYNA1* | 7.7 |
| *MYO1F--AARS2* | 7.7 |
| *MYO1D--MNAT1* | 7.7 |
| *MYO1B--CDK2* | 7.7 |
| *MYO19--MALAT1* | 7.7 |
| *MYO10--SH3GLB1* | 7.7 |
| *MYLK--ZBTB7A* | 7.7 |
| *MYLK--AC092445.1* | 7.7 |
| *MYL12A--PSAP* | 7.7 |
| *MYH9--SOX13* | 7.7 |
| *MYH9--GFOD2* | 7.7 |
| *MYH3--TNNT3* | 7.7 |
| *MYH11--TTC3* | 7.7 |
| *MYH11--SCARNA5* | 7.7 |
| *MYCN--LINC01128* | 7.7 |
| *MYCBP2--NAGK* | 7.7 |
| *MYC--MYH11* | 7.7 |
| *MYBL2--PPP6R1* | 7.7 |
| *MYBBP1A--PRPF8* | 7.7 |
| *MYADM--F5* | 7.7 |
| *MVP--OCIAD1* | 7.7 |
| *MVK--TSKU* | 7.7 |
| *MUC5B--SRRM2* | 7.7 |
| *MUC5B--MARCH8* | 7.7 |
| *MUC5B--CEP57* | 7.7 |
| *MUC4--WWOX* | 7.7 |
| *MUC4--TMSB4X* | 7.7 |
| *MUC4--MAPK13* | 7.7 |
| *MUC4--DDX47* | 7.7 |
| *MUC4--CIT* | 7.7 |
| *MUC4--CALR* | 7.7 |
| *MUC4--ADAM10* | 7.7 |
| *MUC16--TPR* | 7.7 |
| *MUC16--NEAT1* | 7.7 |
| *MUC16--KIAA0100* | 7.7 |
| *MUC16--FUS* | 7.7 |
| *MTREX--USP9X* | 7.7 |
| *MTOR--SYP* | 7.7 |
| *MTMR4--DNAAF2* | 7.7 |
| *MTMR3--HIST1H2AE* | 7.7 |
| *MTMR2--TLE5* | 7.7 |
| *MTMR10--STT3B* | 7.7 |
| *MTMR10--AARS2* | 7.7 |
| *MTHFD2--KANSL3* | 7.7 |
| *MTHFD1--ANO10* | 7.7 |
| *MTFMT--MALAT1* | 7.7 |
| *MTERF4--RPL10A* | 7.7 |
| *MTCH2--CNTNAP4* | 7.7 |
| *MSRB3--FMO5* | 7.7 |
| *MSN--MALAT1* | 7.7 |
| *MSN--APLN* | 7.7 |
| *MSMO1--FTH1* | 7.7 |
| *MSI2--IFT57* | 7.7 |
| *MSI2--DCAF7* | 7.7 |
| *MSH6--REV3L* | 7.7 |
| *MSH6--AL021155.5* | 7.7 |
| *MSH3--PKM* | 7.7 |
| *MSANTD4--ADGRV1* | 7.7 |
| *MRTFA--XPNPEP3* | 7.7 |
| *MRPS27--PIK3C2B* | 7.7 |
| *MRPS26--FXYD6-FXYD2* | 7.7 |
| *MRPS25--HOMER3* | 7.7 |
| *MRPS18A--LLPH-DT* | 7.7 |
| *MRPL45--PPIG* | 7.7 |
| *MRPL39--COL1A2* | 7.7 |
| *MRPL32--MALAT1* | 7.7 |
| *MRPL1--GDI1* | 7.7 |
| *MRFAP1L1--NIPA1* | 7.7 |
| *MREG--SCG2* | 7.7 |
| *MRC1--COMMD2* | 7.7 |
| *MPRIP--CSRP1* | 7.7 |
| *MPP6--QSER1* | 7.7 |
| *MPP6--BBX* | 7.7 |
| *MPC2--MALAT1* | 7.7 |
| *MORF4L1--MALAT1* | 7.7 |
| *MORC4--MALAT1* | 7.7 |
| *MORC3--TTN* | 7.7 |
| *MORC3--HMGN2* | 7.7 |
| *MON2--FTL* | 7.7 |
| *MOB3A--EIF4G3* | 7.7 |
| *MOB1B--EXOC4* | 7.7 |
| *MOAP1--ANP32C* | 7.7 |
| *MNT--MUC4* | 7.7 |
| *MMADHC--DNMT3A* | 7.7 |
| *MLLT3--MALAT1* | 7.7 |
| *MLLT10--CREBZF* | 7.7 |
| *MLH3--CNN3* | 7.7 |
| *MKRN1--NINL* | 7.7 |
| *MKLN1--RPS17* | 7.7 |
| *MKLN1--MALAT1* | 7.7 |
| *MKI67--SAP130* | 7.7 |
| *MKI67--PTPRK* | 7.7 |
| *MKI67--NUCKS1* | 7.7 |
| *MITF--GLIS2* | 7.7 |
| *MIR100HG--PABPC1* | 7.7 |
| *MINDY2--MAPK6* | 7.7 |
| *MICALL1--MALAT1* | 7.7 |
| *MICAL3--MALAT1* | 7.7 |
| *MIAT--SGSH* | 7.7 |
| *MGRN1--FBXO45* | 7.7 |
| *MGA--NSF* | 7.7 |
| *MGA--MALAT1* | 7.7 |
| *MFSD6--WNK1* | 7.7 |
| *MFSD4B--GCNT1* | 7.7 |
| *MFN1--CRYBG3* | 7.7 |
| *MFHAS1--NIPBL* | 7.7 |
| *METTL21A--NEDD4* | 7.7 |
| *MET--KIF21A* | 7.7 |
| *MEP1A--TAF4B* | 7.7 |
| *MEIOB--TLN2* | 7.7 |
| *MEGF10--XIST* | 7.7 |
| *MED28--FREM2* | 7.7 |
| *MED13L--GOT1* | 7.7 |
| *MED12--SERPINA1* | 7.7 |
| *MED12--LINC00632* | 7.7 |
| *MECP2--RYR2* | 7.7 |
| *MEAF6--TFAP2A* | 7.7 |
| *MDN1--AKAP9* | 7.7 |
| *MDM4--CTCF* | 7.7 |
| *MDM2--HSPA9* | 7.7 |
| *MDM2--CWF19L2* | 7.7 |
| *MDM1--EIF4G2* | 7.7 |
| *MDK--SOGA1* | 7.7 |
| *MDC1--LARGE1* | 7.7 |
| *MDC1--GTF2I* | 7.7 |

| | |
|---|---|
| *MCTP2--ZNF292* | 7.7 |
| *MCOLN3--MALAT1* | 7.7 |
| *MCM9--GJB3* | 7.7 |
| *MCM7--AC019117.3* | 7.7 |
| *MCM6--PASK* | 7.7 |
| *MCM3--TRG@* | 7.7 |
| *MCL1--ABCC2* | 7.7 |
| *MCL1--AATF* | 7.7 |
| *MCFD2--RMRP* | 7.7 |
| *MCCC2--DIS3L2* | 7.7 |
| *MCCC1--INTS1* | 7.7 |
| *MBTD1--PTP4A2* | 7.7 |
| *MBP--ODC1* | 7.7 |
| *MBD1--PCDHGA10* | 7.7 |
| *MAX--DTWD1* | 7.7 |
| *MAU2--IDH1* | 7.7 |
| *MATR3--FOXP1* | 7.7 |
| *MASTL--RNF213* | 7.7 |
| *MARS--RMRP* | 7.7 |
| *MARCKS--DDX5* | 7.7 |
| *MARCH6--HSBP1* | 7.7 |
| *MAPKAPK2--PCID2* | 7.7 |
| *MAPK1IP1L--PEX19* | 7.7 |
| *MAP7D2--NOL12* | 7.7 |
| *MAP4K4--PDK2* | 7.7 |
| *MAP3K4--MIR9-3HG* | 7.7 |
| *MAP3K4--MALAT1* | 7.7 |
| *MAP3K13--AVPR1A* | 7.7 |
| *MAP3K10--LINC01833* | 7.7 |
| *MANEA--NINL* | 7.7 |
| *MAN2B1--AC005332.6* | 7.7 |
| *MAN2A2--APEH* | 7.7 |
| *MAN2A1--WDR6* | 7.7 |
| *MAN2A1--RICTOR* | 7.7 |
| *MAN2A1--MALAT1* | 7.7 |
| *MAN1B1--MFSD14B* | 7.7 |
| *MAN1A2--RPS8* | 7.7 |
| *MAN1A2--KNL1* | 7.7 |
| *MAML1--SALL1* | 7.7 |
| *MAML1--IL6ST* | 7.7 |
| *MALT1--ARHGAP35* | 7.7 |
| *MALRD1--AL845552.1* | 7.7 |
| *MALAT1--ZZEF1* | 7.7 |
| *MALAT1--ZNF93* | 7.7 |
| *MALAT1--ZNF708* | 7.7 |
| *MALAT1--ZNF347* | 7.7 |
| *MALAT1--ZNF33A* | 7.7 |
| *MALAT1--ZNF24* | 7.7 |
| *MALAT1--ZNF143* | 7.7 |
| *MALAT1--ZMIZ2* | 7.7 |
| *MALAT1--ZFHX3* | 7.7 |
| *MALAT1--ZFC3H1* | 7.7 |
| *MALAT1--ZEB2* | 7.7 |
| *MALAT1--ZCCHC7* | 7.7 |
| *MALAT1--ZC3HAV1* | 7.7 |
| *MALAT1--ZBTB20* | 7.7 |
| *MALAT1--YWHAZ* | 7.7 |
| *MALAT1--YTHDF3* | 7.7 |
| *MALAT1--YIPF3* | 7.7 |
| *MALAT1--YBX1* | 7.7 |
| *MALAT1--XRN1* | 7.7 |
| *MALAT1--XRCC6* | 7.7 |
| *MALAT1--XRCC5* | 7.7 |
| *MALAT1--XPO6* | 7.7 |
| *MALAT1--XPO1* | 7.7 |
| *MALAT1--WWOX* | 7.7 |
| *MALAT1--WTAP* | 7.7 |
| *MALAT1--WSB1* | 7.7 |
| *MALAT1--WNK1* | 7.7 |
| *MALAT1--WDFY3* | 7.7 |
| *MALAT1--WASHC4* | 7.7 |
| *MALAT1--WASH5P* | 7.7 |
| *MALAT1--WAPL* | 7.7 |
| *MALAT1--VWA8* | 7.7 |
| *MALAT1--VPS8* | 7.7 |
| *MALAT1--VPS50* | 7.7 |
| *MALAT1--VPS13C* | 7.7 |
| *MALAT1--VCAN* | 7.7 |
| *MALAT1--VAPB* | 7.7 |
| *MALAT1--USP7* | 7.7 |
| *MALAT1--USF2* | 7.7 |
| *MALAT1--UQCRB* | 7.7 |
| *MALAT1--UQCC1* | 7.7 |
| *MALAT1--ULK1* | 7.7 |
| *MALAT1--UBR5* | 7.7 |
| *MALAT1--UBR4* | 7.7 |
| *MALAT1--UBE2R2* | 7.7 |
| *MALAT1--UBAP2L* | 7.7 |
| *MALAT1--UBAP2* | 7.7 |
| *MALAT1--TXNRD1* | 7.7 |
| *MALAT1--TUT7* | 7.7 |
| *MALAT1--TTN* | 7.7 |
| *MALAT1--TTC21B* | 7.7 |
| *MALAT1--TTC19* | 7.7 |
| *MALAT1--TTC14* | 7.7 |
| *MALAT1--TRAM1* | 7.7 |
| *MALAT1--TRAK1* | 7.7 |
| *MALAT1--TRA@* | 7.7 |
| *MALAT1--TPD52* | 7.7 |
| *MALAT1--TOPBP1* | 7.7 |
| *MALAT1--TOP1* | 7.7 |
| *MALAT1--TNS3* | 7.7 |
| *MALAT1--TMEM131* | 7.7 |
| *MALAT1--TMED5* | 7.7 |
| *MALAT1--TMED2* | 7.7 |
| *MALAT1--TMCO6* | 7.7 |
| *MALAT1--TMBIM6* | 7.7 |
| *MALAT1--TM4SF1-AS1* | 7.7 |
| *MALAT1--TJP1* | 7.7 |
| *MALAT1--TF* | 7.7 |
| *MALAT1--TET3* | 7.7 |
| *MALAT1--TCERG1* | 7.7 |
| *MALAT1--TC2N* | 7.7 |
| *MALAT1--TAS2R30* | 7.7 |
| *MALAT1--TANGO2* | 7.7 |
| *MALAT1--TALDO1* | 7.7 |
| *MALAT1--TAF6* | 7.7 |
| *MALAT1--TAF1D* | 7.7 |
| *MALAT1--TAF15* | 7.7 |
| *MALAT1--SYT13* | 7.7 |
| *MALAT1--SYNE1* | 7.7 |
| *MALAT1--SYAP1* | 7.7 |
| *MALAT1--SUZ12* | 7.7 |
| *MALAT1--SUPT5H* | 7.7 |
| *MALAT1--STRN4* | 7.7 |
| *MALAT1--STAT1* | 7.7 |
| *MALAT1--STAMBP* | 7.7 |
| *MALAT1--ST5* | 7.7 |
| *MALAT1--SRSF11* | 7.7 |
| *MALAT1--SRSF10* | 7.7 |
| *MALAT1--SRSF1* | 7.7 |
| *MALAT1--SRRM2* | 7.7 |
| *MALAT1--SQSTM1* | 7.7 |
| *MALAT1--SPTAN1* | 7.7 |
| *MALAT1--SPG7* | 7.7 |

| | |
|---|---|
| *MALAT1--SPG11* | 7.7 |
| *MALAT1--SPEN* | 7.7 |
| *MALAT1--SPARC* | 7.7 |
| *MALAT1--SPAG9* | 7.7 |
| *MALAT1--SON* | 7.7 |
| *MALAT1--SOCS4* | 7.7 |
| *MALAT1--SNX29* | 7.7 |
| *MALAT1--SMG5* | 7.7 |
| *MALAT1--SMCHD1* | 7.7 |
| *MALAT1--SMC5* | 7.7 |
| *MALAT1--SLC44A3* | 7.7 |
| *MALAT1--SLC44A1* | 7.7 |
| *MALAT1--SLC40A1* | 7.7 |
| *MALAT1--SLC39A11* | 7.7 |
| *MALAT1--SLC29A1* | 7.7 |
| *MALAT1--SLC25A37* | 7.7 |
| *MALAT1--SLC25A22* | 7.7 |
| *MALAT1--SLC17A5* | 7.7 |
| *MALAT1--SIPA1L1* | 7.7 |
| *MALAT1--SIN3B* | 7.7 |
| *MALAT1--SGPL1* | 7.7 |
| *MALAT1--SF1* | 7.7 |
| *MALAT1--SETD5* | 7.7 |
| *MALAT1--SET* | 7.7 |
| *MALAT1--SERINC1* | 7.7 |
| *MALAT1--SERBP1* | 7.7 |
| *MALAT1--SEMA4B* | 7.7 |
| *MALAT1--SEL1L3* | 7.7 |
| *MALAT1--SEC11C* | 7.7 |
| *MALAT1--SDHC* | 7.7 |
| *MALAT1--SCRN1* | 7.7 |
| *MALAT1--SCN8A* | 7.7 |
| *MALAT1--SCARNA7* | 7.7 |
| *MALAT1--SCARNA13* | 7.7 |
| *MALAT1--SCAPER* | 7.7 |
| *MALAT1--SAMD4B* | 7.7 |
| *MALAT1--SAFB* | 7.7 |
| *MALAT1--RTTN* | 7.7 |
| *MALAT1--RTCA-AS1* | 7.7 |
| *MALAT1--RSRC1* | 7.7 |
| *MALAT1--RRBP1* | 7.7 |
| *MALAT1--RPS9* | 7.7 |
| *MALAT1--RPS8* | 7.7 |
| *MALAT1--RPS27* | 7.7 |
| *MALAT1--RPS11* | 7.7 |

| | |
|---|---|
| *MALAT1--RPS10-NUDT3* | 7.7 |
| *MALAT1--RPRD2* | 7.7 |
| *MALAT1--RPN2* | 7.7 |
| *MALAT1--RPLP1* | 7.7 |
| *MALAT1--RPL5* | 7.7 |
| *MALAT1--RPL35A* | 7.7 |
| *MALAT1--RPL32* | 7.7 |
| *MALAT1--RPL22* | 7.7 |
| *MALAT1--ROBO1* | 7.7 |
| *MALAT1--RNF10* | 7.7 |
| *MALAT1--RN7SL151P* | 7.7 |
| *MALAT1--RN7SL1* | 7.7 |
| *MALAT1--RMND5B* | 7.7 |
| *MALAT1--RIPOR2* | 7.7 |
| *MALAT1--RIMS2* | 7.7 |
| *MALAT1--RIBC1* | 7.7 |
| *MALAT1--RHOBTB3* | 7.7 |
| *MALAT1--REV3L* | 7.7 |
| *MALAT1--RETREG3* | 7.7 |
| *MALAT1--RERE* | 7.7 |
| *MALAT1--REPS1* | 7.7 |
| *MALAT1--REL* | 7.7 |
| *MALAT1--RCAN3* | 7.7 |
| *MALAT1--RBM41* | 7.7 |
| *MALAT1--RBM26* | 7.7 |
| *MALAT1--RBL2* | 7.7 |
| *MALAT1--RBBP4* | 7.7 |
| *MALAT1--RASA1* | 7.7 |
| *MALAT1--RAPGEF4* | 7.7 |
| *MALAT1--RAPGEF2* | 7.7 |
| *MALAT1--RAF1* | 7.7 |
| *MALAT1--RAD21* | 7.7 |
| *MALAT1--RAB5A* | 7.7 |
| *MALAT1--RAB3IP* | 7.7 |
| *MALAT1--RAB22A* | 7.7 |
| *MALAT1--PXK* | 7.7 |
| *MALAT1--PTCH2* | 7.7 |
| *MALAT1--PTCD3* | 7.7 |
| *MALAT1--PTBP2* | 7.7 |
| *MALAT1--PSME3* | 7.7 |
| *MALAT1--PSMD11* | 7.7 |
| *MALAT1--PSIP1* | 7.7 |
| *MALAT1--PRRC2B* | 7.7 |
| *MALAT1--PRR11* | 7.7 |

| | |
|---|---|
| *MALAT1--PROX1* | 7.7 |
| *MALAT1--PRKACA* | 7.7 |
| *MALAT1--PRKAB1* | 7.7 |
| *MALAT1--PRIM1* | 7.7 |
| *MALAT1--PPP4R3B* | 7.7 |
| *MALAT1--PPP1CB* | 7.7 |
| *MALAT1--PPIL4* | 7.7 |
| *MALAT1--PNN* | 7.7 |
| *MALAT1--PMS1* | 7.7 |
| *MALAT1--PLSCR1* | 7.7 |
| *MALAT1--PLOD2* | 7.7 |
| *MALAT1--PLEKHA2* | 7.7 |
| *MALAT1--PLEC* | 7.7 |
| *MALAT1--PLCG2* | 7.7 |
| *MALAT1--PLA2G4B* | 7.7 |
| *MALAT1--PKP4* | 7.7 |
| *MALAT1--PITPNB* | 7.7 |
| *MALAT1--PI4KA* | 7.7 |
| *MALAT1--PHKA2* | 7.7 |
| *MALAT1--PGM5* | 7.7 |
| *MALAT1--PGM3* | 7.7 |
| *MALAT1--PFN2* | 7.7 |
| *MALAT1--PEX13* | 7.7 |
| *MALAT1--PDS5B* | 7.7 |
| *MALAT1--PDIA3* | 7.7 |
| *MALAT1--PDHB* | 7.7 |
| *MALAT1--PCLO* | 7.7 |
| *MALAT1--PBX3* | 7.7 |
| *MALAT1--PARP14* | 7.7 |
| *MALAT1--PAPOLA* | 7.7 |
| *MALAT1--PANK2* | 7.7 |
| *MALAT1--PABPC1L* | 7.7 |
| *MALAT1--PABPC1* | 7.7 |
| *MALAT1--OXNAD1* | 7.7 |
| *MALAT1--OTULIN* | 7.7 |
| *MALAT1--OSBPL3* | 7.7 |
| *MALAT1--OSBPL1A* | 7.7 |
| *MALAT1--ORC3* | 7.7 |
| *MALAT1--ONECUT2* | 7.7 |
| *MALAT1--NUP153* | 7.7 |
| *MALAT1--NSD2* | 7.7 |
| *MALAT1--NRDC* | 7.7 |
| *MALAT1--NQO1* | 7.7 |
| *MALAT1--NPM1* | 7.7 |
| *MALAT1--NPIPB5* | 7.7 |

| | |
|---|---|
| *MALAT1--NPEPPS* | 7.7 |
| *MALAT1--NOTCH2* | 7.7 |
| *MALAT1--NOS3* | 7.7 |
| *MALAT1--NORAD* | 7.7 |
| *MALAT1--NHLRC2* | 7.7 |
| *MALAT1--NF1* | 7.7 |
| *MALAT1--NET1* | 7.7 |
| *MALAT1--NES* | 7.7 |
| *MALAT1--NEK9* | 7.7 |
| *MALAT1--NEBL* | 7.7 |
| *MALAT1--NEB* | 7.7 |
| *MALAT1--NCOA3* | 7.7 |
| *MALAT1--NCLN* | 7.7 |
| *MALAT1--NCL* | 7.7 |
| *MALAT1--NACA* | 7.7 |
| *MALAT1--N4BP2* | 7.7 |
| *MALAT1--MYRF* | 7.7 |
| *MALAT1--MYO9A* | 7.7 |
| *MALAT1--MYL6* | 7.7 |
| *MALAT1--MUC4* | 7.7 |
| *MALAT1--MUC16* | 7.7 |
| *MALAT1--MUC1* | 7.7 |
| *MALAT1--MRC2* | 7.7 |
| *MALAT1--MORF4L2* | 7.7 |
| *MALAT1--MOB1A* | 7.7 |
| *MALAT1--MLX* | 7.7 |
| *MALAT1--MKI67* | 7.7 |
| *MALAT1--MIOS* | 7.7 |
| *MALAT1--MGAT4A* | 7.7 |
| *MALAT1--MED23* | 7.7 |
| *MALAT1--MED15* | 7.7 |
| *MALAT1--MED13L* | 7.7 |
| *MALAT1--MED13* | 7.7 |
| *MALAT1--MBTD1* | 7.7 |
| *MALAT1--MBNL2* | 7.7 |
| *MALAT1--MAVS* | 7.7 |
| *MALAT1--MAT2A* | 7.7 |
| *MALAT1--MAST2* | 7.7 |
| *MALAT1--MAP4K4* | 7.7 |
| *MALAT1--MAP2K4* | 7.7 |
| *MALAT1--MALT1* | 7.7 |
| *MALAT1--LUM* | 7.7 |
| *MALAT1--LUC7L3* | 7.7 |
| *MALAT1--LRP8* | 7.7 |
| *MALAT1--LRBA* | 7.7 |
| *MALAT1--LMO7* | 7.7 |
| *MALAT1--LINC00910* | 7.7 |
| *MALAT1--LEO1* | 7.7 |
| *MALAT1--LDLR* | 7.7 |
| *MALAT1--LDHA* | 7.7 |
| *MALAT1--LARS2* | 7.7 |
| *MALAT1--LAMB1* | 7.7 |
| *MALAT1--LAMA3* | 7.7 |
| *MALAT1--LAMA1* | 7.7 |
| *MALAT1--KRT7* | 7.7 |
| *MALAT1--KRT18* | 7.7 |
| *MALAT1--KPNB1* | 7.7 |
| *MALAT1--KMT2C* | 7.7 |
| *MALAT1--KMT2A* | 7.7 |
| *MALAT1--KIF11* | 7.7 |
| *MALAT1--KIAA2012* | 7.7 |
| *MALAT1--KIAA1841* | 7.7 |
| *MALAT1--KHDRBS1* | 7.7 |
| *MALAT1--KDM3B* | 7.7 |
| *MALAT1--KCNQ4* | 7.7 |
| *MALAT1--KCNQ1OT1* | 7.7 |
| *MALAT1--KCNK6* | 7.7 |
| *MALAT1--KAT6B* | 7.7 |
| *MALAT1--KANSL1* | 7.7 |
| *MALAT1--JUP* | 7.7 |
| *MALAT1--IQGAP1* | 7.7 |
| *MALAT1--IPO7* | 7.7 |
| *MALAT1--INTS2* | 7.7 |
| *MALAT1--INPP1* | 7.7 |
| *MALAT1--ING1* | 7.7 |
| *MALAT1--IL6ST* | 7.7 |
| *MALAT1--IGF1R* | 7.7 |
| *MALAT1--IFT122* | 7.7 |
| *MALAT1--HSPD1* | 7.7 |
| *MALAT1--HSPA9* | 7.7 |
| *MALAT1--HSPA4* | 7.7 |
| *MALAT1--HSP90AA1* | 7.7 |
| *MALAT1--HRG* | 7.7 |
| *MALAT1--HNRNPR* | 7.7 |
| *MALAT1--HNRNPH1* | 7.7 |
| *MALAT1--HNRNPC* | 7.7 |
| *MALAT1--HMGN2* | 7.7 |
| *MALAT1--HMGA1* | 7.7 |
| *MALAT1--HMCN1* | 7.7 |
| *MALAT1--HELZ* | 7.7 |
| *MALAT1--HDLBP* | 7.7 |
| *MALAT1--HCFC2* | 7.7 |
| *MALAT1--GTF3C4* | 7.7 |
| *MALAT1--GRN* | 7.7 |
| *MALAT1--GPBP1* | 7.7 |
| *MALAT1--GPATCH2L* | 7.7 |
| *MALAT1--GOLGA8A* | 7.7 |
| *MALAT1--GOLGA3* | 7.7 |
| *MALAT1--GMNN* | 7.7 |
| *MALAT1--GCLC* | 7.7 |
| *MALAT1--GART* | 7.7 |
| *MALAT1--GAPDH* | 7.7 |
| *MALAT1--GALT* | 7.7 |
| *MALAT1--GABPB2* | 7.7 |
| *MALAT1--FUS* | 7.7 |
| *MALAT1--FTH1* | 7.7 |
| *MALAT1--FRS2* | 7.7 |
| *MALAT1--FREM2* | 7.7 |
| *MALAT1--FOXP1* | 7.7 |
| *MALAT1--FNIP2* | 7.7 |
| *MALAT1--FMNL2* | 7.7 |
| *MALAT1--FLT4* | 7.7 |
| *MALAT1--FEM1A* | 7.7 |
| *MALAT1--FBXW2* | 7.7 |
| *MALAT1--FBXL3* | 7.7 |
| *MALAT1--FBXL13* | 7.7 |
| *MALAT1--FAM95C* | 7.7 |
| *MALAT1--FAM13A* | 7.7 |
| *MALAT1--EXTL3* | 7.7 |
| *MALAT1--EXOC3* | 7.7 |
| *MALAT1--ETV6* | 7.7 |
| *MALAT1--ETS2* | 7.7 |
| *MALAT1--ETF1* | 7.7 |
| *MALAT1--ERGIC2* | 7.7 |
| *MALAT1--ERBIN* | 7.7 |
| *MALAT1--EPRS* | 7.7 |
| *MALAT1--EP400* | 7.7 |
| *MALAT1--ENTPD4* | 7.7 |
| *MALAT1--ENPP2* | 7.7 |
| *MALAT1--EML4* | 7.7 |
| *MALAT1--ELP2* | 7.7 |
| *MALAT1--EIF4G3* | 7.7 |
| *MALAT1--EIF4G2* | 7.7 |
| *MALAT1--EIF4B* | 7.7 |
| *MALAT1--EIF4A3* | 7.7 |

| | |
|---|---|
| *MALAT1--ECT2* | 7.7 |
| *MALAT1--EBF1* | 7.7 |
| *MALAT1--DYRK1A* | 7.7 |
| *MALAT1--DOCK7* | 7.7 |
| *MALAT1--DHCR24* | 7.7 |
| *MALAT1--DGKH* | 7.7 |
| *MALAT1--DERL2* | 7.7 |
| *MALAT1--DEPP1* | 7.7 |
| *MALAT1--CUL9* | 7.7 |
| *MALAT1--CUL1* | 7.7 |
| *MALAT1--CTR9* | 7.7 |
| *MALAT1--CTPS1* | 7.7 |
| *MALAT1--CTDSP2* | 7.7 |
| *MALAT1--CSE1L* | 7.7 |
| *MALAT1--CRIPAK* | 7.7 |
| *MALAT1--CREB3L2* | 7.7 |
| *MALAT1--CPSF6* | 7.7 |
| *MALAT1--COPA* | 7.7 |
| *MALAT1--COL6A3* | 7.7 |
| *MALAT1--COA1* | 7.7 |
| *MALAT1--CNTRL* | 7.7 |
| *MALAT1--CNOT1* | 7.7 |
| *MALAT1--CNKSR2* | 7.7 |
| *MALAT1--CLSTN3* | 7.7 |
| *MALAT1--CLCN7* | 7.7 |
| *MALAT1--CHD2* | 7.7 |
| *MALAT1--CFAP36* | 7.7 |
| *MALAT1--CEP192* | 7.7 |
| *MALAT1--CENPF* | 7.7 |
| *MALAT1--CELSR2* | 7.7 |
| *MALAT1--CELF2-AS2* | 7.7 |
| *MALAT1--CDKL2* | 7.7 |
| *MALAT1--CDH11* | 7.7 |
| *MALAT1--CDH1* | 7.7 |
| *MALAT1--CDC42BPA* | 7.7 |
| *MALAT1--CD63* | 7.7 |
| *MALAT1--CD24* | 7.7 |
| *MALAT1--CCT6A* | 7.7 |
| *MALAT1--CCND3* | 7.7 |
| *MALAT1--CCDC191* | 7.7 |
| *MALAT1--CBLB* | 7.7 |
| *MALAT1--CARD8* | 7.7 |
| *MALAT1--CAPRIN1* | 7.7 |
| *MALAT1--CAMSAP1* | 7.7 |
| *MALAT1--CALD1* | 7.7 |

| | |
|---|---|
| *MALAT1--CADM1* | 7.7 |
| *MALAT1--C3ORF80* | 7.7 |
| *MALAT1--C3AR1* | 7.7 |
| *MALAT1--C15ORF40* | 7.7 |
| *MALAT1--BTN3A2* | 7.7 |
| *MALAT1--BTAF1* | 7.7 |
| *MALAT1--BMP2K* | 7.7 |
| *MALAT1--BIVM-ERCC5* | 7.7 |
| *MALAT1--BCLAF1* | 7.7 |
| *MALAT1--BACH2* | 7.7 |
| *MALAT1--ATXN1* | 7.7 |
| *MALAT1--ATP8B1* | 7.7 |
| *MALAT1--ATP13A3* | 7.7 |
| *MALAT1--ATP11A* | 7.7 |
| *MALAT1--ATM* | 7.7 |
| *MALAT1--ASPM* | 7.7 |
| *MALAT1--ARID4A* | 7.7 |
| *MALAT1--ARID2* | 7.7 |
| *MALAT1--ARHGEF7* | 7.7 |
| *MALAT1--ARHGEF12* | 7.7 |
| *MALAT1--ARHGAP1* | 7.7 |
| *MALAT1--API5* | 7.7 |
| *MALAT1--APC* | 7.7 |
| *MALAT1--AP001888.1* | 7.7 |
| *MALAT1--ANP32D* | 7.7 |
| *MALAT1--ANKRD18A* | 7.7 |
| *MALAT1--ANAPC1* | 7.7 |
| *MALAT1--ALDH5A1* | 7.7 |
| *MALAT1--ALDH1A1* | 7.7 |
| *MALAT1--AL512356.1* | 7.7 |
| *MALAT1--AL360020.1* | 7.7 |
| *MALAT1--AL157400.3* | 7.7 |
| *MALAT1--AL139353.1* | 7.7 |
| *MALAT1--AL136295.4* | 7.7 |
| *MALAT1--AL109811.3* | 7.7 |
| *MALAT1--AL031595.2* | 7.7 |
| *MALAT1--AKR1B10* | 7.7 |
| *MALAT1--AIG1* | 7.7 |
| *MALAT1--AHNAK* | 7.7 |
| *MALAT1--AFMID* | 7.7 |
| *MALAT1--AEBP1* | 7.7 |
| *MALAT1--ADGRV1* | 7.7 |
| *MALAT1--ADGRD1* | 7.7 |
| *MALAT1--ADAM17* | 7.7 |
| *MALAT1--ACTB* | 7.7 |

| | |
|---|---|
| *MALAT1--ACP6* | 7.7 |
| *MALAT1--ACACA* | 7.7 |
| *MALAT1--AC245884.4* | 7.7 |
| *MALAT1--AC245033.1* | 7.7 |
| *MALAT1--AC118549.1* | 7.7 |
| *MALAT1--AC109517.1* | 7.7 |
| *MALAT1--AC106741.1* | 7.7 |
| *MALAT1--AC100830.1* | 7.7 |
| *MALAT1--AC090227.2* | 7.7 |
| *MALAT1--AC023509.1* | 7.7 |
| *MALAT1--AC021087.5* | 7.7 |
| *MALAT1--AC012676.5* | 7.7 |
| *MALAT1--AC006148.1* | 7.7 |
| *MALAT1--ABHD2* | 7.7 |
| *MALAT1--ABCB1* | 7.7 |
| *MALAT1--ABCA5* | 7.7 |
| *MALAT1--ABCA3* | 7.7 |
| *MALAT1--ABCA1* | 7.7 |
| *MALAT1--A1CF* | 7.7 |
| *MAGI3--AHNAK* | 7.7 |
| *MACF1--UMPS* | 7.7 |
| *MACF1--TXNIP* | 7.7 |
| *MACF1--THRAP3* | 7.7 |
| *MACF1--KDM4A* | 7.7 |
| *MACF1--INPP5B* | 7.7 |
| *MACF1--GTF2I* | 7.7 |
| *MACF1--GDI1* | 7.7 |
| *MACF1--FANCI* | 7.7 |
| *MACF1--FAM120A* | 7.7 |
| *MACF1--EFCAB14* | 7.7 |
| *MACF1--AHNAK* | 7.7 |
| *M6PR--XIST* | 7.7 |
| *LXN--CLOCK* | 7.7 |
| *LUC7L3--RBM15* | 7.7 |
| *LUC7L3--MALAT1* | 7.7 |
| *LUC7L3--HSPA8* | 7.7 |
| *LTBP3--CPXM1* | 7.7 |
| *LTA4H--SLC35E1* | 7.7 |
| *LTA4H--HSPB1* | 7.7 |
| *LSP1--UBASH3B* | 7.7 |
| *LSM14B--ZNF638* | 7.7 |
| *LSAMP--MADD* | 7.7 |
| *LRRIQ1--ZNF664* | 7.7 |
| *LRRCC1--TBL1XR1* | 7.7 |
| *LRRC57--CEP170B* | 7.7 |

| | |
|---|---|
| *LRRC52-AS1--GOLGA6L2* | 7.7 |
| *LRRC3B--TAB2* | 7.7 |
| *LRRC37A4P--MALAT1* | 7.7 |
| *LRPPRC--MALAT1* | 7.7 |
| *LRP10--PCNX1* | 7.7 |
| *LRP1--HERC1* | 7.7 |
| *LRMP--PLEC* | 7.7 |
| *LRIG2--MED9* | 7.7 |
| *LRCH3--MALAT1* | 7.7 |
| *LRATD2--NDUFS8* | 7.7 |
| *LPIN1--ALB* | 7.7 |
| *LPGAT1--TPD52* | 7.7 |
| *LOX--RACGAP1* | 7.7 |
| *LONRF1--MLYCD* | 7.7 |
| *LONP2--SYT13* | 7.7 |
| *LONP2--STRIP1* | 7.7 |
| *LONP1--ATP6V1G2-DDX39B* | 7.7 |
| *LNPEP--JPT2* | 7.7 |
| *LMNB2--LAS1L* | 7.7 |
| *LMAN1--STX3* | 7.7 |
| *LINC02234--VPS13B* | 7.7 |
| *LINC02145--CHD9* | 7.7 |
| *LINC00863--NUTM2A* | 7.7 |
| *LINC00630--BAZ2A* | 7.7 |
| *LINC00476--DDX5* | 7.7 |
| *LINC00115--ZNF596* | 7.7 |
| *LIN54--AC048338.2* | 7.7 |
| *LIMCH1--FXYD2* | 7.7 |
| *LIMCH1--CTDSP2* | 7.7 |
| *LILRB5--LILRP1* | 7.7 |
| *LIG3--OSTC* | 7.7 |
| *LIG3--IGKC* | 7.7 |
| *LIG3--IGK@* | 7.7 |
| *LIG3--GOLGA4* | 7.7 |
| *LIG3--AHCYL1* | 7.7 |
| *LHFPL6--NUP98* | 7.7 |
| *LGR4--ITPA* | 7.7 |
| *LGALS8--MALAT1* | 7.7 |
| *LGALS8--LINC01876* | 7.7 |
| *LGALS3BP--NCSTN* | 7.7 |
| *LGALS1--UBAC2* | 7.7 |
| *LFNG--SPECC1L-ADORA2A* | 7.7 |
| *LEPROTL1--SMG1* | 7.7 |
| *LENG8--SERPINB6* | 7.7 |

| | |
|---|---|
| *LENG8--MYCL* | 7.7 |
| *LDLR--PRODH2* | 7.7 |
| *LCP1--CENPJ* | 7.7 |
| *LCORL--CTNNB1* | 7.7 |
| *LBR--EIF4A2* | 7.7 |
| *LATS1--AL021546.1* | 7.7 |
| *LASP1--HSPD1* | 7.7 |
| *LARP4B--FTH1* | 7.7 |
| *LARP4B--AP3B2* | 7.7 |
| *LARP1--RPPH1* | 7.7 |
| *LARP1--ENO2* | 7.7 |
| *LARGE2--NSD1* | 7.7 |
| *LAPTM4A--SZT2* | 7.7 |
| *LAMC3--SLC12A4* | 7.7 |
| *LAMC3--KIF23* | 7.7 |
| *LAMC1--TGFBR2* | 7.7 |
| *LAMC1--PICALM* | 7.7 |
| *LAMB1--MALAT1* | 7.7 |
| *LAMB1--HIST1H1D* | 7.7 |
| *LAMB1--DQX1* | 7.7 |
| *LAMA1--VWDE* | 7.7 |
| *LAMA1--SIN3B* | 7.7 |
| *KTN1--PSMA3* | 7.7 |
| *KRT8--MYO6* | 7.7 |
| *KRT18--NEBL* | 7.7 |
| *KRT18--MALAT1* | 7.7 |
| *KRT18--H6PD* | 7.7 |
| *KRT18--BBS9* | 7.7 |
| *KRT15--DHX9* | 7.7 |
| *KPNB1--WDR74* | 7.7 |
| *KPNB1--VPS54* | 7.7 |
| *KPNB1--SND1* | 7.7 |
| *KPNB1--NSUN4* | 7.7 |
| *KPNB1--NEK4* | 7.7 |
| *KPNB1--MALAT1* | 7.7 |
| *KPNB1--GNA12* | 7.7 |
| *KPNB1--CAPN7* | 7.7 |
| *KPNB1--ARID1A* | 7.7 |
| *KPNB1--AKAP13* | 7.7 |
| *KPNA6--MGAT4B* | 7.7 |
| *KPNA2--AC005833.1* | 7.7 |
| *KNSTRN--GUF1* | 7.7 |
| *KNDC1--GPAT4* | 7.7 |
| *KMT5C--COL1A2* | 7.7 |
| *KMT5B--PDGFC* | 7.7 |

| | |
|---|---|
| *KMT2E--SPON2* | 7.7 |
| *KMT2E--IGK@* | 7.7 |
| *KMT2E--CACNA2D1* | 7.7 |
| *KMT2D--PKN2* | 7.7 |
| *KMT2D--GPI* | 7.7 |
| *KMT2C--TTLL7* | 7.7 |
| *KMT2C--RUFY3* | 7.7 |
| *KMT2C--MALAT1* | 7.7 |
| *KMT2C--CAST* | 7.7 |
| *KMT2A--SMARCA2* | 7.7 |
| *KMT2A--RAB7A* | 7.7 |
| *KLK12--CRY1* | 7.7 |
| *KLHL9--RGS7* | 7.7 |
| *KLHL7--RNF43* | 7.7 |
| *KLHL7--IRF3* | 7.7 |
| *KLHL42--MALAT1* | 7.7 |
| *KLHL28--MUC4* | 7.7 |
| *KLHDC3--MYO1B* | 7.7 |
| *KLHDC2--SMG1* | 7.7 |
| *KLF7--PPIB* | 7.7 |
| *KLF6--MALAT1* | 7.7 |
| *KLC2--ZNF470* | 7.7 |
| *KIFC1--RNF213* | 7.7 |
| *KIF3B--HERC2* | 7.7 |
| *KIF3A--HSPD1* | 7.7 |
| *KIF2A--HNRNPU* | 7.7 |
| *KIF21A--TLK1* | 7.7 |
| *KIF21A--NRAS* | 7.7 |
| *KIF21A--KIAA1522* | 7.7 |
| *KIF1C--PIGT* | 7.7 |
| *KIF1B--RPPH1* | 7.7 |
| *KIF1B--CTNNBIP1* | 7.7 |
| *KIF1A--MED13* | 7.7 |
| *KIF11--MUC4* | 7.7 |
| *KIAA2026--DSP* | 7.7 |
| *KIAA1671--MALAT1* | 7.7 |
| *KIAA1549--MALAT1* | 7.7 |
| *KIAA1324--TNFRSF11B* | 7.7 |
| *KIAA1324--TFG* | 7.7 |
| *KIAA1211L--B4GALT1* | 7.7 |
| *KIAA1109--TUG1* | 7.7 |
| *KIAA1109--SCFD1* | 7.7 |
| *KIAA1109--MAP1B* | 7.7 |
| *KIAA0895L--PTPRA* | 7.7 |
| *KIAA0895--KIF21A* | 7.7 |

| | |
|---|---|
| *KIAA0556--MALAT1* | 7.7 |
| *KIAA0355--KCTD20* | 7.7 |
| *KIAA0100--XIST* | 7.7 |
| *KIAA0100--MALAT1* | 7.7 |
| *KIAA0100--DHCR24* | 7.7 |
| *KHDC4--SOX12* | 7.7 |
| *KDR--SFMBT1* | 7.7 |
| *KDM6A--VCAN* | 7.7 |
| *KDM6A--VANGL2* | 7.7 |
| *KDM5A--PI4KA* | 7.7 |
| *KDM5A--MALAT1* | 7.7 |
| *KDM1A--CNOT7* | 7.7 |
| *KCTD10--SBF2* | 7.7 |
| *KCNQ1OT1--TTN* | 7.7 |
| *KCNQ1OT1--MAP4K4* | 7.7 |
| *KCNMA1--NAGK* | 7.7 |
| *KCNE4--FNDC3A* | 7.7 |
| *KCMF1--PTPRM* | 7.7 |
| *KBTBD8--DSP* | 7.7 |
| *KAT6B--TBL1XR1* | 7.7 |
| *KAT6B--PLEKHO2* | 7.7 |
| *KAT6A--AP1G1* | 7.7 |
| *KANSL1--ARID1A* | 7.7 |
| *JUN--VDAC3* | 7.7 |
| *JPT2--HMCN1* | 7.7 |
| *JMJD1C--P4HA1* | 7.7 |
| *JAK1--PAPSS1* | 7.7 |
| *JAK1--LPAR6* | 7.7 |
| *JAK1--APCDD1* | 7.7 |
| *JAG1--STK36* | 7.7 |
| *IWS1--USP40* | 7.7 |
| *IWS1--PRRC2C* | 7.7 |
| *IVNS1ABP--GLUL* | 7.7 |
| *IVNS1ABP--DEUP1* | 7.7 |
| *ITSN2--TMEM127* | 7.7 |
| *ITSN1--UBC* | 7.7 |
| *ITPR3--SIGLEC1* | 7.7 |
| *ITPR3--ESYT1* | 7.7 |
| *ITPR2--RABGAP1L* | 7.7 |
| *ITPR2--FGFR1OP2* | 7.7 |
| *ITPR1--ARL8B* | 7.7 |
| *ITM2B--RMRP* | 7.7 |
| *ITM2B--ACTB* | 7.7 |
| *ITGB4--DIP2A* | 7.7 |
| *ITGAV--ZFPM2* | 7.7 |
| *ITGAV--MALAT1* | 7.7 |
| *ITGA7--AC004951.1* | 7.7 |
| *ITGA6--TSHZ1* | 7.7 |
| *ITGA4--UBR5* | 7.7 |
| *ITGA1--PISD* | 7.7 |
| *ITGA1--MPHOSPH9* | 7.7 |
| *ISL1--FXR1* | 7.7 |
| *IRF1--SMG1* | 7.7 |
| *IRAK1--SCARNA2* | 7.7 |
| *IQGAP3--SUZ12* | 7.7 |
| *IQGAP1--SIN3A* | 7.7 |
| *IQGAP1--IFI16* | 7.7 |
| *IQGAP1--FTL* | 7.7 |
| *IQCN--DTNA* | 7.7 |
| *IQCA1--TPR* | 7.7 |
| *IQCA1--CMTR1* | 7.7 |
| *IPO9--ATXN10* | 7.7 |
| *IPO5--DLG1* | 7.7 |
| *IPO5--COL1A2* | 7.7 |
| *INVS--FLVCR1* | 7.7 |
| *INTS8--KIAA1522* | 7.7 |
| *INTS7--RMRP* | 7.7 |
| *INTS3--HSP90B1* | 7.7 |
| *INTS10--MALAT1* | 7.7 |
| *INTS1--GALNS* | 7.7 |
| *INO80D--MCM3* | 7.7 |
| *ING5--MALAT1* | 7.7 |
| *IMPAD1--HLCS* | 7.7 |
| *IMMT--PCYT1B* | 7.7 |
| *ILVBL--MKI67* | 7.7 |
| *ILRUN--RBM20* | 7.7 |
| *ILRUN--BICD1* | 7.7 |
| *ILKAP--NF1* | 7.7 |
| *ILF3--SEC24B* | 7.7 |
| *ILF3--FTL* | 7.7 |
| *ILF3--DHX15* | 7.7 |
| *IL6ST--ZFP62* | 7.7 |
| *IL33--ZNF562* | 7.7 |
| *IL20RB--MALAT1* | 7.7 |
| *IL1R1--RPS4X* | 7.7 |
| *IL13RA1--WNK1* | 7.7 |
| *IL10RA--VIM* | 7.7 |
| *IGKV1-5--RBMS1* | 7.7 |
| *IGK@--RBM14* | 7.7 |
| *IGK@--LSP1* | 7.7 |
| *IGK@--HIST1H2BD* | 7.7 |
| *IGK@--HGD* | 7.7 |
| *IGK@--H19* | 7.7 |
| *IGK@--DUX4* | 7.7 |
| *IGK@--COL1A2* | 7.7 |
| *IGK@--CNTRL* | 7.7 |
| *IGK@--CDK6* | 7.7 |
| *IGK@--AC010976.1* | 7.7 |
| *IGHV3-30--CD44* | 7.7 |
| *IGHG1--UGGT2* | 7.7 |
| *IGHG1--SDK1* | 7.7 |
| *IGH@--WWOX* | 7.7 |
| *IGH@--MALT1* | 7.7 |
| *IGH@--CD44* | 7.7 |
| *IGH@--BCL11A* | 7.7 |
| *IGH@--AL137139.2* | 7.7 |
| *IGFN1--APEH* | 7.7 |
| *IGFN1--AL133243.3* | 7.7 |
| *IGFBP2--UBC* | 7.7 |
| *IGF2R--UNC13B* | 7.7 |
| *IGF2R--AK2* | 7.7 |
| *IFT74--BRIP1* | 7.7 |
| *IFT172--AHNAK* | 7.7 |
| *IFNGR2--MALAT1* | 7.7 |
| *IFNAR1--POLR2C* | 7.7 |
| *IFITM3--YAP1* | 7.7 |
| *IFITM1--METAP2* | 7.7 |
| *IDH2--POMP* | 7.7 |
| *IDH2--MAP4* | 7.7 |
| *IDH2--HTATIP2* | 7.7 |
| *IDH1--MALAT1* | 7.7 |
| *IDH1--HECTD4* | 7.7 |
| *ID1--NAP1L1* | 7.7 |
| *ICMT--MALAT1* | 7.7 |
| *ICE2--SLC43A1* | 7.7 |
| *ICE1--SNF8* | 7.7 |
| *IBTK--MALAT1* | 7.7 |
| *IBTK--INHBA* | 7.7 |
| *IBTK--GRN* | 7.7 |
| *IARS2--MALAT1* | 7.7 |
| *IARS--MALAT1* | 7.7 |
| *HYOU1--CASZ1* | 7.7 |
| *HUWE1--SP110* | 7.7 |
| *HUWE1--MALAT1* | 7.7 |
| *HUWE1--LMNB2* | 7.7 |

| | |
|---|---|
| *HUWE1--CDCA5* | 7.7 |
| *HUWE1--AK2* | 7.7 |
| *HUWE1--AC106785.1* | 7.7 |
| *HTATSF1--NCOR2* | 7.7 |
| *HSPG2--RPL8* | 7.7 |
| *HSPG2--MALAT1* | 7.7 |
| *HSPE1-MOB4--HNRNPLL* | 7.7 |
| *HSPD1--UMPS* | 7.7 |
| *HSPD1--NEAT1* | 7.7 |
| *HSPD1--HERC2* | 7.7 |
| *HSPD1--H19* | 7.7 |
| *HSPD1--DSP* | 7.7 |
| *HSPD1--COLGALT1* | 7.7 |
| *HSPA9--SCMH1* | 7.7 |
| *HSPA9--PTPRF* | 7.7 |
| *HSPA9--MATR3* | 7.7 |
| *HSPA9--MALAT1* | 7.7 |
| *HSPA8--PFKM* | 7.7 |
| *HSPA8--PDIA4* | 7.7 |
| *HSPA8--LTBR* | 7.7 |
| *HSPA5--LMNA* | 7.7 |
| *HSPA5--DST* | 7.7 |
| *HSPA5--ARFGAP1* | 7.7 |
| *HSPA4--HNRNPU* | 7.7 |
| *HSPA1B--DIAPH1* | 7.7 |
| *HSPA1A--RPS6KC1* | 7.7 |
| *HSP90B1--TGOLN2* | 7.7 |
| *HSP90B1--MALAT1* | 7.7 |
| *HSP90B1--CDYL* | 7.7 |
| *HSP90B1--BCR* | 7.7 |
| *HSP90AB1--MAP1B* | 7.7 |
| *HSP90AB1--MALAT1* | 7.7 |
| *HSP90AB1--FN1* | 7.7 |
| *HSP90AA1--SYNE2* | 7.7 |
| *HSP90AA1--BCL6* | 7.7 |
| *HSDL2--TMEM86A* | 7.7 |
| *HSDL1--EIF2A* | 7.7 |
| *HSD17B4--UBR4* | 7.7 |
| *HSD17B4--RARS* | 7.7 |
| *HSD17B4--PUS7L* | 7.7 |
| *HSD11B2--SSH1* | 7.7 |
| *HPF1--AC122713.1* | 7.7 |
| *HP1BP3--DZANK1* | 7.7 |
| *HP1BP3--DLGAP5* | 7.7 |
| *HOXD8--RNF213* | 7.7 |
| *HOXD8--HIPK2* | 7.7 |
| *HOOK3--SLC16A1* | 7.7 |
| *HOOK3--LINC00632* | 7.7 |
| *HNRNPUL2-BSCL2--PTPRF* | 7.7 |
| *HNRNPUL2-BSCL2--KANSL1* | 7.7 |
| *HNRNPUL1--GALNT7* | 7.7 |
| *HNRNPUL1--AC021087.5* | 7.7 |
| *HNRNPU--UBC* | 7.7 |
| *HNRNPU--TANC1* | 7.7 |
| *HNRNPU--SSR1* | 7.7 |
| *HNRNPU--SLC25A6* | 7.7 |
| *HNRNPU--RC3H1* | 7.7 |
| *HNRNPU--CNTNAP2* | 7.7 |
| *HNRNPU--AEBP1* | 7.7 |
| *HNRNPU--AC016588.2* | 7.7 |
| *HNRNPR--ITPKB* | 7.7 |
| *HNRNPL--AFG3L1P* | 7.7 |
| *HNRNPK--CYBA* | 7.7 |
| *HNRNPH3--MFN1* | 7.7 |
| *HNRNPH3--AL162171.3* | 7.7 |
| *HNRNPH1--WDR59* | 7.7 |
| *HNRNPH1--ODF2L* | 7.7 |
| *HNRNPH1--MPHOSPH9* | 7.7 |
| *HNRNPH1--IL6ST* | 7.7 |
| *HNRNPH1--CLIC1* | 7.7 |
| *HNRNPH1--ANKRA2* | 7.7 |
| *HNRNPF--CD2AP* | 7.7 |
| *HNRNPDL--SRRM2* | 7.7 |
| *HNRNPDL--ONECUT2* | 7.7 |
| *HNRNPDL--CSNK1D* | 7.7 |
| *HNRNPD--DDX1* | 7.7 |
| *HNRNPC--RALGAPA2* | 7.7 |
| *HNRNPC--MALAT1* | 7.7 |
| *HNRNPC--ICE2* | 7.7 |
| *HNRNPC--HERPUD1* | 7.7 |
| *HNRNPA3--ADHFE1* | 7.7 |
| *HNRNPA2B1--ZFYVE26* | 7.7 |
| *HNRNPA2B1--WWOX* | 7.7 |
| *HNRNPA2B1--SPTAN1* | 7.7 |
| *HNRNPA2B1--SCN8A* | 7.7 |
| *HNRNPA2B1--RMRP* | 7.7 |
| *HNRNPA2B1--MALAT1* | 7.7 |
| *HNRNPA2B1--DENND4B* | 7.7 |
| *HNRNPA2B1--AC093512.2* | 7.7 |
| *HNRNPA1--PRKAA2* | 7.7 |
| *HNRNPA1--GFPT1* | 7.7 |
| *HMGN1--PCSK7* | 7.7 |
| *HMGN1--EEF1A1* | 7.7 |
| *HMGN1--COL5A2* | 7.7 |
| *HMGN1--APP* | 7.7 |
| *HMGCS1--KMT2C* | 7.7 |
| *HMGB3--WDR83* | 7.7 |
| *HMGB2--PHF14* | 7.7 |
| *HMGB2--COL1A2* | 7.7 |
| *HMGB1--UBE4B* | 7.7 |
| *HMGB1--FCN1* | 7.7 |
| *HMGB1--AC008725.1* | 7.7 |
| *HMCN1--VIM* | 7.7 |
| *HMCN1--MTCH2* | 7.7 |
| *HMCN1--HMGB3* | 7.7 |
| *HMCN1--COL5A2* | 7.7 |
| *HLTF--PTK2* | 7.7 |
| *HLTF--GTF3C1* | 7.7 |
| *HLCS--MUC16* | 7.7 |
| *HK1--MALAT1* | 7.7 |
| *HIST1H4L--MPRIP* | 7.7 |
| *HIST1H3C--ZNF720* | 7.7 |
| *HIST1H2BJ--TPM2* | 7.7 |
| *HIST1H2BH--PRSS16* | 7.7 |
| *HIST1H2BF--ZNF592* | 7.7 |
| *HIST1H2BC--ZNRF1* | 7.7 |
| *HIST1H2AL--ISYNA1* | 7.7 |
| *HIST1H2AB--TPR* | 7.7 |
| *HIST1H1E--PSAP* | 7.7 |
| *HIST1H1E--POLR2A* | 7.7 |
| *HIST1H1E--KHNYN* | 7.7 |
| *HIPK3--ATP2A2* | 7.7 |
| *HIPK2--MALAT1* | 7.7 |
| *HIPK1--TTBK2* | 7.7 |
| *HIPK1--NEAT1* | 7.7 |
| *HIPK1--HSP90AB1* | 7.7 |
| *HIGD2A--VPS9D1-AS1* | 7.7 |
| *HIF1A--CACNA1C* | 7.7 |
| *HID1--ADAR* | 7.7 |
| *HGS--SCG5* | 7.7 |
| *HERC4--GALE* | 7.7 |

| | |
|---|---|
| *HERC2--SIGMAR1* | 7.7 |
| *HERC2--PROX1* | 7.7 |
| *HERC2--MALAT1* | 7.7 |
| *HERC2--IGK@* | 7.7 |
| *HERC2--ATXN2* | 7.7 |
| *HERC2--APP* | 7.7 |
| *HERC2--AP4E1* | 7.7 |
| *HERC2--AL138966.2* | 7.7 |
| *HERC1--MALAT1* | 7.7 |
| *HERC1--HELLPAR* | 7.7 |
| *HELZ--TPI1* | 7.7 |
| *HELZ--RNMT* | 7.7 |
| *HELLPAR--TPM3* | 7.7 |
| *HELB--CDC6* | 7.7 |
| *HECTD4--PTCH1* | 7.7 |
| *HECTD4--MALAT1* | 7.7 |
| *HECTD1--URB1* | 7.7 |
| *HECTD1--TLK1* | 7.7 |
| *HECTD1--MALAT1* | 7.7 |
| *HECTD1--AC004805.1* | 7.7 |
| *HDLBP--MAX* | 7.7 |
| *HDGFL2--MIGA2* | 7.7 |
| *HDGF--RBM3* | 7.7 |
| *HDC--TTC14* | 7.7 |
| *HDAC2--ZNF141* | 7.7 |
| *HDAC2--NSD3* | 7.7 |
| *HDAC1--SMARCA4* | 7.7 |
| *HDAC1--PRORP* | 7.7 |
| *HDAC1--KPNA6* | 7.7 |
| *HCCS--SCARNA5* | 7.7 |
| *HBP1--CDK6* | 7.7 |
| *HAUS1--REL* | 7.7 |
| *HASPIN--ELOVL7* | 7.7 |
| *HASPIN--AKAP9* | 7.7 |
| *HAS2--ZNF572* | 7.7 |
| *HADH--NKD1* | 7.7 |
| *HACD4--RSBN1* | 7.7 |
| *HACD3--AC245033.1* | 7.7 |
| *H3F3A--SPEN* | 7.7 |
| *H1F0--HELLS* | 7.7 |
| *H1F0--ALDH7A1* | 7.7 |
| *H19--ZNF639* | 7.7 |
| *H19--ZNF496* | 7.7 |
| *H19--NUP98* | 7.7 |
| *H19--MTHFD1* | 7.7 |
| *H19--MTF1* | 7.7 |
| *H19--MSI2* | 7.7 |
| *H19--HIVEP1* | 7.7 |
| *GUCY1A2--XIST* | 7.7 |
| *GUCD1--ATRAID* | 7.7 |
| *GTPBP1--MOV10* | 7.7 |
| *GTF3C4--RPS11* | 7.7 |
| *GTF3C2--DDX23* | 7.7 |
| *GTF3C1--ACADVL* | 7.7 |
| *GTF2I--PAWR* | 7.7 |
| *GTF2I--NEK9* | 7.7 |
| *GTF2I--MALAT1* | 7.7 |
| *GTF2I--HDLBP* | 7.7 |
| *GTF2I--APH1A* | 7.7 |
| *GSTP1--MALAT1* | 7.7 |
| *GSR--SMG1* | 7.7 |
| *GSPT1--RPS4X* | 7.7 |
| *GSE1--TAF1* | 7.7 |
| *GSE1--MTMR3* | 7.7 |
| *GSE1--MALAT1* | 7.7 |
| *GRSF1--EEF1D* | 7.7 |
| *GRK2--TCN2* | 7.7 |
| *GRIPAP1--GSTK1* | 7.7 |
| *GRHPR--C1QC* | 7.7 |
| *GRAMD2B--USP24* | 7.7 |
| *GPR65--XIST* | 7.7 |
| *GPR108--FN1* | 7.7 |
| *GPI--STAB2* | 7.7 |
| *GPI--EEF2* | 7.7 |
| *GPC6--PRRC2C* | 7.7 |
| *GPBP1L1--DDX58* | 7.7 |
| *GPBP1--COMMD10* | 7.7 |
| *GPATCH2--DDX21* | 7.7 |
| *GOSR2--RANBP2* | 7.7 |
| *GORASP2--BLVRA* | 7.7 |
| *GON4L--SDR9C7* | 7.7 |
| *GOLM1--SDHB* | 7.7 |
| *GOLM1--DA750114* | 7.7 |
| *GOLGB1--MALAT1* | 7.7 |
| *GOLGB1--ITGA3* | 7.7 |
| *GOLGA6L10--XIST* | 7.7 |
| *GOLGA4--LRRFIP2* | 7.7 |
| *GOLGA3--APPBP2* | 7.7 |
| *GOLGA3--AC104472.3* | 7.7 |
| *GNPTAB--MALAT1* | 7.7 |
| *GNPNAT1--FMNL2* | 7.7 |
| *GNL3L--PPIP5K2* | 7.7 |
| *GNL3L--FTL* | 7.7 |
| *GNL3--DSG2* | 7.7 |
| *GNE--RNF213* | 7.7 |
| *GNE--EDEM3* | 7.7 |
| *GNB4--CTNND1* | 7.7 |
| *GNB1--NOP56* | 7.7 |
| *GNB1--AL031777.3* | 7.7 |
| *GNAS--SMG1* | 7.7 |
| *GNAS--AD000090.1* | 7.7 |
| *GMNN--NID1* | 7.7 |
| *GLUL--XPO6* | 7.7 |
| *GLUL--DOCK10* | 7.7 |
| *GLUL--CEP350* | 7.7 |
| *GLRX2--ATXN7L3B* | 7.7 |
| *GLG1--KIAA1324* | 7.7 |
| *GLB1L--EEF2* | 7.7 |
| *GJA1--MALAT1* | 7.7 |
| *GJA1--CD74* | 7.7 |
| *GINS1--MALAT1* | 7.7 |
| *GIGYF2--MALAT1* | 7.7 |
| *GGA1--EP300* | 7.7 |
| *GFPT1--UBA1* | 7.7 |
| *GFPT1--LAPTM4A* | 7.7 |
| *GFM1--MALAT1* | 7.7 |
| *GDPD3--GNB1* | 7.7 |
| *GDNF--CCT3* | 7.7 |
| *GDI2--RPL8* | 7.7 |
| *GDI2--MMP24OS* | 7.7 |
| *GDI2--CSDE1* | 7.7 |
| *GDI2--C21ORF91* | 7.7 |
| *GDI1--DCP2* | 7.7 |
| *GDF7--EEF1A1* | 7.7 |
| *GDE1--AHNAK* | 7.7 |
| *GDAP1--TM9SF2* | 7.7 |
| *GCSH--RANBP2* | 7.7 |
| *GCN1--NDUFB11* | 7.7 |
| *GCN1--MALAT1* | 7.7 |
| *GCLC--SMG1* | 7.7 |
| *GBF1--SENP2* | 7.7 |
| *GBF1--AC004797.1* | 7.7 |
| *GBA2--FHL1* | 7.7 |
| *GBA--LINC00205* | 7.7 |
| *GBA--AC005534.1* | 7.7 |

| | |
|---|---|
| *GASK1B--GPI* | 7.7 |
| *GAS7--PSD3* | 7.7 |
| *GAS5--RERE* | 7.7 |
| *GARS--LRP10* | 7.7 |
| *GARS--LINC00472* | 7.7 |
| *GAPVD1--RNF167* | 7.7 |
| *GAPDH--XRCC6* | 7.7 |
| *GAPDH--MALAT1* | 7.7 |
| *GALNT3--PRRC2B* | 7.7 |
| *GALNT2--ABHD3* | 7.7 |
| *GALNT18--ANKRD17* | 7.7 |
| *GALNT13--GDE1* | 7.7 |
| *GALM--KPNB1* | 7.7 |
| *GALC--NEAT1* | 7.7 |
| *GABRA3--TGFBR3* | 7.7 |
| *GABPB1-AS1--NEB* | 7.7 |
| *GAB2--ARRDC3* | 7.7 |
| *G3BP1--MYH10* | 7.7 |
| *FYTTD1--LRCH3* | 7.7 |
| *FXR1--SLC29A1* | 7.7 |
| *FXR1--RMRP* | 7.7 |
| *FXR1--NUTF2* | 7.7 |
| *FXR1--CCDC59* | 7.7 |
| *FUS--RTBDN* | 7.7 |
| *FUS--NXF1* | 7.7 |
| *FUS--AL021155.5* | 7.7 |
| *FUBP1--USP33* | 7.7 |
| *FUBP1--MALAT1* | 7.7 |
| *FTX--MUC4* | 7.7 |
| *FTSJ1--PCM1* | 7.7 |
| *FTL--RPL11* | 7.7 |
| *FTL--RN7SL1* | 7.7 |
| *FTL--RBM6* | 7.7 |
| *FTL--MALAT1* | 7.7 |
| *FTL--DPYSL2* | 7.7 |
| *FTH1--WASHC5* | 7.7 |
| *FTH1--WAC* | 7.7 |
| *FTH1--VEGFA* | 7.7 |
| *FTH1--PRLHR* | 7.7 |
| *FTH1--PAH* | 7.7 |
| *FTH1--NEAT1* | 7.7 |
| *FTH1--NDEL1* | 7.7 |
| *FTH1--MRM3* | 7.7 |
| *FTH1--MDM2* | 7.7 |
| *FTH1--MALAT1* | 7.7 |

| | |
|---|---|
| *FTH1--LHPP* | 7.7 |
| *FTH1--HMGN2* | 7.7 |
| *FTH1--HIPK2* | 7.7 |
| *FTH1--HECTD3* | 7.7 |
| *FTH1--COA3* | 7.7 |
| *FTH1--CHST15* | 7.7 |
| *FTH1--CACNA2D1* | 7.7 |
| *FTH1--BDP1* | 7.7 |
| *FTH1--ADAM15* | 7.7 |
| *FSTL1--MALAT1* | 7.7 |
| *FST--MALAT1* | 7.7 |
| *FRMD3--XRCC6* | 7.7 |
| *FRK--MALAT1* | 7.7 |
| *FRG1BP--AL161457.2* | 7.7 |
| *FRG1--AL161457.2* | 7.7 |
| *FREM2--PPL* | 7.7 |
| *FPR2--DDX3X* | 7.7 |
| *FOXRED2--VWF* | 7.7 |
| *FOXP1--ZNFX1* | 7.7 |
| *FOXP1--RNF213* | 7.7 |
| *FOXP1--RABEP1* | 7.7 |
| *FOXP1--GOLGA4* | 7.7 |
| *FOXP1--EGR1* | 7.7 |
| *FOXM1--ZC3H13* | 7.7 |
| *FOXK2--ZNF652* | 7.7 |
| *FOXJ3--RMRP* | 7.7 |
| *FOS--MRTFA* | 7.7 |
| *FO538757.1--AC009533.1* | 7.7 |
| *FNDC3B--ROBO1* | 7.7 |
| *FNDC3A--DGKH* | 7.7 |
| *FNDC3A--CD2AP* | 7.7 |
| *FNDC3A--ACTB* | 7.7 |
| *FNBP1L--MALAT1* | 7.7 |
| *FN1--ZFAT* | 7.7 |
| *FN1--WDR74* | 7.7 |
| *FN1--VDAC2* | 7.7 |
| *FN1--UGDH-AS1* | 7.7 |
| *FN1--UBXN7* | 7.7 |
| *FN1--SART3* | 7.7 |
| *FN1--PABPC1* | 7.7 |
| *FN1--NPM1* | 7.7 |
| *FN1--IGF1R* | 7.7 |
| *FN1--HNRNPK* | 7.7 |
| *FN1--FNTA* | 7.7 |

| | |
|---|---|
| *FN1--FBXW2* | 7.7 |
| *FN1--ERBIN* | 7.7 |
| *FN1--CPED1* | 7.7 |
| *FN1--CAST* | 7.7 |
| *FMR1--TASOR2* | 7.7 |
| *FMR1--IREB2* | 7.7 |
| *FMC1-LUC7L2--ZNF189* | 7.7 |
| *FLOT1--MON1B* | 7.7 |
| *FLNC--CHGB* | 7.7 |
| *FLNA--MALAT1* | 7.7 |
| *FLNA--HIST1H2BH* | 7.7 |
| *FLCN--KIAA2026* | 7.7 |
| *FKBP14--ZBED1* | 7.7 |
| *FGG--SLC38A10* | 7.7 |
| *FGFR2--ZMYND11* | 7.7 |
| *FGFR1OP--IPO5* | 7.7 |
| *FGFR1--UBC* | 7.7 |
| *FGD1--MALAT1* | 7.7 |
| *FEN1--HIST1H2AM* | 7.7 |
| *FEM1B--RET* | 7.7 |
| *FEM1B--MALAT1* | 7.7 |
| *FDPS--PARP14* | 7.7 |
| *FCHSD2--TMEM106B* | 7.7 |
| *FCF1--CHCHD5* | 7.7 |
| *FCF1--AC009220.3* | 7.7 |
| *FBXW11--SEMA3C* | 7.7 |
| *FBXO34--MUC4* | 7.7 |
| *FBXO28--ANGEL2* | 7.7 |
| *FBXL3--CDK4* | 7.7 |
| *FBXL18--YBX3* | 7.7 |
| *FBXL18--UTRN* | 7.7 |
| *FBXL18--F5* | 7.7 |
| *FBRSL1--MALAT1* | 7.7 |
| *FBN1--FOXP1* | 7.7 |
| *FBLN1--PI4KA* | 7.7 |
| *FBLN1--MALAT1* | 7.7 |
| *FAT3--BAZ1A* | 7.7 |
| *FAT1--SLC38A2* | 7.7 |
| *FAT1--RAPGEF2* | 7.7 |
| *FAT1--CADPS* | 7.7 |
| *FAT1--ANKRD17* | 7.7 |
| *FASTKD2--LNPEP* | 7.7 |
| *FASN--MALAT1* | 7.7 |
| *FASN--DSP* | 7.7 |
| *FARP2--NUP155* | 7.7 |

| | |
|---|---|
| *FARP1--ZC3HAV1* | 7.7 |
| *FAP--RMRP* | 7.7 |
| *FANCI--ZNF396* | 7.7 |
| *FANCA--CNOT1* | 7.7 |
| *FAM50A--MALAT1* | 7.7 |
| *FAM50A--GOLM1* | 7.7 |
| *FAM50A--BOLA3-AS1* | 7.7 |
| *FAM47DP--FAM47C* | 7.7 |
| *FAM32A--CPLX1* | 7.7 |
| *FAM210B--CNOT1* | 7.7 |
| *FAM20C--MUC4* | 7.7 |
| *FAM182B--AC209154.1* | 7.7 |
| *FAM126A--MALAT1* | 7.7 |
| *FAM120B--STXBP5L* | 7.7 |
| *FAM120B--PSEN1* | 7.7 |
| *FAM120B--AP4B1* | 7.7 |
| *FAM118B--PRRC2A* | 7.7 |
| *FAM117A--AGMO* | 7.7 |
| *FAM110B--LEMD3* | 7.7 |
| *FAM104B--FZD7* | 7.7 |
| *FAM102A--SMG1* | 7.7 |
| *FAF1--SSBP2* | 7.7 |
| *FAF1--PPP1CB* | 7.7 |
| *F5--RPN1* | 7.7 |
| *F3--RIMS1* | 7.7 |
| *F2--DDX17* | 7.7 |
| *F12--IDH1* | 7.7 |
| *EZR--FLNA* | 7.7 |
| *EYA4--PCID2* | 7.7 |
| *EXTL3--MALAT1* | 7.7 |
| *EXTL2--KIAA0100* | 7.7 |
| *EXOC2--MALAT1* | 7.7 |
| *EXOC2--DUSP22* | 7.7 |
| *EXOC2--ANKH* | 7.7 |
| *EVC--SCARNA5* | 7.7 |
| *ETV6--MALAT1* | 7.7 |
| *ETV6--EEF1A1* | 7.7 |
| *ETV6--ATIC* | 7.7 |
| *ETS2--AL391380.1* | 7.7 |
| *ETNK1--LRRC14* | 7.7 |
| *ESCO1--DDX46* | 7.7 |
| *ESCO1--B3GALNT2* | 7.7 |
| *ERRFI1--SAMD9* | 7.7 |
| *ERRFI1--PAICS* | 7.7 |
| *ERRFI1--ANKRD11* | 7.7 |

| | |
|---|---|
| *ERO1A--SLC38A3* | 7.7 |
| *ERO1A--KIF14* | 7.7 |
| *ERLIN1--SCARB2* | 7.7 |
| *ERCC6L2--MALAT1* | 7.7 |
| *ERCC6L2--HECTD1* | 7.7 |
| *ERBIN--EID1* | 7.7 |
| *ERBB4--PERM1* | 7.7 |
| *ERBB3--MTHFD1* | 7.7 |
| *ERBB3--FZD6* | 7.7 |
| *ERAL1--WASHC4* | 7.7 |
| *EPSTI1--IGK@* | 7.7 |
| *EPS15--FAF1* | 7.7 |
| *EPM2AIP1--MALAT1* | 7.7 |
| *EPCAM--PCDHGA11* | 7.7 |
| *EPCAM--NR2C2* | 7.7 |
| *EPCAM--MALAT1* | 7.7 |
| *EPB41L5--RMRP* | 7.7 |
| *EPB41L5--APOC1* | 7.7 |
| *EPB41L3--SRXN1* | 7.7 |
| *EPB41L3--MALAT1* | 7.7 |
| *EPB41L3--INTS7* | 7.7 |
| *EP400--ATP9A* | 7.7 |
| *EP300--TCEA3* | 7.7 |
| *EP300--SOS1-IT1* | 7.7 |
| *EP300--GNA13* | 7.7 |
| *EP300--ERO1B* | 7.7 |
| *ENTPD6--AC005534.1* | 7.7 |
| *ENTPD1--MALAT1* | 7.7 |
| *ENPP2--ADGRV1* | 7.7 |
| *ENPP1--COQ8A* | 7.7 |
| *ENO1--ST3GAL2* | 7.7 |
| *ENO1--PALM2-AKAP2* | 7.7 |
| *ENO1--DENND5A* | 7.7 |
| *ENGASE--MALAT1* | 7.7 |
| *ENC1--CTSD* | 7.7 |
| *ENAH--SP100* | 7.7 |
| *ENAH--PRKDC* | 7.7 |
| *EMP3--GSN* | 7.7 |
| *EMP2--PHIP* | 7.7 |
| *EMP1--ATN1* | 7.7 |
| *EML5--EIF3D* | 7.7 |
| *EML4--POSTN* | 7.7 |
| *EML3--PLAA* | 7.7 |
| *EMC1--MALAT1* | 7.7 |
| *ELOA--EEF1A1* | 7.7 |

| | |
|---|---|
| *ELMO2--FMNL2* | 7.7 |
| *ELMO1--IRF2BP2* | 7.7 |
| *ELL2--AGAP1* | 7.7 |
| *ELK3--ATL3* | 7.7 |
| *ELF3--RAI1* | 7.7 |
| *ELAC2--PRKAA2* | 7.7 |
| *EIF5B--SPTBN1* | 7.7 |
| *EIF5B--CKAP2L* | 7.7 |
| *EIF5--MALAT1* | 7.7 |
| *EIF4G3--RBM26* | 7.7 |
| *EIF4G3--ARID1B* | 7.7 |
| *EIF4G1--GABPB1-IT1* | 7.7 |
| *EIF4ENIF1--NECTIN1* | 7.7 |
| *EIF4B--XRCC6* | 7.7 |
| *EIF4B--LCAT* | 7.7 |
| *EIF4B--ABCG1* | 7.7 |
| *EIF4A2--XIST* | 7.7 |
| *EIF4A2--HSPH1* | 7.7 |
| *EIF4A2--GMPR2* | 7.7 |
| *EIF4A2--CACNA2D1* | 7.7 |
| *EIF4A2--C19ORF44* | 7.7 |
| *EIF4A1--SEC14L1* | 7.7 |
| *EIF4A1--MALAT1* | 7.7 |
| *EIF4A1--KRT8* | 7.7 |
| *EIF4A1--DUSP6* | 7.7 |
| *EIF3H--TRPS1* | 7.7 |
| *EIF3H--SKI* | 7.7 |
| *EIF3E--XIST* | 7.7 |
| *EIF3B--MALAT1* | 7.7 |
| *EIF3A--ZNF292* | 7.7 |
| *EIF3A--SLTM* | 7.7 |
| *EIF3A--SLC12A7* | 7.7 |
| *EIF3A--MALAT1* | 7.7 |
| *EIF2S3--HSP90AA1* | 7.7 |
| *EIF2AK4--AC016876.2* | 7.7 |
| *EIF2AK1--HNRNPUL1* | 7.7 |
| *EIF2AK1--HMGB1* | 7.7 |
| *EIF2A--OTOF* | 7.7 |
| *EHMT1--ZDBF2* | 7.7 |
| *EHBP1L1--LHFPL4* | 7.7 |
| *EHBP1L1--CD276* | 7.7 |
| *EGFR--MALAT1* | 7.7 |
| *EFHD2--SIPA1L1* | 7.7 |
| *EFEMP1--MTO1* | 7.7 |
| *EFCAB14--C14ORF93* | 7.7 |

| | |
|---|---|
| *EEF2--MALAT1* | 7.7 |
| *EEF2--KIFAP3* | 7.7 |
| *EEF1B2--TMED4* | 7.7 |
| *EEF1AKMT1--COL4A1* | 7.7 |
| *EEF1A1--ZNF133* | 7.7 |
| *EEF1A1--XIST* | 7.7 |
| *EEF1A1--WDR74* | 7.7 |
| *EEF1A1--VPS13A* | 7.7 |
| *EEF1A1--UBE3C* | 7.7 |
| *EEF1A1--SYT7* | 7.7 |
| *EEF1A1--RPPH1* | 7.7 |
| *EEF1A1--RMRP* | 7.7 |
| *EEF1A1--PIEZO1* | 7.7 |
| *EEF1A1--MYH9* | 7.7 |
| *EEF1A1--MIR100HG* | 7.7 |
| *EEF1A1--MED13* | 7.7 |
| *EEF1A1--FOXP1* | 7.7 |
| *EEF1A1--FN1* | 7.7 |
| *EEF1A1--FLNA* | 7.7 |
| *EEF1A1--F10* | 7.7 |
| *EEF1A1--EPC1* | 7.7 |
| *EEF1A1--BAZ2B* | 7.7 |
| *EEF1A1--ACOT9* | 7.7 |
| *EED--UBC* | 7.7 |
| *EDRF1--AHNAK* | 7.7 |
| *EDF1--SREBF2* | 7.7 |
| *EDC3--XIST* | 7.7 |
| *ECPAS--ATP2A2* | 7.7 |
| *ECM1--ATN1* | 7.7 |
| *ECD--POMK* | 7.7 |
| *ECD--NT5E* | 7.7 |
| *EBNA1BP2--WBP1L* | 7.7 |
| *EBF1--MALAT1* | 7.7 |
| *EBF1--GNAS* | 7.7 |
| *EBF1--CNN2* | 7.7 |
| *EBF1--API5* | 7.7 |
| *EBAG9--SMG1* | 7.7 |
| *DZIP1L--NCKAP1* | 7.7 |
| *DYRK2--AL590666.2* | 7.7 |
| *DYRK1A--EFHC1* | 7.7 |
| *DYNLL1--SYNPO* | 7.7 |
| *DYNC1LI2--DFFA* | 7.7 |
| *DYNC1LI2--CAVIN1* | 7.7 |
| *DYNC1I1--COL3A1* | 7.7 |
| *DYNC1H1--WFDC2* | 7.7 |

| | |
|---|---|
| *DYNC1H1--RNPS1* | 7.7 |
| *DYNC1H1--NEB* | 7.7 |
| *DYNC1H1--ISCA1* | 7.7 |
| *DYNC1H1--ENAH* | 7.7 |
| *DYNC1H1--CASC3* | 7.7 |
| *DYNC1H1--AXIN2* | 7.7 |
| *DYNC1H1--APP* | 7.7 |
| *DVL3--MALAT1* | 7.7 |
| *DVL3--COL1A2* | 7.7 |
| *DUXAP8--XIST* | 7.7 |
| *DUSP22--KDM6A* | 7.7 |
| *DUOX1--BLOC1S5-TXNDC5* | 7.7 |
| *DTNB--CHM* | 7.7 |
| *DTNA--SERPINA1* | 7.7 |
| *DTNA--PTK2* | 7.7 |
| *DSTYK--SERINC5* | 7.7 |
| *DST--TMEM230* | 7.7 |
| *DST--RMRP* | 7.7 |
| *DST--MAZ* | 7.7 |
| *DST--AK2* | 7.7 |
| *DSP--ZXDC* | 7.7 |
| *DSP--YY1AP1* | 7.7 |
| *DSP--RMRP* | 7.7 |
| *DSP--MALAT1* | 7.7 |
| *DSP--ETV6* | 7.7 |
| *DSP--ATG4B* | 7.7 |
| *DSP--AHNAK* | 7.7 |
| *DSG2--CASC19* | 7.7 |
| *DROSHA--ELK3* | 7.7 |
| *DRG1--HIST1H4E* | 7.7 |
| *DRAXIN--NDUFS2* | 7.7 |
| *DRAIC--NCAPG2* | 7.7 |
| *DPYSL2--UROD* | 7.7 |
| *DPY19L1--IGH@* | 7.7 |
| *DPP3--MALAT1* | 7.7 |
| *DPH3--ANKRD36* | 7.7 |
| *DOP1A--MDH2* | 7.7 |
| *DOP1A--MALAT1* | 7.7 |
| *DOCK7--ZMIZ1* | 7.7 |
| *DOCK7--PEAK1* | 7.7 |
| *DOCK10--EPB41L4B* | 7.7 |
| *DOCK1--MALAT1* | 7.7 |
| *DOCK1--HSPA8* | 7.7 |
| *DOCK1--ETV1* | 7.7 |

| | |
|---|---|
| *DOCK1--AKR1E2* | 7.7 |
| *DNTTIP2--DYRK2* | 7.7 |
| *DNMT1--MAP3K4* | 7.7 |
| *DNM3--MALAT1* | 7.7 |
| *DNER--OAT* | 7.7 |
| *DNASE1L1--ZNF428* | 7.7 |
| *DNAJC5--TOM1L2* | 7.7 |
| *DNAJC5--MALAT1* | 7.7 |
| *DNAJC3--MUC20* | 7.7 |
| *DNAJC21--MTPAP* | 7.7 |
| *DNAJC2--COX20* | 7.7 |
| *DNAJC19--MALAT1* | 7.7 |
| *DNAJB6--OSGIN2* | 7.7 |
| *DNAJA3--MALAT1* | 7.7 |
| *DNAH5--HCFC2* | 7.7 |
| *DNAH11--EEF1A1* | 7.7 |
| *DNA2--CHD9* | 7.7 |
| *DMXL2--MALAT1* | 7.7 |
| *DMXL1--TPD52L2* | 7.7 |
| *DLST--FKBP11* | 7.7 |
| *DLL3--KANSL1* | 7.7 |
| *DLGAP5--EMC1* | 7.7 |
| *DLG1--PUS10* | 7.7 |
| *DLG1--N4BP1* | 7.7 |
| *DLG1--DDX24* | 7.7 |
| *DLEU2--NINL* | 7.7 |
| *DLD--PRKAA2* | 7.7 |
| *DLD--MSI2* | 7.7 |
| *DLD--BAZ2B* | 7.7 |
| *DLAT--METTL21A* | 7.7 |
| *DLAT--CEMIP2* | 7.7 |
| *DKC1--SAMD5* | 7.7 |
| *DKC1--ITPRID2* | 7.7 |
| *DKC1--AP000553.5* | 7.7 |
| *DIP2C--MALAT1* | 7.7 |
| *DICER1--UBE2I* | 7.7 |
| *DICER1--KMT2A* | 7.7 |
| *DIAPH3--MALAT1* | 7.7 |
| *DIAPH1--IGH@* | 7.7 |
| *DIABLO--GRHL2* | 7.7 |
| *DHX57--GPN3* | 7.7 |
| *DHX40--EIF4A2* | 7.7 |
| *DHX35--NDRG1* | 7.7 |
| *DHX30--VWF* | 7.7 |
| *DHX15--DAPK3* | 7.7 |

| | |
|---|---|
| *DHRS13--STIL* | 7.7 |
| *DGCR8--MCM7* | 7.7 |
| *DESI1--ZNF30* | 7.7 |
| *DEPDC5--FTL* | 7.7 |
| *DEPDC1B--ELOVL7* | 7.7 |
| *DEPDC1B--AL157935.2* | 7.7 |
| *DEPDC1--CRTC2* | 7.7 |
| *DENR--MALAT1* | 7.7 |
| *DENR--AL450263.2* | 7.7 |
| *DENND5B--COL5A1* | 7.7 |
| *DENND5A--CBLB* | 7.7 |
| *DENND3--AC124312.3* | 7.7 |
| *DELE1--SASH1* | 7.7 |
| *DEK--SMG1* | 7.7 |
| *DEK--MALAT1* | 7.7 |
| *DEK--GLUL* | 7.7 |
| *DDX60L--IGK@* | 7.7 |
| *DDX6--SSBP1* | 7.7 |
| *DDX6--PASK* | 7.7 |
| *DDX54--UBXN7* | 7.7 |
| *DDX5--RBM14* | 7.7 |
| *DDX5--DARS* | 7.7 |
| *DDX5--COL6A3* | 7.7 |
| *DDX3Y--DDX3P1* | 7.7 |
| *DDX3X--MALAT1* | 7.7 |
| *DDX3X--AC023509.1* | 7.7 |
| *DDX39B--ARHGAP12* | 7.7 |
| *DDX27--TRIO* | 7.7 |
| *DDX23--E4F1* | 7.7 |
| *DDX21--MALAT1* | 7.7 |
| *DDX19A--SNRNP200* | 7.7 |
| *DDX17--RALGAPA2* | 7.7 |
| *DDX17--MALAT1* | 7.7 |
| *DDX17--HBB* | 7.7 |
| *DDX11--PRDM15* | 7.7 |
| *DDR1--YLPM1* | 7.7 |
| *DDR1--RIMS2* | 7.7 |
| *DDOST--KRT8* | 7.7 |
| *DDHD2--NEB* | 7.7 |
| *DDHD1--MAPKAPK5* | 7.7 |
| *DDB1--XIST* | 7.7 |
| *DDB1--CYP17A1* | 7.7 |
| *DCUN1D1--MUC4* | 7.7 |
| *DCTPP1--C21ORF91* | 7.7 |
| *DCTN1--DPY19L4* | 7.7 |
| *DCP1A--RPLP1* | 7.7 |
| *DCLK2--MALAT1* | 7.7 |
| *DCHS2--ZSWIM2* | 7.7 |
| *DCHS1--KLHL24* | 7.7 |
| *DCDC1--PTCH2* | 7.7 |
| *DCDC1--ADAMTS1* | 7.7 |
| *DCBLD2--SYNCRIP* | 7.7 |
| *DCBLD2--CGNL1* | 7.7 |
| *DCAF6--SEC61A1* | 7.7 |
| *DCAF16--EIF4G1* | 7.7 |
| *DCAF13--PTMA* | 7.7 |
| *DBT--ARAP1* | 7.7 |
| *DBI--FHAD1* | 7.7 |
| *DAPK1--PIK3R3* | 7.7 |
| *DAPK1--AKAP9* | 7.7 |
| *DAP3--MON2* | 7.7 |
| *DAP--SELENOF* | 7.7 |
| *DAG1--EIF3E* | 7.7 |
| *DAG1--CCNC* | 7.7 |
| *DAAM1--CLCN7* | 7.7 |
| *CYP17A1--MALAT1* | 7.7 |
| *CYLD--SERPINA1* | 7.7 |
| *CYFIP1--MALAT1* | 7.7 |
| *CYB5A--SEPTIN9* | 7.7 |
| *CXCL8--NAMPT* | 7.7 |
| *CXADR--DHCR24* | 7.7 |
| *CUX1--AKR1C3* | 7.7 |
| *CTSK--SUV39H2* | 7.7 |
| *CTPS1--MALAT1* | 7.7 |
| *CTNND1--ZNF577* | 7.7 |
| *CTNNB1--ARF5* | 7.7 |
| *CTH--ASPM* | 7.7 |
| *CTDSPL2--DDX6* | 7.7 |
| *CTCF--GRHL2* | 7.7 |
| *CTCF--ATF4* | 7.7 |
| *CSTB--VIM* | 7.7 |
| *CSRP1--XIST* | 7.7 |
| *CSNK1G3--VMP1* | 7.7 |
| *CSNK1G1--DLAT* | 7.7 |
| *CSNK1E--EFHC1* | 7.7 |
| *CSNK1D--RAD21* | 7.7 |
| *CSE1L--SPOCD1* | 7.7 |
| *CSE1L--SMG7* | 7.7 |
| *CSE1L--PTPN1* | 7.7 |
| *CSE1L--MALAT1* | 7.7 |
| *CSDE1--RAN* | 7.7 |
| *CSDE1--NEB* | 7.7 |
| *CSDE1--LGALS3BP* | 7.7 |
| *CSDE1--ATP6V1A* | 7.7 |
| *CS--AGO1* | 7.7 |
| *CROCC--NAPG* | 7.7 |
| *CRIPAK--TNRC18* | 7.7 |
| *CREBBP--PAXIP1-AS2* | 7.7 |
| *CREBBP--LASP1* | 7.7 |
| *CREBBP--FOXP2* | 7.7 |
| *CREBBP--COL1A2* | 7.7 |
| *CREB3L2--HSDL2* | 7.7 |
| *CPSF6--ESPN* | 7.7 |
| *CPSF6--DIP2A* | 7.7 |
| *CPOX--COL1A1* | 7.7 |
| *CPLX2--TMEM50A* | 7.7 |
| *CPLX2--KMT2E* | 7.7 |
| *CPLANE1--MALAT1* | 7.7 |
| *CPED1--RAB1B* | 7.7 |
| *CPD--EFR3A* | 7.7 |
| *COX6B1--TMTC2* | 7.7 |
| *COX6B1--SRRM2* | 7.7 |
| *COX6A1--RPL7L1* | 7.7 |
| *COX6A1--BMP2K* | 7.7 |
| *COX20--BIRC6* | 7.7 |
| *COX11--ASPN* | 7.7 |
| *COX10--PABPC1* | 7.7 |
| *COTL1--ZBTB21* | 7.7 |
| *COQ8A--DVL3* | 7.7 |
| *COPS3--EEF2* | 7.7 |
| *COPA--VPS37D* | 7.7 |
| *COPA--SULF2* | 7.7 |
| *COPA--HSP90B1* | 7.7 |
| *COMMD7--CCDC50* | 7.7 |
| *COLGALT1--AL031777.3* | 7.7 |
| *COLEC12--MALAT1* | 7.7 |
| *COL6A3--MORF4L1* | 7.7 |
| *COL6A3--MAP1B* | 7.7 |
| *COL6A3--LINC02012* | 7.7 |
| *COL4A5--FN1* | 7.7 |
| *COL4A2--MALAT1* | 7.7 |
| *COL3A1--WDR74* | 7.7 |
| *COL3A1--TAX1BP1* | 7.7 |
| *COL3A1--RPLP1* | 7.7 |

| | |
|---|---|
| *COL3A1--EGFR* | 7.7 |
| *COL3A1--BCL2* | 7.7 |
| *COL3A1--AC093010.3* | 7.7 |
| *COL1A2--TMBIM6* | 7.7 |
| *COL1A2--RPL36A-HNRNPH2* | 7.7 |
| *COL1A2--PSMA3-AS1* | 7.7 |
| *COL1A2--NISCH* | 7.7 |
| *COL1A2--MTRNR2L12* | 7.7 |
| *COL1A2--MALAT1* | 7.7 |
| *COL1A2--HNRNPH1* | 7.7 |
| *COL1A2--CABIN1* | 7.7 |
| *COL1A1--WDR70* | 7.7 |
| *COL1A1--STT3B* | 7.7 |
| *COL1A1--SHOC2* | 7.7 |
| *COL1A1--RPL15* | 7.7 |
| *COL1A1--LRPPRC* | 7.7 |
| *COL1A1--KLHL42* | 7.7 |
| *COL1A1--HSD17B7* | 7.7 |
| *COL1A1--H19* | 7.7 |
| *COL1A1--EXOSC7* | 7.7 |
| *COL1A1--ENO2* | 7.7 |
| *COL1A1--DDX27* | 7.7 |
| *COL1A1--ASCC3* | 7.7 |
| *COL12A1--TPT1* | 7.7 |
| *COG3--CALR* | 7.7 |
| *COASY--UBR4* | 7.7 |
| *COASY--TAF15* | 7.7 |
| *COA1--KMT2C* | 7.7 |
| *CNPY4--GOLGA8J* | 7.7 |
| *CNPY2--MYSM1* | 7.7 |
| *CNOT6--MAPKAPK2* | 7.7 |
| *CNOT6--LRPAP1* | 7.7 |
| *CNOT1--UBC* | 7.7 |
| *CNOT1--SUGP2* | 7.7 |
| *CNOT1--SMG1* | 7.7 |
| *CNOT1--PHKA2* | 7.7 |
| *CNOT1--KNL1* | 7.7 |
| *CNNM4--MUC4* | 7.7 |
| *CNN3--SON* | 7.7 |
| *CNKSR3--DUSP6* | 7.7 |
| *CNIH4--HOXB3* | 7.7 |
| *CNBP--ACTB* | 7.7 |
| *CMTM6--RBAK* | 7.7 |
| *CMTM4--KMT2D* | 7.7 |

| | |
|---|---|
| *CLYBL--IGH@* | 7.7 |
| *CLU--SLIT2* | 7.7 |
| *CLTC--VPS13A* | 7.7 |
| *CLTC--TMEM94* | 7.7 |
| *CLTC--RNF7* | 7.7 |
| *CLTC--PLEC* | 7.7 |
| *CLTC--LMTK2* | 7.7 |
| *CLSPN--ANKRD7* | 7.7 |
| *CLPTM1--MALAT1* | 7.7 |
| *CLPTM1--DUS4L* | 7.7 |
| *CLMP--COL5A2* | 7.7 |
| *CLIP1--CD63* | 7.7 |
| *CLIC4--FBN1* | 7.7 |
| *CLIC4--C1GALT1* | 7.7 |
| *CLDN4--SERPINA1* | 7.7 |
| *CLASP2--B2M* | 7.7 |
| *CKAP5--TPT1* | 7.7 |
| *CKAP5--DST* | 7.7 |
| *CKAP2--MAN2B2* | 7.7 |
| *CIT--BICDL1* | 7.7 |
| *CIITA--SDHA* | 7.7 |
| *CIC--RTL8C* | 7.7 |
| *CIAO1--VASN* | 7.7 |
| *CIAO1--PRPF18* | 7.7 |
| *CHSY1--MALAT1* | 7.7 |
| *CHST11--SEMA6C* | 7.7 |
| *CHST11--MALAT1* | 7.7 |
| *CHST11--HMCN1* | 7.7 |
| *CHST11--FN1* | 7.7 |
| *CHST11--COL6A1* | 7.7 |
| *CHMP5--AC114402.2* | 7.7 |
| *CHMP1A--UBB* | 7.7 |
| *CHML--DTX3L* | 7.7 |
| *CHM--TTLL7* | 7.7 |
| *CHID1--MALAT1* | 7.7 |
| *CHGB--GTF2I* | 7.7 |
| *CHGB--CFAP221* | 7.7 |
| *CHGA--MALAT1* | 7.7 |
| *CHFR--MALAT1* | 7.7 |
| *CHFR--AC011448.1* | 7.7 |
| *CHD9--PSAP* | 7.7 |
| *CHD9--LRRC49* | 7.7 |
| *CHD9--GGA1* | 7.7 |
| *CHD6--COL27A1* | 7.7 |
| *CHD3--MALAT1* | 7.7 |

| | |
|---|---|
| *CHD2--PPP4R3B* | 7.7 |
| *CHD1--CDC42* | 7.7 |
| *CHD1--AC000123.3* | 7.7 |
| *CHAF1B--GNAS* | 7.7 |
| *CFLAR--AC068896.1* | 7.7 |
| *CFL1--ARHGAP1* | 7.7 |
| *CFDP1--ZCCHC8* | 7.7 |
| *CFAP70--AL451062.3* | 7.7 |
| *CFAP20--ABCC3* | 7.7 |
| *CERT1--ABCC2* | 7.7 |
| *CEP97--TNFAIP3* | 7.7 |
| *CEP85L--XRCC5* | 7.7 |
| *CEP76--LAPTM4A* | 7.7 |
| *CEP70--CCT8* | 7.7 |
| *CEP350--TLN1* | 7.7 |
| *CEP350--KHSRP* | 7.7 |
| *CEP350--IGH@* | 7.7 |
| *CEP295--TFRC* | 7.7 |
| *CEP290--MALAT1* | 7.7 |
| *CEP250--PAXBP1* | 7.7 |
| *CEP250--MALAT1* | 7.7 |
| *CEP250--CCNT2* | 7.7 |
| *CEP192--DIS3* | 7.7 |
| *CEP131--TACC1* | 7.7 |
| *CEP128--CSNK1D* | 7.7 |
| *CEP120--KDM5C* | 7.7 |
| *CEP104--HERC2* | 7.7 |
| *CENPU--AL355987.2* | 7.7 |
| *CENPO--CPNE1* | 7.7 |
| *CENPJ--ZNF217* | 7.7 |
| *CENPF--PGD* | 7.7 |
| *CENPF--NEO1* | 7.7 |
| *CENPF--CNPY4* | 7.7 |
| *CENPF--AC117386.2* | 7.7 |
| *CEMIP2--RPL27A* | 7.7 |
| *CELSR2--COL1A2* | 7.7 |
| *CDYL--AC091230.1* | 7.7 |
| *CDV3--CARMIL1* | 7.7 |
| *CDR2L--SMAD1* | 7.7 |
| *CDON--AP004607.3* | 7.7 |
| *CDKN2B-AS1--POLR3A* | 7.7 |
| *CDKL5--RGL2* | 7.7 |
| *CDK6--SOS1* | 7.7 |
| *CDK6--KMT2A* | 7.7 |
| *CDK6--AC079594.2* | 7.7 |

| | |
|---|---|
| *CDK5RAP2--LUZP1* | 7.7 |
| *CDK5RAP2--HNRNPA3* | 7.7 |
| *CDK2--EEF1A1* | 7.7 |
| *CDK13--MALAT1* | 7.7 |
| *CDH2--FTL* | 7.7 |
| *CDH1--NDUFV1* | 7.7 |
| *CDCP1--ZDHHC7* | 7.7 |
| *CDC42BPG--NACA* | 7.7 |
| *CDC42BPB--LAS1L* | 7.7 |
| *CDC42BPA--RNF213* | 7.7 |
| *CDC42BPA--BMP2K* | 7.7 |
| *CDC27--XIST* | 7.7 |
| *CDC27--PNRC2* | 7.7 |
| *CDC23--MALAT1* | 7.7 |
| *CD81--USP34* | 7.7 |
| *CD81--FTL* | 7.7 |
| *CD55--NAMPT* | 7.7 |
| *CD55--HUWE1* | 7.7 |
| *CD46--OTUD7B* | 7.7 |
| *CD46--HERC2* | 7.7 |
| *CD46--ATP11B* | 7.7 |
| *CD44--RERE* | 7.7 |
| *CD2AP--USP53* | 7.7 |
| *CD2AP--COPG1* | 7.7 |
| *CD24--RHCE* | 7.7 |
| *CD209--MALAT1* | 7.7 |
| *CD164--GDA* | 7.7 |
| *CCT8--SPOCK3* | 7.7 |
| *CCT8--SFPQ* | 7.7 |
| *CCT6A--NCOR1* | 7.7 |
| *CCT5--ZFAND6* | 7.7 |
| *CCT5--KDM4A* | 7.7 |
| *CCT5--BRIX1* | 7.7 |
| *CCT4--SQSTM1* | 7.7 |
| *CCT2--TPT1* | 7.7 |
| *CCSER2--KIDINS220* | 7.7 |
| *CCP110--HMGN2* | 7.7 |
| *CCNT1--MALAT1* | 7.7 |
| *CCNI--SLC25A6* | 7.7 |
| *CCNI--ACTB* | 7.7 |
| *CCNG2--KTN1* | 7.7 |
| *CCNE1--AC027097.2* | 7.7 |
| *CCND3--MALAT1* | 7.7 |
| *CCND3--IGH@* | 7.7 |
| *CCNB1IP1--CAPZB* | 7.7 |
| *CCNB1--NPIPB11* | 7.7 |
| *CCN1--TTN* | 7.7 |
| *CCM2--MALAT1* | 7.7 |
| *CCDC88A--MYO9A* | 7.7 |
| *CCDC88A--ATRX* | 7.7 |
| *CCDC82--RO60* | 7.7 |
| *CCDC186--PHKA2* | 7.7 |
| *CCDC186--MAP4K1* | 7.7 |
| *CCDC174--ARL2-SNX15* | 7.7 |
| *CCDC162P--MUC4* | 7.7 |
| *CCDC15--CUL5* | 7.7 |
| *CCDC14--ZNF148* | 7.7 |
| *CCDC14--DISC1* | 7.7 |
| *CCDC138--CHD7* | 7.7 |
| *CCDC122--MALAT1* | 7.7 |
| *CCDC112--RC3H2* | 7.7 |
| *CCAR2--IFNGR2* | 7.7 |
| *CCAR1--TASOR2* | 7.7 |
| *CCAR1--DMPK* | 7.7 |
| *CBLL1--CASC3* | 7.7 |
| *CBFB--PLEC* | 7.7 |
| *CAST--TCF12* | 7.7 |
| *CAST--PRELID3B* | 7.7 |
| *CASD1--LAMA4* | 7.7 |
| *CARMIL1--SLC17A4* | 7.7 |
| *CARMIL1--PEG10* | 7.7 |
| *CARM1--STK38* | 7.7 |
| *CAPRIN1--PRKD1* | 7.7 |
| *CAPRIN1--ABCC3* | 7.7 |
| *CAPN9--NOTCH2* | 7.7 |
| *CAPN5--FASN* | 7.7 |
| *CANX--TSN* | 7.7 |
| *CAND1--ZNF107* | 7.7 |
| *CAND1--MALAT1* | 7.7 |
| *CAND1--KIF4A* | 7.7 |
| *CALR--STRC* | 7.7 |
| *CALR--PRRC2C* | 7.7 |
| *CALR--PLXNB1* | 7.7 |
| *CALR--PEPD* | 7.7 |
| *CALR--MALAT1* | 7.7 |
| *CALR--CUL4B* | 7.7 |
| *CALM1--ARSG* | 7.7 |
| *CALD1--MGAT4B* | 7.7 |
| *CALCOCO2--VPS28* | 7.7 |
| *CALCA--UTRN* | 7.7 |
| *CALCA--MALAT1* | 7.7 |
| *CALCA--ATRX* | 7.7 |
| *CADPS2--MALAT1* | 7.7 |
| *CAD--GDI1* | 7.7 |
| *CACUL1--METTL9* | 7.7 |
| *CACNA2D1--WDR74* | 7.7 |
| *CACNA2D1--FTL* | 7.7 |
| *CACNA1H--RNF169* | 7.7 |
| *CACNA1D--RPPH1* | 7.7 |
| *C9ORF78--CLTC* | 7.7 |
| *C8ORF33--RREB1* | 7.7 |
| *C6ORF62--RPL30* | 7.7 |
| *C6ORF62--IFI27L2* | 7.7 |
| *C5ORF63--RNF34* | 7.7 |
| *C5--L3MBTL3* | 7.7 |
| *C2CD4B--CEP85* | 7.7 |
| *C2CD3--XIST* | 7.7 |
| *C2CD3--STAT1* | 7.7 |
| *C21ORF58--RACGAP1* | 7.7 |
| *C1RL--REST* | 7.7 |
| *C1QA--STIP1* | 7.7 |
| *C1ORF43--DNAJC2* | 7.7 |
| *C19MC--IGK@* | 7.7 |
| *C18ORF25--CDC5L* | 7.7 |
| *C17ORF80--STOM* | 7.7 |
| *C17ORF80--COL4A2* | 7.7 |
| *C16ORF72--AFG3L2* | 7.7 |
| *C15ORF40--GAPDH* | 7.7 |
| *C12ORF45--MALAT1* | 7.7 |
| *BYSL--MALAT1* | 7.7 |
| *BX322639.1--ZNF99* | 7.7 |
| *BUB3--PPP1R21* | 7.7 |
| *BTF3--SENP7* | 7.7 |
| *BTF3--AL359762.3* | 7.7 |
| *BTBD3--UBC* | 7.7 |
| *BTBD10--COL1A2* | 7.7 |
| *BSN--MALAT1* | 7.7 |
| *BRWD1--SNX9* | 7.7 |
| *BRWD1--OGFRL1* | 7.7 |
| *BRWD1--FAM204A* | 7.7 |
| *BRPF3--KDM2A* | 7.7 |
| *BROX--ADGRD1* | 7.7 |
| *BRI3BP--MALAT1* | 7.7 |
| *BRD8--ILVBL* | 7.7 |
| *BRD7--TSEN54* | 7.7 |

| | |
|---|---|
| *BRD7--SCN4B* | 7.7 |
| *BRD2--RRP1B* | 7.7 |
| *BRD2--RNF10* | 7.7 |
| *BRD2--AHNAK* | 7.7 |
| *BRAF--EFS* | 7.7 |
| *BPTF--XIST* | 7.7 |
| *BPTF--SPP1* | 7.7 |
| *BPTF--MALAT1* | 7.7 |
| *BPTF--IGH@* | 7.7 |
| *BOP1--FOXH1* | 7.7 |
| *BOD1L1--RABIF* | 7.7 |
| *BOD1L1--GJA1* | 7.7 |
| *BNIP3L--GRB2* | 7.7 |
| *BMS1--LRTOMT* | 7.7 |
| *BMPR1A--DSP* | 7.7 |
| *BMP7--AL139300.1* | 7.7 |
| *BLOC1S5-TXNDC5--TUG1* | 7.7 |
| *BIRC6--ZNF706* | 7.7 |
| *BIRC6--RBM39* | 7.7 |
| *BIRC6--CAPN15* | 7.7 |
| *BIRC3--BTAF1* | 7.7 |
| *BICRAL--PLCB1* | 7.7 |
| *BICD1--AHNAK* | 7.7 |
| *BDP1--ZFYVE16* | 7.7 |
| *BDP1--TSPAN13* | 7.7 |
| *BCR--DYNC1H1* | 7.7 |
| *BCR--AL078602.1* | 7.7 |
| *BCLAF1--RPS7* | 7.7 |
| *BCL2L2--COL1A2* | 7.7 |
| *BCL2--KDM6B* | 7.7 |
| *BCL11A--ZNF839* | 7.7 |
| *BCKDHB--HMG20B* | 7.7 |
| *BCKDHB--GLS* | 7.7 |
| *BCAR3--EIF3B* | 7.7 |
| *BBX--SLX4* | 7.7 |
| *BBS2--MALAT1* | 7.7 |
| *BAZ2B--ZZEF1* | 7.7 |
| *BAZ2B--MALAT1* | 7.7 |
| *BAZ1B--MALAT1* | 7.7 |
| *BANK1--COL3A1* | 7.7 |
| *BANF1--BRAP* | 7.7 |
| *BAGE2--CU104787.1* | 7.7 |
| *BAG6--TLK1* | 7.7 |
| *BACH2--SERPING1* | 7.7 |

| | |
|---|---|
| *BABAM2--ZNF384* | 7.7 |
| *B4GALT5--PARN* | 7.7 |
| *B4GALT5--MALAT1* | 7.7 |
| *B4GALT4--HECTD1* | 7.7 |
| *B4GALT3--SAFB* | 7.7 |
| *B4GALT1--C21ORF58* | 7.7 |
| *B3GNTL1--HEATR5B* | 7.7 |
| *B2M--MGAT4B* | 7.7 |
| *B2M--INTS8* | 7.7 |
| *B2M--AREL1* | 7.7 |
| *AZGP1--TUBB4B* | 7.7 |
| *AVL9--ABCC10* | 7.7 |
| *AURKA--ADCY3* | 7.7 |
| *AUP1--SFN* | 7.7 |
| *ATXN2L--RAB5C* | 7.7 |
| *ATXN2L--CSDE1* | 7.7 |
| *ATXN2--SSR1* | 7.7 |
| *ATXN10--NFE2L1* | 7.7 |
| *ATXN1--ADCY6* | 7.7 |
| *ATRX--SMG1* | 7.7 |
| *ATRX--NPAT* | 7.7 |
| *ATRX--MYL6* | 7.7 |
| *ATRX--MAGT1* | 7.7 |
| *ATRNL1--TNRC6B* | 7.7 |
| *ATRNL1--MALAT1* | 7.7 |
| *ATRNL1--IPO9* | 7.7 |
| *ATRN--HERC2* | 7.7 |
| *ATR--MALAT1* | 7.7 |
| *ATP9A--TTC37* | 7.7 |
| *ATP9A--KCNQ1OT1* | 7.7 |
| *ATP9A--CSTF1* | 7.7 |
| *ATP6V1H--RP1L1* | 7.7 |
| *ATP6V1H--PPIG* | 7.7 |
| *ATP6V1G2-DDX39B--MALAT1* | 7.7 |
| *ATP6V1E1--IPO9* | 7.7 |
| *ATP6V1A--NDUFAF5* | 7.7 |
| *ATP6V0E2--MFSD4B* | 7.7 |
| *ATP6V0A1--RPS6* | 7.7 |
| *ATP6V0A1--MDM4* | 7.7 |
| *ATP5PO--LARS* | 7.7 |
| *ATP5ME--FTH1* | 7.7 |
| *ATP5ME--CKAP2L* | 7.7 |
| *ATP5IF1--CHMP4B* | 7.7 |
| *ATP5F1E--KPNB1* | 7.7 |

| | |
|---|---|
| *ATP5F1C--MALAT1* | 7.7 |
| *ATP5F1B--ROCK1* | 7.7 |
| *ATP2C1--AFDN* | 7.7 |
| *ATP2B2--FNBP4* | 7.7 |
| *ATP2B1--MALAT1* | 7.7 |
| *ATP2B1--AC117386.2* | 7.7 |
| *ATP2A2--MALAT1* | 7.7 |
| *ATP2A2--LRPPRC* | 7.7 |
| *ATP2A2--DA750114* | 7.7 |
| *ATP1B1--DYNC1H1* | 7.7 |
| *ATP1B1--CAPNS1* | 7.7 |
| *ATP1A1--YAF2* | 7.7 |
| *ATP1A1--NOTCH2* | 7.7 |
| *ATP1A1--KIDINS220* | 7.7 |
| *ATP13A3--TAX1BP1* | 7.7 |
| *ATP13A3--MALAT1* | 7.7 |
| *ATP13A3--HSP90AB1* | 7.7 |
| *ATP11C--HSPG2* | 7.7 |
| *ATN1--CKS1B* | 7.7 |
| *ATM--EIF3J* | 7.7 |
| *ATM--ACOT9* | 7.7 |
| *ATM--AC120057.2* | 7.7 |
| *ATIC--MALAT1* | 7.7 |
| *ATG16L1--MICU2* | 7.7 |
| *ATF4--ZNF664* | 7.7 |
| *ATF2--PTK2* | 7.7 |
| *ATAD5--MALAT1* | 7.7 |
| *ATAD5--FN1* | 7.7 |
| *ATAD2B--CUL4B* | 7.7 |
| *ATAD2--LDHA* | 7.7 |
| *ATAD2--CLPTM1* | 7.7 |
| *ASXL2--FBLN7* | 7.7 |
| *ASRGL1--PPP2R2C* | 7.7 |
| *ASRGL1--ARL13B* | 7.7 |
| *ASPM--RAB10* | 7.7 |
| *ASPM--MALAT1* | 7.7 |
| *ASPM--FASN* | 7.7 |
| *ASH2L--PTGFRN* | 7.7 |
| *ASH2L--PRKD3* | 7.7 |
| *ASH1L--MALAT1* | 7.7 |
| *ASCL1--AK9* | 7.7 |
| *ASCC3--MALAT1* | 7.7 |
| *ARPC5--SPP1* | 7.7 |
| *ARPC3--ZC3HAV1* | 7.7 |
| *ARPC2--CCAR1* | 7.7 |

| | |
|---|---|
| *ARNT--SYNE1* | 7.7 |
| *ARNT--C2CD2L* | 7.7 |
| *ARMCX6--N4BP2L2* | 7.7 |
| *ARMC8--PAFAH1B2* | 7.7 |
| *ARL8B--ITPR1* | 7.7 |
| *ARL6IP5--DTX3L* | 7.7 |
| *ARL1--MALAT1* | 7.7 |
| *ARIH2--TBX21* | 7.7 |
| *ARIH2--SYPL1* | 7.7 |
| *ARIH2--STT3A* | 7.7 |
| *ARID5B--PHF6* | 7.7 |
| *ARID4B--SLK* | 7.7 |
| *ARID4B--CECR2* | 7.7 |
| *ARID2--SETX* | 7.7 |
| *ARID2--SCAF11* | 7.7 |
| *ARID2--SBF2* | 7.7 |
| *ARID1B--APOB* | 7.7 |
| *ARID1A--KRBOX4* | 7.7 |
| *ARHGEF40--MALAT1* | 7.7 |
| *ARHGEF2--TMEM266* | 7.7 |
| *ARHGEF2--MALAT1* | 7.7 |
| *ARHGEF17--HECTD4* | 7.7 |
| *ARHGEF12--RNF103* | 7.7 |
| *ARHGEF12--ITIH3* | 7.7 |
| *ARHGEF11--FAT1* | 7.7 |
| *ARHGEF10L--UBR4* | 7.7 |
| *ARHGEF10L--FN1* | 7.7 |
| *ARHGDIA--SPG7* | 7.7 |
| *ARHGAP5--MALAT1* | 7.7 |
| *ARHGAP35--NRF1* | 7.7 |
| *ARHGAP31--TP53BP1* | 7.7 |
| *ARHGAP31--MALAT1* | 7.7 |
| *ARHGAP29--CDK6* | 7.7 |
| *ARHGAP27--HECTD4* | 7.7 |
| *ARHGAP26--MUC4* | 7.7 |
| *ARHGAP26--CCNA2* | 7.7 |
| *ARHGAP21--VIM* | 7.7 |
| *ARHGAP11A--PLEC* | 7.7 |
| *ARGLU1--POLR2A* | 7.7 |
| *ARFGEF3--MALAT1* | 7.7 |
| *ARFGEF3--FAM102A* | 7.7 |
| *ARFGEF2--CLTC* | 7.7 |
| *ARF3--CALR* | 7.7 |
| *AREL1--XRN2* | 7.7 |
| *AQP3--SUN1* | 7.7 |

| | |
|---|---|
| *APRT--MFSD10* | 7.7 |
| *APPL1--ECI1* | 7.7 |
| *APOOL--MALAT1* | 7.7 |
| *APOM--SUPT20H* | 7.7 |
| *APOM--SMARCC2* | 7.7 |
| *APOL2--KMT2C* | 7.7 |
| *APOE--MACF1* | 7.7 |
| *APOBEC3C--NUP133* | 7.7 |
| *APOB--RPPH1* | 7.7 |
| *APOB--OBSCN* | 7.7 |
| *APOB--MALAT1* | 7.7 |
| *APOB--LRP10* | 7.7 |
| *APOA2--SETD2* | 7.7 |
| *APLP2--SP100* | 7.7 |
| *APLP2--CBX5* | 7.7 |
| *APLP1--MALAT1* | 7.7 |
| *API5--NEAT1* | 7.7 |
| *API5--MALAT1* | 7.7 |
| *APCDD1--VAPA* | 7.7 |
| *APCDD1--DLG5* | 7.7 |
| *APCDD1--ASH1L* | 7.7 |
| *APC--RALGAPA1* | 7.7 |
| *APBB2--DNAJC13* | 7.7 |
| *AP3S1--LVRN* | 7.7 |
| *AP004607.3--CDON* | 7.7 |
| *AP001273.2--TSSK4* | 7.7 |
| *AP001273.2--SERINC2* | 7.7 |
| *AP001267.5--MALAT1* | 7.7 |
| *AP000781.2--CLIP1* | 7.7 |
| *AP000646.1--P4HA1* | 7.7 |
| *AP000350.2--KLHL5* | 7.7 |
| *ANXA5--CALM2* | 7.7 |
| *ANXA5--AL445222.1* | 7.7 |
| *ANXA3--NUP107* | 7.7 |
| *ANXA2--ACTG1* | 7.7 |
| *ANP32B--TRA@* | 7.7 |
| *ANP32B--MALAT1* | 7.7 |
| *ANP32A--MTHFD1* | 7.7 |
| *ANP32A--AC093525.7* | 7.7 |
| *ANKS1B--PPT1* | 7.7 |
| *ANKRD50--ZBED5* | 7.7 |
| *ANKRD50--B2M* | 7.7 |
| *ANKRD46--WDR74* | 7.7 |
| *ANKRD36--C11ORF24* | 7.7 |
| *ANKRD28--PTPN13* | 7.7 |

| | |
|---|---|
| *ANKRD26--SPATA13* | 7.7 |
| *ANKRD26--RMRP* | 7.7 |
| *ANKRD17--ARHGAP10* | 7.7 |
| *ANKRD12--DNAJC5* | 7.7 |
| *ANKRD11--DCUN1D1* | 7.7 |
| *ANKRD11--CEP192* | 7.7 |
| *ANKRD10--DKC1* | 7.7 |
| *ANKMY1--PRPF8* | 7.7 |
| *ANKIB1--PAM16* | 7.7 |
| *ANKHD1-EIF4EBP3--SYNE2* | 7.7 |
| *ANKHD1-EIF4EBP3--PCNX1* | 7.7 |
| *ANKHD1-EIF4EBP3--FADS1* | 7.7 |
| *ANKHD1--TRA@* | 7.7 |
| *ANKFY1--SEPTIN9* | 7.7 |
| *ANKAR--PMS1* | 7.7 |
| *ANK2--COL3A1* | 7.7 |
| *ANK2--CALD1* | 7.7 |
| *ANK1--BAG5* | 7.7 |
| *ANAPC5--NCEH1* | 7.7 |
| *ANAPC16--AC018521.1* | 7.7 |
| *ANAPC15--OSCP1* | 7.7 |
| *AMZ2--NRIP1* | 7.7 |
| *AMY2B--UBB* | 7.7 |
| *AMBP--ENOPH1* | 7.7 |
| *ALPK3--ZBTB44* | 7.7 |
| *ALMS1--ENOSF1* | 7.7 |
| *ALK--MALAT1* | 7.7 |
| *ALG5--ILF3* | 7.7 |
| *ALG2--MALAT1* | 7.7 |
| *ALDOA--ZBTB20* | 7.7 |
| *ALDH3A2--AMD1* | 7.7 |
| *ALAS1--ZNF638* | 7.7 |
| *AL845552.1--CPED1* | 7.7 |
| *AL671762.1--KTN1* | 7.7 |
| *AL596202.1--ATP2A2* | 7.7 |
| *AL590004.3--RNF123* | 7.7 |
| *AL512637.1--PRIM1* | 7.7 |
| *AL445487.1--KPNA2* | 7.7 |
| *AL445305.1--AP3S1* | 7.7 |
| *AL390728.4--OAS3* | 7.7 |
| *AL365475.1--PUM3* | 7.7 |
| *AL365181.3--CTNNB1* | 7.7 |
| *AL358334.3--PAX5* | 7.7 |
| *AL357075.5--MALAT1* | 7.7 |

| | |
|---|---|
| *AL355987.3--PPIL2* | 7.7 |
| *AL355377.1--SMC5* | 7.7 |
| *AL355297.4--FTL* | 7.7 |
| *AL355075.4--ACTB* | 7.7 |
| *AL354809.1--RP1L1* | 7.7 |
| *AL160286.1--MALAT1* | 7.7 |
| *AL160237.3--PCLO* | 7.7 |
| *AL157392.5--AC026464.4* | 7.7 |
| *AL139300.1--VPS37B* | 7.7 |
| *AL137782.1--VIRMA* | 7.7 |
| *AL133500.1--ITGB1* | 7.7 |
| *AL133353.2--ZNF568* | 7.7 |
| *AL121900.2--GJB2* | 7.7 |
| *AL109827.1--ICAM1* | 7.7 |
| *AL109811.3--RAPGEF2* | 7.7 |
| *AL109811.3--DRAP1* | 7.7 |
| *AL049776.1--TSIX* | 7.7 |
| *AL049697.1--COL3A1* | 7.7 |
| *AL035078.4--NDUFV3* | 7.7 |
| *AL031777.3--UPF1* | 7.7 |
| *AL031681.2--STIL* | 7.7 |
| *AL024498.2--TMEM14DP* | 7.7 |
| *AL022311.1--NEAT1* | 7.7 |
| *AL022238.4--DYRK4* | 7.7 |
| *AL021408.1--CCDC50* | 7.7 |
| *AL021155.5--PHIP* | 7.7 |
| *AL021155.5--MAN2B1* | 7.7 |
| *AL021155.5--MALAT1* | 7.7 |
| *AL021155.5--AL513165.2* | 7.7 |
| *AL020996.2--MKI67* | 7.7 |
| *AKT1--DICER1* | 7.7 |
| *AKR1C3--TSHZ1* | 7.7 |
| *AKR1C3--CEP192* | 7.7 |
| *AKR1C2--MALAT1* | 7.7 |
| *AKR1B10--PTGES3* | 7.7 |
| *AKR1A1--SUGP2* | 7.7 |
| *AKNA--SCARNA7* | 7.7 |
| *AKAP9--ZNF692* | 7.7 |
| *AKAP9--PARG* | 7.7 |
| *AKAP9--KPNB1* | 7.7 |
| *AKAP9--GTF2I* | 7.7 |
| *AKAP9--GPX2* | 7.7 |
| *AKAP9--EIF5B* | 7.7 |
| *AKAP9--CNOT4* | 7.7 |
| *AKAP9--CD74* | 7.7 |
| *AKAP8L--TMTC4* | 7.7 |
| *AKAP6--SRRM2* | 7.7 |
| *AKAP17A--DEPTOR* | 7.7 |
| *AKAP13--HSP90B1* | 7.7 |
| *AKAP11--MALAT1* | 7.7 |
| *AHSA2P--NUP98* | 7.7 |
| *AHSA1--ONECUT2* | 7.7 |
| *AHR--MALAT1* | 7.7 |
| *AHNAK--WDR74* | 7.7 |
| *AHNAK--UCKL1* | 7.7 |
| *AHNAK--TAB3* | 7.7 |
| *AHNAK--STON1* | 7.7 |
| *AHNAK--PBX2* | 7.7 |
| *AHNAK--MMAB* | 7.7 |
| *AHNAK--ASH1L* | 7.7 |
| *AHNAK--ARF3* | 7.7 |
| *AHCYL2--AL162417.1* | 7.7 |
| *AGTRAP--GLIPR2* | 7.7 |
| *AGTPBP1--MALAT1* | 7.7 |
| *AGRN--EFHD2* | 7.7 |
| *AGR2--CCNT2* | 7.7 |
| *AGPS--BCL2* | 7.7 |
| *AGPS--AOC1* | 7.7 |
| *AGO4--ANP32E* | 7.7 |
| *AGO2--SMG1* | 7.7 |
| *AGMO--CXCL5* | 7.7 |
| *AGGF1--TMEM176A* | 7.7 |
| *AGFG1--SLC4A1AP* | 7.7 |
| *AGAP1--RPS6KA5* | 7.7 |
| *AGA--NUCKS1* | 7.7 |
| *AFG3L2--TPR* | 7.7 |
| *AFG3L1P--POLR3A* | 7.7 |
| *AFF4--ACTB* | 7.7 |
| *AFDN--MALAT1* | 7.7 |
| *AEBP1--MALAT1* | 7.7 |
| *AEBP1--CASK* | 7.7 |
| *AEBP1--ARL6IP5* | 7.7 |
| *ADSL--SPTAN1* | 7.7 |
| *ADNP2--ZC3H7B* | 7.7 |
| *ADNP--MALAT1* | 7.7 |
| *ADNP--CACNA2D1* | 7.7 |
| *ADM--TMEM67* | 7.7 |
| *ADK--SLC16A9* | 7.7 |
| *ADK--JAK2* | 7.7 |
| *ADK--DOCK1* | 7.7 |
| *ADH5--NR1D1* | 7.7 |
| *ADGRV1--PFKL* | 7.7 |
| *ADGRV1--MALAT1* | 7.7 |
| *ADGRV1--GAB2* | 7.7 |
| *ADGRV1--B2M* | 7.7 |
| *ADGRL2--MALAT1* | 7.7 |
| *ADD2--ZZEF1* | 7.7 |
| *ADD1--CEMIP2* | 7.7 |
| *ADCY9--MMP25-AS1* | 7.7 |
| *ADARB1--SPATC1L* | 7.7 |
| *ADAR--MALAT1* | 7.7 |
| *ADAR--EPHB3* | 7.7 |
| *ADAR--ARIH2* | 7.7 |
| *ADAMTS9--TRIM72* | 7.7 |
| *ADAM9--MALAT1* | 7.7 |
| *ADAM17--SPG11* | 7.7 |
| *ADAM12--ENO1* | 7.7 |
| *AD000090.1--XIST* | 7.7 |
| *AD000090.1--UBC* | 7.7 |
| *AD000090.1--SRRM2* | 7.7 |
| *AD000090.1--SLTM* | 7.7 |
| *AD000090.1--SEC62* | 7.7 |
| *AD000090.1--NCL* | 7.7 |
| *AD000090.1--MUC4* | 7.7 |
| *AD000090.1--MED13* | 7.7 |
| *AD000090.1--HMCN1* | 7.7 |
| *AD000090.1--H19* | 7.7 |
| *AD000090.1--FTL* | 7.7 |
| *AD000090.1--FN1* | 7.7 |
| *AD000090.1--FLNA* | 7.7 |
| *AD000090.1--EEF1A1* | 7.7 |
| *AD000090.1--DYNC1H1* | 7.7 |
| *AD000090.1--DIP2B* | 7.7 |
| *AD000090.1--CSDE1* | 7.7 |
| *AD000090.1--COL1A2* | 7.7 |
| *AD000090.1--COL1A1* | 7.7 |
| *AD000090.1--AHNAK* | 7.7 |
| *ACVR1B--KMT2E* | 7.7 |
| *ACVR1--SCN9A* | 7.7 |
| *ACTN4--GGNBP2* | 7.7 |
| *ACTN4--DTX3* | 7.7 |
| *ACTG1--UNC13B* | 7.7 |
| *ACTG1--PSAP* | 7.7 |
| *ACTG1--PRKDC* | 7.7 |

| | |
|---|---|
| *ACTG1--NEAT1* | 7.7 |
| *ACTG1--HERC2* | 7.7 |
| *ACTG1--CHD7* | 7.7 |
| *ACTB--ZDHHC18* | 7.7 |
| *ACTB--ZBTB16* | 7.7 |
| *ACTB--WDR74* | 7.7 |
| *ACTB--RDX* | 7.7 |
| *ACTB--MLLT6* | 7.7 |
| *ACTB--KAT6A* | 7.7 |
| *ACTB--HACL1* | 7.7 |
| *ACTB--ERICH6B* | 7.7 |
| *ACTB--CRTC1* | 7.7 |
| *ACTB--AHR* | 7.7 |
| *ACTA2--USP34* | 7.7 |
| *ACTA2--DGCR2* | 7.7 |
| *ACSM3--TM9SF2* | 7.7 |
| *ACOX2--ANXA11* | 7.7 |
| *ACOT9--SLC9B2* | 7.7 |
| *ACOT8--ACTA2* | 7.7 |
| *ACAT2--IGFBP4* | 7.7 |
| *ACAT1--MALAT1* | 7.7 |
| *ACAT1--AFDN* | 7.7 |
| *ACAP2--RNF213* | 7.7 |
| *ACADVL--TPM2* | 7.7 |
| *ACADS--TAPBP* | 7.7 |
| *ACAD9--MALAT1* | 7.7 |
| *ACAD8--MALAT1* | 7.7 |
| *ACACB--MALAT1* | 7.7 |
| *ACACA--VPS13A* | 7.7 |
| *ACACA--RTN4* | 7.7 |
| *ACAA2--MRPS25* | 7.7 |
| *ACAA2--LRRC45* | 7.7 |
| *AC253572.2--MALAT1* | 7.7 |
| *AC245047.6--AL606490.1* | 7.7 |
| *AC245033.1--MACF1* | 7.7 |
| *AC244197.3--PKD1* | 7.7 |
| *AC239859.5--IGK@* | 7.7 |
| *AC138915.2--IGK@* | 7.7 |
| *AC138409.2--LBR* | 7.7 |
| *AC132217.2--VPS13A* | 7.7 |
| *AC128688.1--INAVA* | 7.7 |
| *AC124312.3--RBM39* | 7.7 |
| *AC124312.3--FTH1* | 7.7 |
| *AC118549.1--PRPF8* | 7.7 |

| | |
|---|---|
| *AC110275.1--BCL6* | 7.7 |
| *AC104758.3--GOLGA6L9* | 7.7 |
| *AC104619.3--MALAT1* | 7.7 |
| *AC104581.2--NEB* | 7.7 |
| *AC104046.1--FABP5* | 7.7 |
| *AC104041.1--CCDC162P* | 7.7 |
| *AC100839.2--FRY* | 7.7 |
| *AC098650.1--PPIA* | 7.7 |
| *AC098582.1--ANLN* | 7.7 |
| *AC098483.1--SYNE1* | 7.7 |
| *AC093512.2--MALAT1* | 7.7 |
| *AC092642.1--TXNL4A* | 7.7 |
| *AC092279.1--BAZ2B* | 7.7 |
| *AC091951.1--GOLGA6L2* | 7.7 |
| *AC091551.1--DPYSL2* | 7.7 |
| *AC090360.1--MLLT10* | 7.7 |
| *AC073912.3--NPIPB5* | 7.7 |
| *AC073610.2--ACTG1* | 7.7 |
| *AC073585.2--UBC* | 7.7 |
| *AC067968.1--NFAT5* | 7.7 |
| *AC048338.2--BDP1* | 7.7 |
| *AC036108.2--RPL18* | 7.7 |
| *AC034193.1--GYS1* | 7.7 |
| *AC027644.4--MRPS33* | 7.7 |
| *AC026979.4--NEFM* | 7.7 |
| *AC026362.1--PIGG* | 7.7 |
| *AC024451.3--UBE2N* | 7.7 |
| *AC023934.1--ZNF91* | 7.7 |
| *AC023509.1--RSBN1* | 7.7 |
| *AC022679.1--POLR2J3* | 7.7 |
| *AC022400.4--TIMM23B* | 7.7 |
| *AC022150.4--ZNF816* | 7.7 |
| *AC020661.4--MALAT1* | 7.7 |
| *AC018362.3--SSR2* | 7.7 |
| *AC017015.1--SET* | 7.7 |
| *AC015813.2--MALAT1* | 7.7 |
| *AC015813.1--TPI1* | 7.7 |
| *AC013410.1--HSP90AB1* | 7.7 |
| *AC012254.2--HNRNPA2B1* | 7.7 |
| *AC010197.2--AP002495.1* | 7.7 |
| *AC009951.4--FN1* | 7.7 |
| *AC009220.3--FCF1* | 7.7 |

| | |
|---|---|
| *AC009133.1--AC138028.4* | 7.7 |
| *AC009093.10--BANP* | 7.7 |
| *AC009086.2--SMG1* | 7.7 |
| *AC008581.2--IFITM2* | 7.7 |
| *AC007938.3--MALAT1* | 7.7 |
| *AC007608.4--MALAT1* | 7.7 |
| *AC007608.4--KDM5A* | 7.7 |
| *AC007192.1--NOP14* | 7.7 |
| *AC007192.1--COL1A2* | 7.7 |
| *AC006427.2--TAPT1-AS1* | 7.7 |
| *AC006064.6--PPP3R1* | 7.7 |
| *AC006064.6--MALAT1* | 7.7 |
| *AC006064.6--CAMSAP1* | 7.7 |
| *AC006001.3--UBB* | 7.7 |
| *AC005537.1--ATP8B1* | 7.7 |
| *AC005336.2--POLRMT* | 7.7 |
| *AC005154.2--GOLGA8R* | 7.7 |
| *AC005077.3--GTF2IRD1* | 7.7 |
| *AC004951.1--TIA1* | 7.7 |
| *AC004951.1--LINC00243* | 7.7 |
| *AC004951.1--DOCK5* | 7.7 |
| *AC004148.1--DUSP22* | 7.7 |
| *AC000120.1--MUC4* | 7.7 |
| *ABR--YWHAE* | 7.7 |
| *ABI1--NEK3* | 7.7 |
| *ABHD2--UBC* | 7.7 |
| *ABHD12--APOB* | 7.7 |
| *ABCD1--IGK@* | 7.7 |
| *ABCC6--BLOC1S2* | 7.7 |
| *ABCC4--ATP2A2* | 7.7 |
| *ABCC2--TMX2-CTNND1* | 7.7 |
| *ABCC2--SREBF2* | 7.7 |
| *ABCC2--MALAT1* | 7.7 |
| *ABCB6--HACL1* | 7.7 |
| *ABCB1--TRIM25* | 7.7 |
| *ABCA3--MALAT1* | 7.7 |
| *ABCA1--NUDT6* | 7.7 |
| *ABCA1--CNDP2* | 7.7 |
| *AASDH--ARPP19* | 7.7 |
| *AARS--GSN* | 7.7 |
| *AARS--AP000346.2* | 7.7 |
